# Comparative food-web analysis of bluefin tuna spawning habitats in the eastern Indian Ocean and Gulf of Mexico

**DOI:** 10.64898/2026.03.18.711569

**Authors:** Michael R. Stukel, Michael R. Landry, Moira Décima, Christian K. Fender, Sven A. Kranz, Raul Laiz-Carrion, Estrella Malca, Jose M. Quintanilla, Karen E. Selph, Rasmus Swalethorp, Natalia Yingling

**Affiliations:** Department of Earth, Ocean & Atmospheric Science, Florida State University, Tallahassee, FL 32306, USA; Scripps Institution of Oceanography, University of California at San Diego, La Jolla, CA 92093, USA; Department of Biosciences, Rice University, Houston, TX 77005, USA; Centro Oceanográfico de Málaga, Instituto Español de Oceanografía (IEO-CSIC), Fuengirola 29640, Spain; Cooperative Institute for Marine and Atmospheric Studies, University of Miami, Miami, FL 33146; Department of Oceanography, University of Hawai’i at Manoa, Honolulu, HI 96822, USA

**Keywords:** Indian Ocean, Gulf of Mexico, plankton ecology, marine food web, pelagic ecosystems, larval fish

## Abstract

Using linear inverse ecosystem modeling as a data assimilation tool, we compare spawning grounds of Atlantic and Southern Bluefin Tuna (ABT and SBT, respectively) based on results from field campaigns in the Gulf of Mexico (GoM) and eastern Indian Ocean off northwest Australia (Argo Basin). Both regions are warm, stratified, low-nutrient waters dominated by cyanobacteria (*Prochlorococcus*). Despite these similarities, the Argo Basin is more productive, with ∼1.5X higher net primary production and nearly 2X higher production of top trophic levels in the model (tuna larvae, planktivorous fish, and predatory gelatinous zooplankton). Higher primary production in the Argo Basin is mainly driven by higher N_2_ fixation and storm mixing of new nutrients in the upper and lower euphotic zone, respectively. Increased ecosystem efficiency (secondary production of top trophic levels / primary production) results from differences in plankton food web organization. In the GoM, protistan zooplankton are the direct consumers of nearly all phytoplankton production. In contrast, higher rates of herbivory by crustaceans feeding on nanophytoplankton combines with a higher impact of appendicularians on cyanobacteria to convert plankton production into larval tuna prey more efficiently in the Argo Basin. Despite similarities in the proportions of phytoplankton production mediated by cyanobacteria and other picoplankton in both systems, food web pathways to larval tuna and other planktivorous fish are substantially shorter in the Argo Basin. Our results highlight the impact of distinct zooplankton ecological niches on ecosystem efficiency and suggest a need for better inclusion of plankton food-web structure in models simulating climate impacts on fisheries production.

**HIGHLIGHTS:**

- Developed food web models of tuna spawning habitat (Indian Ocean & Gulf of Mexico)
- Spawning habitats in the Argo Basin and Gulf of Mexico (GoM) are both oligotrophic
- Argo Basin had higher net primary production in part as a result of nitrogen fixation
- Argo Basin had higher rates of direct herbivory by metazoan zooplankton
- This resulted in greater ecosystem efficiency in the Argo Basin.

## 1. Introduction

Bluefin tuna are marine top predators that feed over wide regions of the world oceans. However, they travel long distances to spawn in geographically restricted locations: deep-water regions of the Gulf of Mexico (GoM) and Mediterranean Sea for Atlantic Bluefin Tuna (ABT), and the Argo Abyssal Plain (Argo Basin) in the Indian Ocean north of Australia for Southern Bluefin Tuna (SBT) (Jenkins et al., 1991; Farley and Davis, 1998; Rooker et al., 2007; Teo et al., 2007). These regions are warm, nutrient-poor basins that are hypothesized to be favorable for larval recruitment due to some combination of low predation risk, enhanced larval growth rates, larval retention in favorable habitat, and/or eventual transport to suitable juvenile habitat (Bakun, 2013; Muhling *et al*., 2017; Díaz-Barroso *et al*., 2022; Malca *et al*., 2022; Shropshire *et al*., 2022; Reglero *et al*., 2025). While the exact characteristics that make these sites optimal spawning habitat are uncertain, the importance of these regions to species survival is clear and necessitates a better understanding of how these ecosystems function and might respond to changing climate (Landry et al., 2019).

In this study, we focus on spawning regions in the GoM and Argo Basin (Fig. 1), which were the sites of two recent ecosystem studies by comparable methods (Gerard et al., 2022; Landry et al., this issue-a). The deepwater GoM is an oligotrophic region with low euphotic-zone nutrient concentrations and a deep nitracline (Biggs, 1992; Knapp et al., 2021). Its circulation is heavily influenced by the spatially variable and very warm waters of the Loop Current. Phytoplankton and zooplankton biomasses are persistently low, and a strong deep chlorophyll maximum exists year-round (Green et al., 2014; Shropshire et al., 2020; Selph et al., 2021). Mesoscale eddies (including large eddies shed by the Loop Current) can persist for months to greater than a year as they propagate across the GoM and play important roles in modulating both vertical and horizontal nutrient flux and phytoplankton production (Biggs and Müller-Karger, 1994; Lee-Sánchez et al., 2022), and the larval fish distribution (Lindo-Atichati *et al*., 2012). Phytoplankton communities are dominated by the cyanobacterium *Prochlorococcus*, one of the smallest and most abundant photosynthetic organisms in the ocean (Selph et al., 2021), and phytoplankton growth is closely balanced by protistan grazing with relatively little production available for direct grazing by metazoan zooplankton (Landry et al., 2021b; Landry and Swalethorp, 2021). ABT larvae in the GoM are selective feeders with disproportionate impacts on poecilostomatoid copepods, cladocerans and occasionally appendicularians (Llopiz et al., 2015; Tilley et al., 2016; Shiroza et al., 2022). The distinct ecological roles of these different prey taxa suggest that ecosystem efficiency could be altered by the selection of different food chains linking the food web base to ABT larvae (Landry et al., 2019; Stukel *et al*., 2022) with direct implications on larval growth performance and survival (Quintanilla *et al*., 2024).

**Fig. 1.**
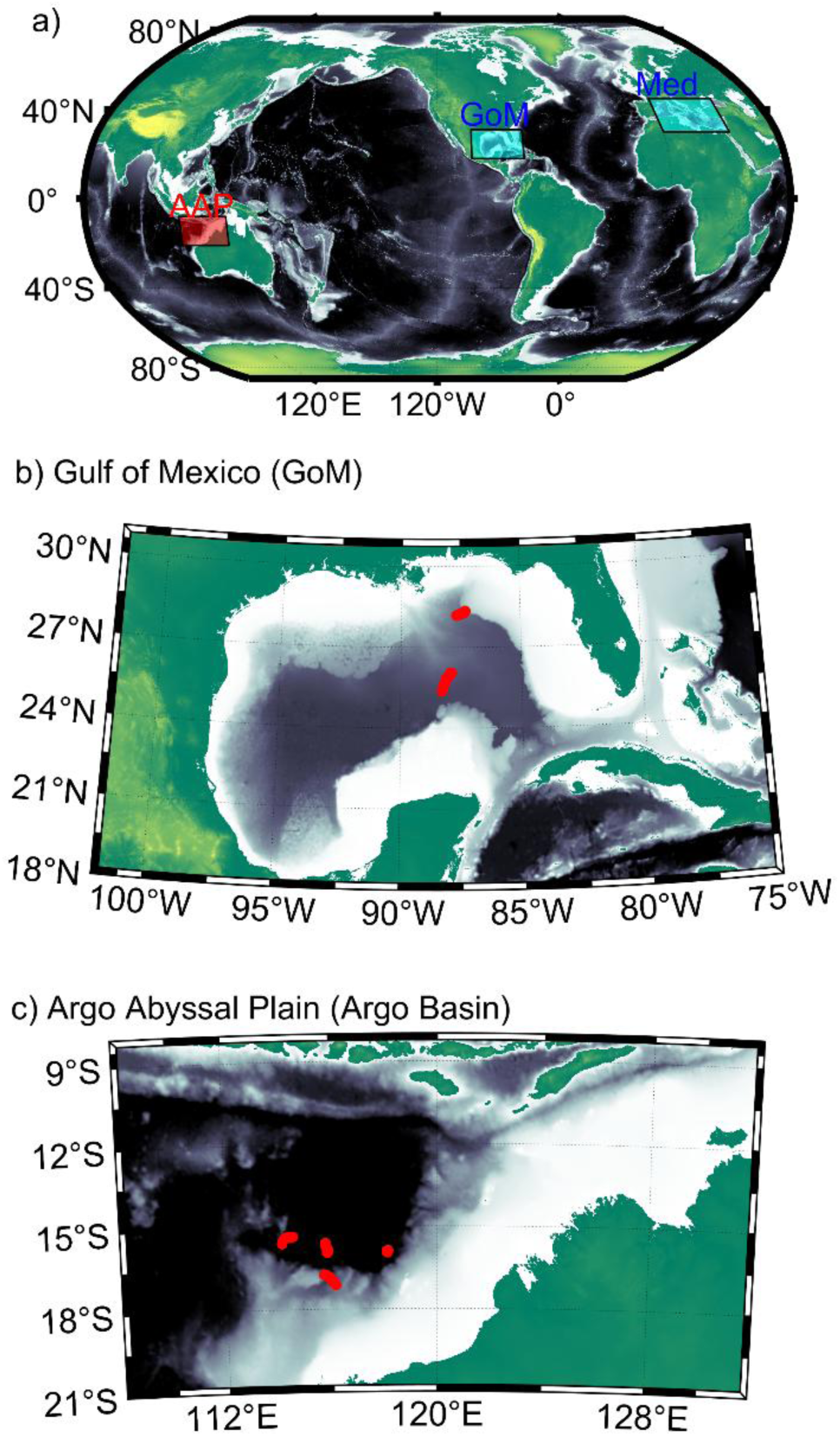
Global map (a) showing spawning sites for Atlantic bluefin tuna (blue) and Southern bluefin tuna (red) and insets focused on study regions in the Gulf of Mexico (b, GoM) and waters overlying the Argo Abyssal Plain (c, Argo Basin). Red symbols in (b) and (c) show Lagrangian experiment locations.

Relative to the GoM, the ecology of the Argo Basin region is substantially less studied. The dominant flow in the region is from the Indonesian Throughflow, the only major tropical connection between the Pacific and Indian Oceans, and into the South Equatorial and Leeuwin Currents. Summertime spawning season of SBT (January – February) coincides with the austral Northwest Monsoon season, when surface temperatures are very warm and the region is strongly stratified although it can be heavily impacted by transient disturbances including tropical cyclones (Quadfasel et al., 1996; Brink et al., 2007; Domingues et al., 2007; Condie and Andrewartha, 2008; Hood et al., 2017). The surface ocean is nutrient-depleted with commensurately low surface chlorophyll and net primary production (Llopiz and Hobday, 2015; Raes et al., 2022; Kehinde et al., 2023). The phytoplankton community is dominated by cyanobacteria and especially *Prochlorococcus*, while a diverse group of heterotrophic and mixotrophic eukaryotes may play an important role in nutrient regeneration (Landry *et al*., 2021a; Raes et al., 2022; Selph *et al*., this issue-a). As with ABT, SBT larvae show strong selectivity for distinct zooplankton taxa including poecilostomatoid copepods and appendicularians (Young and Davis, 1990; Swalethorp *et al*., this issue) together with narrow isotopic niches (Laiz-Carrión *et al*., this issue).

In this study, we synthesize results from paired Lagrangian experiments conducted in the GoM and Argo Basin (Gerard et al., 2022; Landry et al., this issue-a) using a linear inverse ecosystem modeling (LIEM) approach (Vézina and Platt, 1988; van Oevelen *e*t al., 2010). LIEM is a data synthesis tool that assimilates food web rate and standing stock measurements with a priori knowledge of food-web structure and organismal physiology to constraint flows of mass or energy between different living and non-living compartments. Our focus is on elucidating the pathways that connect the base of the euphotic zone (nutrient supply and primary producers) to larval tuna, with daily growth implications (Borrego-Santos *et al*., this issue). We find substantially greater secondary production of secondary consumers in the Argo Basin relative to the GoM and further show that this results from both increased phytoplankton production and a more efficient food web. Increased ecosystem efficiency is driven by differences in food-web structure that allow metazoan zooplankton access to a higher proportion of phytoplankton production through herbivory in the Argo Basin, thus enabling shorter food chains to larval tuna and other planktivorous fish.

## 2. Methods

### 2.1. Field campaigns and in situ data

The data synthesized in this study comes from two large field programs. The BLOOFINZ-GoM (Bluefin Larvae in Oligotrophic Ocean Foodwebs, Investigations of Nutrients to Zooplankton – Gulf of Mexico) cruises were conducted in deep-water regions of the Gulf of Mexico (spawning habitat for ABT) in May 2017 and May 2018 (Gerard et al., 2022). The BLOOFINZ-IO (Indian Ocean) cruise was conducted mostly in February 2022 in the Argo Basin off of northwest Australia (Landry et al., this issue-a). Both programs used a Lagrangian sampling scheme in which water parcels found to have bluefin tuna larvae were tagged with floating arrays that included 3×1-m holey sock drogues centered at 15 m (Landry et al., 2009; Fernández et al., this issue), allowing the water parcel and its plankton community to be sampled repeatedly over periods of 3 to 5 days. Five Lagrangian experiments were conducted as part of BLOOFINZ-GoM, of which only two were in water parcels with ABT; hence, only those two will be included in this study (Fig. 1b). Four Lagrangian experiments were conducted, all with SBT larvae, as part of BLOOFINZ-IO (Fig. 1c). During each of these Lagrangian experiments a suite of measurements was made to determine ecosystem standing stocks and rates from nutrients and biogeochemistry through phytoplankton and zooplankton to larval tuna (Supp. Table S1).

Nutrient concentrations were measured at 6 depths spanning the euphotic zone and additional depths through the mesopelagic zone, and the δ^15^N of nitrate was measured by the denitrifier method (Knapp et al., 2021; Kranz *et al*., this issue). Particulate organic matter (POM) concentrations (organic carbon and nitrogen) and isotopes were measured at similar depths (Kehinde et al., 2023). Drifting sediment trap arrays were utilized to collect sinking particles beneath the mixed layer and at the base of the euphotic zone (Stukel *et al*., 2021; Stukel *et al*., this issue). Sinking particles were assayed to measure organic carbon, nitrogen, δ^15^N, chlorophyll *a* (used to estimate phytoplankton flux), and phaeopigments (used to estimate zooplankton fecal pellet flux). Supply rates of new nitrogen via upwelling and lateral transport of organic matter were constrained by Thorpe scale analysis and remote sensing (Kelly *et al*., 2021; Kehinde et al., 2023).

Daily CTD casts were conducted at 02:00 to collect samples from 6 depths spanning the euphotic zone for a suite of microbial (bacteria and protists) biomass and rate measurements. Picoplankton abundances (*Prochlorococcus*, *Synechococcus*, picoeukaryotic phytoplankton, and heterotrophic bacteria) were determined by flow cytometry (Selph et al., 2021; Selph et al., this issue-a). Flow cytometry counts were converted to biomass assuming cellular carbon contents of 32, 101, 320, and 5 fg C cell^−1^ for *Prochlorococcus*, *Synechococcus*, picoeukaryotic phytoplankton, and heterotrophic bacteria, respectively. Nano- and microplankton phototroph biomasses (flagellates and diatoms) were determined by epifluorescence microscopy (Selph et al., 2021; Yingling *et al*., this issue). *Trichodesmium* biomass was estimated from microscopic analysis of trichomes collected from 6-L samples filtered onto 8-µm pore-size filters directly from CTD bottles (Selph et al., 2021). Because *Trichodesmium* and diatom biomass were determined to be below detection limits on the BLOOFINZ-IO cruise (despite identical methods as those used in BLOOFINZ-GoM), we assumed that their maximum biomass and rates were equal to half of that measured in BLOOFINZ-GoM. However, this assumption has little impact on model results because neither phytoplankton group was a major source of production in either region. Net primary production (NPP) was measured by incorporation of H^13^CO_3_^−^ or H^14^CO_3_^−^ in incubations conducted *in situ* on the drifting array (Yingling *et al*., 2022; Kranz et al., this issue). Nitrate and ammonium uptake rates were measured by incorporation of ^15^N-labeled nutrients (6 depths incubated in situ for nitrate, 3 depths incubated deckboard for ammonium, Yingling et al., 2022; Yingling et al., this issue). Nitrogen fixation was measured by incorporation of ^15^N-labeled dinitrogen gas (Kranz et al., this issue). Taxon-specific phytoplankton growth rates and mortality due to protistan grazing were measured by the protistan grazing dilution method (Landry et al., 2021b; Landry *et al*., this issue-b). Bacterial carbon production and protistan grazing rates on bacteria were similarly measured in the same dilution experiments (Landry *et al*., 2023; Landry et al., this issue-b).

Mesozooplankton were collected with ring nets and bongo nets towed through either the upper euphotic zone (0-30 m, larval bluefin habitat) or the entire euphotic zone (Landry and Swalethorp, 2021; Shiroza et al., 2022; Décima *et al*., this issue; Swalethorp et al., this issue). Whole euphotic zone samples were size-fractionated (five size-classes) and assayed for biomass, isotopes, and grazing rates by the gut pigment method (Landry and Swalethorp, 2021; Décima et al., this issue). Upper euphotic zone samples were sorted to coarse taxonomic resolution to determine average sizes, biomasses, and isotopic compositions of appendicularians, cladocerans, chaetognaths, calanoid copepods, and poecilostomatoid copepods (Shiroza et al., 2022; Swalethorp et al., this issue).

Bluefin larvae were collected with bongo nets towed through the upper 25-m of the water column. The larvae were sorted and sized, and subsets were assayed for stage (pre-flexion, flexion, or post-flexion), isotopic composition, age (via otolith measurements) and gut contents (Malca et al., 2022; Shiroza et al., 2022; Borrego-Santos et al., this issue; Laiz-Carrión et al., this issue; Quintanilla, this issue; Swalethorp et al., this issue). Potential prey items were identified to the following groups: microzooplankton (ciliates), appendicularians, cladocerans, chaetognaths, calanoid copepods, and poecilostomatoid copepods (Shiroza et al., 2022; Swalethorp et al., this issue).

For both BLOOFINZ-GoM and BLOOFINZ-IO, mean values from each Lagrangian experiment were averaged to determine project means, which were then used as inputs to the LIEM (Supp. Table S1).

### 2.2. Food-web structure

The food-web structure was designed following Stukel et al. (2022) to investigate energy and nutrient pathways from the base of the food chain to larval tuna (Fig. 2, Supp. Table S2). The model is nitrogen-based, starting with three inorganic N classes (nitrate, ammonium, and N_2_ gas), and with three pools of non-living organic matter (dissolved organic matter (DOM), small non-sinking detritus, and large detritus). There are four phytoplankton: *Trichodesmium*, picophytoplankton (assumed to be potentially diazotrophic), diatoms, and flagellates (assumed to be potentially mixotrophic, see Selph *et al*., this issue-b). Heterotrophic bacteria consume DOM, while mixotrophic flagellates, heterotrophic nanoflagellates, and microzooplankton comprise three groups of protists which feed on phytoplankton and smaller heterotrophs. Six suspension-feeding mesozooplankton are included: appendicularians (assumed capable of feeding on cyanobacteria and heterotrophic bacteria), vertically-migrating calanoid copepods, non-vertically-migrating calanoid copepods, cladocerans, other non-vertically-migrating herbivorous suspension feeders, and other vertically-migrating herbivorous suspension feeders. There are two small predatory mesozooplankton (chaetognaths and poecilostomatoid copepods) along with four higher trophic levels that serve as closure terms in the model (preflexion bluefin larvae, postflexion bluefin larvae, other planktivorous fish, and predatory gelatinous zooplankton). Bluefin larvae are assumed to feed on microzooplankton, appendicularians, cladocerans, non-vertically-migrating calanoid copepods, and poecilostomatoid copepods. The model includes two layers: upper euphotic zone (0-50 m for GoM, 0-40 m for Argo Basin) and deep euphotic zone (50-100 m for GoM; 50-80 m for Argo Basin). All model compartments are identical between the two layers, except that bluefin larvae only exist in the upper euphotic zone. The two layers are connected through upward nitrate flux, downward sinking particle flux and the movement of vertical migrators, which inhabit both layers during the night, but rest beneath the euphotic zone (i.e., outside the model) during the day. Nitrate, diazotrophy, and lateral advection of POM and DOM are nitrogen inputs to the model that must balance the following closure terms: secondary production of higher trophic levels, sinking of large detritus, sinking of diatoms, sinking of mixotrophic flagellates, and excretion from vertical migratory taxa beneath the euphotic zone. We assume Redfield stoichiometry for all model flows, which allows us to relate respiration to ammonium excretion. All model flows are shown in Supp. Table S2.

**Fig. 2.**
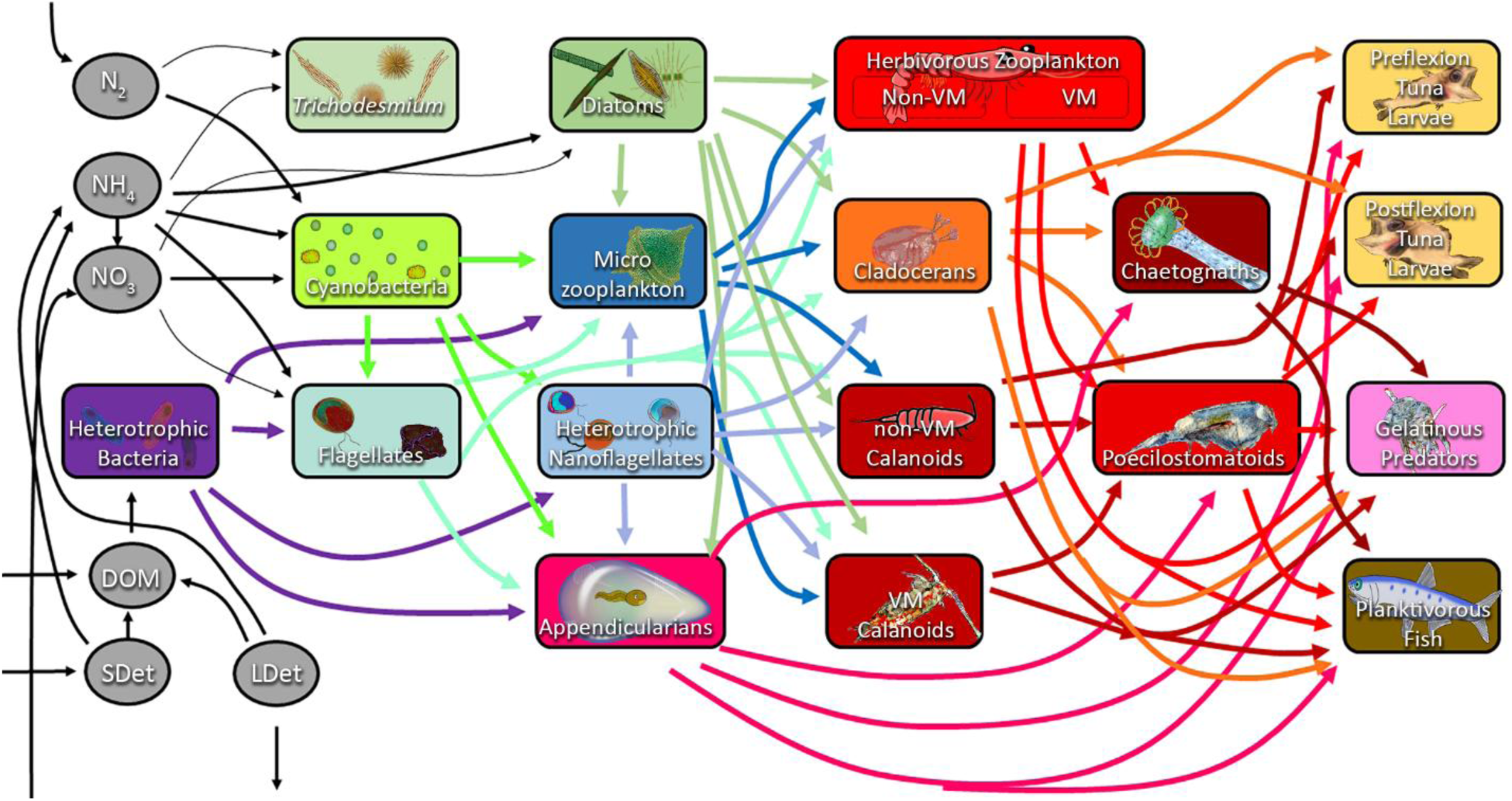
Food web structure. The model has a two-layer structure (∼mixed layer and deep euphotic zone) with this diagram depicting the mixed layer portion of the model (deep euphotic zone is identical except for the absence of tuna larvae). Phytoplankton (cyanobacteria, potentially mixotrophic flagellates, diatoms, and *Trichodesmium*) rely on nitrate (NO_3_), ammonium (NH_4_) or dinitrogen fixation. Primary grazers include protists (heterotrophic nanoflagellates, microzooplankton, mixotrophic flagellates, all of which can feed on phytoplankton and heterotrophic bacteria) and suspension-feeding mesozooplankton (appendicularians, cladocerans, vertically migrating calanoid copepods, non-vertically migrating calanoid copepods, and other vertically migrating and non-vertically migrating herbivores, all of which can consume protists but only appendicularians can consume bacteria-sized prey). Two carnivorous zooplankton (chaetognaths and poecilostomatoids) feed on other mesozooplankton, while pre-flexion bluefin larvae, post-flexion bluefin larvae, other planktivorous fish, and gelatinous predators feed on multiple zooplankton groups (tuna larvae also feed on microzooplankton, but cannot eat vertical migrators). Non-living model compartments also include dissolved organic matter (DOM), small non-sinking detritus (SDet) and large sinking detritus (LDET). All major food web flows between organisms are shown. However, for visual simplicity, we omit production of NH_4_^+^, DOM, and detritus by all living groups as well as consumption of detritus by protistan zooplankton and suspension-feeding metazoans. For all model flows, see Supp. Table 2.

### 2.3. Linear inverse ecosystem model (LIEM) solutions

The linear inverse problem involves 302 unknown ecosystem flows (e.g., nitrogen flux from one compartment to another). These 302 variables are constrained with 44 equations defining mass-balance constraints for each of the model compartments (Fig. 2). We also include 42 approximate equality constraints that encapsulate measured ecosystem rate processes (and their associated uncertainty) including nitrate and ammonium uptake, nitrogen fixation, net primary production, taxon-specific phytoplankton production rates, taxon-specific phytoplankton mortality due to protistan grazing, heterotrophic bacteria production and losses to grazing, taxon-specific ingestion rates of larval tuna, and sinking particle flux (Supp. Table S1). An additional 44 approximate equality constraints are based on ^15^N isotope mass balance. These isotopic mass balance constraints are treated as approximate equalities, because isotopic ratios are not known for all model compartments. Finally, model solutions are bounded by a series of 556 greater than or less than constraints which are taken directly from Stukel et al. (2022) and include simple constraints (e.g., all model flows must be positive), as well as a series of constraints built on basic knowledge of plankton ecosystems (e.g., zooplankton gross growth efficiencies can vary from 10 to 40% and their minimum metabolic requirements are set by published allometric relationships; phytoplankton exudation of DOM is capped at 55% of NPP and maximum growth rates are set to two doublings per day). Although the LIEM remains substantially under constrained, the inequality constraints put bounds on the solution space.

The solution space can be thought of as a 258-dimensional polytope (302 variables minus 44 exact equalities = 258) that includes all combinations of ecosystem flows that satisfy the mass balance and inequality constraints. To sample this solution space and find unique solutions that minimize the mismatch between the 86 approximate equalities, we used the Markov Chain Monte Carlo (MCMC) approach (Kones *et al*., 2009; Van den Meersche *et al*., 2009; van Oevelen et al., 2010). The MCMC approach finds an initial solution in the polytope that satisfies the inequality and equality constraints. It then engages in a directed random walk in which new solutions are computed as slight nudges from the prior solutions and the new and prior solutions are compared with respect to their fit to the approximate equality constraints. If the new solution has a lower misfit to the observations, it is automatically accepted. If it has a higher misfit to the observations, its potential acceptance is based on a random number draw and the relative misfit between the two solutions. If it is rejected, the process is repeated from the prior solution. This process generates a set of millions of solutions that (after a “burn-in” period has been removed) can all be considered equally likely solutions to the LIEM. Mean solutions for each ecosystem flow are calculated as the arithmetic mean of across all solutions, while uncertainty in each flow can then be assessed via standard deviations or 95% confidence intervals. Importantly, the uncertainties encompass uncertainties due to the underconstrained nature of the LIEM as well as uncertainties in observational constraints. The specific implementation of the MCMC approach follows the MCMC+^15^N approach which introduced a second set of unknowns representing the ^15:14^N isotopic ratios of model compartments (Stukel *et al*., 2018a; Stukel *et al*., 2018b), thus allowing nonlinear equations (isotopic mass balance) to further constrain the solutions. The MCMC solution was run for >1.5×10^8^ iterations with the first 20% discarded as a “burn-in” period and the remainder thinned by retaining only every 10,000^th^ solution set (for computational efficiency).

### 2.4. Food-web indices

We computed trophic levels (TL) for all consumers as one plus the ingestion-weighted mean TL of prey (*TL*_*consumer*_ = ∑(*TL*_*prey*,*i*_ × *F*_*prey*,*i*→*consumer*_)⁄∑ *F*_*prey*,*i*→*consumer*_, where *TL*_*prey*,*i*_ is the trophic level of prey i and *F*_*prey*,*i*→*consumer*_ is the rate of feeding of the consumer on prey i). All phytoplankton were assumed TL=1, except mixotrophic flagellates, which had *TL* = (1 − *p*_*phag*_) + *p*_*phag*_(1 + *TL*_*prey*_), where *p_phag_* is the proportion of their nitrogen derived from phagotrophy (rather than dissolved nutrient uptake). Heterotrophic bacteria were assumed to have a TL equal to one plus the TL of the organism producing the organic matter they utilized.

We defined three major pathways of energy and nutrient flow from the base of the food web (following Stukel et al., 2012): i) The herbivorous food chain is the sum of direct nitrogen fluxes from phytoplankton to metazoan zooplankton. ii) The multivorous food chain is the sum of nitrogen fluxes that reach metazoan zooplankton after passing through protistan grazers. iii) The microbial loop is the sum of bacterial respiration and the fraction of protistan respiration supported by bacterial production.

We used indirect food web flow analysis to trace nitrogen flows through the food web. The normalized amount of nitrogen (direct and indirect) that any organism derives from any other organism can be computed following Hannon (1973) as (*I* − *G*)^−1^, where *I* is the identity matrix and *G* is the normalized production matrix (i.e., a matrix giving the percentage of an organism’s nitrogen requirement directly derived from any other organism).

## 3. Results

### 3.1. Physical and biogeochemical environment

Conditions encountered during the BLOOFINZ-GoM and BLOOFINZ-IO cruises were broadly similar. The GoM and Argo Basin were both warm (surface temperatures of 24.5 to 26.5°C in the GoM and 28 to 30.5°C in the Argo Basin). Mixed layers depths ranged from <15 to ∼30 m in both regions. Nutrient (nitrate and ammonium) concentrations were negligible in the mixed layer and the depth of the nitracline (defined as the depth at which nitrate concentration first exceeded 1 µmol L^−1^) was typically near the depth of the deep chlorophyll maximum (DCM). Nitracline and euphotic-zone depths were, however, generally deeper in the GoM (DCM depths ranged from 84 to 127 m) than in the Argo Basin (DCM depths ranged from 67 to 75 m). This matched a general pattern of greater oligotrophy in the GoM relative to the Argo Basin, with NPP averaging 27 mmol C m^−2^ d^−1^ in the GoM compared to 40 mmol C m^−2^ d^−1^. Surface chlorophyll *a* concentrations, however, were similar between regions, ranging from 0.05 to 0.13 in the GoM and 0.07 to 0.09 mg chlorophyll *a* m^−3^ in the Argo Basin. These characteristics are all consistent with expectations for tropical/subtropical oligotrophic basins.

### 3.2. Model-observation agreement

The LIEM exhibited close agreement with observations (Fig. 3). The square root mean squared error, which is basically the average number of standard errors (of the observations) that the LIEM estimates were from the observation, averaged 1.17 for GoM if we considered only the field rate measurements and 1.92 if we include the δ^15^N mass balance equations. The equivalent values for Argo Basin were 1.60 for rate measurements and 0.89 for all approximate equalities. Across both cruises and all observations, by far the largest mismatches (based on square root mean squared error) was for NPP in the Argo Basin. The square root mean squared error was 5.2 for shallow NPP and 4.1 for deep NPP. These large normalized error terms primarily reflected the low uncertainties in observations for these measurements, rather than a massive model-data mismatch. For shallow NPP, the LIEM estimate was 3.9 ± 0.07 mmol N m^−2^ d^−1^ while the observed NPP was 3.5 ± 0.09 mmol N m^−2^ d^−1^. For deep NPP, the LIEM estimate was 2.6 ± 0.09 compared to observed values of 2.3 ± 0.09. These specific model-data mismatches are not particularly surprising. The MCMC method often slightly overestimates NPP. This happens because the multi-dimensional space of possible solutions — defined by the inequality constraints — tends to contain more scenarios where NPP is higher than the observed value, compared to scenarios where it is lower. In other words, the “allowed” space above the observed NPP is larger than the space below it, so the sampling skews high (Stukel et al., 2012).

**Fig. 3.**
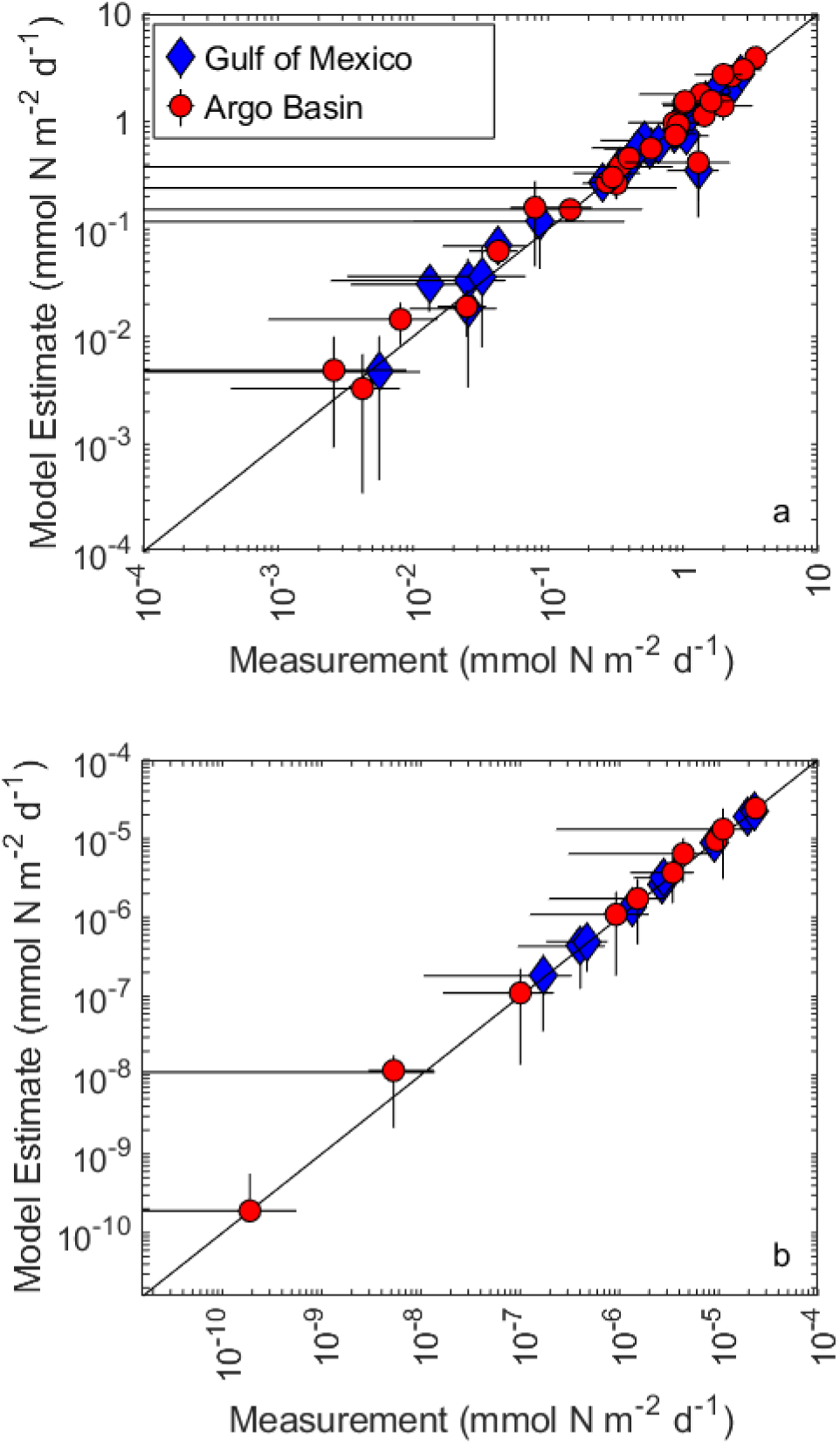
Model-observation comparisons. Comparison of field-measured rates (x-axis) with model estimates (y-axis). Blue diamonds are Gulf of Mexico. Red circles are Argo Basin. Figure is split into two panels because of the large difference in magnitude of flows within the lower food web (a) and flows to larval bluefin tuna (b).

While the LIEM was generally in good agreement with the observations, a few model-data mismatches are notable. First, the LIEM consistently overestimated mesozooplankton grazing rates relative to estimates from the gut pigment measurements. For the GoM, the LIEM overestimated grazing by 24%, while for Argo Basin it overestimated grazing by 44%. These grazing overestimates may lead to slight overestimates of total ecosystem efficiency by passing more energy through the “herbivorous” or “classical” food chain rather than the microbial loop or multivorous food webs. Nevertheless, given that the LIEM also overestimated NPP, this is not a major reapportionment of food-web energy. The other notable model-data mismatch was in the relative balance of production by cyanobacteria and eukaryotes for Argo Basin. The LIEM suggested that the ratio of cyanobacteria NPP to eukaryotic phytoplankton NPP was 2.3 in the upper ocean while the measurements suggested it should be 1.3. This reflected a substantial overestimate of NPP by cyanobacteria and an underestimate of NPP by phototrophic flagellates. This difference did not occur for GoM, where the LIEM overestimated NPP by both groups. The underestimate of flagellate NPP for Argo Basin did not, however, substantially alter the predicted balance of energy flowing from these groups, because the LIEM suggested substantial phagotrophy within the phototrophic (mixotrophic) flagellate group (which is actually supported by independent estimates of mixotrophy by this group, Selph et al., this issue-b). We also note that the observed NPP estimates for flagellates are based on the dilution technique which actually measures growth rate, rather than NPP. Considering the additional growth supported by phagotrophy that is not measured as ^14^C uptake, the LIEM and observations are in much closer agreement.

### 3.3. Ecosystem structure

There are many broad similarities between the GoM and Argo Basin bluefin tuna spawning regions. Recycled nutrients (ammonium) were the dominant source of nutrients for phytoplankton; NPP was moderately higher in the upper euphotic zone than the lower euphotic zone; cyanobacteria were the dominant primary producers with eukaryotic flagellates also playing a substantial role; diatoms and *Trichodesmium* were only minor contributors; and protists were the dominant grazers of phytoplankton. However, we also note some distinct differences between the regions. In the GoM, upwelled nitrate and nitrogen fixation were negligible sources of new nitrogen to the ecosystem. Instead, the ecosystem was supplied primarily by the lateral advection of nutrients and organic matter from nearby productive shelf regions. In the Argo Basin, the ecosystem was supported more evenly by the three processes. In the upper euphotic zone, nitrogen fixation was the dominant source of new nitrogen (introducing 0.56 mmol N m^−2^ d^−1^) to the ecosystem. Importantly, this nitrogen fixation was predominantly mediated by unicellular cyanobacteria, rather than *Trichodesmium,* and hence contributed directly to the microbial food web. In the deep euphotic zone, lateral transport was estimated to be the dominant source of new nitrogen, although nitrogen fixation was still quantitatively important.

Focusing on the upper euphotic zone, which is especially important as the layer in which bluefin larvae are found, a major difference became apparent in the partitioning of energy between the microbial loop, multivorous food web, and herbivorous food chains (Fig. 4). In the GoM, the microbial loop and multivorous food web dominated the upper euphotic zone, processing 1.8 ± 0.3 and 1.4 ± 0.1 mmol N m^−2^ d^−1^, respectively. The herbivorous food chain was much less important, with 0.2 ± 0.06 mmol N m^−2^ d^−1^ passing directly from phytoplankton to metazoan zooplankton. In the Argo Basin, the herbivorous food chain was substantially more important (1.3 ± 0.03 mmol N m^−2^ d^−1^) and comparable in magnitude to the microbial loop (2.0 ± 0.5 mmol N m^−2^ d^−1^) and multivorous food web (1.4 ± 0.08 mmol N m^−2^ d^−1^). This reflects a substantial alteration to the food web structure and allows more efficient energy transfer to higher trophic levels. This LIEM result was also supported by the observations, which showed >2-fold higher mesozooplankton grazing rates in the Argo Basin, relative to the GoM (Supp. Table S1).

**Fig. 4.**
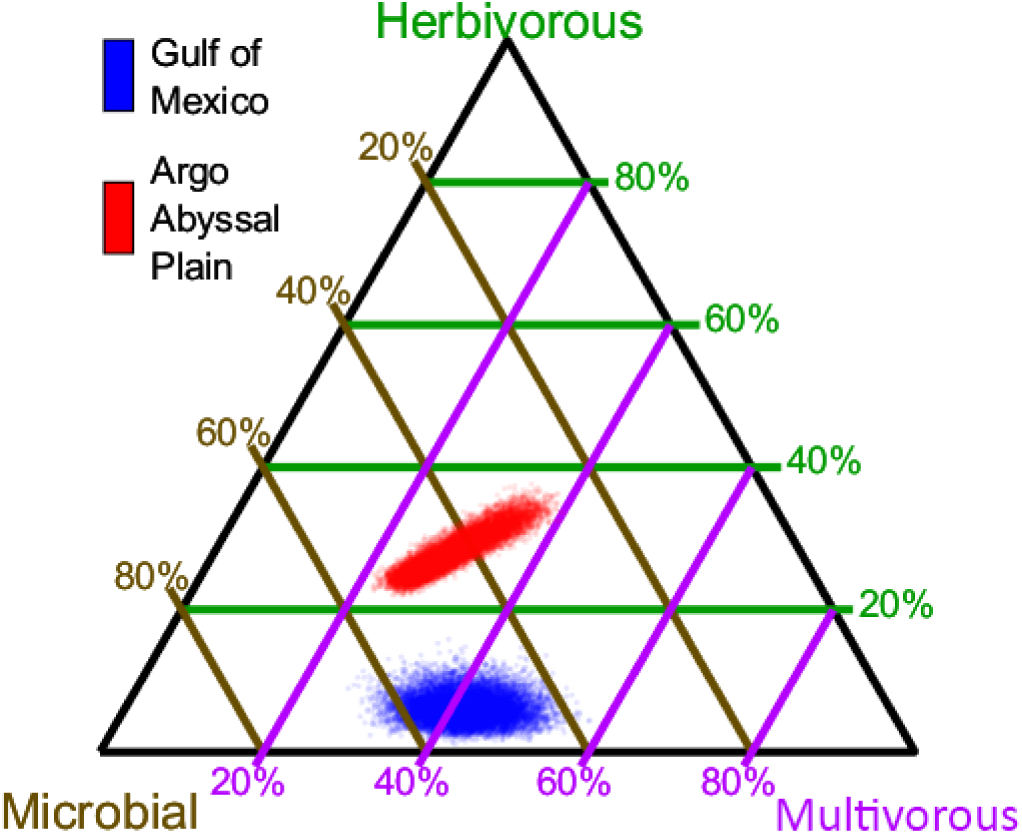
Ternary diagram showing relative importance of different food-web pathways in the upper euphotic zone of the Gulf of Mexico and Argo Basin (individual points are plotted for each MCMC solution vector to show uncertainty range).

Altered ecosystem structure was also evident in LIEM trophic level estimates (Fig. 5). Focusing on higher trophic levels, bluefin tuna larvae and poecilostomatoid copepods (a carnivorous group of copepods) had meaningfully lower trophic positions in the Argo Basin than in the GoM. For instance, the trophic position of preflexion larval tuna was 4.2 ± 0.1 in the GoM and 3.7 ± 0.1 in the Argo Basin, while postflexion larval tuna trophic positions were 4.3 ± 0.1 in the GoM and 3.9 ± 0.1 in the Argo Basin. This difference in trophic position of higher trophic levels in the model was not generally reflected in the trophic positions of suspension-feeding zooplankton. Indeed, calanoid copepods had similar trophic positions in each region, while the LIEM actually predicted higher trophic positions for cladocerans in the Argo Basin than in the GoM. Rather, the differences were primarily driven specifically by dynamics associated with appendicularians. Appendicularians are unique taxa with fine-mesh mucous feeding filters that are able to efficiently capture bacterial size prey. They hence are the only metazoan taxon in the LIEM that feed directly on cyanobacteria. Appendicularians were not abundant in the GoM. However, in the Argo Basin they created an efficient food web pathway linking carnivorous zooplankton and planktivorous fish to cyanobacteria production.

**Fig. 5.**
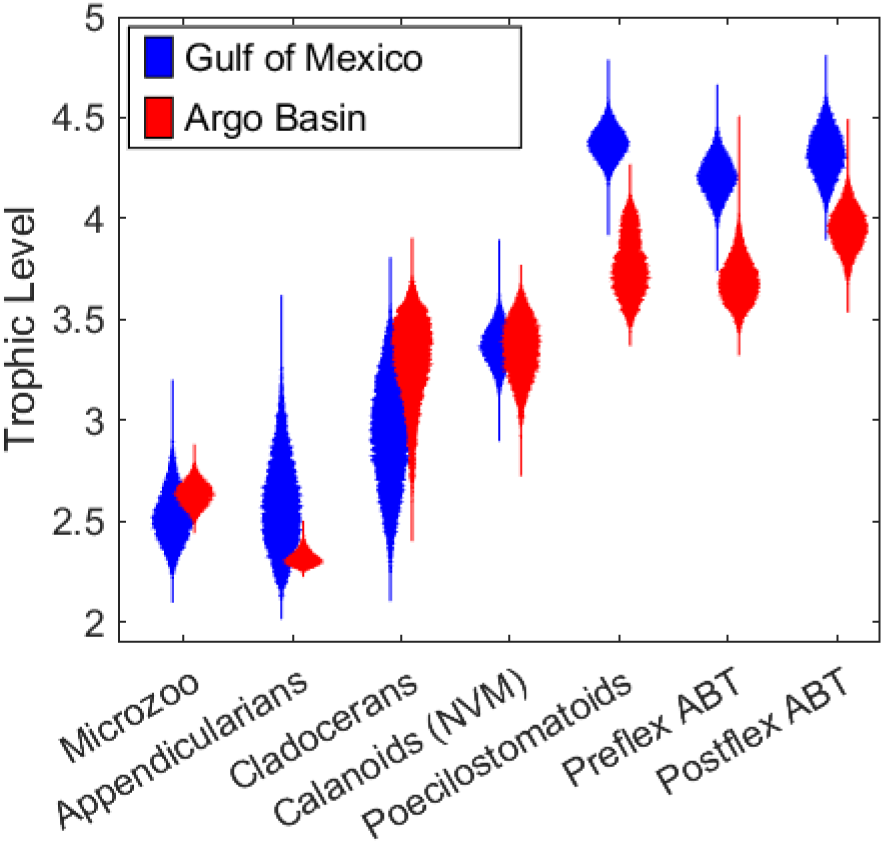
Violin plot showing trophic levels of key taxa.

### 3.4. Larval bluefin tuna

Bluefin tuna larvae exhibited strikingly different diets in the GoM and Argo Basin (Fig. 6), selectively choosing podonid cladocerans as prey in the GoM (while also consuming substantial amounts of abundant calanoid copepods) and selectively consuming appendicularians in the Argo Basin. In the GoM, preflexion larvae derived the majority of their nutrition from calanoid copepods (74%), with appendicularians comprising only 10% of their diet. In the Argo Basin, this shifted to appendicularians providing most of their diet (68%), with calanoid copepods only contributing 19%. These differences were also apparent for flexion and postflexion larvae, which, in the GoM derived most of their nutrition from a combination of cladocerans (35%) and calanoid copepods (41%), with negligible (5%) contributions from appendicularians. In the Argo Basin, appendicularians were the dominant dietary component of flexion and postflexion larvae (48%) with poecilostomatoid and calanoid copepods providing most of the remainder (25% and 19%, respectively). As noted above, the significant contribution of appendicularians to the diets played an important role in lowering the trophic position of SBT relative to ABT in the GoM (Fig. 5). Given typical zooplankton gross growth efficiencies of 30%, this 0.4-0.5 trophic position difference in food chain length equates to a ∼40% decrease in food web transfer efficiency in the GoM, relative to the Argo Basin, mediated in large part by the differences in level of appendicularian contributions to their respective diets.

**Fig. 6.**
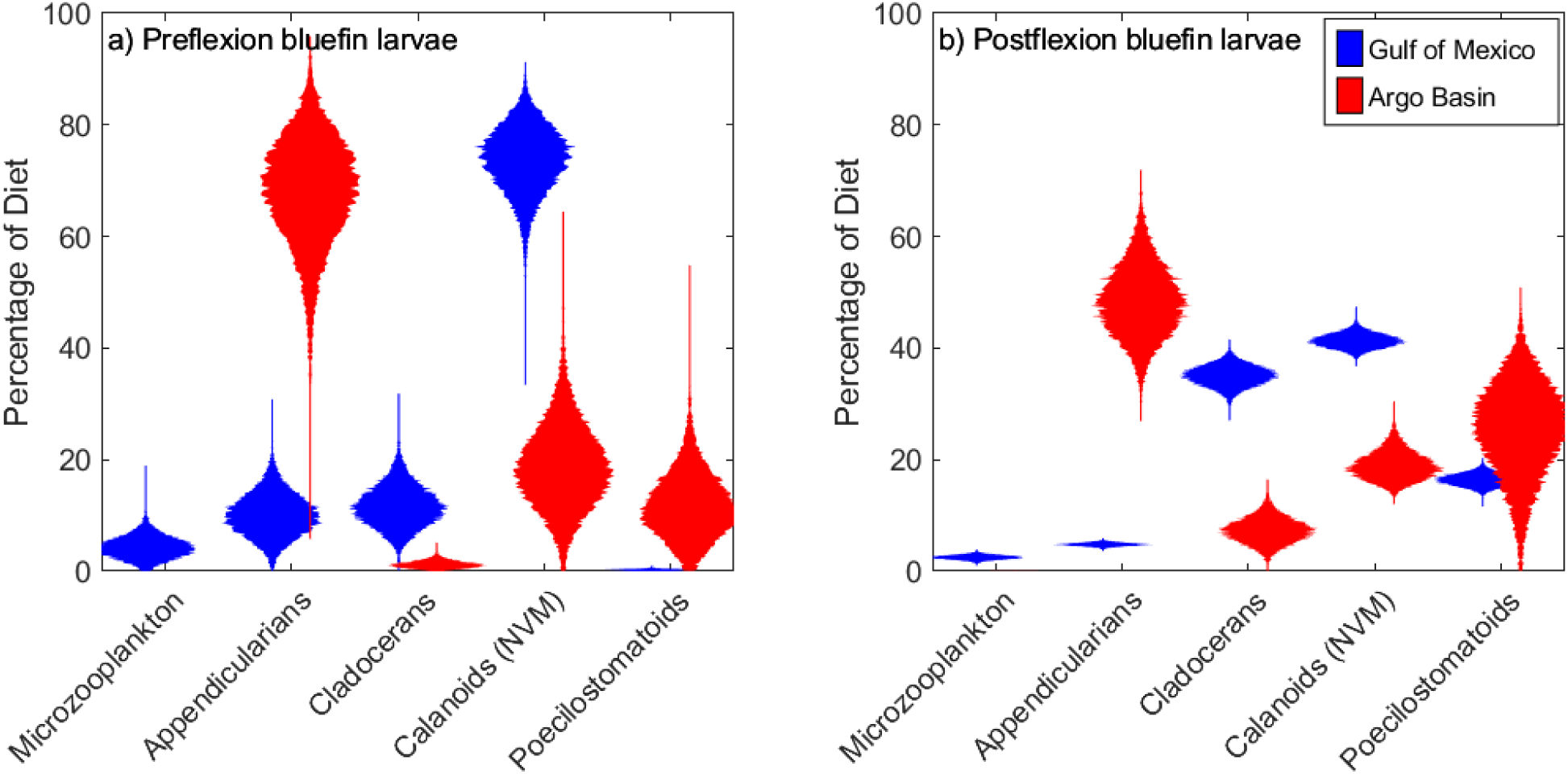
Violin plots showing the percentage of different prey in tuna diets. When prey groups are not visible it indicates that they were not found in larvae guts.

To more directly quantify the importance of this dietary difference, we also used indirect food web flow analysis (subsection 2.4) to track nitrogen through the ecosystem (Fig. 7). These results show a greater reliance of tuna larvae on nitrogen fixation in the Argo Basin than in the GoM. In the Argo Basin, nearly 10% of the nitrogen consumed by tuna larvae stems from the nitrogen fixation of cyanobacteria. The importance of overall cyanobacteria production to tuna larvae diets was relatively similar in the two regions (50 ± 6% in GoM; 62 ± 2% in Argo Basin). However, the ways in which the food web structured the flows from cyanobacteria to larvae was distinctly different. In the Argo Basin, 44 ± 5% of tuna larvae diets was derived from pathways that flowed from cyanobacteria through appendicularians to tuna larvae (note that this was primarily through the direct pathway cyanobacteria ➜ appendicularians ➜ tuna, but also included less direct pathways such as cyanobacteria ➜ protists ➜ appendicularians ➜ tuna). In the GoM, the pathway through appendicularians comprised only 3 ± 1% of larvae diets. Instead, cyanobacterial support of ABT larvae was primarily through less direct pathways that included other suspension-feeding mesozooplankton feeding on protistan grazers. For instance, if we focus on the pathways that support consumption of calanoid copepods by tuna larvae, in the Argo Basin, these were primarily supported by the production of phototrophic flagellates. However, in the GoM, foodweb pathways from cyanobacteria through calanoid copepods to larvae supported a higher portion of the larvae diet (24 ± 3%) than pathways from flagellates through calanoid copepods to larvae (15 ± 2%).

**Fig. 7.**
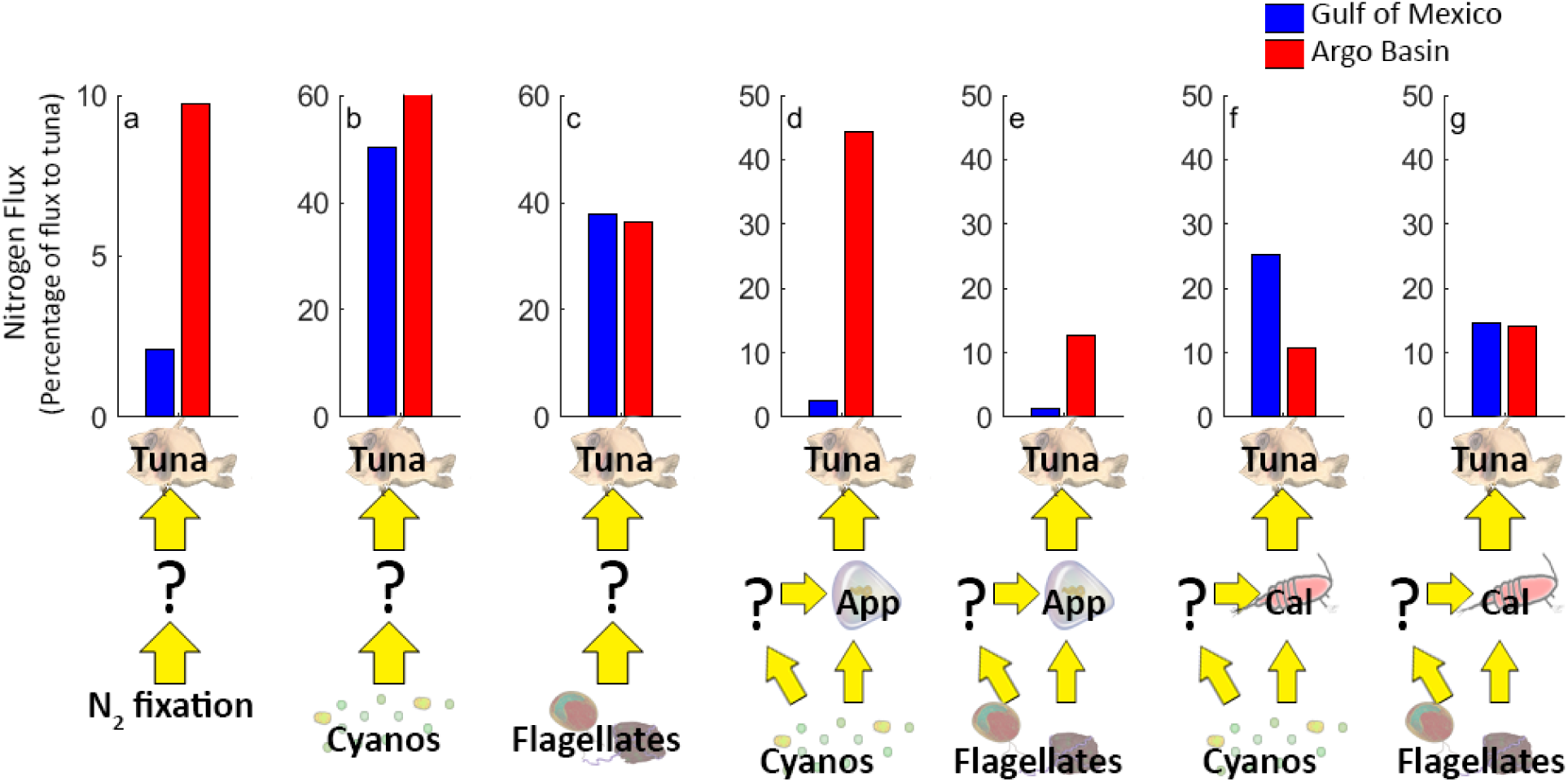
Foodweb pathways to larval tuna. All panels show the percentage of tuna ingestion supported by specific foodweb pathways. a) All pathways from nitrogen fixation to larval tuna. b) All pathways from cyanobacteria to tuna. c) All pathways from mixotrophic flagellates to tuna. d) All pathways from cyanobacteria to appendicularians to tuna. e) All pathways from flagellates to appendicularians to tuna. f) All pathways from cyanobacteria to non-vertically migrating calanoid copepods to tuna. g) All pathways from flagellates to non-vertically migrating calanoid copepods to tuna. Note that d – g include direct food chains from phytoplankton to zooplankton to tuna, as well as many longer food chains in which the phytoplankton are processed by other taxa prior to consumption by the zooplankton.

## 4. Discussion

### 4.1. Nutrient supply

Climate change is expected to lead to surface warming and increased stratification in open ocean environments (Capotondi et al., 2012; Li et al., 2020). Coupled climate models broadly predict a reduction in marine net primary production resulting from reduced vertical mixing and nutrient supply (Fu et al., 2016; Kwiatkowski et al., 2020). However, the response of any specific ecosystem can be complex as a result of different lower trophic level dynamics and the potential for ecosystem reorganization as a result of the disturbance of climate change (Karl et al., 2021; Lomas et al., 2022; Kahru et al., 2023). Mechanistic understanding of bottom-up ecosystem processes is thus crucial for predicting future changes and enabling effective ecosystem-based fisheries management.

Pelagic ecosystems ultimately rely on “new” nutrients to replace the organic matter removed by the biological carbon pump (Eppley and Peterson, 1979). In deepwater, semi-enclosed basins such as the GoM and Argo Basin, the most likely sources of new nutrients include upwelling of nutrient-rich deep water, nitrogen fixation, and lateral transport of nutrient- or organic matter-rich water from coastal regions (Dugdale and Goering, 1967; Zehr and Capone, 2020; Kelly et al., 2021). Our LIEM, which synthesizes the results from multiple distinct observational studies (Kelly et al., 2021; Yingling et al., 2022; Kehinde et al., 2023; Kranz et al., this issue), shows marked differences between the “new” nutrient sources for the GoM and Argo Basin. In the GoM, the nitrogen necessary to support the ecosystem is derived primarily from lateral advection. This lateral advection is the result of a highly active mesoscale eddy environment, including large Loop Current Eddies, which can impinge upon the continental shelf drawing coastal water across the shelf and into the deep GoM (Zhong and Bracco, 2013; Lee-Sánchez et al., 2022). This process brings both organisms and non-living organic matter (detritus and DOM) into the oligotrophic domain (Kelly et al., 2021; Shropshire et al., 2022). Since these lateral transport processes, rather than vertical mixing, are responsible for nitrogen supply to the ecosystem, increased stratification may not be a driving factor in future change in the system. Rather, more research is needed into whether horizontal stirring (i.e., eddy kinetic energy) will increase in the region.

The Argo Basin appears to function differently. While lateral advection is still quantitatively important (Kehinde et al., 2023), it is not the dominant source of new nutrients supplying the upper euphotic zone where SBT spawn and their larvae navigate this critical phase in their development. Instead, the upper euphotic zone is largely supported by nitrogen fixation (Fig. 7a and Kranz et al., this issue). While there is substantial uncertainty in the response of nitrogen-fixing phytoplankton to climate change (Doléac et al., 2025), it is reasonable to surmise that systems relying on nitrogen fixation (which does not rely on upwelled nitrate) will be less sensitive to increased stratification than systems relying on subsurface nitrate supply. Even so, nitrogen fixation ultimately relies on phosphate supply (typically from upwelling) and Fe availability (typically supplied through a combination of upwelling and atmospheric deposition). This suggests that understanding future changes in Argo Basin ecosystem productivity requires a focus on Fe supply mechanisms and N:P ratios of both subsurface and surface source waters, including inflows from the Indonesian Throughflow. Furthermore, the importance of episodic mixing events should not be discounted in the Argo Basin. Unlike ABT, which typically spawn before the start of the tropical-cyclone season in the GoM, SBT spawning occurs during the summer cyclone season when strong storms can drive deep convective mixing events and introduce substantial nutrients into the surface ocean.

### 4.2. Ecosystem structure and secondary production

Fisheries production depends not only on the net primary productivity of an ecosystem, but also on multiple food-web pathways that determine how efficiently phytoplankton production is channeled to higher trophic levels. Here, we use the term “ecosystem efficiency” to represent this concept and define it as the ratio of top trophic level secondary production (gelatinous predators, pre- and postflexion larvae, and other planktivorous fish) to net primary production. Using this definition, we can see that higher secondary production in the Argo Basin relative to the GoM is the result of not only higher net primary production but also increased ecosystem efficiency (Figs. 8a-c).

**Fig. 8.**
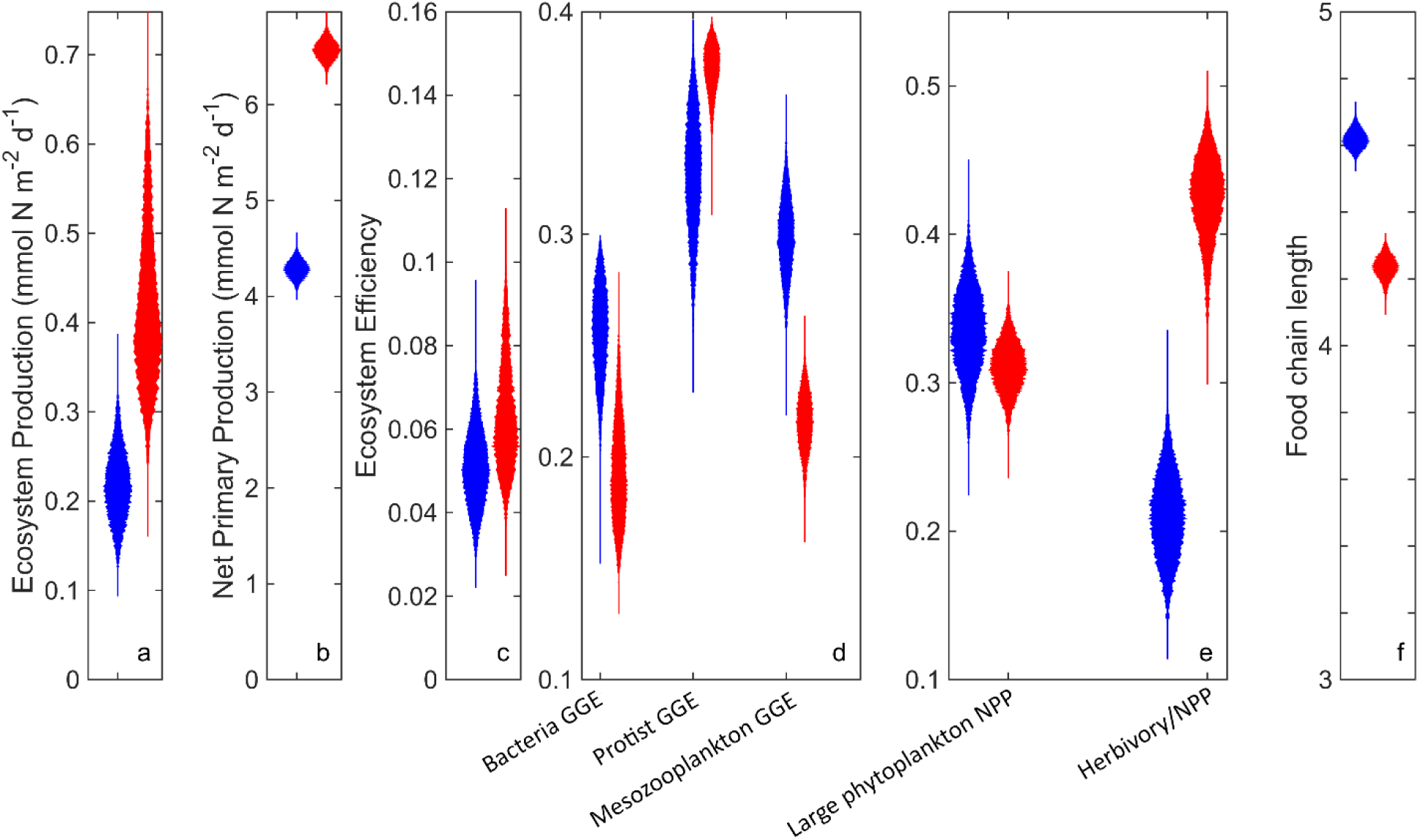
Ecosystem determinants of secondary production. a) Ecosystem production, defined as the secondary production of all top trophic levels (pre-flexion and post-flexion tuna larvae, other planktivorous fish, and gelatinous predators). b) Net primary production (NPP) of all phytoplankton. c) Ecosystem efficiency (defined as ecosystem production / NPP). d) Gross growth efficiencies of bacteria, protists, and mesozooplankton. e) the proportion of net primary production conducted by nano- and micro-phytoplankton (i.e., mixotrophic nanoflagellates + diatoms) and the proportion of NPP that was directly consumed by herbivorous mesozooplankton. f) food chain length (i.e., the production-weighted mean trophic level of top trophic levels). Blue is Gulf of Mexico; red is Argo Basin. All results are summed over the upper and lower euphotic zone.

Understanding the processes that can affect ecosystem efficiency is crucial. The “trophic amplification” hypothesis suggests that ecosystem efficiency will decline globally as a result of climate change (Kwiatkowski et al., 2019; Lotze et al., 2019). These declines in ecosystem efficiency are predicted to result primarily from i) a shift towards smaller phytoplankton and hence food chain elongation and ii) altered zooplankton and fish gross growth efficiencies as warming effects on reduced food availability and increased metabolism (Stock et al., 2014). While the trophic amplification hypothesis is focused on future climate change, our LIEM results give insight into how these processes differ in two similar ecosystems in the present day. Gross growth efficiency is clearly not the primary driver of differences in ecosystem efficiency, because the Argo Basin actually had lower gross growth efficiencies for most taxa (Fig. 8d). Rather, shorter food chain lengths in the Argo Basin, relative to the GoM, drive increased efficiency in the Argo Basin (Fig. 8f). However, contrary to the models on which the trophic amplification hypothesis was elucidated (which typically have limited zooplankton complexity), the relatively short food chains in the Argo Basin were not driven by a higher proportion of large phytoplankton. Indeed, the ratios of cyanobacteria to larger phytoplankton (diatoms + flagellates) were similar between the Argo Basin and GoM (Fig. 8e). Instead, differences between the two systems were driven by appendicularian herbivory on cyanobacteria. In the Argo Basin, appendicularians were preferentially selected by SBT larvae (Swalethorp et al., this issue), which allowed these organisms to maintain relatively low trophic positions (Fig. 7). While ABT in the GoM did show indications of positive selection for appenducularians with increasing availability, overall low availability may have caused larvae to focus on other prey (Shiroza et al., 2022). While results presented herein are derived from a single expedition to the Argo Basin, these shortened food-web pathways may be a general feature of the region. Indeed, zooplankton sampling along 110°E found substantially higher herbivory in tropical waters immediately downstream of the Argo Basin than anywhere else along the transect (Landry *et al*., 2020). Taken together, these results suggest that, despite very warm surface temperatures, the Argo Basin ecosystem efficiently converts phytoplankton production into the secondary production of higher trophic levels.

## 5. Conclusions

Our results show the food-web pathways that support bluefin tuna larvae in the Gulf of Mexico and Argo Basin. Despite many similarities as warm, oligotrophic basins, distinct food-web pathways lead to substantially altered ecosystem efficiency. Direct herbivory of appendicularians on cyanobacteria allows ∼2X higher production of top trophic levels in the Argo Basin, relative to the Gulf of Mexico. These results highlight the importance of understanding zooplankton ecological niches and predator:prey size ratios (Hansen et al., 1994; Fender et al., 2023; Heneghan et al., 2023). Indeed, pelagic tunicates (including appendicularians and salps) have been suggested to increase ecosystem efficiency through their ability to feed at high predator:prey ratios and hence directly access the production of picophytoplankton (Jaspers *et al*., 2023; Stukel *et al*., 2024). Unfortunately, challenges associated with sampling and experimenting with zooplankton make it difficult to assess climate change impacts on diverse zooplankton taxa (Ratnarajah et al., 2023). Furthermore, current climate models typically involve very simplified parameterization of zooplankton taxa and lack the ability to simulate shifts in predator:prey size ratios and ecological niches. Our results highlight the centrality of these concepts in explaining differences in food-web structure and productivity between ecosystems and suggest that they may also play an important role in understanding how these ecosystems will change in the future.

## Supporting information

Code Package

## CRediT authorship contribution statement

**Michael R. Stukel:** Conceptualization, Funding acquisition, Investigation, Writing – original draft, **Raul Laiz-Carrion:** Conceptualization, Funding acquisition, Investigation, Writing – review and editing, **Moira Decima:** Conceptualization, Investigation, Writing – review and editing, **Christian K. Fender:** Investigation, Writing – review and editing, **Sven A. Kranz:** Conceptualization, Funding acquisition, Investigation, Writing – review and editing, **Michael R. Landry:** Conceptualization, Funding acquisition, Investigation, Writing – review and editing, **Estrella Malca:** Conceptualization, Investigation, Writing – review and editing, **Jose M. Quintanilla:** Conceptualization, Investigation, Writing – review and editing, **Karen E Selph:** Conceptualization, Funding acquisition, Investigation, Writing – review and editing, **Rasmus Swalethorp:** Conceptualization, Investigation, Writing – review and editing, **Natalia Yingling:** Investigation, Writing – review and editing

## Acknowledgments

We thank our many collaborators in the BLOOFINZ-IO and BLOOFINZ-GoM research projects and are very grateful to the captain and crew of the R/V Roger Revelle. This study was supported by National Science Foundation Biological Oceanography grants OCE-1851347, OCE-2332036, OCE-1851381 to M.R.S, S.A.K, and K.E.S., respectively, as well as OCE-1851558 and OCE-2404504/OCE-2332036 to M.R.L. and S.A.K., respectively. Funding support was also provided by the INDITUN project PID2021/122862NB/100 UE-FEDER (R.L.-C.) by the Spanish Ministry of Science, Innovation and Universities (MICINN) to R. L.-C. This study was a part of project EP-46 of the 2nd International Indian Ocean Expedition (IIOE-2). Sampling in the Argo-Rowley Terrace Marine Park was done under Australian Government permit AU-COM2021-520 and permit PA2021-00062-1 issued by the Director of National Parks, Australia. The views expressed in this publication do not necessarily represent those of the Director of National Parks or the Australian Government.

## Data availability

Data from this project are available at the Biological and Chemical Oceanography Data Management Office (BCO-DMO) project pages for BLOOFINZ-IO and BLOOFINZ-GoM: https://www.bco-dmo.org/project/819488 and https://www.bco-dmo.org/project/834957

---

Supp. Table 1 Observational results used to constrain the inverse ecosystem model.

Supp. Table 2 Model results showing mean and standard deviation of Markov Chain Monte Carlo simulations for the Gulf of Mexico and Argo Basin.

## References

Bakun, A., 2013. Ocean eddies, predator pits and bluefin tuna: implications of an inferred ‘low risk–limited payoff’reproductive scheme of a (former) archetypical top predator. Fish and Fisheries 14 (3), 424–438.

Biggs, D.C., 1992. Nutrients, plankton, and productivity in a warm-core ring in the western Gulf of Mexico. J. Geophys. Res. Oceans 97 (C2), 2143–2154.

Biggs, D.C., Müller-Karger, F.E., 1994. Ship and satellite observations of chlorophyll stocks in interacting cyclone-anticyclone eddy pairs in the western Gulf of Mexico. J. Geophys. Res. Oceans 99 (C4), 7371–7384.

Borrego-Santos, R., Quintanilla, J.M., Laiz-Carrion, R., García, A., Malca, E., Abascal, F., Die, D., Riveiro, I., Swalethorp, R., Landry, M.R., this issue. Revisiting daily growth and survival insights of southern bluefin tuna (Thunnus maccoyii) larvae in the eastern Indian Ocean. Deep-Sea Res. II.

Brink, K.H., Bahr, F., Shearman, R.K., 2007. Alongshore currents and mesoscale variability near the shelf edge off northwestern Australia. J. Geophys. Res. Oceans 112 (C5).

Capotondi, A., Alexander, M.A., Bond, N.A., Curchitser, E.N., Scott, J.D., 2012. Enhanced upper ocean stratification with climate change in the CMIP3 models. J. Geophys. Res. Oceans 117 (C4), C04031.

Condie, S., Andrewartha, J., 2008. Circulation and connectivity on the Australian North West shelf. Continental Shelf Research 28 (14), 1724–1739.

Décima, M., Swalethorp, R., Cawley, G., Traboni, C., Davies, C.H., Landry, M.R., this issue. Zooplankton trophic processes in the Eastern Indian Ocean Southern Bluefin Tuna spawning region. Deep-Sea Res. II.

Díaz-Barroso, L., Hernández-Carrasco, I., Orfila, A., Reglero, P., Balbín, R., Hidalgo, M., Tintoré, J., Alemany, F., Álvarez-Berastegui, D., 2022. Singularities of surface mixing activity in the Western Mediterranean influence bluefin tuna larval habitats. Mar. Ecol. Prog. Ser. 685, 69–84.

Doléac, S., Lévy, M., El Hourany, R., Bopp, L., 2025. Toward more robust net primary production projections in the North Atlantic Ocean. Biogeosciences 22 (4), 841–862.

Domingues, C.M., Maltrud, M.E., Wijffels, S.E., Church, J.A., Tomczak, M., 2007. Simulated Lagrangian pathways between the Leeuwin Current System and the upper-ocean circulation of the southeast Indian Ocean. Deep-Sea Res. II 54 (8-10), 797–817.

Dugdale, R.C., Goering, J.J., 1967. Uptake of New and Regenerated Forms of Nitrogen in Primary Productivity. Limnol. Oceanogr. 12 (2), 196–206.

Eppley, R.W., Peterson, B.J., 1979. Particulate organic matter flux and planktonic new production in the deep ocean. Nature 282 (5740), 677–680.

Farley, J.H., Davis, T.L.O., 1998. Reproductive dynamics of southern bluefin tuna, Thunnus maccoyii. Fishery Bulletin 96 (2), 223–236.

Fender, C.K., Décima, M., Gutiérrez-Rodriguez, A., Selph, K.E., Yingling, N., Stukel, M.R., 2023. Prey size spectra and predator to prey size ratios of southern ocean salps. Mar. Biol. 170, 40.

Fernández, P.R., Vargas-García, E., Borrego, R., Laiz-Carrión, R., Quintanilla, J.M., Landry, M.R., Kranz, S.A., Stukel, M.R., Kelly, T.B., Vargas-Yáñez, M., this issue. Analysis of environmental data during a lagrangian experiment: The influence of vertical movements. Deep-Sea Res. II.

Fu, W., Randerson, J.T., Moore, J.K., 2016. Climate change impacts on net primary production (NPP) and export production (EP) regulated by increasing stratification and phytoplankton community structure in the CMIP5 models.

Gerard, T., Lamkin, J., Kelly, T.B., Knapp, A.N., Laiz-Carrión, R., Malca, E., Selph, K.E., Shiroza, A., Shropshire, T.A., Stukel, M.R., Swalethorp, R., Yingling, N., Landry, M.R., 2022. Bluefin Larvae in Oligotrophic Ocean Foodwebs, Investigations of Nutrients to Zooplankton: Overview of the BLOOFINZ-Gulf of Mexico program. J. Plankton Res. 44 (5), 600–617.

Green, R.E., Bower, A.S., Lugo-Fernandez, A., 2014. First autonomous bio-optical profiling float in the Gulf of Mexico reveals dynamic biogeochemistry in deep waters. PLoS One 9 (7), e101658.

Hansen, B., Bjornsen, P.K., Hansen, P.J., 1994. The size ratio between planktonic predators and their prey. Limnol. Oceanogr. 39 (2), 395–403.

Heneghan, R.F., Everett, J.D., Blanchard, J.L., Sykes, P., Richardson, A.J., 2023. Climate-driven zooplankton shifts cause large-scale declines in food quality for fish. Nature Climate Change 13 (5), 470–477.

Hood, R.R., Beckley, L.E., Wiggert, J.D., 2017. Biogeochemical and ecological impacts of boundary currents in the Indian Ocean. Prog. Oceanogr. 156, 290–325.

Jaspers, C., Hopcroft, R.R., Kiørboe, T., Lombard, F., López-Urrutia, Á., Everett, J.D., Richardson, A.J., 2023. Gelatinous larvacean zooplankton can enhance trophic transfer and carbon sequestration. Trends in Ecology & Evolution 38 (10), 980–993.

Jenkins, G.P., Young, J.W., Davis, T.L.O., 1991. Density dependence of larval growth of a marine fish, the Southern Bluefin Tuna, *Thunnus maccoyi*. Canadian Journal of Fisheries and Aquatic Sciences 48 (8), 1358–1363.

Kahru, M., Lee, Z., Ohman, M.D., 2023. Multidecadal changes in ocean transparency: Decrease in a coastal upwelling region and increase offshore. Limnol. Oceanogr. 68 (7), 1546–1556.

Karl, D.M., Letelier, R.M., Bidigare, R.R., Björkman, K.M., Church, M.J., Dore, J.E., White, A.E., 2021. Seasonal-to-decadal scale variability in primary production and particulate matter export at Station ALOHA. Prog. Oceanogr. 195, 102563.

Kehinde, O., Bourassa, M., Kranz, S., Landry, M.R., Kelly, T., Stukel, M.R., 2023. Lateral advection of particulate organic matter in the eastern Indian Ocean. J. Geophys. Res. Oceans 128 (5), e2023JC019723.

Kelly, T.B., Knapp, A.N., Landry, M.R., Selph, K.E., Shropshire, T.A., Thomas, R.K., Stukel, M.R., 2021. Lateral advection supports nitrogen export in the oligotrophic open-ocean Gulf of Mexico. Nature Communications 12 (1), 3325.

Knapp, A.N., Thomas, R.K., Stukel, M.R., Kelly, T.B., Landry, M.R., Selph, K.E., Malca, E., Gerard, T., Lamkin, J., 2021. Constraining the sources of nitrogen fueling export production in the Gulf of Mexico using nitrogen isotope budgets. J. Plankton Res.

Kones, J.K., Soetaert, K., van Oevelen, D., Owino, J.O., 2009. Are network indices robust indicators of food web functioning? A Monte Carlo approach. Ecol. Model. 220 (3), 370–382.

Kranz, S.A., Rose, J., Stukel, M.R., Selph, K.E., Yingling, N., Landry, M.R., this issue. Primary productivity and N_2_-fixation in the eastern Indian Ocean: bottom-up support for an ecologically and economically important ecosystem Deep-Sea Res. II.

Kwiatkowski, L., Aumont, O., Bopp, L., 2019. Consistent trophic amplification of marine biomass declines under climate change. Global change biology 25 (1), 218–229.

Kwiatkowski, L., Torres, O., Bopp, L., Aumont, O., Chamberlain, M., Christian, J.R., Dunne, J.P., Gehlen, M., Ilyina, T., John, J.G., 2020. Twenty-first century ocean warming, acidification, deoxygenation, and upper-ocean nutrient and primary production decline from CMIP6 model projections. Biogeosciences 17 (13), 3439–3470.

Laiz-Carrión, R., Borrego-Santos, R., M., Q.J., Quezada-Romegialli, C., Malca, E., Swalethorp, R., Abascal, F., Pennino, M.G., Godoy, M.A., Die, D., Landry, M.R., this issue. Trophic specialization supports better growth performance in southern bluefin, albacore and skipjack tuna larvae from the eastern Indian Ocean. Deep-Sea Res. II.

Landry, M.R., Beckley, L.E., Muhling, B.A., 2019. Climate sensitivities and uncertainties in food-web pathways supporting larval bluefin tuna in subtropical oligotrophic oceans. ICES J. Mar. Sci. 76 (2), 359–369.

Landry, M.R., Hood, R.R., Davies, C.H., 2020. Mesozooplankton biomass and temperature-enhanced grazing along a 110°E transect in the eastern Indian Ocean. Mar. Ecol. Prog. Ser. 649, 1–19.

Landry, M.R., Laiz-Carrión, R., al., e., this issue-a. Overview of BLOOFINZ/INDITUN investigations of the southern bluefin spawning region off northwest Australia, January-March 2022. Deep-Sea Res. II.

Landry, M.R., Ohman, M.D., Goericke, R., Stukel, M.R., Tsyrklevich, K., 2009. Lagrangian studies of phytoplankton growth and grazing relationships in a coastal upwelling ecosystem off Southern California. Prog. Oceanogr. 83, 208–216.

Landry, M.R., Rivera, S.R., Stukel, M.R., Selph, K.E., 2023. Comparison of bacterial carbon production estimates from dilution and 3H-leucine methods across a strong gradient in ocean productivity. Limnol. Oceanogr. Meth. 21 (6), 295–306.

Landry, M.R., Selph, K.E., Hood, R.R., Davies, C.H., Beckley, L.E., 2021a. Low temperature sensitivity of picophytoplankton P: B ratios and growth rates across a natural 10° C temperature gradient in the oligotrophic Indian Ocean. Limnology and Oceanography Letters.

Landry, M.R., Selph, K.E., Stukel, M.R., Swalethorp, R., Kelly, T.B., Beatty, J.L., Quackenbush, C.R., 2021b. Microbial food web dynamics in the oceanic Gulf of Mexico. J. Plankton Res.

Landry, M.R., Stukel, M.R., Yingling, N., Selph, K.E., Kranz, S.A., Fender, C.K., Swalethorp, R., Bhabu, R., this issue-b. Microbial food web dynamics in tropical waters of the bluefin tuna spawning region off northwestern Australia. Deep-Sea Res. II.

Landry, M.R., Swalethorp, R., 2021. Mesozooplankton biomass, grazing and trophic structure in the bluefin tuna spawning area of the oceanic Gulf of Mexico. J. Plankton Res.

Lee-Sánchez, E., Camacho-Ibar, V.F., Velásquez-Aristizábal, J.A., Valencia-Gasti, J.A., Samperio-Ramos, G., 2022. Impacts of mesoscale eddies on the nitrate distribution in the deep-water region of the Gulf of Mexico. J. Mar. Sys. 229, 103721.

Li, G., Cheng, L., Zhu, J., Trenberth, K.E., Mann, M.E., Abraham, J.P., 2020. Increasing ocean stratification over the past half-century. Nature Climate Change 10 (12), 1116–1123.

Lindo-Atichati, D., Bringas, F., Goni, G., Muhling, B., Muller-Karger, F.E., Habtes, S., 2012. Varying mesoscale structures influence larval fish distribution in the northern Gulf of Mexico. Mar. Ecol. Prog. Ser. 463, 245–257.

Llopiz, J.K., Hobday, A.J., 2015. A global comparative analysis of the feeding dynamics and environmental conditions of larval tunas, mackerels, and billfishes. Deep-Sea Res. II 113, 113–124.

Llopiz, J.K., Muhling, B.A., Lamkin, J.T., 2015. Feeding dynamics of Atlantic Bluefin Tuna (*Thunnus thynnus*) larvae in the Gulf of Mexico. Collect. Vol. Sci. Pap. ICCAT 71 (4), 1710–1715.

Lomas, M.W., Bates, N.R., Johnson, R.J., Steinberg, D.K., Tanioka, T., 2022. Adaptive carbon export response to warming in the Sargasso Sea. Nature Communications 13 (1), 1211.

Lotze, H.K., Tittensor, D.P., Bryndum-Buchholz, A., Eddy, T.D., Cheung, W.W., Galbraith, E.D., Barange, M., Barrier, N., Bianchi, D., Blanchard, J.L., 2019. Global ensemble projections reveal trophic amplification of ocean biomass declines with climate change. Proc. Natl. Acad. Sci. U. S. A. 116 (26), 12907–12912.

Malca, E., Shropshire, T., Landry, M.R., Quintanilla, J.M., Laiz-CarriÓn, R., Shiroza, A., Stukel, M.R., Lamkin, J., Gerard, T., Swalethorp, R., 2022. Influence of food quality on larval growth of Atlantic bluefin tuna (Thunnus thynnus) in the Gulf of Mexico. J. Plankton Res. 44 (5), 747–762.

Muhling, B.A., Lamkin, J.T., Alemany, F., García, A., Farley, J., Ingram, G.W., Berastegui, D.A., Reglero, P., Carrion, R.L., 2017. Reproduction and larval biology in tunas, and the importance of restricted area spawning grounds. Rev. Fish. Biol. Fisheries 27 (4), 697–732.

Quadfasel, D., Frische, A., Cresswell, G., 1996. The circulation in the source area of the South Equatorial Current in the eastern Indian Ocean. J. Geophys. Res. Oceans 101 (C5), 12483–12488.

Quintanilla, J.M., this issue. Optimal Maternal Feeding Isotopic Niche: influence of breeder trophic behaviour on larval growth and survival in bluefin tuna species. Deep-Sea Res. II.

Quintanilla, J.M., Borrego-Santos, R., Malca, E., Swalethorp, R., Landry, M.R., Gerard, T., Lamkin, J., García, A., Laiz-Carrión, R., 2024. Maternal Effects and Trophodynamics Drive Interannual Larval Growth Variability of Atlantic Bluefin Tuna (Thunnus thynnus) from the Gulf of Mexico. Animals 14 (9), 1319.

Raes, E.J., Hörstmann, C., Landry, M.R., Beckley, L.E., Marin, M., Thompson, P., Antoine, D., Focardi, A., O’Brien, J., Ostrowski, M., Waite, A.M., 2022. Dynamic change in an ocean desert: Microbial diversity and trophic transfer along the 110 °E meridional in the Indian Ocean. Deep-Sea Res. II 201, 105097.

Ratnarajah, L., Abu-Alhaija, R., Atkinson, A., Batten, S., Bax, N.J., Bernard, K.S., Canonico, G., Cornils, A., Everett, J.D., Grigoratou, M., Ishak, N.H.A., Johns, D., Lombard, F., Muxagata, E., Ostle, C., Pitois, S., Richardson, A.J., Schmidt, K., Stemmann, L., Swadling, K.M., Yang, G., Yebra, L., 2023. Monitoring and modelling marine zooplankton in a changing climate. Nature Communications 14 (1), 564.

Reglero, P., Tugores, M.P., Titelman, J., Santandreu, M., Martin, M., Balbin, R., Alvarez-Berastegui, D., Torres, A.P., Calcina, N., Leyva, L., 2025. Bluefin tuna (Thunnus thynnus) larvae exploit rare food sources to break food limitations in their warm oligotrophic environment. J. Plankton Res. 47 (2), fbaf006.

Rooker, J.R., Bremer, J.R.A., Block, B.A., Dewar, H., De Metrio, G., Corriero, A., Kraus, R.T., Prince, E.D., Rodriguez-Marin, E., Secor, D.H., 2007. Life history and stock structure of Atlantic bluefin tuna (*Thunnus thynnus*). Reviews in Fisheries Science 15 (4), 265–310.

Selph, K.E., Lampe, R.H., Yingling, N., Bhabu, R., Kranz, S.A., Allen, A.E., Landry, M.R., this issue-a. Phytoplankton community composition in the oligotrophic Argo Basin of the eastern Indian Ocean. Deep-Sea Res. II.

Selph, K.E., Swalethorp, R., Stukel, M.R., Kelly, T.B., Knapp, A.N., Fleming, K., Hernandez, T., Landry, M.R., 2021. Phytoplankton community composition and biomass in the oligotrophic Gulf of Mexico. J. Plankton Res.

Selph, K.E., Yingling, N., Traboni, C., Landry, M.R., this issue-b. Acidic vacuole-containing organisms are a majority of the eukaryotic microbial community in oligotrophic Argo Basin waters (eastern Indian Ocean).

Shiroza, A., Malca, E., Lamkin, J., Gerard, T., Landry, M.R., Stukel, M.R., Laiz-Carrión, R., Swalethorp, R., 2022. Active prey selection in developing larvae of Atlantic bluefin tuna (*Thunnus thynnus*) in spawning grounds of the Gulf of Mexico. J. Plankton Res. 44 (5), 728–746.

Shropshire, T.A., Morey, S.L., Chassignet, E.P., Bozec, A., Coles, V.J., Landry, M.R., Swalethorp, R., Zapfe, G., Stukel, M.R., 2020. Quantifying spatiotemporal variability in zooplankton dynamics in the Gulf of Mexico with a physical-biogeochemical model. Biogeosciences 17, 3385–3407.

Shropshire, T.A., Morey, S.L., Chassignet, E.P., Karnauskas, M., Coles, V.J., Malca, E., Laiz-Carrión, R., Fiksen, Ø., Reglero, P., Shiroza, A., Quintanilla, J.M., Gerard, T., Lamkin, J.T., Stukel, M.R., 2022. Trade-offs between risks of predation and starvation in larvae make the shelf break an optimal spawning location for Atlantic bluefin tuna. J. Plankton Res. 44 (5), 782–798.

Stock, C., Dunne, J., John, J., 2014. Drivers of trophic amplification of ocean productivity trends in a changing climate. Biogeosciences 11 (24), 7125–7135.

Stukel, M.R., Biard, T., Décima, M., Fender, C.K., Kehinde, O., Kelly, T.B., Kranz, S.A., Laget, M., Landry, M.R., Yingling, N., this issue. Sinking particle export within and beneath the euphotic zone in the eastern Indian Ocean. Deep-Sea Res. II.

Stukel, M.R., Décima, M., Fender, C.K., Gutierrez-Rodriguez, A., Selph, K.E., 2024. Gelatinous filter feeders increase ecosystem efficiency. Communications Biology 7 (1), 1039.

Stukel, M.R., Décima, M., Kelly, T.B., 2018a. A new approach for incorporating 15N isotopic data into linear inverse ecosystem models with Markov Chain Monte Carlo sampling. PloS one 13 (6), e0199123.

Stukel, M.R., Décima, M., Landry, M.R., Selph, K.E., 2018b. Nitrogen and isotope flows through the Costa Rica Dome upwelling ecosystem: The crucial mesozooplankton role in export flux. Global Biogeochem. Cycles 32, 1815–1832.

Stukel, M.R., Gerard, T., Kelly, T.B., Knapp, A.N., Laiz-Carrión, R., Lamkin, J., Landry, M.R., Malca, E., Shiroza, A., A., S.T., Selph, K.E., Swalethorp, R., 2022. Plankton food webs of the Gulf of Mexico spawning grounds of Atlantic Bluefin tuna. J. Plankton Res. 44 (5), 763–781.

Stukel, M.R., Kelly, T.B., Landry, M.R., Selph, K.E., Swalethorp, R., 2021. Sinking carbon, nitrogen, and pigment flux within and beneath the euphotic zone in the oligotrophic, open-ocean Gulf of Mexico. J. Plankton Res. 44 (5), 711–727.

Stukel, M.R., Landry, M.R., Ohman, M.D., Goericke, R., Samo, T., Benitez-Nelson, C.R., 2012. Do inverse ecosystem models accurately reconstruct plankton trophic flows? Comparing two solution methods using field data from the California Current. J. Mar. Sys. 91 (1), 20–33.

Swalethorp, R., Malca, E., Shiroza, A., Kim, L., Decima, M., Quintanilla, J.M., Borrego-Santos, R., Davies, C.H., Die, D., Beckley, L.E., Traboni, C., Cawley, G., Walsh, K., Landry, M.R., Laiz-Carrión, R., this issue. Selective feeding in Southern Bluefin Tuna (Thunnus maccoyii) larvae in the Argo Basin, Eastern Indian Ocean spawning ground. Deep-Sea Res. II.

Teo, S.L., Boustany, A., Dewar, H., Stokesbury, M.J., Weng, K.C., Beemer, S., Seitz, A.C., Farwell, C.J., Prince, E.D., Block, B.A., 2007. Annual migrations, diving behavior, and thermal biology of Atlantic bluefin tuna, *Thunnus thynnus*, on their Gulf of Mexico breeding grounds. Mar. Biol. 151 (1), 1–18.

Tilley, J.D., Butler, C.M., Suárez-Morales, E., Franks, J.S., Hoffmayer, E.R., Gibson, D.P., Comyns, B.H., Ingram Jr, G.W., Blake, E.M., 2016. Feeding ecology of larval Atlantic bluefin tuna, *Thunnus thynnus*, from the central Gulf of Mexico. Bull. Mar. Sci. 92 (3), 321–334.

Van den Meersche, K., Soetaert, K., Van Oevelen, D., 2009. xSample(): An R function for sampling linear inverse problems. J. Stat. Softw., Code Snippets 30 (1), 1–15.

van Oevelen, D., Van den Meersche, K., Meysman, F.J.R., Soetaert, K., Middelburg, J.J., Vezina, A.F., 2010. Quantifying food web flows using linear inverse models. Ecosystems 13 (1), 32–45.

Vézina, A.F., Platt, T., 1988. Food web dynamics in the ocean .1. Best estimates of flow networks using inverse methods. Mar. Ecol. Prog. Ser. 42 (3), 269–287.

Yingling, N., Kelly, T.B., Selph, K.E., Landry, M.R., Knapp, A.N., Kranz, S.A., Stukel, M.R., 2022. Taxon-specific phytoplankton growth, nutrient limitation, and light limitation in the oligotrophic Gulf of Mexico. J. Plankton Res. 44 (5), 656–676.

Yingling, N., Selph, K.E., Landry, M.R., Kranz, S.A., Johnson, M., Stukel, M.R., this issue. Phytoplankton Nutrient Uptake, Abundance, Biomass and Community Composition in the Oligotrophic Argo Basin, Indian Ocean. Deep-Sea Res. II.

Young, J.W., Davis, T.L.O., 1990. Feeding ecology of larvae of southern bluefin, albacore, and skipjack tunas (Pisces, Scombridae) in the eastern Indian Ocean. Mar. Ecol. Prog. Ser. 61 (1-2), 17–29.

Zehr, J.P., Capone, D.G., 2020. Changing perspectives in marine nitrogen fixation. Science 368 (6492).

Zhong, Y.S., Bracco, A., 2013. Submesoscale impacts on horizontal and vertical transport in the Gulf of Mexico. J. Geophys. Res. Oceans 118 (10), 5651–5668.

