## Supplementary material for "Comparative food-web analysis of bluefin tuna spawning habitats in the eastern Indian Ocean and Gulf of Mexico": Code Package: AddPanelLabels.m

```
function[output]=AddPanelLabels(varargin)

%This function adds panel labels (a,b,c,d, etc.) to the corners of plots

%Call with:
%AddPanelLabels(LABEL,LOCATION) or AddPanelLabels('b','topleft','YPADDING',0.02,'XPADDING',0.05,'FONTSIZE',9)

%LABEL is the label (e.g., "a" or "b" or "c") to add to the panel
%LOCATION is the location ("topleft","topright","bottomleft", or "bottomright")

%Other options are: 
% FONTSIZE to specify the FontSize
% XPADDING to specify the percentage of the figure to move over from chosen x axis (default is 0.1 or 10%)
% YPADDING to specify the percentage of the figure to move over from chosen y axis (default is 0.1 or 10%)

%Specify the other options by adding (for example): AddPanelLabels('a','topright',XPADDING,0.05,FONTSIZE,12)


args=cell(varargin)
nu=numel(args)

default.values={[],'topleft'};
default.values(1:nu)=args;

LABEL = varargin{1};
LOCATION = varargin{2};
XPADDING = 0.05;
YPADDING = 0.05;
FONTSIZE = 10;
col = 'k'

if nu>2
    for i=3:nu
        if strcmp( args{i}, 'XPADDING')
            XPADDING = args{i+1}
        elseif strcmp( args{i}, 'YPADDING')
            YPADDING = args{i+1}
        elseif strcmp( args{i}, 'FONTSIZE')
            FONTSIZE = args{i+1}
        elseif strcmp( args{i}, 'COLOR')
            col = args{i+1}
        end
    end
end

if strcmp(LOCATION,'bottomright')
    TEXTALIGNUPDOWN = 'bottom'
elseif strcmp(LOCATION,'bottomleft')
    TEXTALIGNUPDOWN = 'bottom'
elseif strcmp(LOCATION,'topright')
    TEXTALIGNUPDOWN = 'top'
elseif strcmp(LOCATION,'topleft')
    TEXTALIGNUPDOWN = 'top'
end


%The following code flips things if the y axes are reversed
if strcmp(get(gca,'YDir'),'reverse')
    if strcmp(LOCATION,'bottomright')
        LOCATION = 'topright'
    elseif strcmp(LOCATION,'bottomleft')
        LOCATION = 'topleft'
    elseif strcmp(LOCATION,'topright')
        LOCATION = 'bottomright'
    elseif strcmp(LOCATION,'topleft')
        LOCATION = 'bottomleft'
    end
end

x=get(gca,'XLim')
y=get(gca,'YLim')
if strcmp(get(gca,'XScale'),'linear') & strcmp(get(gca,'YScale'),'linear')
    if strcmp(LOCATION,'topright')
        text( x(2)-(x(2)-x(1))*XPADDING, y(2)-(y(2)-y(1))*YPADDING, LABEL,'FontSize',FONTSIZE,'HorizontalAlignment','right','VerticalAlignment', TEXTALIGNUPDOWN,'Color',col);
    elseif strcmp(LOCATION,'topleft')
        text( x(1)+(x(2)-x(1))*XPADDING, y(2)-(y(2)-y(1))*YPADDING, LABEL,'FontSize',FONTSIZE,'HorizontalAlignment','left','VerticalAlignment', TEXTALIGNUPDOWN,'Color',col);
    elseif strcmp(LOCATION,'bottomleft')
        text( x(1)+(x(2)-x(1))*XPADDING, y(1)+(y(2)-y(1))*YPADDING, LABEL,'FontSize',FONTSIZE,'HorizontalAlignment','left','VerticalAlignment', TEXTALIGNUPDOWN,'Color',col);
    elseif strcmp(LOCATION,'bottomright')
        text( x(2)-(x(2)-x(1))*XPADDING, y(1)+(y(2)-y(1))*YPADDING, LABEL,'FontSize',FONTSIZE,'HorizontalAlignment','right','VerticalAlignment', TEXTALIGNUPDOWN,'Color',col);
    end
elseif strcmp(get(gca,'XScale'),'log') & strcmp(get(gca,'YScale'),'log')
    if strcmp(LOCATION,'topright')
        text( 10.^( log10(x(2))-(log10(x(2))-log10(x(1)))*XPADDING ), 10.^( log10(y(2))-(log10(y(2))-log10(y(1)))*YPADDING ), LABEL,'FontSize',FONTSIZE,'HorizontalAlignment','right','VerticalAlignment', TEXTALIGNUPDOWN,'Color',col);
    elseif strcmp(LOCATION,'topleft')
        text( 10.^( log10(x(1))+(log10(x(2))-log10(x(1)))*XPADDING ), 10.^( log10(y(2))-(log10(y(2))-log10(y(1)))*YPADDING ), LABEL,'FontSize',FONTSIZE,'HorizontalAlignment','left','VerticalAlignment', TEXTALIGNUPDOWN,'Color',col);
    elseif strcmp(LOCATION,'bottomleft')
        text( 10.^( log10(x(1))+(log10(x(2))-log10(x(1)))*XPADDING ), 10.^( log10(y(1))+(log10(y(2))-log10(y(1)))*YPADDING ), LABEL,'FontSize',FONTSIZE,'HorizontalAlignment','left','VerticalAlignment', TEXTALIGNUPDOWN,'Color',col);
    elseif strcmp(LOCATION,'bottomright')
        text( 10.^( log10(x(2))-(log10(x(2))-log10(x(1)))*XPADDING ), 10.^( log10(y(1))+(log10(y(2))-log10(y(1)))*YPADDING ), LABEL,'FontSize',FONTSIZE,'HorizontalAlignment','right','VerticalAlignment', TEXTALIGNUPDOWN,'Color',col);
        [10.^( log10(x(2))-(log10(x(2))-log10(x(1)))*XPADDING ) , 10.^( log10(y(1))+(log10(y(2))-log10(y(1)))*YPADDING )]
    end
elseif strcmp(get(gca,'XScale'),'log') & strcmp(get(gca,'YScale'),'linear')
    if strcmp(LOCATION,'topright')
        text( 10.^( log10(x(2))-(log10(x(2))-log10(x(1)))*XPADDING ),  y(2)-(y(2)-y(1))*YPADDING , LABEL,'FontSize',FONTSIZE,'HorizontalAlignment','right','VerticalAlignment', TEXTALIGNUPDOWN,'Color',col);
    elseif strcmp(LOCATION,'topleft')
        text( 10.^( log10(x(1))+(log10(x(2))-log10(x(1)))*XPADDING ),  y(2)-(y(2)-y(1))*YPADDING , LABEL,'FontSize',FONTSIZE,'HorizontalAlignment','left','VerticalAlignment', TEXTALIGNUPDOWN,'Color',col);
    elseif strcmp(LOCATION,'bottomleft')
        text( 10.^( log10(x(1))+(log10(x(2))-log10(x(1)))*XPADDING ),  y(1)+(y(2)-y(1))*YPADDING , LABEL,'FontSize',FONTSIZE,'HorizontalAlignment','left','VerticalAlignment', TEXTALIGNUPDOWN,'Color',col);
    elseif strcmp(LOCATION,'bottomright')
    'IM in here'
        text( 10.^( log10(x(2))-(log10(x(2))-log10(x(1)))*XPADDING ),  y(1)+(y(2)-y(1))*YPADDING , LABEL,'FontSize',FONTSIZE,'HorizontalAlignment','right','VerticalAlignment', TEXTALIGNUPDOWN,'Color',col);
        [10.^( log10(x(2))-(log10(x(2))-log10(x(1)))*XPADDING ) , y(1)+(y(2)-y(1))*YPADDING ]
    end
elseif strcmp(get(gca,'XScale'),'linear') & strcmp(get(gca,'YScale'),'log')
    if strcmp(LOCATION,'topright')
        text( x(2)-(x(2)-x(1))*XPADDING, 10.^( log10(y(2))-(log10(y(2))-log10(y(1)))*YPADDING ), LABEL,'FontSize',FONTSIZE,'HorizontalAlignment','right','VerticalAlignment', TEXTALIGNUPDOWN,'Color',col);
    elseif strcmp(LOCATION,'topleft')
        text( x(1)+(x(2)-x(1))*XPADDING, 10.^( log10(y(2))-(log10(y(2))-log10(y(1)))*YPADDING ), LABEL,'FontSize',FONTSIZE,'HorizontalAlignment','left','VerticalAlignment', TEXTALIGNUPDOWN,'Color',col);
    elseif strcmp(LOCATION,'bottomleft')
        text( x(1)+(x(2)-x(1))*XPADDING, 10.^( log10(y(1))+(log10(y(2))-log10(y(1)))*YPADDING ), LABEL,'FontSize',FONTSIZE,'HorizontalAlignment','left','VerticalAlignment', TEXTALIGNUPDOWN,'Color',col);
    elseif strcmp(LOCATION,'bottomright')
        text( x(2)-(x(2)-x(1))*XPADDING, 10.^( log10(y(1))+(log10(y(2))-log10(y(1)))*YPADDING ), LABEL,'FontSize',FONTSIZE,'HorizontalAlignment','right','VerticalAlignment', TEXTALIGNUPDOWN,'Color',col);
        [10.^( log10(x(2))-(log10(x(2))-log10(x(1)))*XPADDING ) , 10.^( log10(y(1))+(log10(y(2))-log10(y(1)))*YPADDING )]
    end
end

output=0;
```
