## Supplementary material for "Comparative food-web analysis of bluefin tuna spawning habitats in the eastern Indian Ocean and Gulf of Mexico": Code Package: AppendDoodle3.m

```
function[output]=AppendDoodle3(Bkgd,doodle,location,masktype,mask,flag);
output=0;

warning off

% Bkgd=fig;
% doodle=marsnow;
% location=MarSnowLoc-[szmarsnow(2) szmarsnow(1)]/2;
% masktype=3;
% mask=double(marsnowmask);
% flag=0;


%Masktype==1 is a vector file equivalent to that returned by find command
%Masktype==2 is a matrix of values that tells you how much weight to give
%each pixel from doodle

xdimbkgd=length(Bkgd(1,:,1));
ydimbkgd=length(Bkgd(:,1,1));

xdimdoodle=length(doodle(1,:,1));
ydimdoodle=length(doodle(:,1,1));

xloc=location(1);
yloc=location(2);

if xloc>=xdimbkgd
    output=Bkgd;
    if flag==1
    'Location is out of range'
    end
    return 
end
if yloc>=ydimbkgd
    output=Bkgd;
    if flag==1
    'Location is out of range'
    end
    return
end
if xloc+xdimdoodle<1
    output=Bkgd;
    if flag==1
    'Location is out of range'
    end
    return
end
if yloc+ydimdoodle<1
    output=Bkgd;
    if flag==1
    'Location is out of range'
    end
    return
end


if masktype==2
    if max(max(mask))>1
        mask=double(mask);
        mask=mask/max(max(mask));
    end
end


if masktype==3
    if max(max(mask))>1
        mask=double(mask);
        mask=mask/255;
    end
end

% If doodle sticks out on the high end
if xloc+xdimdoodle>xdimbkgd
    if masktype==1
        doodle1=doodle(:,:,1);
        doodle2=doodle(:,:,2);
        doodle3=doodle(:,:,3);
        doodletemp1=NaN(size(doodle1));
        doodletemp2=NaN(size(doodle2));
        doodletemp3=NaN(size(doodle3));
        doodletemp1(mask)=doodle1(mask);
        doodletemp2(mask)=doodle2(mask);
        doodletemp3(mask)=doodle3(mask);
        doodle=doodletemp1;
        doodle(:,:,2)=doodletemp2;
        doodle(:,:,3)=doodletemp3;
        clear doodle1 doodle2 doodle3 doodletemp1 doodletemp2 doodletemp3
    end
    doodle(:,xdimdoodle-(xloc+xdimdoodle-xdimbkgd)+1:xdimdoodle,:)=[];
    if masktype==1
        mask=find(doodle(:,:,1)>-1000);
    elseif masktype==2 | masktype==3
        size(mask)
        xdimdoodle-(xloc+xdimdoodle-xdimbkgd)+1
        xdimdoodle
        mask(:,xdimdoodle-(xloc+xdimdoodle-xdimbkgd)+1:xdimdoodle,:)=[];
    end
    xdimdoodle=length(doodle(1,:,1));
end
if yloc+ydimdoodle>ydimbkgd
    if masktype==1
        doodle1=doodle(:,:,1);
        doodle2=doodle(:,:,2);
        doodle3=doodle(:,:,3);
        doodletemp1=NaN(size(doodle1));
        doodletemp2=NaN(size(doodle2));
        doodletemp3=NaN(size(doodle3));
        doodletemp1(mask)=doodle1(mask);
        doodletemp2(mask)=doodle2(mask);
        doodletemp3(mask)=doodle3(mask);
        doodle=doodletemp1;
        doodle(:,:,2)=doodletemp2;
        doodle(:,:,3)=doodletemp3;
        clear doodle1 doodle2 doodle3 doodletemp1 doodletemp2 doodletemp3
    end
    doodle(ydimdoodle-(yloc+ydimdoodle-ydimbkgd)+1:ydimdoodle,:,:)=[];
    if masktype==1
        mask=find(doodle(:,:,1)>-1000);
    elseif masktype==2 | masktype==3
        mask(ydimdoodle-(yloc+ydimdoodle-ydimbkgd)+1:ydimdoodle,:,:)=[];
    end
    ydimdoodle=length(doodle(:,1,1));
end


% If doodle sticks out on the low end
if xloc<1
    if masktype==1
        doodle1=doodle(:,:,1);
        doodle2=doodle(:,:,2);
        doodle3=doodle(:,:,3);
        doodletemp1=NaN(size(doodle1));
        doodletemp2=NaN(size(doodle2));
        doodletemp3=NaN(size(doodle3));
        doodletemp1(mask)=doodle1(mask);
        doodletemp2(mask)=doodle2(mask);
        doodletemp3(mask)=doodle3(mask);
        doodle=doodletemp1;
        doodle(:,:,2)=doodletemp2;
        doodle(:,:,3)=doodletemp3;
        clear doodle1 doodle2 doodle3 doodletemp1 doodletemp2 doodletemp3
    end
    doodle(:,1:-xloc,:)=[];
    if masktype==1
        mask=find(doodle(:,:,1)>-1000);
    elseif masktype==2 | masktype==3
        mask(:,1:-xloc,:)=[];
    end
    xdimdoodle=length(doodle(1,:,1));
    xloc=1;
end
if yloc<1
    if masktype==1
        doodle1=doodle(:,:,1);
        doodle2=doodle(:,:,2);
        doodle3=doodle(:,:,3);
        doodletemp1=NaN(size(doodle1));
        doodletemp2=NaN(size(doodle2));
        doodletemp3=NaN(size(doodle3));
        doodletemp1(mask)=doodle1(mask);
        doodletemp2(mask)=doodle2(mask);
        doodletemp3(mask)=doodle3(mask);
        doodle=doodletemp1;
        doodle(:,:,2)=doodletemp2;
        doodle(:,:,3)=doodletemp3;
        clear doodle1 doodle2 doodle3 doodletemp1 doodletemp2 doodletemp3
    end
    doodle(1:-yloc,:,:)=[];
    if masktype==1
        mask=find(doodle(:,:,1)>-1000);
    elseif masktype==2 | masktype==3
        mask(1:-yloc,:,:)=[];
    end
    ydimdoodle=length(doodle(:,1,1));
    yloc=1;
end


if masktype==1
    temp1=Bkgd(yloc:yloc+ydimdoodle-1,xloc:xloc+xdimdoodle-1,1);
    temp2=Bkgd(yloc:yloc+ydimdoodle-1,xloc:xloc+xdimdoodle-1,2);
    temp3=Bkgd(yloc:yloc+ydimdoodle-1,xloc:xloc+xdimdoodle-1,3);
    doodletemp1=doodle(:,:,1);
    doodletemp2=doodle(:,:,2);
    doodletemp3=doodle(:,:,3);
    temp1(mask)=doodletemp1(mask);
    temp2(mask)=doodletemp2(mask);
    temp3(mask)=doodletemp3(mask);
    temp=temp1;
    temp(:,:,2)=temp2;
    temp(:,:,3)=temp3;
    output=Bkgd;
    output(yloc:yloc+ydimdoodle-1,xloc:xloc+xdimdoodle-1,:)=temp;
elseif masktype==2 | masktype==3    
    temp1=double(Bkgd(yloc:min([yloc+ydimdoodle-1,ydimbkgd]),xloc:min([xloc+xdimdoodle-1,xdimbkgd]),1));
    temp2=double(Bkgd(yloc:min([yloc+ydimdoodle-1,ydimbkgd]),xloc:min([xloc+xdimdoodle-1,xdimbkgd]),2));
    temp3=double(Bkgd(yloc:min([yloc+ydimdoodle-1,ydimbkgd]),xloc:min([xloc+xdimdoodle-1,xdimbkgd]),3));
    if xdimdoodle>xdimbkgd & ydimdoodle>ydimbkgd
        doodletemp1=double(doodle(2-yloc:min([1+ydimdoodle-yloc,ydimbkgd]),2-xloc:min([1+xdimdoodle-xloc,xdimbkgd]),1));
        doodletemp2=double(doodle(2-yloc:min([1+ydimdoodle-yloc,ydimbkgd]),2-xloc:min([1+xdimdoodle-xloc,xdimbkgd]),2));
        doodletemp3=double(doodle(2-yloc:min([1+ydimdoodle-yloc,ydimbkgd]),2-xloc:min([1+xdimdoodle-xloc,xdimbkgd]),3));
        mask=mask(2-yloc:min([1+ydimdoodle-yloc,ydimbkgd]),2-xloc:min([1+xdimdoodle-xloc,xdimbkgd]));
    elseif xdimdoodle>xdimbkgd
        doodletemp1=double(doodle(:,2-xloc:min([1+xdimdoodle-xloc,xdimbkgd]),1));
        doodletemp2=double(doodle(:,2-xloc:min([1+xdimdoodle-xloc,xdimbkgd]),2));
        doodletemp3=double(doodle(:,2-xloc:min([1+xdimdoodle-xloc,xdimbkgd]),3));
        mask=mask(:,2-xloc:min([1+xdimdoodle-xloc,xdimbkgd]));
    elseif ydimdoodle>ydimbkgd
        doodletemp1=double(doodle(2-yloc:min([1+ydimdoodle-yloc,ydimbkgd]),:,1));
        doodletemp2=double(doodle(2-yloc:min([1+ydimdoodle-yloc,ydimbkgd]),:,2));
        doodletemp3=double(doodle(2-yloc:min([1+ydimdoodle-yloc,ydimbkgd]),:,3));
        mask=mask(2-yloc:min([1+ydimdoodle-yloc,ydimbkgd]),:);
    else
        doodletemp1=double(doodle(:,:,1));
        doodletemp2=double(doodle(:,:,2));
        doodletemp3=double(doodle(:,:,3));
    end
    temp1=uint8((1-mask).*temp1+mask.*doodletemp1);
    temp2=uint8((1-mask).*temp2+mask.*doodletemp2);
    temp3=uint8((1-mask).*temp3+mask.*doodletemp3);
    temp=temp1;
    temp(:,:,2)=temp2;
    temp(:,:,3)=temp3;
    output=Bkgd;
    output(yloc:min([yloc+ydimdoodle-1,ydimbkgd]),xloc:min([xloc+xdimdoodle-1,xdimbkgd]),:)=temp;
end
```
