## Supplementary material for "Comparative food-web analysis of bluefin tuna spawning habitats in the eastern Indian Ocean and Gulf of Mexico": Code Package: arrow.m

```
function [h,yy,zz] = arrow(varargin)
% ARROW  Draw a line with an arrowhead.
%
%  ARROW(Start,Stop) draws a line with an arrow from Start to Stop (points
%        should be vectors of length 2 or 3, or matrices with 2 or 3
%        columns), and returns the graphics handle of the arrow(s).
%
%  ARROW uses the mouse (click-drag) to create an arrow.
%
%  ARROW DEMO & ARROW DEMO2 show 3-D & 2-D demos of the capabilities of ARROW.
%
%  ARROW may be called with a normal argument list or a property-based list.
%        ARROW(Start,Stop,Length,BaseAngle,TipAngle,Width,Page,CrossDir) is
%        the full normal argument list, where all but the Start and Stop
%        points are optional.  If you need to specify a later argument (e.g.,
%        Page) but want default values of earlier ones (e.g., TipAngle),
%        pass an empty matrix for the earlier ones (e.g., TipAngle=[]).
%
%  ARROW('Property1',PropVal1,'Property2',PropVal2,...) creates arrows with the
%        given properties, using default values for any unspecified or given as
%        'default' or NaN.  Some properties used for line and patch objects are
%        used in a modified fashion, others are passed directly to LINE, PATCH,
%        or SET.  For a detailed properties explanation, call ARROW PROPERTIES.
%
%        Start         The starting points.                     B
%        Stop          The end points.                         /|\           ^
%        Length        Length of the arrowhead in pixels.     /|||\          |
%        BaseAngle     Base angle in degrees (ADE).          //|||\\        L|
%        TipAngle      Tip angle in degrees (ABC).          ///|||\\\       e|
%        Width         Width of the base in pixels.        ////|||\\\\      n|
%        Page          Use hardcopy proportions.          /////|D|\\\\\     g|
%        CrossDir      Vector || to arrowhead plane.     ////  |||  \\\\    t|
%        NormalDir     Vector out of arrowhead plane.   ///    |||    \\\   h|
%        Ends          Which end has an arrowhead.     //<----->||      \\   |
%        ObjectHandles Vector of handles to update.   /   base |||        \  V
%                                                    E    angle||<-------->C
%  ARROW(H,'Prop1',PropVal1,...), where H is a                 |||tipangle
%        vector of handles to previously-created arrows        |||
%        and/or line objects, will update the previously-      |||
%        created arrows according to the current view       -->|A|<-- width
%        and any specified properties, and will convert
%        two-point line objects to corresponding arrows.  ARROW(H) will update
%        the arrows if the current view has changed.  Root, figure, or axes
%        handles included in H are replaced by all descendant Arrow objects.
%
%  A property list can follow any specified normal argument list, e.g.,
%  ARROW([1 2 3],[0 0 0],36,'BaseAngle',60) creates an arrow from (1,2,3) to
%  the origin, with an arrowhead of length 36 pixels and 60-degree base angle.
%
%  Normally, an ARROW is a PATCH object, so any valid PATCH property/value pairs
%  can be passed, e.g., ARROW(Start,Stop,'EdgeColor','r','FaceColor','g').
%  ARROW will use LINE objects when requested by ARROW(...,'Type','line') or,
%  using LINE property/value pairs, ARROW(Start,Stop,'Type','line','Color','b').
%
%  The basic arguments or properties can generally be vectorized to create
%  multiple arrows with the same call.  This is done by passing a property
%  with one row per arrow, or, if all arrows are to have the same property
%  value, just one row may be specified.
%
%  You may want to execute AXIS(AXIS) before calling ARROW so it doesn't change
%  the axes on you; ARROW determines the sizes of arrow components BEFORE the
%  arrow is plotted, so if ARROW changes axis limits, arrows may be malformed.
%
%  This version of ARROW uses features of MATLAB 6.x and is incompatible with
%  earlier MATLAB versions (ARROW for MATLAB 4.2c is available separately);
%  some problems with perspective plots still exist.

% Copyright (c)1995-2016, Dr. Erik A. Johnson <>, 5/25/2016
% http://www.usc.edu/civil_eng/johnsone/

% Revision history:
%    5/25/16  EAJ  Add documentation of 'Type','line'
%                  Add documentation of how to set color
%                  Add 'Color' property (which sets both 'EdgeColor' and 'FaceColor' for patch objects)
%    5/24/16  EAJ  Remove 'EraseMode' in HG2
%    7/16/14  EAJ  R2014b HandleGraphics2 compatibility
%    7/14/14  EAJ  5/20/13 patch extension didn't work right in HG2
%                    so break the arrow along its length instead
%    5/20/13  EAJ  Extend patch line one more segment so EPS/PDF printed versions
%                    have nice rounded tips when the LineWidth is wider
%    2/06/13  EAJ  Add ShortenLength property to shorten length if arrow is short
%    1/24/13  EAJ  Remove some old comments.
%    5/20/09  EAJ  Fix view direction in (3D) demo.
%    6/26/08  EAJ  Replace eval('trycmd','catchcmd') with try, trycmd; catch,
%                    catchcmd; end; -- break's MATLAB 5 compatibility.
%    8/26/03  EAJ  Eliminate OpenGL attempted fix since it didn't fix anyway.
%   11/15/02  EAJ  Accomodate how MATLAB 6.5 handles NaN and logicals
%    7/28/02  EAJ  Tried (but failed) work-around for MATLAB 6.x / OpenGL bug
%                    if zero 'Width' or not double-ended
%   11/10/99  EAJ  Add logical() to eliminate zero index problem in MATLAB 5.3.
%   11/10/99  EAJ  Corrected warning if axis limits changed on multiple axes.
%   11/10/99  EAJ  Update e-mail address.
%    2/10/99  EAJ  Some documentation updating.
%    2/24/98  EAJ  Fixed bug if Start~=Stop but both colinear with viewpoint.
%    8/14/97  EAJ  Added workaround for MATLAB 5.1 scalar logical transpose bug.
%    7/21/97  EAJ  Fixed a few misc bugs.
%    7/14/97  EAJ  Make arrow([],'Prop',...) do nothing (no old handles)
%    6/23/97  EAJ  MATLAB 5 compatible version, release.
%    5/27/97  EAJ  Added Line Arrows back in.  Corrected a few bugs.
%    5/26/97  EAJ  Changed missing Start/Stop to mouse-selected arrows.
%    5/19/97  EAJ  MATLAB 5 compatible version, beta.
%    4/13/97  EAJ  MATLAB 5 compatible version, alpha.
%    1/31/97  EAJ  Fixed bug with multiple arrows and unspecified Z coords.
%   12/05/96  EAJ  Fixed one more bug with log plots and NormalDir specified
%   10/24/96  EAJ  Fixed bug with log plots and NormalDir specified
%   11/13/95  EAJ  Corrected handling for 'reverse' axis directions
%   10/06/95  EAJ  Corrected occasional conflict with SUBPLOT
%    4/24/95  EAJ  A major rewrite.
%    Fall 94  EAJ  Original code.

% Things to be done:
%  - in the arrow_clicks section, prompt by printing to the screen so that
%    the user knows what's going on; also make sure the figure is brought
%    to the front.
%  - segment parsing, computing, and plotting into separate subfunctions
%  - change computing from Xform to Camera paradigms
%     + this will help especially with 3-D perspective plots
%     + if the WarpToFill section works right, remove warning code
%     + when perpsective works properly, remove perspective warning code
%  - add cell property values and struct property name/values (like get/set)
%  - get rid of NaN as the "default" data label
%     + perhaps change userdata to a struct and don't include (or leave
%       empty) the values specified as default; or use a cell containing
%       an empty matrix for a default value
%  - add functionality of GET to retrieve current values of ARROW properties
% 
% New list of things to be done:
%  - rewrite as a graphics or class object that updates itself in real time
%    (but have a 'Static' or 'DoNotUpdate' property to avoid updating)

% Permission is granted to distribute ARROW with the toolboxes for the book
% "Solving Solid Mechanics Problems with MATLAB 5", by F. Golnaraghi et al.
% (Prentice Hall, 1999).

% Permission is granted to Dr. Josef Bigun to distribute ARROW with his
% software to reproduce the figures in his image analysis text.

% global variable initialization
persistent ARROW_PERSP_WARN ARROW_STRETCH_WARN ARROW_AXLIMITS ARROW_AX
if isempty(ARROW_PERSP_WARN  ), ARROW_PERSP_WARN  =1; end;
if isempty(ARROW_STRETCH_WARN), ARROW_STRETCH_WARN=1; end;

% Handle callbacks
if (nargin>0 & isstr(varargin{1}) & strcmp(lower(varargin{1}),'callback')),
	arrow_callback(varargin{2:end}); return;
end;

% Are we doing the demo?
c = sprintf('\n');
if (nargin==1 & isstr(varargin{1})),
	arg1 = lower(varargin{1});
	if strncmp(arg1,'prop',4), arrow_props;
	elseif strncmp(arg1,'demo',4)
		clf reset
		demo_info = arrow_demo;
		if ~strncmp(arg1,'demo2',5),
			hh=arrow_demo3(demo_info);
		else,
			hh=arrow_demo2(demo_info);
		end;
		if (nargout>=1), h=hh; end;
	elseif strncmp(arg1,'fixlimits',3),
		arrow_fixlimits(ARROW_AX,ARROW_AXLIMITS);
		ARROW_AXLIMITS=[]; ARROW_AX=[];
	elseif strncmp(arg1,'help',4),
		disp(help(mfilename));
	else,
		error([upper(mfilename) ' got an unknown single-argument string ''' deblank(arg1) '''.']);
	end;
	return;
end;

% Check # of arguments
if (nargout>3), error([upper(mfilename) ' produces at most 3 output arguments.']); end;

% find first property number
firstprop = nargin+1;
for k=1:length(varargin), if ~isnumeric(varargin{k}) && ~all(ishandle(varargin{k})), firstprop=k; break; end; end; %eaj 5/24/16   for k=1:length(varargin), if ~isnumeric(varargin{k}), firstprop=k; break; end; end;
lastnumeric = firstprop-1;

% check property list
if (firstprop<=nargin),
	for k=firstprop:2:nargin,
		curarg = varargin{k};
		if ~isstr(curarg) | sum(size(curarg)>1)>1,
			error([upper(mfilename) ' requires that a property name be a single string.']);
		end;
	end;
	if (rem(nargin-firstprop,2)~=1),
		error([upper(mfilename) ' requires that the property ''' ...
		       varargin{nargin} ''' be paired with a property value.']);
	end;
end;

% default output
if (nargout>0), h=[]; end;
if (nargout>1), yy=[]; end;
if (nargout>2), zz=[]; end;

% set values to empty matrices
start      = [];
stop       = [];
len        = [];
baseangle  = [];
tipangle   = [];
wid        = [];
page       = [];
crossdir   = [];
ends       = [];
shorten    = [];
ax         = [];
oldh       = [];
ispatch    = [];
defstart      = [NaN NaN NaN];
defstop       = [NaN NaN NaN];
deflen        = 16;
defbaseangle  = 90;
deftipangle   = 16;
defwid        = 0;
defpage       = 0;
defcrossdir   = [NaN NaN NaN];
defends       = 1;
defshorten    = 0;
defoldh       = [];
defispatch    = 1;

% The 'Tag' we'll put on our arrows
ArrowTag = 'Arrow';

% check for oldstyle arguments
if (firstprop==2),
	% assume arg1 is a set of handles
	oldh = varargin{1}(:);
	if isempty(oldh), return; end;
elseif (firstprop>9),
	error([upper(mfilename) ' takes at most 8 non-property arguments.']);
elseif (firstprop>2),
	{start,stop,len,baseangle,tipangle,wid,page,crossdir};
	args = [varargin(1:firstprop-1) cell(1,length(ans)-(firstprop-1))];
	[start,stop,len,baseangle,tipangle,wid,page,crossdir] = deal(args{:});
end;

% parse property pairs
extraprops={};
for k=firstprop:2:nargin,
	prop = varargin{k};
	val  = varargin{k+1};
	prop = [lower(prop(:)') '      '];
	if     strncmp(prop,'start'  ,5),   start      = val;
	elseif strncmp(prop,'stop'   ,4),   stop       = val;
	elseif strncmp(prop,'len'    ,3),   len        = val(:);
	elseif strncmp(prop,'base'   ,4),   baseangle  = val(:);
	elseif strncmp(prop,'tip'    ,3),   tipangle   = val(:);
	elseif strncmp(prop,'wid'    ,3),   wid        = val(:);
	elseif strncmp(prop,'page'   ,4),   page       = val;
	elseif strncmp(prop,'cross'  ,5),   crossdir   = val;
	elseif strncmp(prop,'norm'   ,4),   if (isstr(val)), crossdir=val; else, crossdir=val*sqrt(-1); end;
	elseif strncmp(prop,'end'    ,3),   ends       = val;
	elseif strncmp(prop,'shorten',5),   shorten    = val;
	elseif strncmp(prop,'object' ,6),   oldh       = val(:);
	elseif strncmp(prop,'handle' ,6),   oldh       = val(:);
	elseif strncmp(prop,'type'   ,4),   ispatch    = val;
	elseif strncmp(prop,'userd'  ,5),   %ignore it
	else,
		% make sure it is a valid patch or line property
		try
			get(0,['DefaultPatch' varargin{k}]);
		catch
			errstr = lasterr;
			try
				get(0,['DefaultLine' varargin{k}]);
			catch
				errstr(1:max(find(errstr==char(13)|errstr==char(10)))) = '';
				error([upper(mfilename) ' got ' errstr]);
			end
		end;
		extraprops={extraprops{:},varargin{k},val};
	end;
end;

% Check if we got 'default' values
start     = arrow_defcheck(start    ,defstart    ,'Start'        );
stop      = arrow_defcheck(stop     ,defstop     ,'Stop'         );
len       = arrow_defcheck(len      ,deflen      ,'Length'       );
baseangle = arrow_defcheck(baseangle,defbaseangle,'BaseAngle'    );
tipangle  = arrow_defcheck(tipangle ,deftipangle ,'TipAngle'     );
wid       = arrow_defcheck(wid      ,defwid      ,'Width'        );
crossdir  = arrow_defcheck(crossdir ,defcrossdir ,'CrossDir'     );
page      = arrow_defcheck(page     ,defpage     ,'Page'         );
ends      = arrow_defcheck(ends     ,defends     ,''             );
shorten   = arrow_defcheck(shorten  ,defshorten  ,''             );
oldh      = arrow_defcheck(oldh     ,[]          ,'ObjectHandles');
ispatch   = arrow_defcheck(ispatch  ,defispatch  ,''             );

% check transpose on arguments
[m,n]=size(start   );   if any(m==[2 3])&(n==1|n>3),   start    = start';      end;
[m,n]=size(stop    );   if any(m==[2 3])&(n==1|n>3),   stop     = stop';       end;
[m,n]=size(crossdir);   if any(m==[2 3])&(n==1|n>3),   crossdir = crossdir';   end;

% convert strings to numbers
if ~isempty(ends) & isstr(ends),
	endsorig = ends;
	[m,n] = size(ends);
	col = lower([ends(:,1:min(3,n)) ones(m,max(0,3-n))*' ']);
	ends = NaN*ones(m,1);
	oo = ones(1,m);
	ii=find(all(col'==['non']'*oo)'); if ~isempty(ii), ends(ii)=ones(length(ii),1)*0; end;
	ii=find(all(col'==['sto']'*oo)'); if ~isempty(ii), ends(ii)=ones(length(ii),1)*1; end;
	ii=find(all(col'==['sta']'*oo)'); if ~isempty(ii), ends(ii)=ones(length(ii),1)*2; end;
	ii=find(all(col'==['bot']'*oo)'); if ~isempty(ii), ends(ii)=ones(length(ii),1)*3; end;
	if any(isnan(ends)),
		ii = min(find(isnan(ends)));
		error([upper(mfilename) ' does not recognize ''' deblank(endsorig(ii,:)) ''' as a valid ''Ends'' value.']);
	end;
else,
	ends = ends(:);
end;
if ~isempty(ispatch) & isstr(ispatch),
	col = lower(ispatch(:,1));
	patchchar='p'; linechar='l'; defchar=' ';
	mask = col~=patchchar & col~=linechar & col~=defchar;
	if any(mask),
		error([upper(mfilename) ' does not recognize ''' deblank(ispatch(min(find(mask)),:)) ''' as a valid ''Type'' value.']);
	end;
	ispatch = (col==patchchar)*1 + (col==linechar)*0 + (col==defchar)*defispatch;
else,
	ispatch = ispatch(:);
end;
oldh = oldh(:);

% check object handles
if ~all(ishandle(oldh)), error([upper(mfilename) ' got invalid object handles.']); end;

% expand root, figure, and axes handles
if ~isempty(oldh),
	ohtype = get(oldh,'Type');
	mask = strcmp(ohtype,'root') | strcmp(ohtype,'figure') | strcmp(ohtype,'axes');
	if any(mask),
		oldh = num2cell(oldh);
		for ii=find(mask)',
			oldh(ii) = {findobj(oldh{ii},'Tag',ArrowTag)};
		end;
		oldh = cat(1,oldh{:});
		if isempty(oldh), return; end; % no arrows to modify, so just leave
	end;
end;

% largest argument length
[mstart,junk]=size(start); [mstop,junk]=size(stop); [mcrossdir,junk]=size(crossdir);
argsizes = [length(oldh) mstart mstop                              ...
            length(len) length(baseangle) length(tipangle)         ...
			length(wid) length(page) mcrossdir length(ends) length(shorten)];
args=['length(ObjectHandle)  '; ...
      '#rows(Start)          '; ...
      '#rows(Stop)           '; ...
      'length(Length)        '; ...
      'length(BaseAngle)     '; ...
      'length(TipAngle)      '; ...
      'length(Width)         '; ...
      'length(Page)          '; ...
      '#rows(CrossDir)       '; ...
      '#rows(Ends)           '; ...
      'length(ShortenLength) '];
if (any(imag(crossdir(:))~=0)),
	args(9,:) = '#rows(NormalDir)      ';
end;
if isempty(oldh),
	narrows = max(argsizes);
else,
	narrows = length(oldh);
end;
if (narrows<=0), narrows=1; end;

% Check size of arguments
ii = find((argsizes~=0)&(argsizes~=1)&(argsizes~=narrows));
if ~isempty(ii),
	s = args(ii',:);
	while ((size(s,2)>1)&((abs(s(:,size(s,2)))==0)|(abs(s(:,size(s,2)))==abs(' ')))),
		s = s(:,1:size(s,2)-1);
	end;
	s = [ones(length(ii),1)*[upper(mfilename) ' requires that  '] s ...
	     ones(length(ii),1)*['  equal the # of arrows (' num2str(narrows) ').' c]];
	s = s';
	s = s(:)';
	s = s(1:length(s)-1);
	error(setstr(s));
end;

% check element length in Start, Stop, and CrossDir
if ~isempty(start),
	[m,n] = size(start);
	if (n==2),
		start = [start NaN*ones(m,1)];
	elseif (n~=3),
		error([upper(mfilename) ' requires 2- or 3-element Start points.']);
	end;
end;
if ~isempty(stop),
	[m,n] = size(stop);
	if (n==2),
		stop = [stop NaN*ones(m,1)];
	elseif (n~=3),
		error([upper(mfilename) ' requires 2- or 3-element Stop points.']);
	end;
end;
if ~isempty(crossdir),
	[m,n] = size(crossdir);
	if (n<3),
		crossdir = [crossdir NaN*ones(m,3-n)];
	elseif (n~=3),
		if (all(imag(crossdir(:))==0)),
			error([upper(mfilename) ' requires 2- or 3-element CrossDir vectors.']);
		else,
			error([upper(mfilename) ' requires 2- or 3-element NormalDir vectors.']);
		end;
	end;
end;

% fill empty arguments
if isempty(start     ),   start      = [Inf Inf Inf];      end;
if isempty(stop      ),   stop       = [Inf Inf Inf];      end;
if isempty(len       ),   len        = Inf;                end;
if isempty(baseangle ),   baseangle  = Inf;                end;
if isempty(tipangle  ),   tipangle   = Inf;                end;
if isempty(wid       ),   wid        = Inf;                end;
if isempty(page      ),   page       = Inf;                end;
if isempty(crossdir  ),   crossdir   = [Inf Inf Inf];      end;
if isempty(ends      ),   ends       = Inf;                end;
if isempty(shorten   ),   shorten    = Inf;                end;
if isempty(ispatch   ),   ispatch    = Inf;                end;

% expand single-column arguments
o = ones(narrows,1);
if (size(start     ,1)==1),   start      = o * start     ;   end;
if (size(stop      ,1)==1),   stop       = o * stop      ;   end;
if (length(len       )==1),   len        = o * len       ;   end;
if (length(baseangle )==1),   baseangle  = o * baseangle ;   end;
if (length(tipangle  )==1),   tipangle   = o * tipangle  ;   end;
if (length(wid       )==1),   wid        = o * wid       ;   end;
if (length(page      )==1),   page       = o * page      ;   end;
if (size(crossdir  ,1)==1),   crossdir   = o * crossdir  ;   end;
if (length(ends      )==1),   ends       = o * ends      ;   end;
if (length(shorten   )==1),   shorten    = o * shorten   ;   end;
if (length(ispatch   )==1),   ispatch    = o * ispatch   ;   end;
ax = repmat(gca,narrows,1);   %eaj 7/16/14  ax=gca; if ~isnumeric(ax), ax=double(ax); end; ax=o*ax;

% if we've got handles, get the defaults from the handles
if ~isempty(oldh),
	for k=1:narrows,
		oh = oldh(k);
		ud = get(oh,'UserData');
		ax(k) = get(oh,'Parent');   %eaj 7/16/14  get(oh,'Parent'); if ~isnumeric(ans), double(ans); end; ax(k)=ans;
		ohtype = get(oh,'Type');
		if strcmp(get(oh,'Tag'),ArrowTag), % if it's an arrow already
			if isinf(ispatch(k)), ispatch(k)=strcmp(ohtype,'patch'); end;
			% arrow UserData format: [start' stop' len base tip wid page crossdir' ends shorten]
			start0 = ud(1:3);
			stop0  = ud(4:6);
			if (isinf(len(k))),           len(k)        = ud( 7);   end;
			if (isinf(baseangle(k))),     baseangle(k)  = ud( 8);   end;
			if (isinf(tipangle(k))),      tipangle(k)   = ud( 9);   end;
			if (isinf(wid(k))),           wid(k)        = ud(10);   end;
			if (isinf(page(k))),          page(k)       = ud(11);   end;
			if (isinf(crossdir(k,1))),    crossdir(k,1) = ud(12);   end;
			if (isinf(crossdir(k,2))),    crossdir(k,2) = ud(13);   end;
			if (isinf(crossdir(k,3))),    crossdir(k,3) = ud(14);   end;
			if (isinf(ends(k))),          ends(k)       = ud(15);   end;
			if (isinf(shorten(k))),       shorten(k)    = ud(16);   end;
		elseif strcmp(ohtype,'line')|strcmp(ohtype,'patch'), % it's a non-arrow line or patch
			convLineToPatch = 1; %set to make arrow patches when converting from lines.
			if isinf(ispatch(k)), ispatch(k)=convLineToPatch|strcmp(ohtype,'patch'); end;
			x=get(oh,'XData');  x=x(~isnan(x(:)));  if isempty(x), x=NaN; end;
			y=get(oh,'YData');  y=y(~isnan(y(:)));  if isempty(y), y=NaN; end;
			z=get(oh,'ZData');  z=z(~isnan(z(:)));  if isempty(z), z=NaN; end;
			start0 = [x(1)   y(1)   z(1)  ];
			stop0  = [x(end) y(end) z(end)];
		else,
			error([upper(mfilename) ' cannot convert ' ohtype ' objects.']);
		end;
		ii=find(isinf(start(k,:)));  if ~isempty(ii),  start(k,ii)=start0(ii);  end;
		ii=find(isinf(stop( k,:)));  if ~isempty(ii),  stop( k,ii)=stop0( ii);  end;
	end;
end;

% convert Inf's to NaN's
start(     isinf(start    )) = NaN;
stop(      isinf(stop     )) = NaN;
len(       isinf(len      )) = NaN;
baseangle( isinf(baseangle)) = NaN;
tipangle(  isinf(tipangle )) = NaN;
wid(       isinf(wid      )) = NaN;
page(      isinf(page     )) = NaN;
crossdir(  isinf(crossdir )) = NaN;
ends(      isinf(ends     )) = NaN;
shorten(   isinf(shorten  )) = NaN;
ispatch(   isinf(ispatch  )) = NaN;

% set up the UserData data (here so not corrupted by log10's and such)
ud = [start stop len baseangle tipangle wid page crossdir ends shorten];

% Set Page defaults
page = ~isnan(page) & trueornan(page);

% Get axes limits, range, min; correct for aspect ratio and log scale
axm       = zeros(3,narrows);
axr       = zeros(3,narrows);
axrev     = zeros(3,narrows);
ap        = zeros(2,narrows);
xyzlog    = zeros(3,narrows);
limmin    = zeros(2,narrows);
limrange  = zeros(2,narrows);
oldaxlims = zeros(6,narrows);
oneax = all(ax==ax(1));
if (oneax),
	T    = zeros(4,4);
	invT = zeros(4,4);
else,
	T    = zeros(16,narrows);
	invT = zeros(16,narrows);
end;
axnotdone = true(size(ax));
while (any(axnotdone)),
	ii = find(axnotdone,1);
	curax = ax(ii);
	curpage = page(ii);
	% get axes limits and aspect ratio
	axl = [get(curax,'XLim'); get(curax,'YLim'); get(curax,'ZLim')];
	ax==curax; oldaxlims(:,ans)=repmat(reshape(axl',[],1),1,sum(ans));
	% get axes size in pixels (points)
	u = get(curax,'Units');
	axposoldunits = get(curax,'Position');
	really_curpage = curpage & strcmp(u,'normalized');
	if (really_curpage),
		curfig = get(curax,'Parent');
		pu = get(curfig,'PaperUnits');
		set(curfig,'PaperUnits','points');
		pp = get(curfig,'PaperPosition');
		set(curfig,'PaperUnits',pu);
		set(curax,'Units','pixels');
		curapscreen = get(curax,'Position');
		set(curax,'Units','normalized');
		curap = pp.*get(curax,'Position');
	else,
		set(curax,'Units','pixels');
		curapscreen = get(curax,'Position');
		curap = curapscreen;
	end;
	set(curax,'Units',u);
	set(curax,'Position',axposoldunits);
	% handle non-stretched axes position
	str_stretch = { 'DataAspectRatioMode'    ; ...
	                'PlotBoxAspectRatioMode' ; ...
	                'CameraViewAngleMode'      };
	str_camera  = { 'CameraPositionMode'  ; ...
	                'CameraTargetMode'    ; ...
	                'CameraViewAngleMode' ; ...
	                'CameraUpVectorMode'    };
	notstretched = strcmp(get(curax,str_stretch),'manual');
	manualcamera = strcmp(get(curax,str_camera),'manual');
	if ~arrow_WarpToFill(notstretched,manualcamera,curax),
		% give a warning that this has not been thoroughly tested
		if 0 & ARROW_STRETCH_WARN,
			ARROW_STRETCH_WARN = 0;
			strs = {str_stretch{1:2},str_camera{:}};
			strs = [char(ones(length(strs),1)*sprintf('\n    ')) char(strs)]';
			warning([upper(mfilename) ' may not yet work quite right ' ...
			         'if any of the following are ''manual'':' strs(:).']);
		end;
		% find the true pixel size of the actual axes
		texttmp = text(axl(1,[1 2 2 1 1 2 2 1]), ...
		               axl(2,[1 1 2 2 1 1 2 2]), ...
		               axl(3,[1 1 1 1 2 2 2 2]),'');
		set(texttmp,'Units','points');
		textpos = get(texttmp,'Position');
		delete(texttmp);
		textpos = cat(1,textpos{:});
		textpos = max(textpos(:,1:2)) - min(textpos(:,1:2));
		% adjust the axes position
		if (really_curpage),
			% adjust to printed size
			textpos = textpos * min(curap(3:4)./textpos);
			curap = [curap(1:2)+(curap(3:4)-textpos)/2 textpos];
		else,
			% adjust for pixel roundoff
			textpos = textpos * min(curapscreen(3:4)./textpos);
			curap = [curap(1:2)+(curap(3:4)-textpos)/2 textpos];
		end;
	end;
	if ARROW_PERSP_WARN & ~strcmp(get(curax,'Projection'),'orthographic'),
		ARROW_PERSP_WARN = 0;
		warning([upper(mfilename) ' does not yet work right for 3-D perspective projection.']);
	end;
	% adjust limits for log scale on axes
	curxyzlog = strcmp(get(curax,{'XScale' 'YScale' 'ZScale'})','log');
	if (any(curxyzlog)),
		ii = find([curxyzlog;curxyzlog]);
		if (any(axl(ii)<=0)),
			error([upper(mfilename) ' does not support non-positive limits on log-scaled axes.']);
		else,
			axl(ii) = log10(axl(ii));
		end;
	end;
	% correct for 'reverse' direction on axes;
	curreverse = strcmp(get(curax,{'XDir' 'YDir' 'ZDir'})','reverse');
	ii = find(curreverse);
	if ~isempty(ii),
		axl(ii,[1 2])=-axl(ii,[2 1]);
	end;
	% compute the range of 2-D values
	try, curT=get(curax,'Xform'); catch, num2cell(get(curax,'View')); curT=viewmtx(ans{:}); end;
	lim = curT*[0 1 0 1 0 1 0 1;0 0 1 1 0 0 1 1;0 0 0 0 1 1 1 1;1 1 1 1 1 1 1 1];
	lim = lim(1:2,:)./([1;1]*lim(4,:));
	curlimmin = min(lim')';
	curlimrange = max(lim')' - curlimmin;
	curinvT = inv(curT);
	if (~oneax),
		curT = curT.';
		curinvT = curinvT.';
		curT = curT(:);
		curinvT = curinvT(:);
	end;
	% check which arrows to which cur corresponds
	ii = find((ax==curax)&(page==curpage));
	oo = ones(1,length(ii));
	axr(:,ii)      = diff(axl')' * oo;
	axm(:,ii)      = axl(:,1)    * oo;
	axrev(:,ii)    = curreverse  * oo;
	ap(:,ii)       = curap(3:4)' * oo;
	xyzlog(:,ii)   = curxyzlog   * oo;
	limmin(:,ii)   = curlimmin   * oo;
	limrange(:,ii) = curlimrange * oo;
	if (oneax),
		T    = curT;
		invT = curinvT;
	else,
		T(:,ii)    = curT    * oo;
		invT(:,ii) = curinvT * oo;
	end;
	axnotdone(ii) = zeros(1,length(ii));
end;

% correct for log scales
curxyzlog = xyzlog.';
ii = find(curxyzlog(:));
if ~isempty(ii),
	start(   ii) = real(log10(start(   ii)));
	stop(    ii) = real(log10(stop(    ii)));
	if (all(imag(crossdir)==0)), % pulled (ii) subscript on crossdir, 12/5/96 eaj
		crossdir(ii) = real(log10(crossdir(ii)));
	end;
end;

% correct for reverse directions
ii = find(axrev.');
if ~isempty(ii),
	start(   ii) = -start(   ii);
	stop(    ii) = -stop(    ii);
	crossdir(ii) = -crossdir(ii);
end;

% transpose start/stop values
start     = start.';
stop      = stop.';

% take care of defaults, page was done above
ii=find(isnan(start(:)       ));  if ~isempty(ii),  start(ii)       = axm(ii)+axr(ii)/2;                end;
ii=find(isnan(stop(:)        ));  if ~isempty(ii),  stop(ii)        = axm(ii)+axr(ii)/2;                end;
ii=find(isnan(crossdir(:)    ));  if ~isempty(ii),  crossdir(ii)    = zeros(length(ii),1);              end;
ii=find(isnan(len            ));  if ~isempty(ii),  len(ii)         = ones(length(ii),1)*deflen;        end;
ii=find(isnan(baseangle      ));  if ~isempty(ii),  baseangle(ii)   = ones(length(ii),1)*defbaseangle;  end;
ii=find(isnan(tipangle       ));  if ~isempty(ii),  tipangle(ii)    = ones(length(ii),1)*deftipangle;   end;
ii=find(isnan(wid            ));  if ~isempty(ii),  wid(ii)         = ones(length(ii),1)*defwid;        end;
ii=find(isnan(ends           ));  if ~isempty(ii),  ends(ii)        = ones(length(ii),1)*defends;       end;
ii=find(isnan(shorten        ));  if ~isempty(ii),  shorten(ii)     = ones(length(ii),1)*defshorten;    end;

% transpose rest of values
len       = len.';
baseangle = baseangle.';
tipangle  = tipangle.';
wid       = wid.';
page      = page.';
crossdir  = crossdir.';
ends      = ends.';
shorten   = shorten.';
ax        = ax.';

% given x, a 3xN matrix of points in 3-space;
% want to convert to X, the corresponding 4xN 2-space matrix
%
%   tmp1=[(x-axm)./axr; ones(1,size(x,1))];
%   if (oneax), X=T*tmp1;
%   else, tmp1=[tmp1;tmp1;tmp1;tmp1]; tmp1=T.*tmp1;
%         tmp2=zeros(4,4*N); tmp2(:)=tmp1(:);
%         X=zeros(4,N); X(:)=sum(tmp2)'; end;
%   X = X ./ (ones(4,1)*X(4,:));

% for all points with start==stop, start=stop-(verysmallvalue)*(up-direction);
ii = find(all(start==stop));
if ~isempty(ii),
	% find an arrowdir vertical on screen and perpendicular to viewer
	%	transform to 2-D
		tmp1 = [(stop(:,ii)-axm(:,ii))./axr(:,ii);ones(1,length(ii))];
		if (oneax), twoD=T*tmp1;
		else, tmp1=[tmp1;tmp1;tmp1;tmp1]; tmp1=T(:,ii).*tmp1;
		      tmp2=zeros(4,4*length(ii)); tmp2(:)=tmp1(:);
		      twoD=zeros(4,length(ii)); twoD(:)=sum(tmp2)'; end;
		twoD=twoD./(ones(4,1)*twoD(4,:));
	%	move the start point down just slightly
		tmp1 = twoD + [0;-1/1000;0;0]*(limrange(2,ii)./ap(2,ii));
	%	transform back to 3-D
		if (oneax), threeD=invT*tmp1;
		else, tmp1=[tmp1;tmp1;tmp1;tmp1]; tmp1=invT(:,ii).*tmp1;
		      tmp2=zeros(4,4*length(ii)); tmp2(:)=tmp1(:);
		      threeD=zeros(4,length(ii)); threeD(:)=sum(tmp2)'; end;
		start(:,ii) = (threeD(1:3,:)./(ones(3,1)*threeD(4,:))).*axr(:,ii)+axm(:,ii);
end;

% compute along-arrow points
%	transform Start points
	tmp1=[(start-axm)./axr;ones(1,narrows)];
	if (oneax), X0=T*tmp1;
	else, tmp1=[tmp1;tmp1;tmp1;tmp1]; tmp1=T.*tmp1;
	      tmp2=zeros(4,4*narrows); tmp2(:)=tmp1(:);
	      X0=zeros(4,narrows); X0(:)=sum(tmp2)'; end;
	X0=X0./(ones(4,1)*X0(4,:));
%	transform Stop points
	tmp1=[(stop-axm)./axr;ones(1,narrows)];
	if (oneax), Xf=T*tmp1;
	else, tmp1=[tmp1;tmp1;tmp1;tmp1]; tmp1=T.*tmp1;
	      tmp2=zeros(4,4*narrows); tmp2(:)=tmp1(:);
	      Xf=zeros(4,narrows); Xf(:)=sum(tmp2)'; end;
	Xf=Xf./(ones(4,1)*Xf(4,:));
%	compute pixel distance between points
	D = sqrt(sum(((Xf(1:2,:)-X0(1:2,:)).*(ap./limrange)).^2));
	D = D + (D==0);  %eaj new 2/24/98
%       shorten the length if requested % added 2/6/2013
	numends = (ends==1) + (ends==2) + 2*(ends==3);
	mask = shorten & D<len.*numends;
	len(mask) = D(mask) ./ numends(mask);
%	compute and modify along-arrow distances
	len1 = len;
	len2 = len - (len.*tan(tipangle/180*pi)-wid/2).*tan((90-baseangle)/180*pi);
	slen0 = zeros(1,narrows);
	slen1 = len1 .* ((ends==2)|(ends==3));
	slen2 = len2 .* ((ends==2)|(ends==3));
	len0 = zeros(1,narrows);
	len1  = len1 .* ((ends==1)|(ends==3));
	len2  = len2 .* ((ends==1)|(ends==3));
	%	for no start arrowhead
		ii=find((ends==1)&(D<len2));
		if ~isempty(ii),
			slen0(ii) = D(ii)-len2(ii);
		end;
	%	for no end arrowhead
		ii=find((ends==2)&(D<slen2));
		if ~isempty(ii),
			len0(ii) = D(ii)-slen2(ii);
		end;
	len1  = len1  + len0;
	len2  = len2  + len0;
	slen1 = slen1 + slen0;
	slen2 = slen2 + slen0;
 	% note:  the division by D below will probably not be accurate if both
 	%        of the following are true:
 	%           1. the ratio of the line length to the arrowhead
 	%              length is large
 	%           2. the view is highly perspective.
%	compute stoppoints
	tmp1=X0.*(ones(4,1)*(len0./D))+Xf.*(ones(4,1)*(1-len0./D));
	if (oneax), tmp3=invT*tmp1;
	else, tmp1=[tmp1;tmp1;tmp1;tmp1]; tmp1=invT.*tmp1;
	      tmp2=zeros(4,4*narrows); tmp2(:)=tmp1(:);
	      tmp3=zeros(4,narrows); tmp3(:)=sum(tmp2)'; end;
	stoppoint = tmp3(1:3,:)./(ones(3,1)*tmp3(4,:)).*axr+axm;
%	compute tippoints
	tmp1=X0.*(ones(4,1)*(len1./D))+Xf.*(ones(4,1)*(1-len1./D));
	if (oneax), tmp3=invT*tmp1;
	else, tmp1=[tmp1;tmp1;tmp1;tmp1]; tmp1=invT.*tmp1;
	      tmp2=zeros(4,4*narrows); tmp2(:)=tmp1(:);
	      tmp3=zeros(4,narrows); tmp3(:)=sum(tmp2)'; end;
	tippoint = tmp3(1:3,:)./(ones(3,1)*tmp3(4,:)).*axr+axm;
%	compute basepoints
	tmp1=X0.*(ones(4,1)*(len2./D))+Xf.*(ones(4,1)*(1-len2./D));
	if (oneax), tmp3=invT*tmp1;
	else, tmp1=[tmp1;tmp1;tmp1;tmp1]; tmp1=invT.*tmp1;
	      tmp2=zeros(4,4*narrows); tmp2(:)=tmp1(:);
	      tmp3=zeros(4,narrows); tmp3(:)=sum(tmp2)'; end;
	basepoint = tmp3(1:3,:)./(ones(3,1)*tmp3(4,:)).*axr+axm;
%	compute startpoints
	tmp1=X0.*(ones(4,1)*(1-slen0./D))+Xf.*(ones(4,1)*(slen0./D));
	if (oneax), tmp3=invT*tmp1;
	else, tmp1=[tmp1;tmp1;tmp1;tmp1]; tmp1=invT.*tmp1;
	      tmp2=zeros(4,4*narrows); tmp2(:)=tmp1(:);
	      tmp3=zeros(4,narrows); tmp3(:)=sum(tmp2)'; end;
	startpoint = tmp3(1:3,:)./(ones(3,1)*tmp3(4,:)).*axr+axm;
%	compute stippoints
	tmp1=X0.*(ones(4,1)*(1-slen1./D))+Xf.*(ones(4,1)*(slen1./D));
	if (oneax), tmp3=invT*tmp1;
	else, tmp1=[tmp1;tmp1;tmp1;tmp1]; tmp1=invT.*tmp1;
	      tmp2=zeros(4,4*narrows); tmp2(:)=tmp1(:);
	      tmp3=zeros(4,narrows); tmp3(:)=sum(tmp2)'; end;
	stippoint = tmp3(1:3,:)./(ones(3,1)*tmp3(4,:)).*axr+axm;
%	compute sbasepoints
	tmp1=X0.*(ones(4,1)*(1-slen2./D))+Xf.*(ones(4,1)*(slen2./D));
	if (oneax), tmp3=invT*tmp1;
	else, tmp1=[tmp1;tmp1;tmp1;tmp1]; tmp1=invT.*tmp1;
	      tmp2=zeros(4,4*narrows); tmp2(:)=tmp1(:);
	      tmp3=zeros(4,narrows); tmp3(:)=sum(tmp2)'; end;
	sbasepoint = tmp3(1:3,:)./(ones(3,1)*tmp3(4,:)).*axr+axm;

% compute cross-arrow directions for arrows with NormalDir specified
if (any(imag(crossdir(:))~=0)),
	ii = find(any(imag(crossdir)~=0));
	crossdir(:,ii) = cross((stop(:,ii)-start(:,ii))./axr(:,ii), ...
	                       imag(crossdir(:,ii))).*axr(:,ii);
end;

% compute cross-arrow directions
basecross  = crossdir + basepoint;
tipcross   = crossdir + tippoint;
sbasecross = crossdir + sbasepoint;
stipcross  = crossdir + stippoint;
ii = find(all(crossdir==0)|any(isnan(crossdir)));
if ~isempty(ii),
	numii = length(ii);
	%	transform start points
		tmp1 = [basepoint(:,ii) tippoint(:,ii) sbasepoint(:,ii) stippoint(:,ii)];
		tmp1 = (tmp1-axm(:,[ii ii ii ii])) ./ axr(:,[ii ii ii ii]);
		tmp1 = [tmp1; ones(1,4*numii)];
		if (oneax), X0=T*tmp1;
		else, tmp1=[tmp1;tmp1;tmp1;tmp1]; tmp1=T(:,[ii ii ii ii]).*tmp1;
		      tmp2=zeros(4,16*numii); tmp2(:)=tmp1(:);
		      X0=zeros(4,4*numii); X0(:)=sum(tmp2)'; end;
		X0=X0./(ones(4,1)*X0(4,:));
	%	transform stop points
		tmp1 = [(2*stop(:,ii)-start(:,ii)-axm(:,ii))./axr(:,ii);ones(1,numii)];
		tmp1 = [tmp1 tmp1 tmp1 tmp1];
		if (oneax), Xf=T*tmp1;
		else, tmp1=[tmp1;tmp1;tmp1;tmp1]; tmp1=T(:,[ii ii ii ii]).*tmp1;
		      tmp2=zeros(4,16*numii); tmp2(:)=tmp1(:);
		      Xf=zeros(4,4*numii); Xf(:)=sum(tmp2)'; end;
		Xf=Xf./(ones(4,1)*Xf(4,:));
	%	compute perpendicular directions
		pixfact = ((limrange(1,ii)./limrange(2,ii)).*(ap(2,ii)./ap(1,ii))).^2;
		pixfact = [pixfact pixfact pixfact pixfact];
		pixfact = [pixfact;1./pixfact];
		[dummyval,jj] = max(abs(Xf(1:2,:)-X0(1:2,:)));
		jj1 = ((1:4)'*ones(1,length(jj))==ones(4,1)*jj);
		jj2 = ((1:4)'*ones(1,length(jj))==ones(4,1)*(3-jj));
		jj3 = jj1(1:2,:);
		Xf(jj1)=Xf(jj1)+(Xf(jj1)-X0(jj1)==0); %eaj new 2/24/98
		Xp = X0;
		Xp(jj2) = X0(jj2) + ones(sum(jj2(:)),1);
		Xp(jj1) = X0(jj1) - (Xf(jj2)-X0(jj2))./(Xf(jj1)-X0(jj1)) .* pixfact(jj3);
	%	inverse transform the cross points
		if (oneax), Xp=invT*Xp;
		else, tmp1=[Xp;Xp;Xp;Xp]; tmp1=invT(:,[ii ii ii ii]).*tmp1;
		      tmp2=zeros(4,16*numii); tmp2(:)=tmp1(:);
		      Xp=zeros(4,4*numii); Xp(:)=sum(tmp2)'; end;
		Xp=(Xp(1:3,:)./(ones(3,1)*Xp(4,:))).*axr(:,[ii ii ii ii])+axm(:,[ii ii ii ii]);
		basecross(:,ii)  = Xp(:,0*numii+(1:numii));
		tipcross(:,ii)   = Xp(:,1*numii+(1:numii));
		sbasecross(:,ii) = Xp(:,2*numii+(1:numii));
		stipcross(:,ii)  = Xp(:,3*numii+(1:numii));
end;

% compute all points
%	compute start points
	axm11 = [axm axm axm axm axm axm axm axm axm axm axm];
	axr11 = [axr axr axr axr axr axr axr axr axr axr axr];
	st = [stoppoint tippoint basepoint sbasepoint stippoint startpoint stippoint sbasepoint basepoint tippoint stoppoint];
	tmp1 = (st - axm11) ./ axr11;
	tmp1 = [tmp1; ones(1,size(tmp1,2))];
	if (oneax), X0=T*tmp1;
	else, tmp1=[tmp1;tmp1;tmp1;tmp1]; tmp1=[T T T T T T T T T T T].*tmp1;
	      tmp2=zeros(4,44*narrows); tmp2(:)=tmp1(:);
	      X0=zeros(4,11*narrows); X0(:)=sum(tmp2)'; end;
	X0=X0./(ones(4,1)*X0(4,:));
%	compute stop points
	tmp1 = ([start tipcross basecross sbasecross stipcross stop stipcross sbasecross basecross tipcross start] ...
	     - axm11) ./ axr11;
	tmp1 = [tmp1; ones(1,size(tmp1,2))];
	if (oneax), Xf=T*tmp1;
	else, tmp1=[tmp1;tmp1;tmp1;tmp1]; tmp1=[T T T T T T T T T T T].*tmp1;
	      tmp2=zeros(4,44*narrows); tmp2(:)=tmp1(:);
	      Xf=zeros(4,11*narrows); Xf(:)=sum(tmp2)'; end;
	Xf=Xf./(ones(4,1)*Xf(4,:));
%	compute lengths
	len0  = len.*((ends==1)|(ends==3)).*tan(tipangle/180*pi);
	slen0 = len.*((ends==2)|(ends==3)).*tan(tipangle/180*pi);
	le = [zeros(1,narrows) len0 wid/2 wid/2 slen0 zeros(1,narrows) -slen0 -wid/2 -wid/2 -len0 zeros(1,narrows)];
	aprange = ap./limrange;
	aprange = [aprange aprange aprange aprange aprange aprange aprange aprange aprange aprange aprange];
	D = sqrt(sum(((Xf(1:2,:)-X0(1:2,:)).*aprange).^2));
	Dii=find(D==0); if ~isempty(Dii), D=D+(D==0); le(Dii)=zeros(1,length(Dii)); end; %should fix DivideByZero warnings
	tmp1 = X0.*(ones(4,1)*(1-le./D)) + Xf.*(ones(4,1)*(le./D));
%	inverse transform
	if (oneax), tmp3=invT*tmp1;
	else, tmp1=[tmp1;tmp1;tmp1;tmp1]; tmp1=[invT invT invT invT invT invT invT invT invT invT invT].*tmp1;
	      tmp2=zeros(4,44*narrows); tmp2(:)=tmp1(:);
	      tmp3=zeros(4,11*narrows); tmp3(:)=sum(tmp2)'; end;
	pts = tmp3(1:3,:)./(ones(3,1)*tmp3(4,:)) .* axr11 + axm11;

% correct for ones where the crossdir was specified
ii = find(~(all(crossdir==0)|any(isnan(crossdir))));
if ~isempty(ii),
	D1 = [pts(:,1*narrows+ii)-pts(:,9*narrows+ii) ...
	      pts(:,2*narrows+ii)-pts(:,8*narrows+ii) ...
	      pts(:,3*narrows+ii)-pts(:,7*narrows+ii) ...
	      pts(:,4*narrows+ii)-pts(:,6*narrows+ii) ...
	      pts(:,6*narrows+ii)-pts(:,4*narrows+ii) ...
	      pts(:,7*narrows+ii)-pts(:,3*narrows+ii) ...
	      pts(:,8*narrows+ii)-pts(:,2*narrows+ii) ...
	      pts(:,9*narrows+ii)-pts(:,1*narrows+ii)]/2;
	ii = ii'*ones(1,8) + ones(length(ii),1)*[1:4 6:9]*narrows;
	ii = ii(:)';
	pts(:,ii) = st(:,ii) + D1;
end;


% readjust for reverse directions
iicols=(1:narrows)'; iicols=iicols(:,ones(1,11)); iicols=iicols(:).';
tmp1=axrev(:,iicols);
ii = find(tmp1(:)); if ~isempty(ii), pts(ii)=-pts(ii); end;

% change from starting/ending at the stop point to doing it at the midpoint %eaj 7/14/2014
(pts(:,2*narrows+1:3*narrows)+pts(:,3*narrows+1:4*narrows))/2;              %eaj 7/14/2014
pts = [ans pts(:,[3*narrows+1:end narrows+1:3*narrows]) ans];               %eaj 7/14/2014

% readjust for log scale on axes
tmp1=xyzlog(:,iicols);
ii = find(tmp1(:)); if ~isempty(ii), pts(ii)=10.^pts(ii); end;

% compute the x,y,z coordinates of the patches;
ii = narrows*(0:size(pts,2)/narrows-1)'*ones(1,narrows) + ones(size(pts,2)/narrows,1)*(1:narrows);
ii = ii(:)';
x = zeros(size(pts,2)/narrows,narrows);
y = zeros(size(pts,2)/narrows,narrows);
z = zeros(size(pts,2)/narrows,narrows);
x(:) = pts(1,ii)';
y(:) = pts(2,ii)';
z(:) = pts(3,ii)';

% do the output
if (nargout<=1),
%	% create or modify the patches
	newpatch = trueornan(ispatch) & (isempty(oldh)|~strcmp(get(oldh,'Type'),'patch'));
	newline = ~trueornan(ispatch) & (isempty(oldh)|~strcmp(get(oldh,'Type'),'line'));
	if isempty(oldh), H=zeros(narrows,1); else, H=oldh; end;
%	% make or modify the arrows
	for k=1:narrows,
		if all(isnan(ud(k,[3 6])))&arrow_is2DXY(ax(k)), zz=[]; else, zz=z(:,k); end;
		xx=x(:,k); yy=y(:,k);
		if (0), % this fix didn't work, so let's not use it -- 8/26/03
			% try to work around a MATLAB 6.x OpenGL bug -- 7/28/02
			  mask=any([ones(1,2+size(zz,2));diff([xx yy zz],[],1)],2);
			  xx=xx(mask); yy=yy(mask); if ~isempty(zz), zz=zz(mask); end;
		end;
		% plot the patch or line
		if newpatch(k) || trueornan(ispatch(k)) %eaj 7/14/2014, 5/25/2016
			% patch is closed so don't need endpoints %eaj 7/14/2014
			if ~isempty(xx), xx(end)=[]; end; %eaj 7/14/2014
			if ~isempty(yy), yy(end)=[]; end; %eaj 7/14/2014
			if ~isempty(zz), zz(end)=[]; end; %eaj 7/14/2014
		end %eaj 7/14/2014
		xyz = {'XData',xx,'YData',yy,'ZData',zz,'Tag',ArrowTag};
		if newpatch(k)|newline(k),
			if newpatch(k),
				H(k) = patch(xyz{:});
			else,
				H(k) = line(xyz{:});
			end;
			if ~isempty(oldh), arrow_copyprops(oldh(k),H(k)); end;
		else,
			if strcmp(get(H(k),'Type'),'patch') %eaj 5/25/16  if ispatch(k)
				xyz = {xyz{:},'CData',[]};
			end;
			set(H(k),xyz{:});
		end;
	end;
	if ~isempty(oldh), delete(oldh(oldh~=H)); end;
%	% additional properties
	set(H,'Clipping','off');
	set(H,{'UserData'},num2cell(ud,2));
	if length(extraprops)>0
		ii = find(strcmpi(extraprops(1:2:end),'color')); %eaj 5/25/16
		ispatch = strcmp(get(H,'Type'),'patch');
		%eaj start 5/25/16
			while ~isempty(ii) && any(ispatch)
				if ii>1, set(H,extraprops{1:2*ii-2}); end;
				c = extraprops{2*ii};
				extraprops(1:2*ii) = [];
				ii(1) = [];
				if all(ispatch) || ischar(c)&&size(c,1)==1 || isnumeric(c)&&isequal(size(c),[1 3])
					set(H,'EdgeColor',c,'FaceColor',c)
				elseif iscell(c) && numel(c)~=numel(H)
					set(H(ispatch),'EdgeColor',c(ispatch),'FaceColor',c(ispatch));
					set(H(~ispatch),'Color',c(~ispatch));
				elseif isnumeric(c) && isequal(size(c),[numel(H) 3])
					set(H(ispatch),'EdgeColor',num2cell(c(ispatch,:),2),'FaceColor',num2cell(c(ispatch,:),2));
					set(H(~ispatch),'Color',num2cell(c(~ispatch,:),2));
				else
					warning('ignoring unknown or invalid ''Color'' specification');
				end
			end
		if ~isempty(extraprops)
		%eaj end   5/25/16
			set(H,extraprops{:});
		end %eaj   5/25/16
	end
	% handle choosing arrow Start and/or Stop locations if unspecified
	[H,oldaxlims,errstr] = arrow_clicks(H,ud,x,y,z,ax,oldaxlims);
	if ~isempty(errstr), error([upper(mfilename) ' got ' errstr]); end;
	% set the output
	if (nargout>0), h=H; end;
	% make sure the axis limits did not change
	if isempty(oldaxlims),
		ARROW_AXLIMITS = [];
		ARROW_AX = [];
	else,
		lims = get(ax(:),{'XLim','YLim','ZLim'})';
		lims = reshape(cat(2,lims{:}),6,size(lims,2));
		mask = arrow_is2DXY(ax(:));
		oldaxlims(5:6,mask) = lims(5:6,mask);
		% store them for possible restoring
		mask = any(oldaxlims~=lims,1); ARROW_AX=ax(mask); ARROW_AXLIMITS=oldaxlims(:,mask);
		if any(mask),
			warning(arrow_warnlimits(ARROW_AX,narrows));
		end;
	end;
else,
	% don't create the patch, just return the data
	h=x;
	yy=y;
	zz=z;
end;


function out = arrow_defcheck(in,def,prop)
% check if we got 'default' values
	out = in;
	if ~isstr(in), return; end;
	if size(in,1)==1 & strncmp(lower(in),'def',3),
		out = def;
	elseif ~isempty(prop),
		error([upper(mfilename) ' does not recognize ''' in(:)' ''' as a valid ''' prop ''' string.']);
	end;


function [H,oldaxlims,errstr] = arrow_clicks(H,ud,x,y,z,ax,oldaxlims)
% handle choosing arrow Start and/or Stop locations if necessary
	errstr = '';
	if isempty(H)|isempty(ud)|isempty(x), return; end;
	% determine which (if any) need Start and/or Stop
	needStart = all(isnan(ud(:,1:3)'))';
	needStop  = all(isnan(ud(:,4:6)'))';
	mask = any(needStart|needStop);
	if ~any(mask), return; end;
	ud(~mask,:)=[]; ax(:,~mask)=[];
	x(:,~mask)=[]; y(:,~mask)=[]; z(:,~mask)=[];
	% make them invisible for the time being
	set(H,'Visible','off');
	% save the current axes and limits modes; set to manual for the time being
	oldAx  = gca;
	limModes=get(ax(:),{'XLimMode','YLimMode','ZLimMode'});
	set(ax(:),{'XLimMode','YLimMode','ZLimMode'},{'manual','manual','manual'});
	% loop over each arrow that requires attention
	jj = find(mask);
	for ii=1:length(jj),
		h = H(jj(ii));
		axes(ax(ii));
		% figure out correct call
		if needStart(ii), prop='Start'; else, prop='Stop'; end;
		[wasInterrupted,errstr] = arrow_click(needStart(ii)&needStop(ii),h,prop,ax(ii));
		% handle errors and control-C
		if wasInterrupted,
			delete(H(jj(ii:end)));
			H(jj(ii:end))=[];
			oldaxlims(jj(ii:end),:)=[];
			break;
		end;
	end;
	% restore the axes and limit modes
	axes(oldAx);
	set(ax(:),{'XLimMode','YLimMode','ZLimMode'},limModes);

function [wasInterrupted,errstr] = arrow_click(lockStart,H,prop,ax)
% handle the clicks for one arrow
	fig = get(ax,'Parent');
	% save some things
	oldFigProps = {'Pointer','WindowButtonMotionFcn','WindowButtonUpFcn'};
	oldFigValue = get(fig,oldFigProps);
	oldArrowProps = {'EraseMode'};
	if ~isnumeric(fig), oldArrowProps={}; end %eaj 5/24/16 % only use in HG2
	oldArrowValue = get(H,oldArrowProps);
	if isnumeric(fig), %eaj 5/24/16
		set(H,'EraseMode','background'); %because 'xor' makes shaft invisible unless Width>1 -- only use in HG2
	end %eaj 5/24/16
	global ARROW_CLICK_H ARROW_CLICK_PROP ARROW_CLICK_AX ARROW_CLICK_USE_Z
	ARROW_CLICK_H=H; ARROW_CLICK_PROP=prop; ARROW_CLICK_AX=ax;
	ARROW_CLICK_USE_Z=~arrow_is2DXY(ax)|~arrow_planarkids(ax);
	set(fig,'Pointer','crosshair');
	% set up the WindowButtonMotion so we can see the arrow while moving around
	set(fig,'WindowButtonUpFcn','set(gcf,''WindowButtonUpFcn'','''')', ...
	        'WindowButtonMotionFcn','');
	if ~lockStart,
		set(H,'Visible','on');
		set(fig,'WindowButtonMotionFcn',[mfilename '(''callback'',''motion'');']);
	end;
	% wait for the button to be pressed
	[wasKeyPress,wasInterrupted,errstr] = arrow_wfbdown(fig);
	% if we wanted to click-drag, set the Start point
	if lockStart & ~wasInterrupted,
		pt = arrow_point(ARROW_CLICK_AX,ARROW_CLICK_USE_Z);
		feval(mfilename,H,'Start',pt,'Stop',pt);
		set(H,'Visible','on');
		ARROW_CLICK_PROP='Stop';
		set(fig,'WindowButtonMotionFcn',[mfilename '(''callback'',''motion'');']);
		% wait for the mouse button to be released
		try
			waitfor(fig,'WindowButtonUpFcn','');
		catch
			errstr = lasterr;
			wasInterrupted = 1;
		end;
	end;
	if ~wasInterrupted, feval(mfilename,'callback','motion'); end;
	% restore some things
	set(gcf,oldFigProps,oldFigValue);
	set(H,oldArrowProps,oldArrowValue);

function arrow_callback(varargin)
% handle redrawing callbacks
	if nargin==0, return; end;
	str = varargin{1};
	if ~isstr(str), error([upper(mfilename) ' got an invalid Callback command.']); end;
	s = lower(str);
	if strcmp(s,'motion'),
		% motion callback
		global ARROW_CLICK_H ARROW_CLICK_PROP ARROW_CLICK_AX ARROW_CLICK_USE_Z
		feval(mfilename,ARROW_CLICK_H,ARROW_CLICK_PROP,arrow_point(ARROW_CLICK_AX,ARROW_CLICK_USE_Z));
		drawnow;
	else,
		error([upper(mfilename) ' does not recognize ''' str(:).' ''' as a valid Callback option.']);
	end;

function out = arrow_point(ax,use_z)
% return the point on the given axes
	if nargin==0, ax=gca; end;
	if nargin<2, use_z=~arrow_is2DXY(ax)|~arrow_planarkids(ax); end;
	out = get(ax,'CurrentPoint');
	out = out(1,:);
	if ~use_z, out=out(1:2); end;

function [wasKeyPress,wasInterrupted,errstr] = arrow_wfbdown(fig)
% wait for button down ignoring object ButtonDownFcn's
	if nargin==0, fig=gcf; end;
	errstr = '';
	% save ButtonDownFcn values
	objs = findobj(fig);
	buttonDownFcns = get(objs,'ButtonDownFcn');
	mask=~strcmp(buttonDownFcns,''); objs=objs(mask); buttonDownFcns=buttonDownFcns(mask);
	set(objs,'ButtonDownFcn','');
	% save other figure values
	figProps = {'KeyPressFcn','WindowButtonDownFcn'};
	figValue = get(fig,figProps);
	% do the real work
	set(fig,'KeyPressFcn','set(gcf,''KeyPressFcn'','''',''WindowButtonDownFcn'','''');', ...
	        'WindowButtonDownFcn','set(gcf,''WindowButtonDownFcn'','''')');
	lasterr('');
	try
		waitfor(fig,'WindowButtonDownFcn','');
		wasInterrupted = 0;
	catch
		wasInterrupted = 1;
	end
	wasKeyPress = ~wasInterrupted & strcmp(get(fig,'KeyPressFcn'),'');
	if wasInterrupted, errstr=lasterr; end;
	% restore ButtonDownFcn and other figure values
	set(objs,'ButtonDownFcn',buttonDownFcns);
	set(fig,figProps,figValue);


function [out,is2D] = arrow_is2DXY(ax)
% check if axes are 2-D X-Y plots
	% may not work for modified camera angles, etc.
	out = false(size(ax)); % 2-D X-Y plots
	is2D = out;                     % any 2-D plots
	views = get(ax(:),{'View'});
	views = cat(1,views{:});
	out(:) = abs(views(:,2))==90;
	is2D(:) = out(:) | all(rem(views',90)==0)';

function out = arrow_planarkids(ax)
% check if axes descendents all have empty ZData (lines,patches,surfaces)
	out = true(size(ax));
	allkids = get(ax(:),{'Children'});
	for k=1:length(allkids),
		kids = get([findobj(allkids{k},'flat','Type','line')
		            findobj(allkids{k},'flat','Type','patch')
		            findobj(allkids{k},'flat','Type','surface')],{'ZData'});
		for j=1:length(kids),
			if ~isempty(kids{j}), out(k)=logical(0); break; end;
		end;
	end;


function arrow_fixlimits(ax,lims)
% reset the axis limits as necessary
	if isempty(ax) || isempty(lims), disp([upper(mfilename) ' does not remember any axis limits to reset.']); end;
	for k=1:numel(ax),
		if any(get(ax(k),'XLim')~=lims(1:2,k)'), set(ax(k),'XLim',lims(1:2,k)'); end;
		if any(get(ax(k),'YLim')~=lims(3:4,k)'), set(ax(k),'YLim',lims(3:4,k)'); end;
		if any(get(ax(k),'ZLim')~=lims(5:6,k)'), set(ax(k),'ZLim',lims(5:6,k)'); end;
	end;


function out = arrow_WarpToFill(notstretched,manualcamera,curax)
% check if we are in "WarpToFill" mode.
	out = strcmp(get(curax,'WarpToFill'),'on');
	% 'WarpToFill' is undocumented, so may need to replace this by
	% out = ~( any(notstretched) & any(manualcamera) );


function out = arrow_warnlimits(ax,narrows)
% create a warning message if we've changed the axis limits
	msg = '';
	switch (numel(ax))
		case 1, msg='';
		case 2, msg='on two axes ';
		otherwise, msg='on several axes ';
	end;
	msg = [upper(mfilename) ' changed the axis limits ' msg ...
	       'when adding the arrow'];
	if (narrows>1), msg=[msg 's']; end;
	out = [msg '.' sprintf('\n') '         Call ' upper(mfilename) ...
	       ' FIXLIMITS to reset them now.'];


function arrow_copyprops(fm,to)
% copy line properties to patches
	props  = {'EraseMode','LineStyle','LineWidth','Marker','MarkerSize',...
	          'MarkerEdgeColor','MarkerFaceColor','ButtonDownFcn',      ...
	          'Clipping','DeleteFcn','BusyAction','HandleVisibility',   ...
	          'Selected','SelectionHighlight','Visible'};
	if ~isnumeric(findobj('Type','root')), props(strcmp(props,'EraseMode'))=[]; end; %eaj 5/24/16
	lineprops  = {'Color',    props{:}};
	patchprops = {'EdgeColor',props{:}};
	patch2props = {'FaceColor',patchprops{:}};
	fmpatch = strcmp(get(fm,'Type'),'patch');
	topatch = strcmp(get(to,'Type'),'patch');
	set(to( fmpatch& topatch),patch2props,get(fm( fmpatch& topatch),patch2props)); %p->p
	set(to(~fmpatch&~topatch),lineprops,  get(fm(~fmpatch&~topatch),lineprops  )); %l->l
	set(to( fmpatch&~topatch),lineprops,  get(fm( fmpatch&~topatch),patchprops )); %p->l
	set(to(~fmpatch& topatch),patchprops, get(fm(~fmpatch& topatch),lineprops)  ,'FaceColor','none'); %l->p


function arrow_props
% display further help info about ARROW properties
	c = sprintf('\n');
	disp([c ...
	'ARROW Properties:  Default values are given in [square brackets], and other' c ...
	'                   acceptable equivalent property names are in (parenthesis).' c c ...
	'  Start           The starting points. For N arrows,            B' c ...
	'                  this should be a Nx2 or Nx3 matrix.          /|\           ^' c ...
	'  Stop            The end points. For N arrows, this          /|||\          |' c ...
	'                  should be a Nx2 or Nx3 matrix.             //|||\\        L|' c ...
	'  Length          Length of the arrowhead (in pixels on     ///|||\\\       e|' c ...
	'                  screen, points on a page). [16] (Len)    ////|||\\\\      n|' c ...
	'  BaseAngle       Angle (degrees) of the base angle       /////|D|\\\\\     g|' c ...
	'                  ADE.  For a simple stick arrow, use    ////  |||  \\\\    t|' c ...
	'                  BaseAngle=TipAngle. [90] (Base)       ///    |||    \\\   h|' c ...
	'  TipAngle        Angle (degrees) of tip angle ABC.    //<----->||      \\   |' c ...
	'                  [16] (Tip)                          /   base |||        \  V' c ...
	'  Width           Width of the base in pixels.  Not  E   angle ||<-------->C' c ...
	'                  the ''LineWidth'' prop. [0] (Wid)            |||tipangle' c ...
	'  Page            If provided, non-empty, and not NaN,         |||' c ...
	'                  this causes ARROW to use hardcopy            |||' c ...
	'                  rather than onscreen proportions.             A' c ...
	'                  This is important if screen aspect        -->   <-- width' c ...
	'                  ratio and hardcopy aspect ratio are    ----CrossDir---->' c ...
	'                  vastly different. []' c...
	'  CrossDir        A vector giving the direction towards which the fletches' c ...
	'                  on the arrow should go.  [computed such that it is perpen-' c ...
	'                  dicular to both the arrow direction and the view direction' c ...
	'                  (i.e., as if it was pasted on a normal 2-D graph)]  (Note' c ...
	'                  that CrossDir is a vector.  Also note that if an axis is' c ...
	'                  plotted on a log scale, then the corresponding component' c ...
	'                  of CrossDir must also be set appropriately, i.e., to 1 for' c ...
	'                  no change in that direction, >1 for a positive change, >0' c ...
	'                  and <1 for negative change.)' c ...
	'  NormalDir       A vector normal to the fletch direction (CrossDir is then' c ...
	'                  computed by the vector cross product [Line]x[NormalDir]). []' c ...
	'                  (Note that NormalDir is a vector.  Unlike CrossDir,' c ...
	'                  NormalDir is used as is regardless of log-scaled axes.)' c ...
	'  Ends            Set which end has an arrowhead.  Valid values are ''none'',' c ...
	'                  ''stop'', ''start'', and ''both''. [''stop''] (End)' c...
	'  ShortenLength   Shorten length of arrowhead(s) if line is too short' c ...
	'  ObjectHandles   Vector of handles to previously-created arrows to be' c ...
	'                  updated or line objects to be converted to arrows.' c ...
	'                  [] (Object,Handle)' c ...
	'  Type            ''patch'' creates the arrow with a PATCH object (the default)' c ...
	'                  and ''line'' creates it with a LINE object [''patch''].' c ...
	'  Color           For patch arrows (the default), set both ''FaceColor'' and' c ...
	'                  ''EdgeColor'' to the given value.  For line arrows, set' c ...
	'                  the ''Color'' property to the given value.' c ...
	]);


function out = arrow_demo
 	% demo
	% create the data
	[x,y,z] = peaks;
	[ddd,out.iii]=max(z(:));
	out.axlim = [min(x(:)) max(x(:)) min(y(:)) max(y(:)) min(z(:)) max(z(:))];
	
	% modify it by inserting some NaN's
	[m,n] = size(z);
	m = floor(m/2);
	n = floor(n/2);
	z(1:m,1:n) = NaN*ones(m,n);
	
	% graph it
	clf('reset');
	out.hs=surf(x,y,z);
	out.x=x; out.y=y; out.z=z;
	xlabel('x'); ylabel('y');
			
function h = arrow_demo3(in)
	% set the view
	axlim = in.axlim;
	axis(axlim);
	zlabel('z');
	%set(in.hs,'FaceColor','interp');
	view(3); % view(viewmtx(-37.5,30,20));
	title(['Demo of the capabilities of the ARROW function in 3-D']);
	
	% Normal blue arrow
	h1 = feval(mfilename,[axlim(1) axlim(4) 4],[-.8 1.2 4], ...
	           'EdgeColor','b','FaceColor','b');
	
	% Normal white arrow, clipped by the surface
	h2 = feval(mfilename,axlim([1 4 6]),[0 2 4]);
	t=text(-2.4,2.7,7.7,'arrow clipped by surf');
	
	% Baseangle<90
	h3 = feval(mfilename,[3 .125 3.5],[1.375 0.125 3.5],30,50);
	t2=text(3.1,.125,3.5,'local maximum');
	
	% Baseangle<90, fill and edge colors different
	h4 = feval(mfilename,axlim(1:2:5)*.5,[0 0 0],36,60,25, ...
	           'EdgeColor','b','FaceColor','c');
	t3=text(axlim(1)*.5,axlim(3)*.5,axlim(5)*.5-.75,'origin');
	set(t3,'HorizontalAlignment','center');
	
	% Baseangle>90, black fill
	h5 = feval(mfilename,[-2.9 2.9 3],[-1.3 .4 3.2],30,120,[],6, ...
	           'EdgeColor','r','FaceColor','k','LineWidth',2);
	
	% Baseangle>90, no fill
	h6 = feval(mfilename,[-2.9 2.9 1.3],[-1.3 .4 1.5],30,120,[],6, ...
	           'EdgeColor','r','FaceColor','none','LineWidth',2);
	
	% Stick arrow
	h7 = feval(mfilename,[-1.6 -1.65 -6.5],[0 -1.65 -6.5],[],16,16);
	t4=text(-1.5,-1.65,-7.25,'global mininum');
	set(t4,'HorizontalAlignment','center');
	
	% Normal, black fill
	h8 = feval(mfilename,[-1.4 0 -7.2],[-1.4 0 -3],'FaceColor','k');
	t5=text(-1.5,0,-7.75,'local minimum');
	set(t5,'HorizontalAlignment','center');
	
	% Gray fill, crossdir specified, 'LineStyle' --
	h9 = feval(mfilename,[-3 2.2 -6],[-3 2.2 -.05],36,[],27,6,[],[0 -1 0], ...
	           'EdgeColor','k','FaceColor',.75*[1 1 1],'LineStyle','--');
	
	% a series of normal arrows, linearly spaced, crossdir specified
	h10y=(0:4)'/3;
	h10 = feval(mfilename,[-3*ones(size(h10y)) h10y -6.5*ones(size(h10y))], ...
	            [-3*ones(size(h10y)) h10y -.05*ones(size(h10y))], ...
	            12,[],[],[],[],[0 -1 0]);
	
	% a series of normal arrows, linearly spaced
	h11x=(1:.33:2.8)';
	h11 = feval(mfilename,[h11x -3*ones(size(h11x)) 6.5*ones(size(h11x))], ...
	            [h11x -3*ones(size(h11x)) -.05*ones(size(h11x))]);
	
	% series of magenta arrows, radially oriented, crossdir specified
	h12x=2; h12y=-3; h12z=axlim(5)/2; h12xr=1; h12zr=h12z; ir=.15;or=.81;
	h12t=(0:11)'/6*pi;
	h12 = feval(mfilename,                                           ...
	            [h12x+h12xr*cos(h12t)*ir h12y*ones(size(h12t))       ...
	             h12z+h12zr*sin(h12t)*ir],[h12x+h12xr*cos(h12t)*or   ...
	             h12y*ones(size(h12t)) h12z+h12zr*sin(h12t)*or],     ...
	            10,[],[],[],[],                                      ...
	            [-h12xr*sin(h12t) zeros(size(h12t)) h12zr*cos(h12t)],...
	            'FaceColor','none','EdgeColor','m');
	
	% series of normal arrows, tangentially oriented, crossdir specified
	or13=.91; h13t=(0:.5:12)'/6*pi;
	locs = [h12x+h12xr*cos(h13t)*or13 h12y*ones(size(h13t)) h12z+h12zr*sin(h13t)*or13];
	h13 = feval(mfilename,locs(1:end-1,:),locs(2:end,:),6);
	
	% arrow with no line ==> oriented downwards
	h14 = feval(mfilename,[3 3 .100001],[3 3 .1],30);
	t6=text(3,3,3.6,'no line'); set(t6,'HorizontalAlignment','center');
	
	% arrow with arrowheads at both ends
	h15 = feval(mfilename,[-.5 -3 -3],[1 -3 -3],'Ends','both','FaceColor','g', ...
	            'Length',20,'Width',3,'CrossDir',[0 0 1],'TipAngle',25);
	
	h=[h1;h2;h3;h4;h5;h6;h7;h8;h9;h10;h11;h12;h13;h14;h15];

function h = arrow_demo2(in)
	axlim = in.axlim;
	dolog = 1;
	if (dolog), set(in.hs,'YData',10.^get(in.hs,'YData')); end;
	shading('interp');
	view(2);
	title(['Demo of the capabilities of the ARROW function in 2-D']);
	hold on; [C,H]=contour(in.x,in.y,in.z,20,'-'); hold off;
	for k=H',
		set(k,'ZData',(axlim(6)+1)*ones(size(get(k,'XData'))),'Color','k');
		if (dolog), set(k,'YData',10.^get(k,'YData')); end;
	end;
	if (dolog), axis([axlim(1:2) 10.^axlim(3:4)]); set(gca,'YScale','log');
	else,       axis(axlim(1:4)); end;
	
	% Normal blue arrow
	start = [axlim(1) axlim(4) axlim(6)+2];
	stop  = [in.x(in.iii) in.y(in.iii) axlim(6)+2];
	if (dolog), start(:,2)=10.^start(:,2); stop(:,2)=10.^stop(:,2); end;
	h1 = feval(mfilename,start,stop,'EdgeColor','b','FaceColor','b');
	
	% three arrows with varying fill, width, and baseangle
	start = [-3   -3   10; -3   -1.5 10; -1.5 -3   10];
	stop  = [-.03 -.03 10; -.03 -1.5 10; -1.5 -.03 10];
	if (dolog), start(:,2)=10.^start(:,2); stop(:,2)=10.^stop(:,2); end;
	h2 = feval(mfilename,start,stop,24,[90;60;120],[],[0;0;4],'Ends',str2mat('both','stop','stop'));
	set(h2(2),'EdgeColor',[0 .35 0],'FaceColor',[0 .85 .85]);
	set(h2(3),'EdgeColor','r','FaceColor',[1 .5 1]);
	h=[h1;h2];

function out = trueornan(x)
if isempty(x),
	out=x;
else,
	out = isnan(x);
	out(~out) = x(~out);
end;
```
