## Supplementary material for "Comparative food-web analysis of bluefin tuna spawning habitats in the eastern Indian Ocean and Gulf of Mexico": Code Package: CalcTL.m

```
function [TL] = CalcTL(flows,del15N,d15NInputs,Ae,InputCol)

% load('N15GoMInverseCycle1.mat')
% load('N15GoMInverseCycle1Routputs.mat')
% data = MCMCmat;
% flows=data(end,:)./wts';
% Ae=Ae0;

[NO3s_col,NH4s_col,Cyas_col,Tris_col,Dtms_col,Flags_col,HNFs_col,MICs_col,HerbNVMs_col,Apps_col,Clads_col,nvmCals_col,Chaetos_col,Poecils_col,GelPreds_col,Plankfishs_col,Preflexs_col,Postflexs_col,Bacs_col,Dets_col,Ldets_col,Dons_col,HerbVM_col,vmCal_col,NO3d_col,NH4d_col,Cyad_col,Trid_col,Dtmd_col,Flagd_col,HNFd_col,MICd_col,HerbNVMd_col,Appd_col,Cladd_col,nvmCald_col,Chaetod_col,Poecild_col,GelPredd_col,Plankfishd_col,Bacd_col,Detd_col,Ldetd_col,Dond_col] = GetColumns(Ae,flows,del15N(1,:),d15NInputs(1,InputCol),d15NInputs(2,InputCol),d15NInputs(3,InputCol),d15NInputs(4,InputCol),d15NInputs(5,InputCol),d15NInputs(6,InputCol),d15NInputs(7,InputCol));

Consumers = [HNFs_col,MICs_col,HerbNVMs_col,Apps_col,Clads_col,nvmCals_col,Chaetos_col,Poecils_col,GelPreds_col,Plankfishs_col,Preflexs_col,Postflexs_col,HerbVM_col,vmCal_col,HNFd_col,MICd_col,HerbNVMd_col,Appd_col,Cladd_col,nvmCald_col,Chaetod_col,Poecild_col,GelPredd_col,Plankfishd_col];
Dets = [Dets_col,Ldets_col,Detd_col,Ldetd_col];
Dons = [Dons_col,Dond_col];
Flags = [Flags_col,Flagd_col];
HBacs = [Bacs_col,Bacd_col];
Phys=[Cyas_col,Tris_col,Dtms_col,Cyad_col,Trid_col,Dtmd_col];
Nuts = [NO3s_col,NH4s_col,NH4s_col,NH4d_col];

%for i=1:length(data(:,1))
    %flows=data(i,:);
    TL=ones(length(Ae(:,1)),1)*2;
    for iter=1:3
        TL(Phys)=1;
        TL(Nuts)=0;
        for j=1:length(Flags)
            ToFlags = find(Ae(Flags(j),:)==1);
            total=0;
            temp=0;
            for k=1:length(ToFlags)
                ind=find(Ae(:,ToFlags(k))==-1);
                if ind~=Flags(1)
                    total=total+flows(ToFlags(k));
                    temp=temp+flows(ToFlags(k))*TL(ind);
                end
            end
            TL(Flags(j),1)=temp./total+1;
        end
        for j=1:length(Consumers)
            ToConsumers = find(Ae(Consumers(j),:)==1);
            total=0;
            temp=0;
            for k=1:length(ToConsumers)
                ind=find(Ae(:,ToConsumers(k))==-1);
                total=total+flows(ToConsumers(k));
                temp=temp+flows(ToConsumers(k))*TL(ind);
            end
            TL(Consumers(j),1)=temp./total+1;
        end
        for j=1:length(Dons)
            ToDons = find(Ae(Dons(j),:)==1);
            total=0;
            temp=0;
            for k=1:length(ToDons)
                ind=find(Ae(:,ToDons(k))==-1);
                if length(ind)==1  %Note this is necessary to avoid issues associated with external DOM not having a TL.
                    total=total+flows(ToDons(k));
                    temp=temp+flows(ToDons(k))*TL(ind);
                end
            end
            TL(Dons(j),1)=temp./total;
        end
        for j=1:length(HBacs)
            ToHBacs = find(Ae(HBacs(j),:)==1);
            total=0;
            temp=0;
            for k=1:length(ToHBacs)
                ind=find(Ae(:,ToHBacs(k))==-1);
                total=total+flows(ToHBacs(k));
                temp=temp+flows(ToHBacs(k))*TL(ind);
            end
            TL(HBacs(j),1)=temp./total+1;
        end
        for j=1:length(Dets)
            ToDets = find(Ae(Dets(j),:)==1);
            total=0;
            temp=0;
            for k=1:length(ToDets)
                ind=find(Ae(:,ToDets(k))==-1);
                if length(ind)==1  %Note this is necessary to avoid issues associated with external Det not having a TL.
                    total=total+flows(ToDets(k));
                    if length(intersect(ind,[Phys,Flags]))==1
                        temp=temp+flows(ToDets(k))*TL(ind);
                    else
                        temp=temp+flows(ToDets(k))*(TL(ind)-1);   %Note assuming that egesta has a TL equal to your food
                    end
                end
            end
            TL(Dets(j),1)=temp./total;
        end
    end
%end
        
```
