## Supplementary material for "Comparative food-web analysis of bluefin tuna spawning habitats in the eastern Indian Ocean and Gulf of Mexico": Code Package: ContinueN15InverseRWGoM.R

```
#Note that first you have to set the directory: Session>Set Working Directory
rm(list=ls())
library(R.matlab)
library(limSolve)
library(MASS)
source('xsampleN15outputs.r')
source('ExternalFunctionsGoM.r')
Cycle <- 0

tmp <- paste('N15GoMInverseCycle',Cycle,'.mat',sep="")
data <- readMat(tmp, fixNames=TRUE)
h <- data[['h']]
b <- data[['b']]
ba <- data[['ba']]
be <- data[['be']]
Ae <- data[['Ae']]
Aa <- data[['Aa']]
Aa0 <- data[['Aa0']]
wts <- data[['wts']]
G <- data[['G']]
InputCol <- data[['InputCol']]
A <- data[['A']]
Inputs <- data[['Inputs']]
W <- rep(1,39)
IterLength <- 2000000
OutputLength <- IterLength/10000

filein <- paste('N15GoMInverseCycle',Cycle,'Routputs.mat',sep="")
previous <- readMat(filein, fixNames=TRUE)
MCMCmat <- previous[['MCMCmat']]
MCMCmatplain <- previous[['MCMCmatplain']]
del15N <- previous[['del15N']]
del15Nknown <- previous[['del15Nknown']]
jmpLength <- previous[['jmpLength']]
jmpLength15 <- previous[['jmpLength15']]
sdba <- previous[['sdba']]

# upNO315N=d15NInputs[1,InputCol]
# pre15N=d15NInputs[2,InputCol]
# post15N=d15NInputs[3,InputCol]
# LDetS15N=d15NInputs[4,InputCol]
# LDetD15N=d15NInputs[5,InputCol]
# del15Nknown = c(upNO315N,pre15N,post15N,LDetS15N,LDetD15N)

for (i in 1:1000)  {
print(i)
d15N0 <- del15N[nrow(del15N),]
Startpt <- MCMCmat[nrow(MCMCmat),]


test2 <- xsampleN15outputs(A = Aa0, B = ba, E = Ae, F = be, G = G,
                           H = h, sdB = sdba, wts=wts, iter = IterLength, outputlength = OutputLength, type="mirror", jmp=jmpLength+runif(1, 0, 1)*jmpLength/5, jmp2=jmpLength15, x0 = Startpt, del15N1=d15N0, del15Nknown=del15Nknown, fulloutput='TRUE')


MCMCmat2 <- test2[['X']]
del15N2 <- test2[['del15Ntrack']]
MCMCmat <- rbind(MCMCmat,MCMCmat2)
del15N <- rbind(del15N,del15N2)
randomnumber <- test2[['randomnumber']]
p <- test2[['p']]
err <- test2[['err']]
acceptedratio <- test2[['acceptedratio']]
fileout <- paste('N15GoMInverseCycle',Cycle,'Routputs.mat',sep="")
#writeMat(fileout, MCMCmat = MCMCmat, MCMCmatplain = MCMCmatplain, del15N=del15N, Aa = Aa, ba = ba, Ae = Ae, be = be, G = G, h = h, sdba = sdba, Inputs = Inputs, InputCol = InputCol, jmpLength = jmpLength, jmpLength15 = jmpLength15, acceptedratio = acceptedratio, Startpt = Startpt, p=p, err=err)
#writeMat(fileout, MCMCmat = MCMCmat, MCMCmat_rejects = MCMCmat_rejects, MCMCmatplain = MCMCmatplain, del15N=del15N, del15N_rejects=del15N_rejects, Aa = Aa, ba = ba, Ae = Ae, be = be, G = G, h = h, sdba = sdba, Inputs = Inputs, InputCol = InputCol, jmpLength = jmpLength, jmpLength15 = jmpLength15, acceptedratio = acceptedratio, Startpt = Startpt, centralval = centralval, p=p, p_rejects=p_rejects, err=err, del15Nknown=del15Nknown, lseisol=lseisol, wts=wts)
writeMat(fileout, MCMCmat = MCMCmat, MCMCmatplain = MCMCmatplain, del15N=del15N, Aa = Aa, ba = ba, Ae = Ae, be = be, G = G, h = h, sdba = sdba, Inputs = Inputs, InputCol = InputCol, jmpLength = jmpLength, jmpLength15 = jmpLength15, acceptedratio = acceptedratio, Startpt = Startpt, p=p, err=err, del15Nknown=del15Nknown, wts=wts)

}
```
