## Supplementary material for "Comparative food-web analysis of bluefin tuna spawning habitats in the eastern Indian Ocean and Gulf of Mexico": Code Package: CreateSuppTable1.m

```
clearvars
close all


for Cycle=[1,5]
    clearvars -except Cycle tempmeantable
    
    load(['N15GoMInverseCycle',num2str(Cycle),'.mat'])
    load(['N15GoMInverseCycle',num2str(Cycle),'Routputs.mat'])
    clearvars -except Ae0 MCMCmat Aa d15NInputs del15N InputCol wts Cycle tempmeantable
    num=length(MCMCmat(:,1));
    MCMCmat(1:round(num*0.2),:)=[];
    del15N(1:round(num*0.2),:)=[];
    len=length(MCMCmat(:,1));
    [num,txt,raw]  = xlsread('N15InverseModelRW.GOM.xlsx','Sheet1','B6:B307');
    Columns = txt;
    meanvals = mean(MCMCmat)'./wts;
    stdvals = std(MCMCmat)'./wts;
    if Cycle==1
        tempmeantable(:,1)=meanvals;
        tempmeantable(:,2)=stdvals;
    elseif Cycle==5
        tempmeantable(:,3)=meanvals;
        tempmeantable(:,4)=stdvals;
    end
    
    
end

meantable = array2table(tempmeantable,'VariableNames',{'Mean1','Std1','Mean5','Std5'});
columntable = array2table(Columns);
SuppTable1 = [columntable,meantable];
```
