## Supplementary material for "Comparative food-web analysis of bluefin tuna spawning habitats in the eastern Indian Ocean and Gulf of Mexico": Code Package: ExcelCol.m

```
function output=ExcelCol(x)
	%EXCELCOL  Converts between column name and number for Excel representation
	%   OUT=ExcelCol(X) takes the input X, which may be a number or vector
	%   and converts it to the other representation
	%
	%   If IN is numeric, output will be a column cell of the column name
	%   If IN is char or cell, output will be a number or column vector,
	%      ignoring any numberic part which may be included in input
	%
	%   EXAMPLES:
	%   ExcelCol(254174985)                        %Number to column name
	%
	% $ Author: Ragaar
	% $ Copyright: 2012, Shadowstar LLC.
	% $ Original Date: 5/24/2012
	
	ABC = 'A':'Z'; % Create an array from A to Z
	col_id = dec2base(x,26); % Convert to base 26
	ABC_id = isletter(col_id); % Find any alphabetic valued outputs
	
	buffer(ABC_id) = col_id(ABC_id)-'A'+10; % Compensate for the offset
	
	% This is used to reference all numeric indices against a character
	M = num2cell(col_id(~ABC_id)); % Convert to a Cell-array
	buffer(~ABC_id) = cellfun(@str2double,M); % For-each str2double(M(i))
	
	output = ABC(buffer); % Return the string
end
```
