## Supplementary material for "Comparative food-web analysis of bluefin tuna spawning habitats in the eastern Indian Ocean and Gulf of Mexico": Code Package: ExternalFunctionsGoM.R

```
ResetRN15 <- function(Aa,flows,wts,d15N,del15NupNO3,del15NPreflexs,del15NPostflexs,del15NLdets,del15NLdetd,del15NDONs,del15NDONd,del15NApp,del15NCal,del15NChaeto,del15NHerbVM,del15NHerbNVM,del15NPoecil,del15NClad){
  
  flows <- flows/wts
  
  Eps_NO3up <- -5
  Eps_NH4up <- -10
  Eps_Nit <- -14
  Eps_ExcL <- -5
  Eps_EgL <- -1.5
  Eps_ExcS <- -1
  Eps_EgS <- -1
  Eps_ExcB <- -5
  Eps_UpB <- -1
  Eps_Remin <- 0
  RN2 <- 0
  
  col0 <- 40
  NO3s_col <- 1
  NH4s_col <- 2
  Cyas_col <- 3
  Tris_col <- 4
  Dtms_col <- 5
  Flags_col <- 6
  HNFs_col <- 7
  MICs_col <- 8
  HerbNVMs_col <- 9
  Apps_col <- 10
  Clads_col <- 11
  nvmCals_col <- 12
  Chaetos_col <- 13
  Poecils_col <- 14
  GelPreds_col <- 15
  Plankfishs_col <- 16
  Preflexs_col <- 17
  Postflexs_col <- 18
  Bacs_col <- 19
  Dets_col <- 20
  Ldets_col <- 21
  Dons_col <- 22
  HerbVM_col <- 23
  vmCal_col <- 24
  NO3d_col <- 25
  NH4d_col <- 26
  Cyad_col <- 27
  Trid_col <- 28
  Dtmd_col <- 29
  Flagd_col <- 30
  HNFd_col <- 31
  MICd_col <- 32
  HerbNVMd_col <- 33
  Appd_col <- 34
  Cladd_col <- 35
  nvmCald_col <- 36
  Chaetod_col <- 37
  Poecild_col <- 38
  GelPredd_col <- 39
  Plankfishd_col <- 40
  Bacd_col <- 41
  Detd_col <- 42
  Ldetd_col <- 43
  Dond_col <- 44
  RSDetInputs_col <- 45
  RSDetInputd_col <- 46
  
  RNO3s <- d15N[NO3s_col]
  RNH4s <- d15N[NH4s_col]
  RCyas <- d15N[Cyas_col]
  RTris <- d15N[Tris_col]
  RDtms <- d15N[Dtms_col]
  RFlags <- d15N[Flags_col]
  RHNFs <- d15N[HNFs_col]
  RMICs <- d15N[MICs_col]
  #RHerbNVMs <- d15N[HerbNVMs_col]
  #RApps <- d15N[Apps_col]
  #RClads <- d15N[Clads_col]
  #RnvmCals <- d15N[nvmCals_col]
  #RChaetos <- d15N[Chaetos_col]
  #RPoecils <- d15N[Poecils_col]
  RGelPreds <- d15N[GelPreds_col]
  RPlankfishs <- d15N[Plankfishs_col]
  #RPreflexs <- d15N[Preflexs_col]
  #RPostflexs <- d15N[Postflexs_col]
  RBacs <- d15N[Bacs_col]
  RDets <- d15N[Dets_col]
  #RLdets <- d15N[Ldets_col]
  #RDons <- d15N[Dons_col]
  #RHerbVM <- d15N[HerbVM_col]
  #RvmCal <- d15N[vmCal_col]
  RNO3d <- d15N[NO3d_col]
  RNH4d <- d15N[NH4d_col]
  RCyad <- d15N[Cyad_col]
  RTrid <- d15N[Trid_col]
  RDtmd <- d15N[Dtmd_col]
  RFlagd <- d15N[Flagd_col]
  RHNFd <- d15N[HNFd_col]
  RMICd <- d15N[MICd_col]
  #RHerbNVMd <- d15N[HerbNVMd_col]
  #RAppd <- d15N[Appd_col]
  #RCladd <- d15N[Cladd_col]
  #RnvmCald <- d15N[nvmCald_col]
  #RChaetod <- d15N[Chaetod_col]
  #RPoecild <- d15N[Poecild_col]
  RGelPredd <- d15N[GelPredd_col]
  RPlankfishd <- d15N[Plankfishd_col]
  RBacd <- d15N[Bacd_col]
  RDetd <- d15N[Detd_col]
  #RLdetd <- d15N[Ldetd_col]
  #RDond <- d15N[Dond_col]
  RSDetInputs <- d15N[RSDetInputs_col]
  RSDetInputd <- d15N[RSDetInputd_col]
  
    
  RPreflexs <- del15NPreflexs
  RPostflexs <- del15NPostflexs
  RLdets <- del15NLdets
  RLdetd <- del15NLdetd
  RupNO3 <- del15NupNO3
  RDONInputs <- del15NDONs
  RDons <- del15NDONs
  RDONInputd <- del15NDONd
  RDond <- del15NDONd
  RAppd <- del15NApp
  RApps <- del15NApp
  RCald <- del15NCal
  RnvmCals <- del15NCal
  RnvmCald <- del15NCal
  RvmCal <- del15NCal
  RChaetos <- del15NChaeto
  RChaetod <- del15NChaeto
  RHerbVM <- del15NHerbVM
  RHerbNVMs <- del15NHerbNVM
  RHerbNVMd <- del15NHerbNVM
  RClads <- del15NClad
  RCladd <- del15NClad
  # Note the following code is necessary because Poecilostomatoid 15N was not measured on Cycle 1
  if (identical(del15NPoecil,NaN)){
    RPoecils <- d15N[Poecils_col]
    RPoecild <- d15N[Poecild_col]
  }else{
    RPoecils <- del15NPoecil
    RPoecild <- del15NPoecil
  }
  
  
  # RN2 <- 0.0036765
  # RupNO3 <- del15NupNO3/1000*RN2+RN2
  # RNO3s <- del15NNO3s/1000*RN2+RN2
  # RNH4s <- del15NNH4s/1000*RN2+RN2
  # RCyas <- del15NCyas/1000*RN2+RN2
  # RTris <- del15NTris/1000*RN2+RN2
  # RDtms <- del15NDtms/1000*RN2+RN2
  # RFlags <- del15NFlags/1000*RN2+RN2
  # RHNFs <- del15NHNFs/1000*RN2+RN2
  # RMICs <- del15NMICs/1000*RN2+RN2
  # RHerbNVMs <- del15NHerbNVMs/1000*RN2+RN2
  # RApps <- del15NApps/1000*RN2+RN2
  # RClads <- del15NClads/1000*RN2+RN2
  # RnvmCals <- del15NnvmCals/1000*RN2+RN2
  # RChaetos <- del15NChaetos/1000*RN2+RN2
  # RPoecils <- del15NPoecils/1000*RN2+RN2
  # RGelPreds <- del15NGelPreds/1000*RN2+RN2
  # RPlankfishs <- del15NPlankfishs/1000*RN2+RN2
  # RPreflexs <- del15NPreflexs/1000*RN2+RN2
  # RPostflexs <- del15NPostflexs/1000*RN2+RN2
  # RBacs <- del15NBacs/1000*RN2+RN2
  # RDets <- del15NDets/1000*RN2+RN2
  # RLdets <- del15NLdets/1000*RN2+RN2
  # RDons <- del15NDons/1000*RN2+RN2
  # RHerbVM <- del15HerbVM/1000*RN2+RN2
  # RvmCal <- del15NvmCal/1000*RN2+RN2
  # RNO3d <- del15NNO3d/1000*RN2+RN2
  # RNH4d <- del15NNH4d/1000*RN2+RN2
  # RCyad <- del15NCyad/1000*RN2+RN2
  # RTrid <- del15NTrid/1000*RN2+RN2
  # RDtmd <- del15NDtmd/1000*RN2+RN2
  # RFlagd <- del15NFlagd/1000*RN2+RN2
  # RHNFd <- del15NHNFd/1000*RN2+RN2
  # RMICd <- del15NMICd/1000*RN2+RN2
  # RHerbNVMd <- del15NHerbNVMd/1000*RN2+RN2
  # RAppd <- del15NAppd/1000*RN2+RN2
  # RCladd <- del15NCladd/1000*RN2+RN2
  # RnvmCald <- del15NnvmCald/1000*RN2+RN2
  # RChaetod <- del15NChaetod/1000*RN2+RN2
  # RPoecild <- del15NPoecild/1000*RN2+RN2
  # RGelPredd <- del15NGelPredd/1000*RN2+RN2
  # RPlankfishd <- del15NPlankfishd/1000*RN2+RN2
  # RBacd <- del15NBacd/1000*RN2+RN2
  # RDetd <- del15NDetd/1000*RN2+RN2
  # RLdetd <- del15NLdetd/1000*RN2+RN2
  # RDond <- del15NDond/1000*RN2+RN2


  
  Upwellings <- 1
  LateralInputDONs <- 2
  LateralInputPONs <- 3
  NH4toNO3s <- 4
  NfixCyanos <- 5
  NO3toCyanos <- 6
  NH4toCyanos <- 7
  CyanoTOhnfs <- 8
  CyanoTOmics <- 9
  CyanoTOflags <- 10
  CyanoTOapps <- 11
  CyanoTOdoms <- 12
  CyanoTOsdets <- 13
  NfixTrichos <- 14
  NO3toTrichos <- 15
  NH4toTrichos <- 16
  TrichoTOdoms <- 17
  TrichoTOldets <- 18
  NO3toDTMs <- 19
  NH4toDTMs <- 20
  dtmTOmics <- 21
  dtmTOherbnvms <- 22
  dtmTOherbvms <- 23
  dtmTOapps <- 24
  dtmTOclads <- 25
  dtmTOnvmCals <- 26
  dtmTOvmCals <- 27
  dtmTOdoms <- 28
  dtmTOldets <- 29
  NO3TOflags <- 30
  NH4TOflags <- 31
  flagTOmics <- 32
  flagTOherbnvms <- 33
  flagTOherbvms <- 34
  flagTOapps <- 35
  flagTOclads <- 36
  flagTOnvmCals <- 37
  flagTOvmCals <- 38
  flagTOdoms <- 39
  flagTOsdets <- 40
  hnfTOmics <- 41
  hnfTOapps <- 42
  hnfTOherbnvms <- 43
  hnfTOherbvms <- 44
  hnfTOclads <- 45
  hnfTOnvmCals <- 46
  hnfTOvmCals <- 47
  hnfTOnh4s <- 48
  hnfTOdoms <- 49
  hnfTOsdets <- 50
  micTOherbnvms <- 51
  micTOherbvms <- 52
  micTOclads <- 53
  micTOnvmCals <- 54
  micTOvmCals <- 55
  micTOpreflexs <- 56
  micTOpostflexs <- 57
  micTOnh4s <- 58
  micTOdoms <- 59
  micTOsdets <- 60
  herbnvmTOchaetos <- 61
  herbnvmTOpoecils <- 62
  herbnvmTOgels <- 63
  herbnvmTOplankfishs <- 64
  herbnvmTOnh4s <- 65
  herbnvmTOdoms <- 66
  herbnvmTOldets <- 67
  appTOchaetos <- 68
  appTOpoecils <- 69
  appTOgels <- 70
  appTOplankfishs <- 71
  appTOpreflexs <- 72
  appTOpostflexs <- 73
  appTOnh4s <- 74
  appTOdoms <- 75
  appTOldets <- 76
  cladTOchaetos <- 77
  cladTOpoecils <- 78
  cladTOgels <- 79
  cladTOplankfishs <- 80
  cladTOpreflexs <- 81
  cladTOpostflexs <- 82
  cladTOnh4s <- 83
  cladTOdoms <- 84
  cladTOldets <- 85
  nvmCalTOchaetos <- 86
  nvmCalTOpoecils <- 87
  nvmCalTOgels <- 88
  nvmCalTOplankfishs <- 89
  nvmCalTOpreflexs <- 90
  nvmCalTOpostflexs <- 91
  nvmCalTOnh4s <- 92
  nvmCalTOdoms <- 93
  nvmCalTOldets <- 94
  chaetoTOgels <- 95
  chaetoTOplankfishs <- 96
  chaetoTOnh4s <- 97
  chaetoTOdoms <- 98
  chaetoTOldets <- 99
  poecilTOgels <- 100
  poecilTOplankfishs <- 101
  poecilTOpreflexs <- 102
  poecilTOpostflexs <- 103
  poecilTOnh4s <- 104
  poecilTOdoms <- 105
  poecilTOldets <- 106
  gelTOhtls <- 107
  gelTOnh4s <- 108
  gelTOdoms <- 109
  gelTOldets <- 110
  plankfishTOhtls <- 111
  plankfishTOnh4s <- 112
  plankfishTOdoms <- 113
  plankfishTOpoops <- 114
  preflexTOgrowths <- 115
  preflexTOmorts <- 116
  preflexTOnh4s <- 117
  preflexTOdoms <- 118
  preflexTOpoops <- 119
  postflexTOgrowths <- 120
  postflexTOmorts <- 121
  postflexTOnh4s <- 122
  postflexTOdoms <- 123
  postflexTOpoops <- 124
  bacTOflags <- 125
  bacTOhnfs <- 126
  bacTOmics <- 127
  bacTOapps <- 128
  bacTOnh4s <- 129
  domTObacs <- 130
  sdetTOhnfs <- 131
  sdetTOmics <- 132
  sdetTOapps <- 133
  sdetTOdoms <- 134
  ldetTOherbnvms <- 135
  ldetTOclads <- 136
  ldetTOnvmCals <- 137
  ldetTOherbvms <- 138
  ldetTOvmCals <- 139
  ldetTOdoms <- 140
  herbvmTOschaetom <- 141
  herbvmTOspoecilm <- 142
  herbvmTOsgelm <- 143
  herbvmTOsplankfishm <- 144
  herbvmTOsnh4m <- 145
  herbvmTOsdomm <- 146
  herbvmTOsldetm <- 147
  herbvmTOdchaetom <- 148
  herbvmTOdpoecilm <- 149
  herbvmTOdgelm <- 150
  herbvmTOdplankfishm <- 151
  herbvmTOdnh4m <- 152
  herbvmTOddomm <- 153
  herbvmTOdldetm <- 154
  herbvmTOexportNH4m <- 155
  herbvmTOexportDOMm <- 156
  herbvmTOexportdetm <- 157
  herbvmTOexportmortm <- 158
  vmCalTOschaetom <- 159
  vmCalTOspoecilm <- 160
  vmCalTOsgelm <- 161
  vmCalTOsplankfishm <- 162
  vmCalTOsnh4m <- 163
  vmCalTOsdomm <- 164
  vmCalTOsldetm <- 165
  vmCalTOdchaetom <- 166
  vmCalTOdpoecilm <- 167
  vmCalTOdgelm <- 168
  vmCalTOdplankfishm <- 169
  vmCalTOdnh4m <- 170
  vmCalTOddomm <- 171
  vmCalTOdldetm <- 172
  vmCalTOexportNH4m <- 173
  vmCalTOexportDOMm <- 174
  vmCalTOexportdetm <- 175
  vmCalTOexportmortm <- 176
  diatomSinkm <- 177
  FlagSinkm <- 178
  dldetSinkm <- 179
  Upwellingd <- 180
  LateralInputDONd <- 181
  LateralInputPONd <- 182
  NH4toNO3d <- 183
  NfixCyanod <- 184
  NO3toCyanod <- 185
  NH4toCyanod <- 186
  CyanoTOhnfd <- 187
  CyanoTOmicd <- 188
  CyanoTOflagd <- 189
  CyanoTOappd <- 190
  CyanoTODOCd <- 191
  CyanoTOsdetd <- 192
  NfixTrichod <- 193
  NO3toTrichod <- 194
  NH4toTrichod <- 195
  TrichoTOdomd <- 196
  TrichoTOldetd <- 197
  NO3toDTMd <- 198
  NH4toDTMd <- 199
  dtmTOmicd <- 200
  dtmTOherbnvmd <- 201
  dtmTOherbvmd <- 202
  dtmTOappd <- 203
  dtmTOcladd <- 204
  dtmTOnvmCald <- 205
  dtmTOvmCald <- 206
  dtmTOdomd <- 207
  dtmTOldetd <- 208
  NO3TOflagd <- 209
  NH4TOflagd <- 210
  flagTOmicd <- 211
  flagTOherbnvmd <- 212
  flagTOherbvmd <- 213
  flagTOappd <- 214
  flagTOcladd <- 215
  flagTOnvmCald <- 216
  flagTOvmCald <- 217
  flagTOdomd <- 218
  flagTOsdetd <- 219
  hnfTOmicd <- 220
  hnfTOappd <- 221
  hnfTOherbnvmd <- 222
  hnfTOherbvmd <- 223
  hnfTOcladd <- 224
  hnfTOnvmCald <- 225
  hnfTOvmCald <- 226
  hnfTOnh4d <- 227
  hnfTOdomd <- 228
  hnfTOsdetd <- 229
  micTOherbnvmd <- 230
  micTOherbvmd <- 231
  micTOcladd <- 232
  micTOnvmCald <- 233
  micTOvmCald <- 234
  micTOnh4d <- 235
  micTOdomd <- 236
  micTOsdetd <- 237
  herbnvmTOchaetod <- 238
  herbnvmTOpoecild <- 239
  herbnvmTOgeld <- 240
  herbnvmTOplankfishd <- 241
  herbnvmTOnh4d <- 242
  herbnvmTOldetd <- 243
  herbnvmTOdomd <- 244
  appTOchaetod <- 245
  appTOpoecild <- 246
  appTOgeld <- 247
  appTOplankfishd <- 248
  appTOnh4d <- 249
  appTOdomd <- 250
  appTOldetd <- 251
  cladTOchaetod <- 252
  cladTOpoecild <- 253
  cladTOgeld <- 254
  cladTOplankfishd <- 255
  cladTOnh4d <- 256
  cladTOdomd <- 257
  cladTOldetd <- 258
  nvmCalTOchaetod <- 259
  nvmCalTOpoecild <- 260
  nvmCalTOgeld <- 261
  nvmCalTOplankfishd <- 262
  nvmCalTOnh4d <- 263
  nvmCalTOdomd <- 264
  nvmCalTOldetd <- 265
  chaetoTOgeld <- 266
  chaetoTOplankfishd <- 267
  chaetoTOnh4d <- 268
  chaetoTOdomd <- 269
  chaetoTOldetd <- 270
  poecilTOgeld <- 271
  poecilTOplankfishd <- 272
  poecilTOnh4d <- 273
  poecilTOdomd <- 274
  poecilTOldetd <- 275
  gelTOhtld <- 276
  gelTOnh4d <- 277
  gelTOdomd <- 278
  gelTOldetd <- 279
  plankfishTOhtld <- 280
  plankfishTOnh4d <- 281
  plankfishTOdomd <- 282
  plankfishTOpoopd <- 283
  bacTOflagd <- 284
  bacTOhnfd <- 285
  bacTOmicd <- 286
  bacTOappd <- 287
  bacTOnh4d <- 288
  domTObacd <- 289
  sdetTOhnfd <- 290
  sdetTOmicd <- 291
  sdetTOappd <- 292
  sdetTOdomd <- 293
  ldetTOherbnvmd <- 294
  ldetTOcladd <- 295
  ldetTOnvmCald <- 296
  ldetTOherbvmd <- 297
  ldetTOvmCald <- 298
  ldetTOdomd <- 299
  DiatomSinkd <- 300
  FlagSinkd <- 301
  DlDetSinkd <- 302
  
  #Adding up the total amount consumed by each group
  HNFscons <- flows[CyanoTOhnfs] + flows[bacTOhnfs] + flows[sdetTOhnfs]
  MICscons <- flows[CyanoTOmics] + flows[dtmTOmics] +flows[flagTOmics] + flows[hnfTOmics] +
                  flows[bacTOmics] + flows[sdetTOmics]
  HerbNVMscons <- flows[dtmTOherbnvms] + flows[flagTOherbnvms] +
                      flows[hnfTOherbnvms] + flows[micTOherbnvms] + flows[ldetTOherbnvms]
  Appscons <- flows[CyanoTOapps] + flows[dtmTOapps] + flows[flagTOapps] + flows[hnfTOapps] +
                  flows[bacTOapps] + flows[sdetTOapps]
  Cladscons <- flows[dtmTOclads] + flows[flagTOclads] +
                   flows[hnfTOclads] + flows[micTOclads] + flows[ldetTOclads]
  nvmCalscons <- flows[dtmTOnvmCals] + flows[flagTOnvmCals] +
                     flows[hnfTOnvmCals] + flows[micTOnvmCals] + flows[ldetTOnvmCals]
  Chaetoscons <- flows[herbnvmTOchaetos] + flows[appTOchaetos] + flows[cladTOchaetos] + flows[nvmCalTOchaetos] + 
                     flows[herbvmTOschaetom] + flows[vmCalTOschaetom]
  Poecilscons <- flows[herbnvmTOpoecils] + flows[appTOpoecils] + flows[cladTOpoecils] + flows[nvmCalTOpoecils] + 
                     flows[herbvmTOspoecilm] + flows[vmCalTOspoecilm]
  GelPredscons <- flows[herbnvmTOgels] + flows[appTOgels] + flows[cladTOgels] + flows[nvmCalTOgels] + 
                      flows[chaetoTOgels] + flows[poecilTOgels] + flows[herbvmTOsgelm] + flows[vmCalTOsgelm]
  Plankfishscons <- flows[herbnvmTOplankfishs] + flows[appTOplankfishs] + flows[cladTOplankfishs] + flows[nvmCalTOplankfishs] + 
                        flows[chaetoTOplankfishs] + flows[poecilTOplankfishs] + flows[herbvmTOsplankfishm] + flows[vmCalTOsplankfishm]
  Preflexscons <- flows[micTOpreflexs] + flows[appTOpreflexs] + 
                      flows[cladTOpreflexs] + flows[nvmCalTOpreflexs] + flows[poecilTOpreflexs]
  Postflexscons <- flows[micTOpostflexs] + flows[appTOpostflexs] + 
                       flows[cladTOpostflexs] + flows[nvmCalTOpostflexs] + flows[poecilTOpostflexs]
  HerbVMcons <- flows[dtmTOherbvms] + flows[flagTOherbvms] + flows[hnfTOherbvms] + flows[micTOherbvms] + 
                    flows[ldetTOherbvms] + flows[dtmTOherbvmd] + flows[flagTOherbvmd] + flows[hnfTOherbvmd] + 
                    flows[micTOherbvmd] + flows[ldetTOherbvmd]
  vmCalcons <- flows[dtmTOvmCals] + flows[flagTOvmCals] + flows[hnfTOvmCals] + flows[micTOvmCals] + 
                   flows[ldetTOvmCals] + flows[dtmTOvmCald] + flows[flagTOvmCald] + flows[hnfTOvmCald] + 
                   flows[micTOvmCald] + flows[ldetTOvmCald]
  HNFdcons <- flows[CyanoTOhnfd] + flows[bacTOhnfd] + flows[sdetTOhnfd]
  MICdcons <- flows[CyanoTOmicd] + flows[dtmTOmicd] + flows[flagTOmicd] + flows[hnfTOmicd] +
                  flows[bacTOmicd] + flows[sdetTOmicd]
  HerbNVMdcons <- flows[dtmTOherbnvmd] + flows[flagTOherbnvmd] + flows[hnfTOherbnvmd] + flows[micTOherbnvmd] + 
                      flows[ldetTOherbnvmd]
  Appdcons <- flows[CyanoTOappd] + flows[dtmTOappd] + flows[flagTOappd] + flows[hnfTOappd] +
                  flows[bacTOappd] + flows[sdetTOappd]
  Claddcons <- flows[dtmTOcladd] + flows[flagTOcladd] +
                   flows[hnfTOcladd] + flows[micTOcladd] + flows[ldetTOcladd]
  nvmCaldcons <- flows[dtmTOnvmCald] + flows[flagTOnvmCald] +
                     flows[hnfTOnvmCald] + flows[micTOnvmCald] + flows[ldetTOnvmCald]
  Chaetodcons <- flows[herbnvmTOchaetod] + flows[appTOchaetod] + flows[cladTOchaetod] + flows[nvmCalTOchaetod] + 
                     flows[herbvmTOdchaetom] + flows[vmCalTOdchaetom]
  Poecildcons <- flows[herbnvmTOpoecild] + flows[appTOpoecild] + flows[cladTOpoecild] + flows[nvmCalTOpoecild] + 
                     flows[herbvmTOdpoecilm] + flows[vmCalTOdpoecilm]
  GelPreddcons <- flows[herbnvmTOgeld] + flows[appTOgeld] + flows[cladTOgeld] + flows[nvmCalTOgeld] + 
                      flows[chaetoTOgeld] + flows[poecilTOgeld] + flows[herbvmTOdgelm] + flows[vmCalTOdgelm]
  Plankfishdcons <- flows[herbnvmTOplankfishd] + flows[appTOplankfishd] + flows[cladTOplankfishd] + flows[nvmCalTOplankfishd] + 
                        flows[chaetoTOplankfishd] + flows[poecilTOplankfishd] + flows[herbvmTOdplankfishm] + flows[vmCalTOdplankfishm]
  
  #Calculating the isotopic composition of the food of each group
  RHNFsfood <- (RCyas*flows[CyanoTOhnfs] + RBacs*flows[bacTOhnfs] + 
                  RDets*flows[sdetTOhnfs])/HNFscons
  RMICsfood <- (RCyas*flows[CyanoTOmics] + RDtms*flows[dtmTOmics] +
                  RFlags*flows[flagTOmics] + RHNFs*flows[hnfTOmics] +
                  RBacs*flows[bacTOmics] + 
                  RDets*flows[sdetTOmics])/MICscons
  RHerbNVMsfood <- (RDtms*flows[dtmTOherbnvms] + RFlags*flows[flagTOherbnvms] +
                  RHNFs*flows[hnfTOherbnvms] + RMICs*flows[micTOherbnvms] + 
                  RLdets*flows[ldetTOherbnvms])/HerbNVMscons
  RAppsfood <- (RCyas*flows[CyanoTOapps] + RDtms*flows[dtmTOapps] + 
                  RFlags*flows[flagTOapps] + RHNFs*flows[hnfTOapps] +
                  RBacs*flows[bacTOapps] + RDets*flows[sdetTOapps])/Appscons 
  RCladsfood <- (RDtms*flows[dtmTOclads] + RFlags*flows[flagTOclads] +
                      RHNFs*flows[hnfTOclads] + RMICs*flows[micTOclads] + 
                      RLdets*flows[ldetTOclads])/Cladscons
  RnvmCalsfood <- (RDtms*flows[dtmTOnvmCals] + RFlags*flows[flagTOnvmCals] +
                   RHNFs*flows[hnfTOnvmCals] + RMICs*flows[micTOnvmCals] + 
                   RLdets*flows[ldetTOnvmCals])/nvmCalscons
  RChaetosfood <- (RHerbNVMs*flows[herbnvmTOchaetos] + RApps*flows[appTOchaetos] + 
                     RClads*flows[cladTOchaetos] + RnvmCals*flows[nvmCalTOchaetos] + 
                     RHerbVM*flows[herbvmTOschaetom] + RvmCal*flows[vmCalTOschaetom])/Chaetoscons
  RPoecilsfood <- (RHerbNVMs*flows[herbnvmTOpoecils] + RApps*flows[appTOpoecils] + 
                     RClads*flows[cladTOpoecils] + RnvmCals*flows[nvmCalTOpoecils] + 
                     RHerbVM*flows[herbvmTOspoecilm] + RvmCal*flows[vmCalTOspoecilm])/Poecilscons
  RGelPredsfood <- (RHerbNVMs*flows[herbnvmTOgels] + RApps*flows[appTOgels] + 
                     RClads*flows[cladTOgels] + RnvmCals*flows[nvmCalTOgels] + 
                     RChaetos*flows[chaetoTOgels] + RPoecils*flows[poecilTOgels] + 
                     RHerbVM*flows[herbvmTOsgelm] + RvmCal*flows[vmCalTOsgelm])/GelPredscons
  RPlankfishsfood <- (RHerbNVMs*flows[herbnvmTOplankfishs] + RApps*flows[appTOplankfishs] + 
                      RClads*flows[cladTOplankfishs] + RnvmCals*flows[nvmCalTOplankfishs] + 
                      RChaetos*flows[chaetoTOplankfishs] + RPoecils*flows[poecilTOplankfishs] + 
                      RHerbVM*flows[herbvmTOsplankfishm] + RvmCal*flows[vmCalTOsplankfishm])/Plankfishscons
  RPreflexsfood <- (RMICs*flows[micTOpreflexs] + RApps*flows[appTOpreflexs] + 
                        RClads*flows[cladTOpreflexs] + RnvmCals*flows[nvmCalTOpreflexs] + 
                        RPoecils*flows[poecilTOpreflexs])/Preflexscons
  RPostflexsfood <- (RMICs*flows[micTOpostflexs] + RApps*flows[appTOpostflexs] + 
                      RClads*flows[cladTOpostflexs] + RnvmCals*flows[nvmCalTOpostflexs] + 
                      RPoecils*flows[poecilTOpostflexs])/Postflexscons
  RHerbVMfood <- (RDtms*flows[dtmTOherbvms] + RFlags*flows[flagTOherbvms] +
                      RHNFs*flows[hnfTOherbvms] + RMICs*flows[micTOherbvms] + 
                      RLdets*flows[ldetTOherbvms] + RDtmd*flows[dtmTOherbvmd] + 
                    RFlagd*flows[flagTOherbvmd] + RHNFd*flows[hnfTOherbvmd] + 
                    RMICd*flows[micTOherbvmd] + RLdetd*flows[ldetTOherbvmd])/HerbVMcons
  RvmCalfood <- (RDtms*flows[dtmTOvmCals] + RFlags*flows[flagTOvmCals] +
                    RHNFs*flows[hnfTOvmCals] + RMICs*flows[micTOvmCals] + 
                    RLdets*flows[ldetTOvmCals] + RDtmd*flows[dtmTOvmCald] + 
                    RFlagd*flows[flagTOvmCald] + RHNFd*flows[hnfTOvmCald] + 
                    RMICd*flows[micTOvmCald] + RLdetd*flows[ldetTOvmCald])/vmCalcons
  RHNFdfood <- (RCyad*flows[CyanoTOhnfd] + RBacd*flows[bacTOhnfd] + 
                  RDetd*flows[sdetTOhnfd])/HNFdcons
  RMICdfood <- (RCyad*flows[CyanoTOmicd] + RDtmd*flows[dtmTOmicd] +
                  RFlagd*flows[flagTOmicd] + RHNFd*flows[hnfTOmicd] +
                  RBacd*flows[bacTOmicd] + 
                  RDetd*flows[sdetTOmicd])/MICdcons
  RHerbNVMdfood <- (RDtmd*flows[dtmTOherbnvmd] + RFlagd*flows[flagTOherbnvmd] +
                      RHNFd*flows[hnfTOherbnvmd] + RMICd*flows[micTOherbnvmd] + 
                      RLdetd*flows[ldetTOherbnvmd])/HerbNVMdcons
  RAppdfood <- (RCyad*flows[CyanoTOappd] + RDtmd*flows[dtmTOappd] + 
                  RFlagd*flows[flagTOappd] + RHNFd*flows[hnfTOappd] +
                  RBacd*flows[bacTOappd] + RDetd*flows[sdetTOappd])/Appdcons 
  RCladdfood <- (RDtmd*flows[dtmTOcladd] + RFlagd*flows[flagTOcladd] +
                   RHNFd*flows[hnfTOcladd] + RMICd*flows[micTOcladd] + 
                   RLdetd*flows[ldetTOcladd])/Claddcons
  RnvmCaldfood <- (RDtmd*flows[dtmTOnvmCald] + RFlagd*flows[flagTOnvmCald] +
                     RHNFd*flows[hnfTOnvmCald] + RMICd*flows[micTOnvmCald] + 
                     RLdetd*flows[ldetTOnvmCald])/nvmCaldcons
  RChaetodfood <- (RHerbNVMd*flows[herbnvmTOchaetod] + RAppd*flows[appTOchaetod] + 
                     RCladd*flows[cladTOchaetod] + RnvmCald*flows[nvmCalTOchaetod] + 
                     RHerbVM*flows[herbvmTOdchaetom] + RvmCal*flows[vmCalTOdchaetom])/Chaetodcons
  RPoecildfood <- (RHerbNVMd*flows[herbnvmTOpoecild] + RAppd*flows[appTOpoecild] + 
                     RCladd*flows[cladTOpoecild] + RnvmCald*flows[nvmCalTOpoecild] + 
                     RHerbVM*flows[herbvmTOdpoecilm] + RvmCal*flows[vmCalTOdpoecilm])/Poecildcons
  RGelPreddfood <- (RHerbNVMd*flows[herbnvmTOgeld] + RAppd*flows[appTOgeld] + 
                      RCladd*flows[cladTOgeld] + RnvmCald*flows[nvmCalTOgeld] + 
                      RChaetod*flows[chaetoTOgeld] + RPoecild*flows[poecilTOgeld] + 
                      RHerbVM*flows[herbvmTOdgelm] + RvmCal*flows[vmCalTOdgelm])/GelPreddcons
  RPlankfishdfood <- (RHerbNVMd*flows[herbnvmTOplankfishd] + RAppd*flows[appTOplankfishd] + 
                        RCladd*flows[cladTOplankfishd] + RnvmCald*flows[nvmCalTOplankfishd] + 
                        RChaetod*flows[chaetoTOplankfishd] + RPoecild*flows[poecilTOplankfishd] + 
                        RHerbVM*flows[herbvmTOdplankfishm] + RvmCal*flows[vmCalTOdplankfishm])/Plankfishdcons
  
 #Computing the sum of each row for normalization purposes
  totflowNO3s <- sum(flows[c(Upwellings,NH4toNO3s)])
  totflowNH4s <- sum(flows[c(hnfTOnh4s,micTOnh4s,herbnvmTOnh4s,appTOnh4s,cladTOnh4s,nvmCalTOnh4s,chaetoTOnh4s,poecilTOnh4s,gelTOnh4s,plankfishTOnh4s,preflexTOnh4s,postflexTOnh4s,bacTOnh4s,herbvmTOsnh4m,vmCalTOsnh4m)])
  totflowCyas <- sum(flows[c(NfixCyanos,NO3toCyanos,NH4toCyanos)])
  totflowTris <- sum(flows[c(NfixTrichos,NO3toTrichos,NH4toTrichos)])
  totflowDtms <- sum(flows[c(NO3toDTMs,NH4toDTMs)])
  totflowFlags <- sum(flows[c(CyanoTOflags,NO3TOflags,NH4TOflags,bacTOflags)])
  totflowHNFs <- sum(flows[c(CyanoTOhnfs,bacTOhnfs,sdetTOhnfs)])
  totflowMICs <- sum(flows[c(CyanoTOmics,dtmTOmics,flagTOmics,hnfTOmics,bacTOmics,sdetTOmics)])
  totflowHerbNVMs <- sum(flows[c(dtmTOherbnvms,flagTOherbnvms,hnfTOherbnvms,micTOherbnvms,ldetTOherbnvms)])
  totflowApps <- sum(flows[c(CyanoTOapps,dtmTOapps,flagTOapps,hnfTOapps,bacTOapps,sdetTOapps)])
  totflowClads <- sum(flows[c(dtmTOclads,flagTOclads,hnfTOclads,micTOclads,ldetTOclads)])
  totflownvmCals <- sum(flows[c(dtmTOnvmCals,flagTOnvmCals,hnfTOnvmCals,micTOnvmCals,ldetTOnvmCals)])
  totflowChaetos <- sum(flows[c(herbnvmTOchaetos,appTOchaetos,cladTOchaetos,nvmCalTOchaetos,herbvmTOschaetom,vmCalTOschaetom)])
  totflowPoecils <- sum(flows[c(herbnvmTOpoecils,appTOpoecils,cladTOpoecils,nvmCalTOpoecils,herbvmTOspoecilm,vmCalTOspoecilm)])
  totflowGelPreds <- sum(flows[c(herbnvmTOgels,appTOgels,cladTOgels,nvmCalTOgels,chaetoTOgels,poecilTOgels,herbvmTOsgelm,vmCalTOsgelm)])
  totflowPlankfishs <- sum(flows[c(herbnvmTOplankfishs,appTOplankfishs,cladTOplankfishs,nvmCalTOplankfishs,chaetoTOplankfishs,poecilTOplankfishs,herbvmTOsplankfishm,vmCalTOsplankfishm)])
  totflowPreflexs <- sum(flows[c(micTOpreflexs,appTOpreflexs,cladTOpreflexs,nvmCalTOpreflexs,poecilTOpreflexs)])
  totflowPostflexs <- sum(flows[c(micTOpostflexs,appTOpostflexs,cladTOpostflexs,nvmCalTOpostflexs,poecilTOpostflexs)])
  totflowBacs <- sum(flows[c(domTObacs)])
  totflowDets <- sum(flows[c(LateralInputPONs,CyanoTOsdets,hnfTOsdets,micTOsdets)])
  totflowLdets <- sum(flows[c(TrichoTOldets,dtmTOldets,flagTOsdets,herbnvmTOldets,appTOldets,cladTOldets,nvmCalTOldets,chaetoTOldets,poecilTOldets,gelTOldets,herbvmTOsldetm,vmCalTOsldetm)])
  totflowDons <- sum(flows[c(domTObacs)])
  totflowHerbVM <- sum(flows[c(dtmTOherbvms,flagTOherbvms,hnfTOherbvms,micTOherbvms,ldetTOherbvms,dtmTOherbvmd,flagTOherbvmd,hnfTOherbvmd,micTOherbvmd,ldetTOherbvmd)])
  totflowvmCal <- sum(flows[c(dtmTOvmCals,flagTOvmCals,hnfTOvmCals,micTOvmCals,ldetTOvmCals,dtmTOvmCald,flagTOvmCald,hnfTOvmCald,micTOvmCald,ldetTOvmCald)])
  totflowNO3d <- sum(flows[c(Upwellingd,NH4toNO3d)])
  totflowNH4d <- sum(flows[c(herbvmTOdnh4m,vmCalTOdnh4m,hnfTOnh4d,micTOnh4d,herbnvmTOnh4d,appTOnh4d,cladTOnh4d,nvmCalTOnh4d,chaetoTOnh4d,poecilTOnh4d,gelTOnh4d,plankfishTOnh4d,bacTOnh4d)])
  totflowCyad <- sum(flows[c(NfixCyanod,NO3toCyanod,NH4toCyanod)])
  totflowTrid <- sum(flows[c(NfixTrichod,NO3toTrichod,NH4toTrichod)])
  totflowDtmd <- sum(flows[c(NO3toDTMd,NH4toDTMd)])
  totflowFlagd <- sum(flows[c(FlagSinkm,CyanoTOflagd,NO3TOflagd,NH4TOflagd,bacTOflagd)])
  totflowHNFd <- sum(flows[c(CyanoTOhnfd,bacTOhnfd,sdetTOhnfd)])
  totflowMICd <- sum(flows[c(CyanoTOmicd,dtmTOmicd,flagTOmicd,hnfTOmicd,bacTOmicd,sdetTOmicd)])
  totflowHerbNVMd <- sum(flows[c(dtmTOherbnvmd,flagTOherbnvmd,hnfTOherbnvmd,micTOherbnvmd,ldetTOherbnvmd)])
  totflowAppd <- sum(flows[c(CyanoTOappd,dtmTOappd,flagTOappd,hnfTOappd,bacTOappd,sdetTOappd)])
  totflowCladd <- sum(flows[c(dtmTOcladd,flagTOcladd,hnfTOcladd,micTOcladd,ldetTOcladd)])
  totflownvmCald <- sum(flows[c(dtmTOnvmCald,flagTOnvmCald,hnfTOnvmCald,micTOnvmCald,ldetTOnvmCald)])
  totflowChaetod <- sum(flows[c(herbvmTOdchaetom,vmCalTOdchaetom,herbnvmTOchaetod,appTOchaetod,cladTOchaetod,nvmCalTOchaetod)])
  totflowPoecild <- sum(flows[c(herbvmTOdpoecilm,vmCalTOdpoecilm,herbnvmTOpoecild,appTOpoecild,cladTOpoecild,nvmCalTOpoecild)])
  totflowGelPredd <- sum(flows[c(herbvmTOdgelm,vmCalTOdgelm,herbnvmTOgeld,appTOgeld,cladTOgeld,nvmCalTOgeld,chaetoTOgeld,poecilTOgeld)])
  totflowPlankfishd <- sum(flows[c(herbvmTOdplankfishm,vmCalTOdplankfishm,herbnvmTOplankfishd,appTOplankfishd,cladTOplankfishd,nvmCalTOplankfishd,chaetoTOplankfishd,poecilTOplankfishd)])
  totflowBacd <- sum(flows[c(domTObacd)])
  totflowDetd <- sum(flows[c(LateralInputPONd,CyanoTOsdetd,hnfTOsdetd,micTOsdetd)])
  totflowLdetd <- sum(flows[c(herbvmTOdldetm,vmCalTOdldetm,dldetSinkm,TrichoTOldetd,dtmTOldetd,flagTOsdetd,herbnvmTOldetd,appTOldetd,cladTOldetd,nvmCalTOldetd,chaetoTOldetd,poecilTOldetd,gelTOldetd)])
  totflowDond <- sum(flows[c(domTObacd)])
  

  Aa[col0+NO3s_col,Upwellings] <- RNO3d
  Aa[col0+NO3s_col,NH4toNO3s] <- RNH4s+Eps_Nit
  Aa[col0+NO3s_col,NO3toCyanos] <- -(RNO3s+Eps_NO3up)
  Aa[col0+NO3s_col,NO3toTrichos] <- -(RNO3s+Eps_NO3up)
  Aa[col0+NO3s_col,NO3toDTMs] <- -(RNO3s+Eps_NO3up)
  Aa[col0+NO3s_col,NO3TOflags] <- -(RNO3s+Eps_NO3up)
  Aa[col0+NH4s_col,NH4toNO3s] <- -(RNH4s+Eps_Nit)
  Aa[col0+NH4s_col,NH4toCyanos] <- -(RNH4s+Eps_NH4up)
  Aa[col0+NH4s_col,NH4toTrichos] <- -(RNH4s+Eps_NH4up)
  Aa[col0+NH4s_col,NH4toDTMs] <- -(RNH4s+Eps_NH4up)
  Aa[col0+NH4s_col,NH4TOflags] <- -(RNH4s+Eps_NH4up)
  Aa[col0+NH4s_col,hnfTOnh4s] <- RHNFs+Eps_ExcS
  Aa[col0+NH4s_col,micTOnh4s] <- RMICs+Eps_ExcS
  Aa[col0+NH4s_col,herbnvmTOnh4s] <- RHerbNVMs+Eps_ExcL
  Aa[col0+NH4s_col,appTOnh4s] <- RApps+Eps_ExcL
  Aa[col0+NH4s_col,cladTOnh4s] <- RClads+Eps_ExcL
  Aa[col0+NH4s_col,nvmCalTOnh4s] <- RnvmCals+Eps_ExcL
  Aa[col0+NH4s_col,chaetoTOnh4s] <- RChaetos+Eps_ExcL
  Aa[col0+NH4s_col,poecilTOnh4s] <- RPoecils+Eps_ExcL
  Aa[col0+NH4s_col,gelTOnh4s] <- RGelPreds+Eps_ExcL
  Aa[col0+NH4s_col,plankfishTOnh4s] <- RPlankfishs+Eps_ExcL
  Aa[col0+NH4s_col,preflexTOnh4s] <- RPreflexs+Eps_ExcL
  Aa[col0+NH4s_col,postflexTOnh4s] <- RPostflexs+Eps_ExcL
  Aa[col0+NH4s_col,bacTOnh4s] <- RBacs+Eps_ExcB
  Aa[col0+NH4s_col,herbvmTOsnh4m] <- RHerbVM+Eps_ExcL
  Aa[col0+NH4s_col,vmCalTOsnh4m] <- RvmCal+Eps_ExcL
  Aa[col0+Cyas_col,NfixCyanos] <- RN2
  Aa[col0+Cyas_col,NO3toCyanos] <- (RNO3s+Eps_NO3up)
  Aa[col0+Cyas_col,NH4toCyanos] <- (RNH4s+Eps_NH4up)
  Aa[col0+Cyas_col,CyanoTOhnfs] <- -RCyas
  Aa[col0+Cyas_col,CyanoTOmics] <- -RCyas
  Aa[col0+Cyas_col,CyanoTOflags] <- -RCyas
  Aa[col0+Cyas_col,CyanoTOapps] <- -RCyas
  Aa[col0+Cyas_col,CyanoTOdoms] <- -RCyas
  Aa[col0+Cyas_col,CyanoTOsdets] <- -RCyas
  Aa[col0+Tris_col,NfixTrichos] <- RN2
  Aa[col0+Tris_col,NO3toTrichos] <- (RNO3s+Eps_NO3up)
  Aa[col0+Tris_col,NH4toTrichos] <- (RNH4s+Eps_NH4up)
  Aa[col0+Tris_col,TrichoTOdoms] <- -RTris
  Aa[col0+Tris_col,TrichoTOldets] <- -RTris
  Aa[col0+Dtms_col,NO3toDTMs] <- (RNO3s+Eps_NO3up)
  Aa[col0+Dtms_col,NH4toDTMs] <- (RNH4s+Eps_NH4up)
  Aa[col0+Dtms_col,dtmTOmics] <- -RDtms
  Aa[col0+Dtms_col,dtmTOherbnvms] <- -RDtms
  Aa[col0+Dtms_col,dtmTOherbvms] <- -RDtms
  Aa[col0+Dtms_col,dtmTOapps] <- -RDtms
  Aa[col0+Dtms_col,dtmTOclads] <- -RDtms
  Aa[col0+Dtms_col,dtmTOnvmCals] <- -RDtms
  Aa[col0+Dtms_col,dtmTOvmCals] <- -RDtms
  Aa[col0+Dtms_col,dtmTOdoms] <- -RDtms
  Aa[col0+Dtms_col,dtmTOldets] <- -RDtms
  Aa[col0+Dtms_col,diatomSinkm] <- -RDtms
  Aa[col0+Flags_col,CyanoTOflags] <- RCyas
  Aa[col0+Flags_col,NO3TOflags] <- (RNO3s+Eps_NO3up)
  Aa[col0+Flags_col,NH4TOflags] <- (RNH4s+Eps_NH4up)
  Aa[col0+Flags_col,flagTOmics] <- -RFlags
  Aa[col0+Flags_col,flagTOherbnvms] <- -RFlags
  Aa[col0+Flags_col,flagTOherbvms] <- -RFlags
  Aa[col0+Flags_col,flagTOapps] <- -RFlags
  Aa[col0+Flags_col,flagTOclads] <- -RFlags
  Aa[col0+Flags_col,flagTOnvmCals] <- -RFlags
  Aa[col0+Flags_col,flagTOvmCals] <- -RFlags
  Aa[col0+Flags_col,flagTOdoms] <- -RFlags
  Aa[col0+Flags_col,flagTOsdets] <- -RFlags
  Aa[col0+Flags_col,bacTOflags] <- RBacs
  Aa[col0+Flags_col,FlagSinkm] <- -RFlags
  Aa[col0+HNFs_col,CyanoTOhnfs] <- RCyas
  Aa[col0+HNFs_col,hnfTOmics] <- -RHNFs
  Aa[col0+HNFs_col,hnfTOapps] <- -RHNFs
  Aa[col0+HNFs_col,hnfTOherbnvms] <- -RHNFs
  Aa[col0+HNFs_col,hnfTOherbvms] <- -RHNFs
  Aa[col0+HNFs_col,hnfTOclads] <- -RHNFs
  Aa[col0+HNFs_col,hnfTOnvmCals] <- -RHNFs
  Aa[col0+HNFs_col,hnfTOvmCals] <- -RHNFs
  Aa[col0+HNFs_col,hnfTOnh4s] <- -(RHNFs+Eps_ExcS)
  Aa[col0+HNFs_col,hnfTOdoms] <- -(RHNFs+Eps_ExcS)
  Aa[col0+HNFs_col,hnfTOsdets] <- -(RHNFsfood+Eps_EgS)
  Aa[col0+HNFs_col,bacTOhnfs] <- RBacs
  Aa[col0+HNFs_col,sdetTOhnfs] <- RDets
  Aa[col0+MICs_col,CyanoTOmics] <- RCyas
  Aa[col0+MICs_col,dtmTOmics] <- RDtms
  Aa[col0+MICs_col,flagTOmics] <- RFlags
  Aa[col0+MICs_col,hnfTOmics] <- RHNFs
  Aa[col0+MICs_col,micTOherbnvms] <- -RMICs
  Aa[col0+MICs_col,micTOherbvms] <- -RMICs
  Aa[col0+MICs_col,micTOclads] <- -RMICs
  Aa[col0+MICs_col,micTOnvmCals] <- -RMICs
  Aa[col0+MICs_col,micTOvmCals] <- -RMICs
  Aa[col0+MICs_col,micTOpreflexs] <- -RMICs
  Aa[col0+MICs_col,micTOpostflexs] <- -RMICs
  Aa[col0+MICs_col,micTOnh4s] <- -(RMICs+Eps_ExcS)
  Aa[col0+MICs_col,micTOdoms] <- -(RMICs+Eps_ExcS)
  Aa[col0+MICs_col,micTOsdets] <- -(RMICsfood+Eps_EgS)
  Aa[col0+MICs_col,bacTOmics] <-  RBacs
  Aa[col0+MICs_col,sdetTOmics] <-  RDets
  Aa[col0+HerbNVMs_col,dtmTOherbnvms] <- RDtms
  Aa[col0+HerbNVMs_col,flagTOherbnvms] <- RFlags
  Aa[col0+HerbNVMs_col,hnfTOherbnvms] <- RHNFs
  Aa[col0+HerbNVMs_col,micTOherbnvms] <- RMICs
  Aa[col0+HerbNVMs_col,herbnvmTOchaetos] <- -RHerbNVMs
  Aa[col0+HerbNVMs_col,herbnvmTOpoecils] <- -RHerbNVMs
  Aa[col0+HerbNVMs_col,herbnvmTOgels] <- -RHerbNVMs
  Aa[col0+HerbNVMs_col,herbnvmTOplankfishs] <- -RHerbNVMs
  Aa[col0+HerbNVMs_col,herbnvmTOnh4s] <- -(RHerbNVMs+Eps_ExcL)
  Aa[col0+HerbNVMs_col,herbnvmTOdoms] <- -(RHerbNVMs+Eps_ExcL)
  Aa[col0+HerbNVMs_col,herbnvmTOldets] <- -(RHerbNVMsfood+Eps_EgL)
  Aa[col0+HerbNVMs_col,ldetTOherbnvms] <- RLdets
  Aa[col0+Apps_col,CyanoTOapps] <- RCyas
  Aa[col0+Apps_col,dtmTOapps] <- RDtms
  Aa[col0+Apps_col,flagTOapps] <- RFlags
  Aa[col0+Apps_col,hnfTOapps] <- RHNFs
  Aa[col0+Apps_col,appTOchaetos] <- -RApps
  Aa[col0+Apps_col,appTOpoecils] <- -RApps
  Aa[col0+Apps_col,appTOgels] <- -RApps
  Aa[col0+Apps_col,appTOplankfishs] <- -RApps
  Aa[col0+Apps_col,appTOpreflexs] <- -RApps
  Aa[col0+Apps_col,appTOpostflexs] <- -RApps
  Aa[col0+Apps_col,appTOnh4s] <- -(RApps+Eps_ExcL)
  Aa[col0+Apps_col,appTOdoms] <- -(RApps+Eps_ExcL)
  Aa[col0+Apps_col,appTOldets] <- -(RAppsfood+Eps_EgL)
  Aa[col0+Apps_col,bacTOapps] <- RBacs
  Aa[col0+Apps_col,sdetTOapps] <- RDets
  Aa[col0+Clads_col,dtmTOclads] <- RDtms
  Aa[col0+Clads_col,flagTOclads] <- RFlags
  Aa[col0+Clads_col,hnfTOclads] <- RHNFs
  Aa[col0+Clads_col,micTOclads] <- RMICs
  Aa[col0+Clads_col,cladTOchaetos] <- -RClads
  Aa[col0+Clads_col,cladTOpoecils] <- -RClads
  Aa[col0+Clads_col,cladTOgels] <- -RClads
  Aa[col0+Clads_col,cladTOplankfishs] <- -RClads
  Aa[col0+Clads_col,cladTOpreflexs] <- -RClads
  Aa[col0+Clads_col,cladTOpostflexs] <- -RClads
  Aa[col0+Clads_col,cladTOnh4s] <- -(RClads+Eps_ExcL)
  Aa[col0+Clads_col,cladTOdoms] <- -(RClads+Eps_ExcL)
  Aa[col0+Clads_col,cladTOldets] <- -(RCladsfood+Eps_EgL)
  Aa[col0+Clads_col,ldetTOclads] <- RLdets
  Aa[col0+nvmCals_col,dtmTOnvmCals] <- RDtms
  Aa[col0+nvmCals_col,flagTOnvmCals] <- RFlags
  Aa[col0+nvmCals_col,hnfTOnvmCals] <- RHNFs
  Aa[col0+nvmCals_col,micTOnvmCals] <- RMICs
  Aa[col0+nvmCals_col,nvmCalTOchaetos] <- -RnvmCals
  Aa[col0+nvmCals_col,nvmCalTOpoecils] <- -RnvmCals
  Aa[col0+nvmCals_col,nvmCalTOgels] <- -RnvmCals
  Aa[col0+nvmCals_col,nvmCalTOplankfishs] <- -RnvmCals
  Aa[col0+nvmCals_col,nvmCalTOpreflexs] <- -RnvmCals
  Aa[col0+nvmCals_col,nvmCalTOpostflexs] <- -RnvmCals
  Aa[col0+nvmCals_col,nvmCalTOnh4s] <- -(RnvmCals+Eps_ExcL)
  Aa[col0+nvmCals_col,nvmCalTOdoms] <- -(RnvmCals+Eps_ExcL)
  Aa[col0+nvmCals_col,nvmCalTOldets] <- -(RnvmCalsfood+Eps_EgL)
  Aa[col0+nvmCals_col,ldetTOnvmCals] <- RLdets
  Aa[col0+Chaetos_col,herbnvmTOchaetos] <- RHerbNVMs
  Aa[col0+Chaetos_col,appTOchaetos] <- RApps
  Aa[col0+Chaetos_col,cladTOchaetos] <- RClads
  Aa[col0+Chaetos_col,nvmCalTOchaetos] <- RnvmCals
  Aa[col0+Chaetos_col,chaetoTOgels] <- -RChaetos
  Aa[col0+Chaetos_col,chaetoTOplankfishs] <- -RChaetos
  Aa[col0+Chaetos_col,chaetoTOnh4s] <- -(RChaetos+Eps_ExcL)
  Aa[col0+Chaetos_col,chaetoTOdoms] <- -(RChaetos+Eps_ExcL)
  Aa[col0+Chaetos_col,chaetoTOldets] <- -(RChaetosfood+Eps_EgL)
  Aa[col0+Chaetos_col,herbvmTOschaetom] <- RHerbVM
  Aa[col0+Chaetos_col,vmCalTOschaetom] <- RvmCal
  Aa[col0+Poecils_col,herbnvmTOpoecils] <- RHerbNVMs
  Aa[col0+Poecils_col,appTOpoecils] <- RApps
  Aa[col0+Poecils_col,cladTOpoecils] <- RClads
  Aa[col0+Poecils_col,nvmCalTOpoecils] <- RnvmCals
  Aa[col0+Poecils_col,poecilTOgels] <- -RPoecils
  Aa[col0+Poecils_col,poecilTOplankfishs] <- -RPoecils
  Aa[col0+Poecils_col,poecilTOpreflexs] <- -RPoecils
  Aa[col0+Poecils_col,poecilTOpostflexs] <- -RPoecils
  Aa[col0+Poecils_col,poecilTOnh4s] <- -(RPoecils+Eps_ExcL)
  Aa[col0+Poecils_col,poecilTOdoms] <- -(RPoecils+Eps_ExcL)
  Aa[col0+Poecils_col,poecilTOldets] <- -(RPoecilsfood+Eps_EgL)
  Aa[col0+Poecils_col,herbvmTOspoecilm] <- RHerbVM
  Aa[col0+Poecils_col,vmCalTOspoecilm] <- RvmCal
  Aa[col0+GelPreds_col,herbnvmTOgels] <- RHerbNVMs
  Aa[col0+GelPreds_col,appTOgels] <- RApps
  Aa[col0+GelPreds_col,cladTOgels] <- RClads
  Aa[col0+GelPreds_col,nvmCalTOgels] <- RnvmCals
  Aa[col0+GelPreds_col,chaetoTOgels] <- RChaetos
  Aa[col0+GelPreds_col,poecilTOgels] <- RPoecils
  Aa[col0+GelPreds_col,gelTOhtls] <- -RGelPreds
  Aa[col0+GelPreds_col,gelTOnh4s] <- -(RGelPreds+Eps_ExcL)
  Aa[col0+GelPreds_col,gelTOdoms] <- -(RGelPreds+Eps_ExcL)
  Aa[col0+GelPreds_col,gelTOldets] <- -(RGelPredsfood+Eps_EgL)
  Aa[col0+GelPreds_col,herbvmTOsgelm] <- RHerbVM
  Aa[col0+GelPreds_col,vmCalTOsgelm] <- RvmCal
  Aa[col0+Plankfishs_col,herbnvmTOplankfishs] <- RHerbNVMs
  Aa[col0+Plankfishs_col,appTOplankfishs] <- RApps
  Aa[col0+Plankfishs_col,cladTOplankfishs] <- RClads
  Aa[col0+Plankfishs_col,nvmCalTOplankfishs] <- RnvmCals
  Aa[col0+Plankfishs_col,chaetoTOplankfishs] <- RChaetos
  Aa[col0+Plankfishs_col,poecilTOplankfishs] <- RPoecils
  Aa[col0+Plankfishs_col,plankfishTOhtls] <- -RPlankfishs
  Aa[col0+Plankfishs_col,plankfishTOnh4s] <- -(RPlankfishs+Eps_ExcL)
  Aa[col0+Plankfishs_col,plankfishTOdoms] <- -(RPlankfishs+Eps_ExcL)
  Aa[col0+Plankfishs_col,plankfishTOpoops] <- -(RPlankfishsfood+Eps_EgL)
  Aa[col0+Plankfishs_col,herbvmTOsplankfishm] <- RHerbVM
  Aa[col0+Plankfishs_col,vmCalTOsplankfishm] <- RvmCal
  Aa[col0+Preflexs_col,micTOpreflexs] <- RMICs
  Aa[col0+Preflexs_col,appTOpreflexs] <- RApps
  Aa[col0+Preflexs_col,cladTOpreflexs] <- RClads
  Aa[col0+Preflexs_col,nvmCalTOpreflexs] <- RnvmCals
  Aa[col0+Preflexs_col,poecilTOpreflexs] <- RPoecils
  Aa[col0+Preflexs_col,preflexTOgrowths] <- -RPreflexs
  Aa[col0+Preflexs_col,preflexTOmorts] <- -RPreflexs
  Aa[col0+Preflexs_col,preflexTOnh4s] <- -(RPreflexs+Eps_ExcL)
  Aa[col0+Preflexs_col,preflexTOdoms] <- -(RPreflexs+Eps_ExcL)
  Aa[col0+Preflexs_col,preflexTOpoops] <- -(RPreflexsfood+Eps_EgL)
  Aa[col0+Postflexs_col,micTOpostflexs] <- RMICs
  Aa[col0+Postflexs_col,appTOpostflexs] <- RApps
  Aa[col0+Postflexs_col,cladTOpostflexs] <- RClads
  Aa[col0+Postflexs_col,nvmCalTOpostflexs] <- RnvmCals
  Aa[col0+Postflexs_col,poecilTOpostflexs] <- RPoecils
  Aa[col0+Postflexs_col,postflexTOgrowths] <- -RPostflexs
  Aa[col0+Postflexs_col,postflexTOmorts] <- -RPostflexs
  Aa[col0+Postflexs_col,postflexTOnh4s] <- -(RPostflexs+Eps_ExcL)
  Aa[col0+Postflexs_col,postflexTOdoms] <- -(RPostflexs+Eps_ExcL)
  Aa[col0+Postflexs_col,postflexTOpoops] <- -(RPostflexsfood+Eps_EgL)
  Aa[col0+Bacs_col,bacTOflags] <- -RBacs
  Aa[col0+Bacs_col,bacTOhnfs] <- -RBacs
  Aa[col0+Bacs_col,bacTOmics] <- -RBacs
  Aa[col0+Bacs_col,bacTOapps] <- -RBacs
  Aa[col0+Bacs_col,bacTOnh4s] <- -(RBacs+Eps_ExcB)
  Aa[col0+Bacs_col,domTObacs] <- RDons+Eps_UpB
  Aa[col0+Dets_col,LateralInputPONs] <- RSDetInputs
  Aa[col0+Dets_col,CyanoTOsdets] <- RCyas
  Aa[col0+Dets_col,flagTOsdets] <- RFlags
  Aa[col0+Dets_col,hnfTOsdets] <- (RHNFsfood+Eps_EgS)
  Aa[col0+Dets_col,micTOsdets] <- (RMICsfood+Eps_EgS)
  Aa[col0+Dets_col,sdetTOhnfs] <- -RDets
  Aa[col0+Dets_col,sdetTOmics] <- -RDets
  Aa[col0+Dets_col,sdetTOapps] <- -RDets
  Aa[col0+Dets_col,sdetTOdoms] <- -(RDets+Eps_Remin)
  Aa[col0+Ldets_col,TrichoTOldets] <- RTris
  Aa[col0+Ldets_col,dtmTOldets] <- RDtms
  Aa[col0+Ldets_col,herbnvmTOldets] <- (RHerbNVMsfood+Eps_EgL)
  Aa[col0+Ldets_col,appTOldets] <- (RAppsfood+Eps_EgL)
  Aa[col0+Ldets_col,cladTOldets] <- (RCladsfood+Eps_EgL)
  Aa[col0+Ldets_col,nvmCalTOldets] <- (RnvmCalsfood+Eps_EgL)
  Aa[col0+Ldets_col,chaetoTOldets] <- (RChaetosfood+Eps_EgL)
  Aa[col0+Ldets_col,poecilTOldets] <- (RPoecilsfood+Eps_EgL)
  Aa[col0+Ldets_col,gelTOldets] <- (RGelPredsfood+Eps_EgL)
  Aa[col0+Ldets_col,ldetTOherbnvms] <- -RLdets
  Aa[col0+Ldets_col,ldetTOclads] <- -RLdets
  Aa[col0+Ldets_col,ldetTOnvmCals] <- -RLdets
  Aa[col0+Ldets_col,ldetTOherbvms] <- -RLdets
  Aa[col0+Ldets_col,ldetTOvmCals] <- -RLdets
  Aa[col0+Ldets_col,ldetTOdoms] <- -(RLdets+Eps_Remin)
  Aa[col0+Ldets_col,herbvmTOsldetm] <- (RHerbVMfood+Eps_EgL)
  Aa[col0+Ldets_col,vmCalTOsldetm] <- (RvmCalfood+Eps_EgL)
  Aa[col0+Ldets_col,dldetSinkm] <- -RLdets
  Aa[col0+Dons_col,LateralInputDONs] <- RDONInputs
  Aa[col0+Dons_col,CyanoTOdoms] <- RCyas
  Aa[col0+Dons_col,TrichoTOdoms] <- RTris
  Aa[col0+Dons_col,dtmTOdoms] <- RDtms
  Aa[col0+Dons_col,flagTOdoms] <- RFlags
  Aa[col0+Dons_col,hnfTOdoms] <- (RHNFs+Eps_ExcS)
  Aa[col0+Dons_col,micTOdoms] <- (RMICs+Eps_ExcS)
  Aa[col0+Dons_col,herbnvmTOdoms] <- (RHerbNVMs+Eps_ExcL)
  Aa[col0+Dons_col,appTOdoms] <- (RApps+Eps_ExcL)
  Aa[col0+Dons_col,cladTOdoms] <- (RClads+Eps_ExcL)
  Aa[col0+Dons_col,nvmCalTOdoms] <- (RnvmCals+Eps_ExcL)
  Aa[col0+Dons_col,chaetoTOdoms] <- (RChaetos+Eps_ExcL)
  Aa[col0+Dons_col,poecilTOdoms] <- (RPoecils+Eps_ExcL)
  Aa[col0+Dons_col,gelTOdoms] <- (RGelPreds+Eps_ExcL)
  Aa[col0+Dons_col,plankfishTOdoms] <- (RPlankfishs+Eps_ExcL)
  Aa[col0+Dons_col,preflexTOdoms] <- (RPreflexs+Eps_ExcL)
  Aa[col0+Dons_col,postflexTOdoms] <-  (RPostflexs+Eps_ExcL)
  Aa[col0+Dons_col,domTObacs] <- -(RDons+Eps_UpB)
  Aa[col0+Dons_col,sdetTOdoms] <- (RDets+Eps_Remin)
  Aa[col0+Dons_col,ldetTOdoms] <- (RLdets+Eps_Remin)
  Aa[col0+Dons_col,herbvmTOsdomm] <- (RHerbVM+Eps_ExcL)
  Aa[col0+Dons_col,vmCalTOsdomm] <- (RvmCal+Eps_ExcL)
  Aa[col0+HerbVM_col,dtmTOherbvms] <- RDtms
  Aa[col0+HerbVM_col,flagTOherbvms] <- RFlags
  Aa[col0+HerbVM_col,hnfTOherbvms] <- RHNFs
  Aa[col0+HerbVM_col,micTOherbvms] <- RMICs
  Aa[col0+HerbVM_col,ldetTOherbvms] <- RLdets
  Aa[col0+HerbVM_col,herbvmTOschaetom] <- -RHerbVM
  Aa[col0+HerbVM_col,herbvmTOspoecilm] <- -RHerbVM
  Aa[col0+HerbVM_col,herbvmTOsgelm] <- -RHerbVM
  Aa[col0+HerbVM_col,herbvmTOsplankfishm] <- -RHerbVM
  Aa[col0+HerbVM_col,herbvmTOsnh4m] <- -(RHerbVM+Eps_ExcL)
  Aa[col0+HerbVM_col,herbvmTOsdomm] <- -(RHerbVM+Eps_ExcL)
  Aa[col0+HerbVM_col,herbvmTOsldetm] <- -(RHerbVMfood+Eps_EgL)
  Aa[col0+HerbVM_col,herbvmTOdchaetom] <- -RHerbVM
  Aa[col0+HerbVM_col,herbvmTOdpoecilm] <- -RHerbVM
  Aa[col0+HerbVM_col,herbvmTOdgelm] <- -RHerbVM
  Aa[col0+HerbVM_col,herbvmTOdplankfishm] <- -RHerbVM
  Aa[col0+HerbVM_col,herbvmTOdnh4m] <- -(RHerbVM+Eps_ExcL)
  Aa[col0+HerbVM_col,herbvmTOddomm] <- -(RHerbVM+Eps_ExcL)
  Aa[col0+HerbVM_col,herbvmTOdldetm] <- -(RHerbVMfood+Eps_EgL)
  Aa[col0+HerbVM_col,herbvmTOexportNH4m] <- -(RHerbVM+Eps_ExcL)
  Aa[col0+HerbVM_col,herbvmTOexportDOMm] <- -(RHerbVM+Eps_ExcL)
  Aa[col0+HerbVM_col,herbvmTOexportdetm] <- -(RHerbVMfood+Eps_EgL)
  Aa[col0+HerbVM_col,herbvmTOexportmortm] <- -RHerbVM
  Aa[col0+HerbVM_col,dtmTOherbvmd] <- -RDtmd
  Aa[col0+HerbVM_col,flagTOherbvmd] <- -RFlagd
  Aa[col0+HerbVM_col,hnfTOherbvmd] <- -RHNFd
  Aa[col0+HerbVM_col,micTOherbvmd] <- -RMICd
  Aa[col0+HerbVM_col,ldetTOherbvmd] <- -RLdetd
  Aa[col0+vmCal_col,dtmTOvmCals] <- RDtms
  Aa[col0+vmCal_col,flagTOvmCals] <- RFlags
  Aa[col0+vmCal_col,hnfTOvmCals] <- RHNFs
  Aa[col0+vmCal_col,micTOvmCals] <- RMICs
  Aa[col0+vmCal_col,ldetTOvmCals] <- RLdets
  Aa[col0+vmCal_col,vmCalTOschaetom] <- -RvmCal
  Aa[col0+vmCal_col,vmCalTOspoecilm] <- -RvmCal
  Aa[col0+vmCal_col,vmCalTOsgelm] <- -RvmCal
  Aa[col0+vmCal_col,vmCalTOsplankfishm] <- -RvmCal
  Aa[col0+vmCal_col,vmCalTOsnh4m] <- -(RvmCal+Eps_ExcL)
  Aa[col0+vmCal_col,vmCalTOsdomm] <- -(RvmCal+Eps_ExcL)
  Aa[col0+vmCal_col,vmCalTOsldetm] <- -(RvmCalfood+Eps_EgL)
  Aa[col0+vmCal_col,vmCalTOdchaetom] <- -RvmCal
  Aa[col0+vmCal_col,vmCalTOdpoecilm] <- -RvmCal
  Aa[col0+vmCal_col,vmCalTOdgelm] <- -RvmCal
  Aa[col0+vmCal_col,vmCalTOdplankfishm] <- -RvmCal
  Aa[col0+vmCal_col,vmCalTOdnh4m] <- -(RvmCal+Eps_ExcL)
  Aa[col0+vmCal_col,vmCalTOddomm] <- -(RvmCal+Eps_ExcL)
  Aa[col0+vmCal_col,vmCalTOdldetm] <- -(RvmCalfood+Eps_EgL)
  Aa[col0+vmCal_col,vmCalTOexportNH4m] <- -(RvmCal+Eps_ExcL)
  Aa[col0+vmCal_col,vmCalTOexportDOMm] <- -(RvmCal+Eps_ExcL)
  Aa[col0+vmCal_col,vmCalTOexportdetm] <- -(RvmCalfood+Eps_EgL)
  Aa[col0+vmCal_col,vmCalTOexportmortm] <- -RvmCal
  Aa[col0+vmCal_col,dtmTOvmCald] <- RDtmd
  Aa[col0+vmCal_col,flagTOvmCald] <- RFlagd
  Aa[col0+vmCal_col,hnfTOvmCald] <- RHNFd
  Aa[col0+vmCal_col,micTOvmCald] <- RMICd
  Aa[col0+vmCal_col,ldetTOvmCald] <- RLdetd
  Aa[col0+NO3d_col,Upwellings] <- -RNO3d
  Aa[col0+NO3d_col,Upwellingd] <- RupNO3
  Aa[col0+NO3d_col,NH4toNO3d] <- RNH4d+Eps_Nit
  Aa[col0+NO3d_col,NO3toCyanod] <- -(RNO3d+Eps_NO3up)
  Aa[col0+NO3d_col,NO3toTrichod] <- -(RNO3d+Eps_NO3up)
  Aa[col0+NO3d_col,NO3toDTMd] <- -(RNO3d+Eps_NO3up)
  Aa[col0+NO3d_col,NO3TOflagd] <- -(RNO3d+Eps_NO3up)
  Aa[col0+NH4d_col,herbvmTOdnh4m] <- RHerbVM+Eps_ExcL
  Aa[col0+NH4d_col,vmCalTOdnh4m] <- RvmCal+Eps_ExcL
  Aa[col0+NH4d_col,NH4toNO3d] <- -(RNH4d+Eps_Nit)
  Aa[col0+NH4d_col,NH4toCyanod] <- -(RNH4d+Eps_NH4up)
  Aa[col0+NH4d_col,NH4toTrichod] <- -(RNH4d+Eps_NH4up)
  Aa[col0+NH4d_col,NH4toDTMd] <- -(RNH4d+Eps_NH4up)
  Aa[col0+NH4d_col,NH4TOflagd] <- -(RNH4d+Eps_NH4up)
  Aa[col0+NH4d_col,hnfTOnh4d] <- RHNFd+Eps_ExcS
  Aa[col0+NH4d_col,micTOnh4d] <- RMICd+Eps_ExcS
  Aa[col0+NH4d_col,herbnvmTOnh4d] <- RHerbNVMd+Eps_ExcL
  Aa[col0+NH4d_col,appTOnh4d] <- RAppd+Eps_ExcL
  Aa[col0+NH4d_col,cladTOnh4d] <- RCladd+Eps_ExcL
  Aa[col0+NH4d_col,nvmCalTOnh4d] <- RnvmCald+Eps_ExcL
  Aa[col0+NH4d_col,chaetoTOnh4d] <- RChaetod+Eps_ExcL
  Aa[col0+NH4d_col,poecilTOnh4d] <- RPoecild+Eps_ExcL
  Aa[col0+NH4d_col,gelTOnh4d] <- RGelPredd+Eps_ExcL
  Aa[col0+NH4d_col,plankfishTOnh4d] <- RPlankfishd+Eps_ExcL
  Aa[col0+NH4d_col,bacTOnh4d] <- RBacd+Eps_ExcB
  Aa[col0+Cyad_col,NfixCyanod] <- RN2
  Aa[col0+Cyad_col,NO3toCyanod] <- (RNO3d+Eps_NO3up)
  Aa[col0+Cyad_col,NH4toCyanod] <- (RNH4d+Eps_NH4up)
  Aa[col0+Cyad_col,CyanoTOhnfd] <- -RCyad
  Aa[col0+Cyad_col,CyanoTOmicd] <- -RCyad
  Aa[col0+Cyad_col,CyanoTOflagd] <- -RCyad
  Aa[col0+Cyad_col,CyanoTOappd] <- -RCyad
  Aa[col0+Cyad_col,CyanoTODOCd] <- -RCyad
  Aa[col0+Cyad_col,CyanoTOsdetd] <- -RCyad
  Aa[col0+Trid_col,NfixTrichod] <- RN2
  Aa[col0+Trid_col,NO3toTrichod] <- (RNO3d+Eps_NO3up)
  Aa[col0+Trid_col,NH4toTrichod] <- (RNH4d+Eps_NH4up)
  Aa[col0+Trid_col,TrichoTOdomd] <- -RTrid
  Aa[col0+Trid_col,TrichoTOldetd] <- -RTrid
  Aa[col0+Dtmd_col,diatomSinkm] <- RDtms
  Aa[col0+Dtmd_col,NO3toDTMd] <- (RNO3d+Eps_NO3up)
  Aa[col0+Dtmd_col,NH4toDTMd] <- (RNH4d+Eps_NH4up)
  Aa[col0+Dtmd_col,dtmTOmicd] <- -RDtmd
  Aa[col0+Dtmd_col,dtmTOherbnvmd] <- -RDtmd
  Aa[col0+Dtmd_col,dtmTOherbvmd] <- -RDtmd
  Aa[col0+Dtmd_col,dtmTOappd] <- -RDtmd
  Aa[col0+Dtmd_col,dtmTOcladd] <- -RDtmd
  Aa[col0+Dtmd_col,dtmTOnvmCald] <- -RDtmd
  Aa[col0+Dtmd_col,dtmTOvmCald] <- -RDtmd
  Aa[col0+Dtmd_col,dtmTOdomd] <- -RDtmd
  Aa[col0+Dtmd_col,dtmTOldetd] <- -RDtmd
  Aa[col0+Dtmd_col,DiatomSinkd] <- -RDtmd
  Aa[col0+Flagd_col,FlagSinkm] <- RFlags
  Aa[col0+Flagd_col,CyanoTOflagd] <- RCyad
  Aa[col0+Flagd_col,NO3TOflagd] <- (RNO3d+Eps_NO3up)
  Aa[col0+Flagd_col,NH4TOflagd] <- (RNH4d+Eps_NH4up)
  Aa[col0+Flagd_col,flagTOmicd] <- -RFlagd
  Aa[col0+Flagd_col,flagTOherbnvmd] <- -RFlagd
  Aa[col0+Flagd_col,flagTOherbvmd] <- -RFlagd
  Aa[col0+Flagd_col,flagTOappd] <- -RFlagd
  Aa[col0+Flagd_col,flagTOcladd] <- -RFlagd
  Aa[col0+Flagd_col,flagTOnvmCald] <- -RFlagd
  Aa[col0+Flagd_col,flagTOvmCald] <- -RFlagd
  Aa[col0+Flagd_col,flagTOdomd] <- -RFlagd
  Aa[col0+Flagd_col,flagTOsdetd] <- -RFlagd
  Aa[col0+Flagd_col,bacTOflagd] <- RBacd
  Aa[col0+Flagd_col,FlagSinkd] <- -RFlagd
  Aa[col0+HNFd_col,CyanoTOhnfd] <- RCyad
  Aa[col0+HNFd_col,hnfTOmicd] <- -RHNFd
  Aa[col0+HNFd_col,hnfTOappd] <- -RHNFd
  Aa[col0+HNFd_col,hnfTOherbnvmd] <- -RHNFd
  Aa[col0+HNFd_col,hnfTOherbvmd] <- -RHNFd
  Aa[col0+HNFd_col,hnfTOcladd] <- -RHNFd
  Aa[col0+HNFd_col,hnfTOnvmCald] <- -RHNFd
  Aa[col0+HNFd_col,hnfTOvmCald] <- -RHNFd
  Aa[col0+HNFd_col,hnfTOnh4d] <- -(RHNFd+Eps_ExcS)
  Aa[col0+HNFd_col,hnfTOdomd] <- -(RHNFd+Eps_ExcS)
  Aa[col0+HNFd_col,hnfTOsdetd] <- -(RHNFdfood+Eps_EgS)
  Aa[col0+HNFd_col,bacTOhnfd] <- RBacd
  Aa[col0+HNFd_col,sdetTOhnfd] <- RDetd
  Aa[col0+MICd_col,CyanoTOmicd] <- RCyad
  Aa[col0+MICd_col,dtmTOmicd] <- RDtmd
  Aa[col0+MICd_col,flagTOmicd] <- RFlagd
  Aa[col0+MICd_col,hnfTOmicd] <- RHNFd
  Aa[col0+MICd_col,micTOherbnvmd] <- -RMICd
  Aa[col0+MICd_col,micTOherbvmd] <- -RMICd
  Aa[col0+MICd_col,micTOcladd] <- -RMICd
  Aa[col0+MICd_col,micTOnvmCald] <- -RMICd
  Aa[col0+MICd_col,micTOvmCald] <- -RMICd
  Aa[col0+MICd_col,micTOnh4d] <- -(RMICd+Eps_ExcS)
  Aa[col0+MICd_col,micTOdomd] <- -(RMICd+Eps_ExcS)
  Aa[col0+MICd_col,micTOsdetd] <- -(RMICdfood+Eps_EgS)
  Aa[col0+MICd_col,bacTOmicd] <- RBacd
  Aa[col0+MICd_col,sdetTOmicd] <- RDetd
  Aa[col0+HerbNVMd_col,dtmTOherbnvmd] <- RDtmd
  Aa[col0+HerbNVMd_col,flagTOherbnvmd] <- RFlagd
  Aa[col0+HerbNVMd_col,hnfTOherbnvmd] <- RHNFd
  Aa[col0+HerbNVMd_col,micTOherbnvmd] <- RMICd
  Aa[col0+HerbNVMd_col,herbnvmTOchaetod] <- -RHerbNVMd
  Aa[col0+HerbNVMd_col,herbnvmTOpoecild] <- -RHerbNVMd
  Aa[col0+HerbNVMd_col,herbnvmTOgeld] <- -RHerbNVMd
  Aa[col0+HerbNVMd_col,herbnvmTOplankfishd] <- -RHerbNVMd
  Aa[col0+HerbNVMd_col,herbnvmTOnh4d] <- -(RHerbNVMd+Eps_ExcL)
  Aa[col0+HerbNVMd_col,herbnvmTOldetd] <- -(RHerbNVMdfood+Eps_EgL)
  Aa[col0+HerbNVMd_col,herbnvmTOdomd] <- -(RHerbNVMd+Eps_ExcL)
  Aa[col0+HerbNVMd_col,ldetTOherbnvmd] <- RLdetd
  Aa[col0+Appd_col,CyanoTOappd] <- RCyad
  Aa[col0+Appd_col,dtmTOappd] <- RDtmd
  Aa[col0+Appd_col,flagTOappd] <- RFlagd
  Aa[col0+Appd_col,hnfTOappd] <- RHNFd
  Aa[col0+Appd_col,appTOchaetod] <- -RAppd
  Aa[col0+Appd_col,appTOpoecild] <- -RAppd
  Aa[col0+Appd_col,appTOgeld] <- -RAppd
  Aa[col0+Appd_col,appTOplankfishd] <- -RAppd
  Aa[col0+Appd_col,appTOnh4d] <- -(RAppd+Eps_ExcL)
  Aa[col0+Appd_col,appTOdomd] <- -(RAppd+Eps_ExcL)
  Aa[col0+Appd_col,appTOldetd] <- -(RAppdfood+Eps_EgL)
  Aa[col0+Appd_col,bacTOappd] <- RBacd
  Aa[col0+Appd_col,sdetTOappd] <- RDetd
  Aa[col0+Cladd_col,dtmTOcladd] <- RDtmd
  Aa[col0+Cladd_col,flagTOcladd] <- RFlagd
  Aa[col0+Cladd_col,hnfTOcladd] <- RHNFd
  Aa[col0+Cladd_col,micTOcladd] <- RMICd
  Aa[col0+Cladd_col,cladTOchaetod] <- -RCladd
  Aa[col0+Cladd_col,cladTOpoecild] <- -RCladd
  Aa[col0+Cladd_col,cladTOgeld] <- -RCladd
  Aa[col0+Cladd_col,cladTOplankfishd] <- -RCladd
  Aa[col0+Cladd_col,cladTOnh4d] <- -(RCladd+Eps_ExcL)
  Aa[col0+Cladd_col,cladTOdomd] <- -(RCladd+Eps_ExcL)
  Aa[col0+Cladd_col,cladTOldetd] <- -(RCladdfood+Eps_EgL)
  Aa[col0+Cladd_col,ldetTOcladd] <- RLdetd
  Aa[col0+nvmCald_col,dtmTOnvmCald] <- RDtmd
  Aa[col0+nvmCald_col,flagTOnvmCald] <- RFlagd
  Aa[col0+nvmCald_col,hnfTOnvmCald] <- RHNFd
  Aa[col0+nvmCald_col,micTOnvmCald] <- RMICd
  Aa[col0+nvmCald_col,nvmCalTOchaetod] <- -RnvmCald
  Aa[col0+nvmCald_col,nvmCalTOpoecild] <- -RnvmCald
  Aa[col0+nvmCald_col,nvmCalTOgeld] <- -RnvmCald
  Aa[col0+nvmCald_col,nvmCalTOplankfishd] <- -RnvmCald
  Aa[col0+nvmCald_col,nvmCalTOnh4d] <- -(RnvmCald+Eps_ExcL)
  Aa[col0+nvmCald_col,nvmCalTOdomd] <- -(RnvmCald+Eps_ExcL)
  Aa[col0+nvmCald_col,nvmCalTOldetd] <- -(RnvmCaldfood+Eps_EgL)
  Aa[col0+nvmCald_col,ldetTOnvmCald] <- RLdetd
  Aa[col0+Chaetod_col,herbvmTOdchaetom] <- RHerbVM
  Aa[col0+Chaetod_col,vmCalTOdchaetom] <- RvmCal
  Aa[col0+Chaetod_col,herbnvmTOchaetod] <- RHerbNVMd
  Aa[col0+Chaetod_col,appTOchaetod] <- RAppd
  Aa[col0+Chaetod_col,cladTOchaetod] <- RCladd
  Aa[col0+Chaetod_col,nvmCalTOchaetod] <- RnvmCald
  Aa[col0+Chaetod_col,chaetoTOgeld] <- -RChaetod
  Aa[col0+Chaetod_col,chaetoTOplankfishd] <- -RChaetod
  Aa[col0+Chaetod_col,chaetoTOnh4d] <- -(RChaetod+Eps_ExcL)
  Aa[col0+Chaetod_col,chaetoTOdomd] <- -(RChaetod+Eps_ExcL)
  Aa[col0+Chaetod_col,chaetoTOldetd] <- -(RChaetodfood+Eps_EgL)
  Aa[col0+Poecild_col,herbvmTOdpoecilm] <- RHerbVM
  Aa[col0+Poecild_col,vmCalTOdpoecilm] <- RvmCal
  Aa[col0+Poecild_col,herbnvmTOpoecild] <- RHerbNVMd
  Aa[col0+Poecild_col,appTOpoecild] <- RAppd
  Aa[col0+Poecild_col,cladTOpoecild] <- RCladd
  Aa[col0+Poecild_col,nvmCalTOpoecild] <- RnvmCald
  Aa[col0+Poecild_col,poecilTOgeld] <- -RPoecild
  Aa[col0+Poecild_col,poecilTOplankfishd] <- -RPoecild
  Aa[col0+Poecild_col,poecilTOnh4d] <- -(RPoecild+Eps_ExcL)
  Aa[col0+Poecild_col,poecilTOdomd] <- -(RPoecild+Eps_ExcL)
  Aa[col0+Poecild_col,poecilTOldetd] <- -(RPoecildfood+Eps_EgL)
  Aa[col0+GelPredd_col,herbvmTOdgelm] <- RHerbVM
  Aa[col0+GelPredd_col,vmCalTOdgelm] <- RvmCal
  Aa[col0+GelPredd_col,herbnvmTOgeld] <- RHerbNVMd
  Aa[col0+GelPredd_col,appTOgeld] <- RAppd
  Aa[col0+GelPredd_col,cladTOgeld] <- RCladd
  Aa[col0+GelPredd_col,nvmCalTOgeld] <- RnvmCald
  Aa[col0+GelPredd_col,chaetoTOgeld] <- RChaetod
  Aa[col0+GelPredd_col,poecilTOgeld] <- RPoecild
  Aa[col0+GelPredd_col,gelTOhtld] <- -RGelPredd
  Aa[col0+GelPredd_col,gelTOnh4d] <- -(RGelPredd+Eps_ExcL)
  Aa[col0+GelPredd_col,gelTOdomd] <- -(RGelPredd+Eps_ExcL)
  Aa[col0+GelPredd_col,gelTOldetd] <- -(RGelPreddfood+Eps_EgL)
  Aa[col0+Plankfishd_col,herbvmTOdplankfishm] <- RHerbVM
  Aa[col0+Plankfishd_col,vmCalTOdplankfishm] <- RvmCal
  Aa[col0+Plankfishd_col,herbnvmTOplankfishd] <- RHerbNVMd
  Aa[col0+Plankfishd_col,appTOplankfishd] <- RAppd
  Aa[col0+Plankfishd_col,cladTOplankfishd] <- RCladd
  Aa[col0+Plankfishd_col,nvmCalTOplankfishd] <- RnvmCald
  Aa[col0+Plankfishd_col,chaetoTOplankfishd] <- RChaetod
  Aa[col0+Plankfishd_col,poecilTOplankfishd] <- RPoecild
  Aa[col0+Plankfishd_col,plankfishTOhtld] <- -RPlankfishd
  Aa[col0+Plankfishd_col,plankfishTOnh4d] <- -(RPlankfishd+Eps_ExcL)
  Aa[col0+Plankfishd_col,plankfishTOdomd] <- -(RPlankfishd+Eps_ExcL)
  Aa[col0+Plankfishd_col,plankfishTOpoopd] <- -(RPlankfishdfood+Eps_EgL)
  Aa[col0+Bacd_col,bacTOflagd] <- -RBacd
  Aa[col0+Bacd_col,bacTOhnfd] <- -RBacd
  Aa[col0+Bacd_col,bacTOmicd] <- -RBacd
  Aa[col0+Bacd_col,bacTOappd] <- -RBacd
  Aa[col0+Bacd_col,bacTOnh4d] <- -(RBacd+Eps_ExcB)
  Aa[col0+Bacd_col,domTObacd] <- RDond+Eps_UpB
  Aa[col0+Detd_col,LateralInputPONd] <- RSDetInputd
  Aa[col0+Detd_col,CyanoTOsdetd] <- RCyad
  Aa[col0+Detd_col,flagTOsdetd] <- RFlagd
  Aa[col0+Detd_col,hnfTOsdetd] <- (RHNFdfood+Eps_EgS)
  Aa[col0+Detd_col,micTOsdetd] <- (RMICdfood+Eps_EgS)
  Aa[col0+Detd_col,sdetTOhnfd] <- -RDets
  Aa[col0+Detd_col,sdetTOmicd] <- -RDets
  Aa[col0+Detd_col,sdetTOappd] <- -RDets
  Aa[col0+Detd_col,sdetTOdomd] <- -(RDetd+Eps_Remin)
  Aa[col0+Ldetd_col,herbvmTOdldetm] <- (RHerbVMfood+Eps_EgL)
  Aa[col0+Ldetd_col,vmCalTOdldetm] <- (RvmCalfood+Eps_EgL)
  Aa[col0+Ldetd_col,dldetSinkm] <- RLdets
  Aa[col0+Ldetd_col,TrichoTOldetd] <- RTrid
  Aa[col0+Ldetd_col,dtmTOldetd] <- RDtmd
  Aa[col0+Ldetd_col,herbnvmTOldetd] <- (RHerbNVMdfood+Eps_EgL)
  Aa[col0+Ldetd_col,appTOldetd] <- (RAppdfood+Eps_EgL)
  Aa[col0+Ldetd_col,cladTOldetd] <- (RCladdfood+Eps_EgL)
  Aa[col0+Ldetd_col,nvmCalTOldetd] <- (RnvmCaldfood+Eps_EgL)
  Aa[col0+Ldetd_col,chaetoTOldetd] <- (RChaetodfood+Eps_EgL)
  Aa[col0+Ldetd_col,poecilTOldetd] <- (RPoecildfood+Eps_EgL)
  Aa[col0+Ldetd_col,gelTOldetd] <- (RGelPreddfood+Eps_EgL)
  Aa[col0+Ldetd_col,ldetTOherbnvmd] <- -RLdetd
  Aa[col0+Ldetd_col,ldetTOcladd] <- -RLdetd
  Aa[col0+Ldetd_col,ldetTOnvmCald] <- -RLdetd
  Aa[col0+Ldetd_col,ldetTOherbvmd] <- -RLdetd
  Aa[col0+Ldetd_col,ldetTOvmCald] <- -RLdetd
  Aa[col0+Ldetd_col,ldetTOdomd] <- -(RLdetd+Eps_Remin)
  Aa[col0+Ldetd_col,DlDetSinkd] <- -RLdetd
  Aa[col0+Dond_col,LateralInputDONd] <- RDONInputd
  Aa[col0+Dond_col,herbvmTOddomm] <- (RHerbVM+Eps_ExcL)
  Aa[col0+Dond_col,vmCalTOddomm] <- (RvmCal+Eps_ExcL)
  Aa[col0+Dond_col,CyanoTODOCd] <- RCyad
  Aa[col0+Dond_col,TrichoTOdomd] <- RTrid
  Aa[col0+Dond_col,dtmTOdomd] <- RDtmd
  Aa[col0+Dond_col,flagTOdomd] <- RFlagd
  Aa[col0+Dond_col,hnfTOdomd] <- (RHNFd+Eps_ExcS)
  Aa[col0+Dond_col,micTOdomd] <- (RMICd+Eps_ExcS)
  Aa[col0+Dond_col,herbnvmTOdomd] <- (RHerbNVMd+Eps_ExcL)
  Aa[col0+Dond_col,appTOdomd] <- (RAppd+Eps_ExcL)
  Aa[col0+Dond_col,cladTOdomd] <- (RCladd+Eps_ExcL)
  Aa[col0+Dond_col,nvmCalTOdomd] <- (RnvmCald+Eps_ExcL)
  Aa[col0+Dond_col,chaetoTOdomd] <- (RChaetod+Eps_ExcL)
  Aa[col0+Dond_col,poecilTOdomd] <- (RPoecild+Eps_ExcL)
  Aa[col0+Dond_col,gelTOdomd] <- (RGelPredd+Eps_ExcL)
  Aa[col0+Dond_col,plankfishTOdomd] <- (RPlankfishd+Eps_ExcL)
  Aa[col0+Dond_col,domTObacd] <- -(RDond+Eps_UpB)
  Aa[col0+Dond_col,sdetTOdomd] <- (RDetd+Eps_Remin)
  Aa[col0+Dond_col,ldetTOdomd] <- (RLdetd+Eps_Remin)
      
  Aa[col0+NO3s_col,] <- Aa[col0+NO3s_col,]/totflowNO3s
  Aa[col0+NH4s_col,] <- Aa[col0+NH4s_col,]/totflowNH4s
  Aa[col0+Cyas_col,] <- Aa[col0+Cyas_col,]/totflowCyas
  Aa[col0+Tris_col,] <- Aa[col0+Tris_col,]/totflowTris
  Aa[col0+Dtms_col,] <- Aa[col0+Dtms_col,]/totflowDtms
  Aa[col0+Flags_col,] <- Aa[col0+Flags_col,]/totflowFlags
  Aa[col0+HNFs_col,] <- Aa[col0+HNFs_col,]/totflowHNFs
  Aa[col0+MICs_col,] <- Aa[col0+MICs_col,]/totflowMICs
  Aa[col0+HerbNVMs_col,] <- Aa[col0+HerbNVMs_col,]/totflowHerbNVMs
  Aa[col0+Apps_col,] <- Aa[col0+Apps_col,]/totflowApps
  Aa[col0+Clads_col,] <- Aa[col0+Clads_col,]/totflowClads
  Aa[col0+nvmCals_col,] <- Aa[col0+nvmCals_col,]/totflownvmCals
  Aa[col0+Chaetos_col,] <- Aa[col0+Chaetos_col,]/totflowChaetos
  Aa[col0+Poecils_col,] <- Aa[col0+Poecils_col,]/totflowPoecils
  Aa[col0+GelPreds_col,] <- Aa[col0+GelPreds_col,]/totflowGelPreds
  Aa[col0+Plankfishs_col,] <- Aa[col0+Plankfishs_col,]/totflowPlankfishs
  Aa[col0+Preflexs_col,] <- Aa[col0+Preflexs_col,]/totflowPreflexs
  Aa[col0+Postflexs_col,] <- Aa[col0+Postflexs_col,]/totflowPostflexs
  Aa[col0+Bacs_col,] <- Aa[col0+Bacs_col,]/totflowBacs
  Aa[col0+Dets_col,] <- Aa[col0+Dets_col,]/totflowDets
  Aa[col0+Ldets_col,] <- Aa[col0+Ldets_col,]/totflowLdets
  Aa[col0+Dons_col,] <- Aa[col0+Dons_col,]/totflowDons
  Aa[col0+HerbVM_col,] <- Aa[col0+HerbVM_col,]/totflowHerbVM
  Aa[col0+vmCal_col,] <- Aa[col0+vmCal_col,]/totflowvmCal
  Aa[col0+NO3d_col,] <- Aa[col0+NO3d_col,]/totflowNO3d
  Aa[col0+NH4d_col,] <- Aa[col0+NH4d_col,]/totflowNH4d
  Aa[col0+Cyad_col,] <- Aa[col0+Cyad_col,]/totflowCyad
  Aa[col0+Trid_col,] <- Aa[col0+Trid_col,]/totflowTrid
  Aa[col0+Dtmd_col,] <- Aa[col0+Dtmd_col,]/totflowDtmd
  Aa[col0+Flagd_col,] <- Aa[col0+Flagd_col,]/totflowFlagd
  Aa[col0+HNFd_col,] <- Aa[col0+HNFd_col,]/totflowHNFd
  Aa[col0+MICd_col,] <- Aa[col0+MICd_col,]/totflowMICd
  Aa[col0+HerbNVMd_col,] <- Aa[col0+HerbNVMd_col,]/totflowHerbNVMd
  Aa[col0+Appd_col,] <- Aa[col0+Appd_col,]/totflowAppd
  Aa[col0+Cladd_col,] <- Aa[col0+Cladd_col,]/totflowCladd
  Aa[col0+nvmCald_col,] <- Aa[col0+nvmCald_col,]/totflownvmCald
  Aa[col0+Chaetod_col,] <- Aa[col0+Chaetod_col,]/totflowChaetod
  Aa[col0+Poecild_col,] <- Aa[col0+Poecild_col,]/totflowPoecild
  Aa[col0+GelPredd_col,] <- Aa[col0+GelPredd_col,]/totflowGelPredd
  Aa[col0+Plankfishd_col,] <- Aa[col0+Plankfishd_col,]/totflowPlankfishd
  Aa[col0+Bacd_col,] <- Aa[col0+Bacd_col,]/totflowBacd
  Aa[col0+Detd_col,] <- Aa[col0+Detd_col,]/totflowDetd
  Aa[col0+Ldetd_col,] <- Aa[col0+Ldetd_col,]/totflowLdetd
  Aa[col0+Dond_col,] <- Aa[col0+Dond_col,]/totflowDond
  
  for (i in 1:length(wts))  {
    Aa[,i] <- Aa[,i]/wts[i]
  }
  #print(Aa[13,10])
  return(Aa)
}


### Notify Function ###
pb = txtProgressBar(0, 1, style=3)
notify = function(i, n) {
  setTxtProgressBar(pb, i/n)
}
```
