## Supplementary material for "Comparative food-web analysis of bluefin tuna spawning habitats in the eastern Indian Ocean and Gulf of Mexico": Code Package: FindRanges.R

```
#Note that first you have to set the directory: Session>Set Working Directory
rm(list=ls())
library(R.matlab)
library(limSolve)
library(MASS)
source('xsampleN15outputs.r')
source('ExternalFunctions.r')
Cycle <- 1

tmp <- paste('N15GoMInverseCycle',Cycle,'.mat',sep="")
data <- readMat(tmp, fixNames=TRUE)
h <- data[['h']]
b <- data[['b']]
ba <- data[['ba']]
be <- data[['be']]
Ae <- data[['Ae']]
Aa <- data[['Aa']]
Aa0 <- data[['Aa0']]
G <- data[['G']]
sdba <- data[['sdba']]
InputCol <- data[['InputCol']]
A <- data[['A']]
Inputs <- data[['Inputs']]
d15NInputs <- data[['d15NInputs']]
wts <- data[['wts']]
#W <- rep(1,39)
BurninLength1 <- 10000
BurninLength2 <- 10000
IterLength <- 300000
OutputLength <- IterLength/1000
jmpLength <- 0.001
jmpLength15 <- 0.03

Aa2 <- Aa[1:36,]
ba2 <- ba[1:36]
sdba2 <- sdba[1:36]
#Aa2 <- Aa[2:3,]
#ba2 <- ba[2:3]
#sdba2 <- sdba[2:3]
#G <- G[1:50,]
#h <- h[1:50]

d15N0 <- c(0,0,0,0,0,0,0,0,0,0,
           0,0,0,0,0,0,0,0,0,0,
           0,0,0,0,0,0,0,0,0,0,
           0,0,0,0,0,0,0,0,0,0,
           0,0,0,0,0,0)
d15N0 <- d15N0 + d15NInputs[1,InputCol]


upNO315N=d15NInputs[1,InputCol]
pre15N=d15NInputs[2,InputCol]
post15N=d15NInputs[3,InputCol]
LDetS15N=d15NInputs[4,InputCol]
LDetD15N=d15NInputs[5,InputCol]
DONS15N=d15NInputs[6,InputCol]
DOND15N=d15NInputs[7,InputCol]
del15Nknown = c(upNO315N,pre15N,post15N,LDetS15N,LDetD15N,DONS15N,DOND15N)


#G <- G[76:183,]
#h <- h[76:183]

testranges <- xranges(E = Ae, F = be, G = G, H = h,
        ispos = FALSE, tol = 1e-8, central = FALSE, full=FALSE)


#L2MN Solution
test <- lsei(A = Aa2, B = ba2, E = Ae, F = be, G = G, H = h, type=2)
X <- test[['X']]
solNorm <- test[['solutionNorm']]
lseisol <- as.matrix(X)
rm(test, X)
fileout <- paste('N15GoMInverseLSEICycle',Cycle,'Routputs.mat',sep="")
writeMat(fileout, Aa = Aa2, ba = ba2, Ae = Ae, be = be, G = G, h = h, lseisol = lseisol, wts=wts)


#Central Value Solution
center <- xranges(E = Ae, F = be, G = G, H = h,
                  ispos = FALSE, tol = 1e-8, central = TRUE, full=FALSE)
centralval <- center[,3]
```
