## Supplementary material for "Comparative food-web analysis of bluefin tuna spawning habitats in the eastern Indian Ocean and Gulf of Mexico": Code Package: FoodwebIndices.m

```
clearvars
close all


for Cycle=[0,10]
    clearvars -except Cycle

    load(['N15GoMInverseCycle',num2str(Cycle),'.mat'])
    load(['N15GoMInverseCycle',num2str(Cycle),'Routputs.mat'])
    clearvars -except Ae0 MCMCmat Aa d15NInputs del15N InputCol wts Cycle
    num=length(MCMCmat(:,1));
    MCMCmat(1:round(num*0.2),:)=[];
    del15N(1:round(num*0.2),:)=[];
    len=length(MCMCmat(:,1));
    [num,txt,raw]  = xlsread('N15InverseModelRW.GOM.xlsx','Sheet1','B6:B307');
    Columns = txt;
    meanvals = mean(MCMCmat)'./wts;
    stdvals = std(MCMCmat)'./wts;
    meantable = table(Columns,meanvals,stdvals);

    [NO3s_col,NH4s_col,Cyas_col,Tris_col,Dtms_col,Flags_col,HNFs_col,MICs_col,HerbNVMs_col,Apps_col,Clads_col,nvmCals_col,Chaetos_col,Poecils_col,GelPreds_col,Plankfishs_col,Preflexs_col,Postflexs_col,Bacs_col,Dets_col,Ldets_col,Dons_col,HerbVM_col,vmCal_col,NO3d_col,NH4d_col,Cyad_col,Trid_col,Dtmd_col,Flagd_col,HNFd_col,MICd_col,HerbNVMd_col,Appd_col,Cladd_col,nvmCald_col,Chaetod_col,Poecild_col,GelPredd_col,Plankfishd_col,Bacd_col,Detd_col,Ldetd_col,Dond_col] = GetColumns(Aa,MCMCmat,del15N(1,:),d15NInputs(1,InputCol),d15NInputs(2,InputCol),d15NInputs(3,InputCol),d15NInputs(4,InputCol),d15NInputs(5,InputCol),d15NInputs(6,InputCol),d15NInputs(7,InputCol));
    ToMesozoos = sort([find(Ae0(HerbNVMs_col,:)==1),find(Ae0(Apps_col,:)==1),find(Ae0(Clads_col,:)==1),find(Ae0(nvmCals_col,:)==1),find(Ae0(Chaetos_col,:)==1),find(Ae0(Poecils_col,:)==1),find(Ae0(HerbVM_col,:)==1),find(Ae0(vmCal_col,:)==1)]);
    ToMesozood = sort([find(Ae0(HerbNVMd_col,:)==1),find(Ae0(Appd_col,:)==1),find(Ae0(Cladd_col,:)==1),find(Ae0(nvmCald_col,:)==1),find(Ae0(Chaetod_col,:)==1),find(Ae0(Poecild_col,:)==1),find(Ae0(HerbVM_col,:)==1),find(Ae0(vmCal_col,:)==1)]);
    FromMesozoos = sort([find(Ae0(HerbNVMs_col,:)==-1),find(Ae0(Apps_col,:)==-1),find(Ae0(Clads_col,:)==-1),find(Ae0(nvmCals_col,:)==-1),find(Ae0(Chaetos_col,:)==-1),find(Ae0(Poecils_col,:)==-1),find(Ae0(HerbVM_col,:)==-1),find(Ae0(vmCal_col,:)==-1)]);
    FromMesozood = sort([find(Ae0(HerbNVMd_col,:)==-1),find(Ae0(Appd_col,:)==-1),find(Ae0(Cladd_col,:)==-1),find(Ae0(nvmCald_col,:)==-1),find(Ae0(Chaetod_col,:)==-1),find(Ae0(Poecild_col,:)==-1),find(Ae0(HerbVM_col,:)==-1),find(Ae0(vmCal_col,:)==-1)]);
    FromHTLs = sort([find(Ae0(GelPreds_col,:)==-1),find(Ae0(Preflexs_col,:)==-1),find(Ae0(Postflexs_col,:)==-1),find(Ae0(Plankfishs_col,:)==-1)]);
    FromHTLd = sort([find(Ae0(HerbNVMd_col,:)==-1),find(Ae0(Plankfishd_col,:)==-1)]);
    FromPhys = sort([find(Ae0(Cyas_col,:)==-1),find(Ae0(Tris_col,:)==-1),find(Ae0(Dtms_col,:)==-1)]);
    FromPhyd = sort([find(Ae0(Cyad_col,:)==-1),find(Ae0(Trid_col,:)==-1),find(Ae0(Dtmd_col,:)==-1)]);
    FromMixs = find(Ae0(Flags_col,:)==-1);
    FromMixd = find(Ae0(Flagd_col,:)==-1);
    ToMixs = find(Ae0(Flags_col,:)==1);
    ToMixd = find(Ae0(Flagd_col,:)==1);
    FromCyas = find(Ae0(Cyas_col,:)==-1);
    FromCyad = find(Ae0(Cyad_col,:)==-1);
    ToCyas = find(Ae0(Cyas_col,:)==1);
    ToCyad = find(Ae0(Cyad_col,:)==1);
    FromNO3s = find(Ae0(NO3s_col,:)==-1);
    FromNO3d = find(Ae0(NO3d_col,:)==-1);
    FromNH4s = find(Ae0(NH4s_col,:)==-1);
    FromNH4d = find(Ae0(NH4d_col,:)==-1);
    ToNH4s = find(Ae0(NH4s_col,:)==1);
    ToNH4d = find(Ae0(NH4d_col,:)==1);
    ToDons = find(Ae0(Dons_col,:)==1);
    ToDond = find(Ae0(Dond_col,:)==1);
    FromBacs = find(Ae0(Bacs_col,:)==-1);
    FromBacd = find(Ae0(Bacd_col,:)==-1);
    FromProtists = sort([find(Ae0(HNFs_col,:)==-1),find(Ae0(MICs_col,:)==-1)]);
    FromProtistd = sort([find(Ae0(HNFd_col,:)==-1),find(Ae0(MICd_col,:)==-1)]);
    ToProtists = sort([find(Ae0(HNFs_col,:)==1),find(Ae0(MICs_col,:)==1)]);
    ToProtistd = sort([find(Ae0(HNFd_col,:)==1),find(Ae0(MICd_col,:)==1)]);
    FromDtms = find(Ae0(Dtms_col,:)==-1);
    FromDtmd = find(Ae0(Dtmd_col,:)==-1);
    ToDtms = find(Ae0(Dtms_col,:)==1);
    ToDtmd = find(Ae0(Dtmd_col,:)==1);
    FromTris = find(Ae0(Tris_col,:)==-1);
    FromTrid = find(Ae0(Trid_col,:)==-1);
    ToTris = find(Ae0(Tris_col,:)==1);
    ToTrid = find(Ae0(Trid_col,:)==1);

    PHYtoMESOZOOs = intersect(ToMesozoos,FromPhys);
    PHYtoMESOZOOd = intersect(ToMesozood,FromPhyd);
    FLAGtoMESOZOOs = intersect(ToMesozoos,FromMixs);
    FLAGtoMESOZOOd = intersect(ToMesozood,FromMixd);
    BACtoFLAGs = intersect(FromBacs,ToMixs);
    BACtoFLAGd = intersect(FromBacd,ToMixd);
    CYAtoFLAGs = intersect(FromCyas,ToMixs);
    CYAtoFLAGd = intersect(FromCyad,ToMixd);
    NUTtoFLAGs = intersect([FromNO3s,FromNH4s],ToMixs);
    NUTtoFLAGd = intersect([FromNO3d,FromNH4d],ToMixd);
    FLAGtoDONs = intersect(FromMixs,ToDons);
    FLAGtoDONd = intersect(FromMixd,ToDond);
    CYAtoDONs = intersect(FromCyas,ToDons);
    CYAtoDONd = intersect(FromCyad,ToDond);
    DTMtoDONs = intersect(FromDtms,ToDons);
    DTMtoDONd = intersect(FromDtmd,ToDond);
    TRICHOtoDONs = intersect(FromTris,ToDons);
    TRICHOtoDONd = intersect(FromTrid,ToDond);
    NUTtoDTMs = intersect([FromNO3s,FromNH4s],ToDtms);
    NUTtoDTMd = intersect([FromNO3d,FromNH4d],ToDtmd);
    DTMtoDONs = intersect(FromDtms,ToDons);
    DTMtoDONd = intersect(FromDtmd,ToDond);
    NUTtoCYAs = intersect([FromNO3s,FromNH4s],ToCyas);
    NUTtoCYAd = intersect([FromNO3d,FromNH4d],ToCyad);
    FLAGtoCYAs = intersect(FromCyas,ToDons);
    FLAGtoCYAd = intersect(FromCyad,ToDond);
    NUTtoTRICHOs = intersect([FromNO3s,FromNH4s],ToTris);
    NUTtoTRICHOd = intersect([FromNO3d,FromNH4d],ToTrid);
    FLAGtoTRICHOs = intersect(FromTris,ToDons);
    FLAGtoTRICHOd = intersect(FromTrid,ToDond);
    PROTISTtoMESOZOOs = intersect(ToMesozoos,FromProtists);
    PROTISTtoMESOZOOd = intersect(ToMesozood,FromProtistd);
    BACtoNH4s = intersect(FromBacs,ToNH4s);
    BACtoNH4d = intersect(FromBacd,ToNH4d);
    BACtoPROTISTs = intersect(FromBacs,ToProtists);
    BACtoPROTISTd = intersect(FromBacd,ToProtistd);
    PROTISTtoDONs = intersect(FromProtists,ToDons);
    PROTISTtoDONd = intersect(FromProtistd,ToDond);
    PROTISTtoNH4s = intersect(FromProtists,ToNH4s);
    PROTISTtoNH4d = intersect(FromProtistd,ToNH4d);
    NO3toPHYs = intersect(FromNO3s,[ToDtms,ToCyas,ToTris,ToMixs]);
    NO3toPHYd = intersect(FromNO3d,[ToDtmd,ToCyad,ToTris,ToMixd]);
    NH4toPHYs = intersect(FromNH4s,[ToDtms,ToCyas,ToTris,ToMixs]);
    NH4toPHYd = intersect(FromNH4d,[ToDtmd,ToCyad,ToTris,ToMixd]);
    PHYtoPROTIST = intersect([FromPhys,FromMixs,FromPhyd,FromMixd],[ToProtists,ToProtistd]);
    PHYtoDONs = intersect(FromPhys,ToDons);
    MIXtoDONs = intersect(FromMixs,ToDons);
    MESOZOOtoDONs = intersect(FromMesozoos,ToDons);
    HTLtoDONs = intersect(FromHTLs,ToDons);
    BACtoNH4s = intersect(FromBacs,ToNH4s);
    MIXtoNH4s = intersect(FromMixs,ToNH4s);
    MESOZOOtoNH4s = intersect(FromMesozoos,ToNH4s);
    HTLtoNH4s = intersect(FromHTLs,ToNH4s);


    for i=1:len
        i
        flows=MCMCmat(i,:)./wts';

        CYAnpps(i) = sum(flows(ToCyas)) - flows(CYAtoDONs);
        TRInpps(i) = sum(flows(ToTris)) - flows(TRICHOtoDONs);
        DTMnpps(i) = sum(flows(NUTtoDTMs)) - flows(DTMtoDONs);
        FLAGnpps(i) = sum(flows(NUTtoFLAGs)) - flows(FLAGtoDONs);
        CYAnppd(i) = sum(flows(ToCyad)) - flows(CYAtoDONd);
        TRInppd(i) = sum(flows(ToTrid)) - flows(TRICHOtoDONd);
        DTMnppd(i) = sum(flows(NUTtoDTMd)) - flows(DTMtoDONd);
        FLAGnppd(i) = sum(flows(NUTtoFLAGd)) - flows(FLAGtoDONd);

        FLAGmixs = flows(BACtoFLAGs)+flows(CYAtoFLAGs);
        HerbFoodChains(i) = sum(flows(PHYtoMESOZOOs)) + sum(flows(FLAGtoMESOZOOs))*FLAGnpps(i)/(FLAGnpps(i)+FLAGmixs);
        FLAGmixd = flows(BACtoFLAGd)+flows(CYAtoFLAGd);
        HerbFoodChaind(i) = sum(flows(PHYtoMESOZOOd)) + sum(flows(FLAGtoMESOZOOd))*FLAGnppd(i)/(FLAGnppd(i)+FLAGmixd);

        MultivFoodChains(i) = sum(flows(PROTISTtoMESOZOOs)) + sum(flows(FLAGtoMESOZOOs))*sum(flows(CYAtoFLAGs))/(FLAGnpps(i)+FLAGmixs);
        MultivFoodChaind(i) = sum(flows(PROTISTtoMESOZOOd)) + sum(flows(FLAGtoMESOZOOd))*sum(flows(CYAtoFLAGd))/(FLAGnppd(i)+FLAGmixd);

        ProtistRespirationPercents = sum(flows(PROTISTtoNH4s))/sum(flows(ToProtists));
        MicrobialLoops(i) = flows(BACtoNH4s) + sum(flows(BACtoPROTISTs))*ProtistRespirationPercents;
        ProtistRespirationPercentd = sum(flows(PROTISTtoNH4d))/sum(flows(ToProtistd));
        MicrobialLoopd(i) = flows(BACtoNH4d) + sum(flows(BACtoPROTISTd))*ProtistRespirationPercentd;

        NPPs(i) = CYAnpps(i) + TRInpps(i) + DTMnpps(i) + FLAGnpps(i);
        NPPd(i) = CYAnppd(i) + TRInppd(i) + DTMnppd(i) + FLAGnppd(i);

        NO3UPTAKEs(i) = sum(flows(NO3toPHYs));
        NO3UPTAKEd(i) = sum(flows(NO3toPHYd));
        NH4UPTAKEs(i) = sum(flows(NH4toPHYs));
        NH4UPTAKEd(i) = sum(flows(NH4toPHYd));
        N2FIXs(i) = sum(flows([ToTris,ToCyas])) - sum(flows([NUTtoCYAs,NUTtoTRICHOs]));
        N2FIXd(i) = sum(flows([ToTrid,ToCyad])) - sum(flows([NUTtoCYAd,NUTtoTRICHOd]));

        PROTISTGRAZING(i) = sum(flows(PHYtoPROTIST));
        %MESOZOOGRAZING(i) = sum(flows(PHYtoMESOZOOs)) + sum(flows(FLAGtoMESOZOOs)) + sum(flows(PHYtoMESOZOOd)) + sum(flows(FLAGtoMESOZOOd));

        PhytoplanktonDONproductions(i)=sum(flows(PHYtoDONs)) + sum(flows(MIXtoDONs))*FLAGnpps(i)/(FLAGnpps(i)+FLAGmixs);
        ProtistDONproductions(i)=sum(flows(PROTISTtoDONs)) + sum(flows(MIXtoDONs))*FLAGmixs/(FLAGnpps(i)+FLAGmixs);
        MesozooDONproductions(i)=sum(flows(MESOZOOtoDONs));
        HTLDONproductions(i)=sum(flows(HTLtoDONs));
        TotalDONproductions(i)=sum(flows(ToDons));

        BacNH4productions(i)=sum(flows(BACtoNH4s));
        ProtistNH4productions(i)=sum(flows(PROTISTtoNH4s)) + sum(flows(MIXtoNH4s))*FLAGmixs/(FLAGnpps(i)+FLAGmixs);
        MesozooNH4productions(i)=sum(flows(MESOZOOtoNH4s));
        HTLNH4productions(i)=sum(flows(HTLtoNH4s));
        TotalNH4productions(i)=sum(flows(ToNH4s));
    end

    %---------------------------------------------------------------
    %------------Equivalent of Figure from Original Paper-----------
    %---------------------------------------------------------------


colVertex1 = [0 0.6 0];
colVertex2 = [0.7 0 1];
colVertex3 = [0.4 0.3 0];

    %-------------------------Raw Flux--------------------------------
    for j=1:2
        if j==1
            if Cycle==0
                fighandle = figure(1);
                fighandle.Units = 'inches';
                fighandle.Position = [1 1 6.5 6.5];
                subplot(2,2,1)
            else
                figure(1)
                subplot(2,2,3)
            end
        else
            if Cycle==0
                fighandle = figure(2);
                fighandle.Units = 'inches';
                fighandle.Position = [2.6 1 6.5 3];
                subplot(1,2,1)
            else
                figure(2)
                subplot(1,2,2)
            end
        end
        hold on
        ViolinPlot(1,0.5,HerbFoodChains,200,colVertex1)
        ViolinPlot(2,0.5,MultivFoodChains,200,colVertex2)
        ViolinPlot(3,0.5,MicrobialLoops,200,colVertex3)
        if Cycle==0
            plot([4,4],[0,3],'-k')
        else
            plot([4,4],[0,4],'-k')
        end

        ViolinPlot(5,0.5,HerbFoodChaind,200,colVertex1)
        ViolinPlot(6,0.5,MultivFoodChaind,200,colVertex2)
        ViolinPlot(7,0.5,MicrobialLoopd,200,colVertex3)
        set(gca,'box','on')
        set(gca,'XTick',[2,6])
        set(gca,'XTickLabel',{['Shallow Eup Zone'],'Deep Chl Max'})
        set(gca,'XTickLabelRotation',20)
        ylabel('Flux (mmol N m^-^2 d^-^1)')
        set(gca,'FontSize',9)

        if j==2 & Cycle==10
            subplot(1,2,1)
            AddPanelLabels('b','bottomright')
            AddPanelLabels('a','bottomright','YPADDING',0.05,'XPADDING',0.54)
            subplot(1,2,2)
            AddPanelLabels('d','bottomright')
            AddPanelLabels('c','bottomright','YPADDING',0.05,'XPADDING',0.54)
        end

        if Cycle==10 & j==2
            fill([4.1 4.1 7.9 7.9 4.1],[2.75 3.95 3.95 2.75 2.75],'w')
            h=fill([4.2 4.2 4.4 4.4 4.2],[3.6 3.9 3.9 3.6 3.6],'w')
            set(h,'FaceColor',colVertex1)
            text(4.5,3.75,'Herbiv Food Chain','FontSize',8)
            h=fill([4.2 4.2 4.4 4.4 4.2],[3.5 3.2 3.2 3.5 3.5],'w')
            set(h,'FaceColor',colVertex2)
            text(4.5,3.35,'Multiv Food Chain','FontSize',8)
            h=fill([4.2 4.2 4.4 4.4 4.2],[3.1 2.8 2.8 3.1 3.1],'w')
            set(h,'FaceColor',colVertex3)
            text(4.5,2.95,'Microbial Loop','FontSize',8)
        end
        if Cycle==0
            title('Gulf of Mexico')
        else
            title('Argo Basin')
        end
    end

    %--------------------Normalized Flux---------------------------------
    if Cycle==0
        figure(1)
        subplot(2,2,2)
    else
        figure(1)
        subplot(2,2,4)
    end
    hold on
    ViolinPlot(1,0.5,HerbFoodChains./NPPs,200,colVertex1)
    ViolinPlot(2,0.5,MultivFoodChains./NPPs,200,colVertex2)
    ViolinPlot(3,0.5,MicrobialLoops./NPPs,200,colVertex3)
    plot([4,4],[0,3],'-k')
    ViolinPlot(5,0.5,HerbFoodChaind./NPPd,200,colVertex1)
    ViolinPlot(6,0.5,MultivFoodChaind./NPPd,200,colVertex2)
    ViolinPlot(7,0.5,MicrobialLoopd./NPPd,200,colVertex3)
    if Cycle==10
        fill([4.1 4.1 7.9 7.9 4.1],[0.94 1.19 1.19 0.94 0.94],'w')
        h=fill([4.2 4.2 4.4 4.4 4.2],[1.18 1.11 1.11 1.18 1.18],'w')
        set(h,'FaceColor',colVertex1)
        text(4.5,1.15,'Herbiv Food Chain','FontSize',8)
        h=fill([4.2 4.2 4.4 4.4 4.2],[1.1 1.03 1.03 1.1 1.1],'w')
        set(h,'FaceColor',colVertex2)
        text(4.5,1.07,'Multiv Food Chain','FontSize',8)
        h=fill([4.2 4.2 4.4 4.4 4.2],[1.02 0.95 0.95 1.02 1.02],'w')
        set(h,'FaceColor',colVertex3)
        text(4.5,0.98,'Microbial Loop','FontSize',8)
    end
    set(gca,'box','on')
    set(gca,'XTick',[2,6])
    set(gca,'XTickLabel',{['Shallow Eup Zone'],'Deep Chl Max'})
    %set(gca,'XTickLabelRotation',45)
    ylabel('Flux / NPP')
    set(gca,'FontSize',9)
    ylim([0 1.2])
    if Cycle==0
        title('Gulf of Mexico')
    else
        title('Argo Basin')
    end


    %---------------------------------------------------------------
    %------------Ternary Plot---------------------------------------
    %---------------------------------------------------------------
    

    if Cycle==0
        fighandle = figure(101);
        fighandle.Units = 'inches';
        fighandle.Position = [1 1 3.5 3.5];
        col = [0 0 1];
    else
        figure(101)
        col = [1 0 0];
    end
    %TernaryPlot(Vertex1,Vertex2,Vertex3,Labels,'Circle',[1 0 0],10,12,9)
    TernaryPlot(mean(HerbFoodChains),mean(MultivFoodChains),mean(MicrobialLoops),{'Herbivorous','Multivorous','Microbial'},'Circle',col,10,12,9,0,1)
    hold on
    if Cycle==0
        TernaryPlot(mean(HerbFoodChaind),mean(MultivFoodChaind),mean(MicrobialLoopd),{'Herbivorous','Multivorous','Microbial'},'Diamond',col,10,12,9,0,1)
    else
        TernaryPlot(mean(HerbFoodChaind),mean(MultivFoodChaind),mean(MicrobialLoopd),{'Herbivorous','Multivorous','Microbial'},'Diamond',col,10,12,9,1,1)
    end


    %---------------------------------------------------------------
    %------------Ternary Plot with dots---------------------------------------
    %---------------------------------------------------------------
    

    if Cycle==0
        fighandle = figure(102);
        fighandle.Units = 'inches';
        fighandle.Position = [1 5 3.5 3.5];
        col = [0 0 1];
    else
        figure(102)
        col = [1 0 0];
    end
    %TernaryPlot(Vertex1,Vertex2,Vertex3,Labels,'Circle',[1 0 0],10,12,9)
    TernaryPlot(HerbFoodChains,MultivFoodChains,MicrobialLoops,{'Herbivorous','Multivorous','Microbial'},'Dot',col,2,12,9,0,0.1)
    hold on
    % if Cycle==0
    %     TernaryPlot(mean(HerbFoodChaind),mean(MultivFoodChaind),mean(MicrobialLoopd),{'Herbivorous','Multivorous','Microbial'},'Diamond',col,10,12,9,0)
    % else
    %     TernaryPlot(mean(HerbFoodChaind),mean(MultivFoodChaind),mean(MicrobialLoopd),{'Herbivorous','Multivorous','Microbial'},'Diamond',col,10,12,9,1)
    % end


    h=fill([0 0.05 0.05 0 0]-0.7,[0 0 0.1 0.1 0]+0.9,'b')
    text(-0.62,0.95,['Gulf of',char(10),'Mexico'],'FontSize',9)
    h=fill([0 0.05 0.05 0 0]-0.7,[0 0 0.1 0.1 0]+0.9-0.2,'r')
    text(-0.62,0.95-0.2,['Argo',char(10),'Basin'],'FontSize',9)


    NH4UPTAKEpercents = sort(NH4UPTAKEs./(NH4UPTAKEs+NO3UPTAKEs+N2FIXs))*100;
    NO3UPTAKEpercents = sort(NO3UPTAKEs./(NH4UPTAKEs+NO3UPTAKEs+N2FIXs))*100;
    N2FIXpercents = sort(N2FIXs./(NH4UPTAKEs+NO3UPTAKEs+N2FIXs))*100;
    ['NH4 was the dominant source of nitrogen to phytoplankton',char(10),'(mean = ',num2str(mean(NH4UPTAKEpercents),2),...
        '%; 95% C.I. = ',num2str(NH4UPTAKEpercents(round(0.025*len)),2),' - ',num2str(NH4UPTAKEpercents(round(0.975*len)),2),...
        '%) in the shallow eup zone (0 - 50 m)']

    ['NO3 uptake (',num2str(mean(NO3UPTAKEpercents),2),...
        '%; ',num2str(NO3UPTAKEpercents(round(0.025*len)),2),' - ',num2str(NO3UPTAKEpercents(round(0.975*len)),2),...
        '%) and N2 fixation (',num2str(mean(N2FIXpercents),2),...
        '%; ',num2str(N2FIXpercents(round(0.025*len)),2),' - ',num2str(N2FIXpercents(round(0.975*len)),2),...
        '%)',char(10),'were comparatively less important']

    NH4UPTAKEpercentd = sort(NH4UPTAKEd./(NH4UPTAKEd+NO3UPTAKEd+N2FIXd))*100;
    NO3UPTAKEpercentd = sort(NO3UPTAKEd./(NH4UPTAKEd+NO3UPTAKEd+N2FIXd))*100;
    N2FIXpercentd = sort(N2FIXd./(NH4UPTAKEd+NO3UPTAKEd+N2FIXd))*100;
    ['NH4 was also dominant in the deep euphotic zone',char(10),'(',num2str(mean(NH4UPTAKEpercentd),2),...
        '%; ',num2str(NH4UPTAKEpercentd(round(0.025*len)),2),' - ',num2str(NH4UPTAKEpercentd(round(0.975*len)),2),...
        '%) in the deep chlorophyll max range (>50 m)']
    ['NO3 uptake (',num2str(mean(NO3UPTAKEpercentd),2),...
        '%; ',num2str(NO3UPTAKEpercentd(round(0.025*len)),2),' - ',num2str(NO3UPTAKEpercentd(round(0.975*len)),2),...
        '%) and N2 fixation (',num2str(mean(N2FIXpercentd),2),...
        '%; ',num2str(N2FIXpercentd(round(0.025*len)),2),' - ',num2str(N2FIXpercentd(round(0.975*len)),2),...
        '%)',char(10),'were comparatively less important']

    NPPs = sort(NPPs);
    NPPd = sort(NPPd);
    ['Mean NPP in the shallow euphotic zone was (',num2str(mean(NPPs),2),'; ',num2str(NPPs(round(0.025*len)),2),...
        ' - ',num2str(NPPs(round(0.975*len)),2),')']
    ['Mean NPP in the deep euphotic zone was (',num2str(mean(NPPd),2),'; ',num2str(NPPd(round(0.025*len)),2),...
        ' - ',num2str(NPPd(round(0.975*len)),2),')']

    DTMNPPpercents = sort(DTMnpps./NPPs)*100;
    FLAGNPPpercents = sort(FLAGnpps./NPPs)*100;
    CYANPPpercents = sort(CYAnpps./NPPs)*100;
    TRICHONPPpercents = sort(TRInpps./NPPs)*100;
    ['Picophytoplankton NPP was ',num2str(mean(CYANPPpercents),2),'% (',num2str(CYANPPpercents(0.025*len),2),' - ',num2str(CYANPPpercents(0.975*len),2),'% in the shallow eup zone)']
    ['Flagellate NPP was ',num2str(mean(FLAGNPPpercents),2),'% (',num2str(FLAGNPPpercents(0.025*len),2),' - ',num2str(FLAGNPPpercents(0.975*len),2),'% in the shallow eup zone)']
    ['Diatom NPP was ',num2str(mean(DTMNPPpercents),2),'% (',num2str(DTMNPPpercents(0.025*len),2),' - ',num2str(DTMNPPpercents(0.975*len),2),'% in the shallow eup zone)']


    DTMNPPpercentd = sort(DTMnpps./NPPd)*100;
    FLAGNPPpercentd = sort(FLAGnpps./NPPd)*100;
    CYANPPpercentd = sort(CYAnpps./NPPd)*100;
    TRICHONPPpercentd = sort(TRInpps./NPPd)*100;
    ['Picophytoplankton NPP was ',num2str(mean(CYANPPpercentd),2),'% (',num2str(CYANPPpercentd(0.025*len),2),' - ',num2str(CYANPPpercentd(0.975*len),2),'% in the deep eup zone)']
    ['Flagellate NPP was ',num2str(mean(FLAGNPPpercentd),2),'% (',num2str(FLAGNPPpercentd(0.025*len),2),' - ',num2str(FLAGNPPpercentd(0.975*len),2),'% in the shallow deep zone)']
    ['Diatom NPP was ',num2str(mean(DTMNPPpercentd),2),'% (',num2str(DTMNPPpercentd(0.025*len),2),' - ',num2str(DTMNPPpercentd(0.975*len),2),'% in the shallow deep zone)']

    ProtistGrazingPercent = sort(PROTISTGRAZING./(NPPs+NPPd))*100;
    ['Protists (including mixotrophic flagellates) consumed ',num2str(mean(ProtistGrazingPercent),2),'% (',num2str(ProtistGrazingPercent(round(0.025*len)),2),' - ',num2str(ProtistGrazingPercent(round(0.975*len)),2),'%) of phytoplankton production.']
    MesozooGrazingPercent = sort((HerbFoodChains+HerbFoodChaind)./(NPPs+NPPd))*100;
    ['Metazoan zooplankton consumed ',num2str(mean(MesozooGrazingPercent),2),'% (',num2str(MesozooGrazingPercent(round(0.025*len)),2),' - ',num2str(MesozooGrazingPercent(round(0.975*len)),2),'%) of phytoplankton production.']


    HerbFoodChainPercents=sort(HerbFoodChains./NPPs)*100;
    MultivFoodChainPercents=sort(MultivFoodChains./NPPs)*100;
    MicrobialLoopPercents=sort(MicrobialLoops./NPPs)*100;
    HerbFoodChainPercentd=sort(HerbFoodChaind./NPPd)*100;
    MultivFoodChainPercentd=sort(MultivFoodChaind./NPPd)*100;
    MicrobialLoopPercentd=sort(MicrobialLoopd./NPPd)*100;
    ['The microbial loop processed ',num2str(mean(MicrobialLoopPercents),2),'% (',num2str(MicrobialLoopPercents(round(0.025*len)),2),' - ',num2str(MicrobialLoopPercents(round(0.975*len)),3),'%) of net primary production',char(10),'in the shallow eup zone and ',...
        num2str(mean(MicrobialLoopPercentd),2),'% (',num2str(MicrobialLoopPercentd(round(0.025*len)),2),' - ',num2str(MicrobialLoopPercentd(round(0.975*len)),2),'%) of net primary production in the deep eup']

    ['The herbivorous food chain processed ',num2str(mean(HerbFoodChainPercents),2),'% and the multivorous food chain processed ',num2str(mean(MultivFoodChainPercents),2),'%',char(10),'of NPP in the shallow eup zone',...
        ' and ',num2str(mean(HerbFoodChainPercentd),2),'% and ',num2str(mean(MultivFoodChainPercentd),2),'%, respectively, of NPP in the deep eup zone']


    PhytoplanktonDONproductionpercents = sort(PhytoplanktonDONproductions./TotalDONproductions)*100;
    ProtistDONproductionpercents = sort(ProtistDONproductions./TotalDONproductions)*100;
    MesozooDONproductionpercents = sort(MesozooDONproductions./TotalDONproductions)*100;
    HTLDONproductionpercents = sort(HTLDONproductions./TotalDONproductions)*100;
    TotalDONproductions = sort(TotalDONproductions./TotalDONproductions)*100;
    ['In the shallow euphotic zone, phytoplankton exudation was',char(10),'responsible for ',num2str(mean(PhytoplanktonDONproductionpercents),2),'% of DON production, compared to',char(10),...
        num2str(mean(ProtistDONproductionpercents),2),'% produced by protistan zooplankton excretion, and ',char(10),num2str(mean(MesozooDONproductionpercents),2),'% by mesozoo excretion']


    BacNH4productionpercents = sort(BacNH4productions./TotalNH4productions)*100;
    ProtistNH4productionpercents = sort(ProtistNH4productions./TotalNH4productions)*100;
    MesozooNH4productionpercents = sort(MesozooNH4productions./TotalNH4productions)*100;
    HTLNH4productionpercents = sort(HTLNH4productions./TotalNH4productions)*100;
    ['In the shallow euphotic zone, bacterial excretion was',char(10),'responsible for ',num2str(mean(BacNH4productionpercents),2),'% of NH4 regeneration, compared to',char(10),...
        num2str(mean(ProtistNH4productionpercents),2),'% produced by protistan zooplankton excretion, and ',char(10),num2str(mean(MesozooNH4productionpercents),2),'% by mesozoo excretion']


    fn = 'ManuscriptValues.txt'
    fileID = fopen(fn,'a');
    if Cycle==0
        formatSpec =  '\n FoodwebIndices.m \n ';
        fprintf(fileID,formatSpec)
    end
    formatSpec =  'For Cycle %f in the UEZ the microbial loop processed %f +/- %f \n the multivorous food web processed %f +/- %f \n and the herbivorous food chain processed %f +/- %f mmol N m-2 d-1 \n';
    fprintf(fileID,formatSpec,[Cycle, mean(MicrobialLoops), std(MicrobialLoops), mean(MultivFoodChains), std(MultivFoodChains), mean(HerbFoodChains), std(HerbFoodChains)])
    fclose(fileID)

end

figure(1)
fn = ['FoodwebIndices.AllPanels'];
exportgraphics(gcf,['../../Figures/',fn,'.png'])
figure(2)
fn = ['FoodwebIndices.FluxOnly'];
exportgraphics(gcf,['../../Figures/',fn,'.png'])
figure(101)
fn = ['FoodwebIndices.Ternary'];
exportgraphics(gcf,['../../Figures/',fn,'.png'])
figure(102)
fn = ['FoodwebIndices.UEZonly'];
exportgraphics(gcf,['../../Figures/',fn,'.png'])
```
