## Supplementary material for "Comparative food-web analysis of bluefin tuna spawning habitats in the eastern Indian Ocean and Gulf of Mexico": Code Package: FoodWebIndirectFigure.m

```
function [output] = FoodWebIndirectFigure(Taxon,PerHerb,PerCal,PerClad,PerApp,PerPoecil)

Cycle=1

load(['N15GoMInverseCycle',num2str(Cycle),'.mat'])
load(['N15GoMInverseCycle',num2str(Cycle),'Routputs.mat'])


MCMC=mean(MCMCmat)./wts';

[NO3s_col,NH4s_col,Cyas_col,Tris_col,Dtms_col,Flags_col,HNFs_col,MICs_col,HerbNVMs_col,Apps_col,Clads_col,nvmCals_col,Chaetos_col,Poecils_col,GelPreds_col,Plankfishs_col,Preflexs_col,Postflexs_col,Bacs_col,Dets_col,Ldets_col,Dons_col,HerbVM_col,vmCal_col,NO3d_col,NH4d_col,Cyad_col,Trid_col,Dtmd_col,Flagd_col,HNFd_col,MICd_col,HerbNVMd_col,Appd_col,Cladd_col,nvmCald_col,Chaetod_col,Poecild_col,GelPredd_col,Plankfishd_col,Bacd_col,Detd_col,Ldetd_col,Dond_col] = GetColumns(Aa,MCMCmat,del15N(1,:),d15NInputs(1,InputCol),d15NInputs(2,InputCol),d15NInputs(3,InputCol),d15NInputs(4,InputCol),d15NInputs(5,InputCol),d15NInputs(6,InputCol),d15NInputs(7,InputCol));
bkgd=imread(['..\..\Figures\FoodWeb2BFT\Food Web to BFT-2 ',Taxon,'.png']);
piemask=imread('..\..\Figures\FoodWeb2BFT\PieMask2.jpg');
piemask_Lg=imread('..\..\Figures\FoodWeb2BFT\PieMask2_Lg.jpg');


ToMix=find(Ae(Flags_col,:)==1);
ToMic=find(Ae(MICs_col,:)==1);
ToHnf=find(Ae(HNFs_col,:)==1);
ToHerbNVM=find(Ae(HerbNVMs_col,:)==1);
ToApp=find(Ae(Apps_col,:)==1);
ToClad=find(Ae(Clads_col,:)==1);
TonvmCal=find(Ae(nvmCals_col,:)==1);
ToPoecil=find(Ae(Poecils_col,:)==1);
ToBft=find(Ae(Postflexs_col,:)==1);
ToHerbVM=find(Ae(HerbVM_col,:)==1);
TovmCal=find(Ae(vmCal_col,:)==1);
ToHerb=[ToHerbNVM,ToHerbVM];
ToCal=[TovmCal,TonvmCal];

FromDtm=find(Ae(Dtms_col,:)==-1);
FromHBac=find(Ae(Bacs_col,:)==-1);
FromCya=find(Ae(Cyas_col,:)==-1);
FromMix=find(Ae(Flags_col,:)==-1);
FromMic=find(Ae(MICs_col,:)==-1);
FromHnf=find(Ae(HNFs_col,:)==-1);
FromHerbNVM=find(Ae(HerbNVMs_col,:)==-1);
FromHerbVM=find(Ae(HerbVM_col,:)==-1);
FromnvmCal=find(Ae(nvmCals_col,:)==-1);
FromvmCal=find(Ae(vmCal_col,:)==-1);
FromClad=find(Ae(Clads_col,:)==-1);
FromApp=find(Ae(Apps_col,:)==-1);
FromPoecil=find(Ae(Poecils_col,:)==-1);
FromHerb=[FromHerbNVM,FromHerbVM];
FromCal=[FromvmCal,FromnvmCal];
FromNO3=find(Ae(NO3s_col,:)==-1);
FromNH4=find(Ae(NH4s_col,:)==-1);
FromNuts=[FromNO3,FromNH4];


HBac2MixArrow=[880,660;880,560];
Cyano2MixArrow=[600,660;800,560];
HBac2MicArrow=[800,660;540,530];
Cyano2MicArrow=[520,660;520,530];
Dtm2MicArrow=[350,660;500,530];
Mix2MicArrow=[680,530;575,495];
Hnf2MicArrow=[140,490;450,490];
Cyano2HnfArrow=[460,660;140,530];
HBac2HnfArrow=[680,665;140,520];
Dtm2AppArrow = [170,660;170,350];
Cyano2AppArrow = [480,660;180,350];
HBac2AppArrow = [680,660;190,350];
Hnf2AppArrow = [90,470;90,350];
Mix2AppArrow = [720,500;230,330];
Dtm2CladArrow = [200,660;350,350];
Hnf2CladArrow = [100,470;340,350];
Mic2CladArrow = [470,465;370,350];
Mix2CladArrow = [790,500;420,350];
Dtm2CalArrow = [240,660;590,350];
Hnf2CalArrow = [120,470;580,340];
Mic2CalArrow = [550,465;610,350];
Mix2CalArrow = [810,500;630,350];
Dtm2HerbArrow = [400,660;800,380];
Hnf2HerbArrow = [140,480;780,350];
Mic2HerbArrow = [580,465;780,365];
Mix2HerbArrow = [850,500;850,390];
App2PoecilArrow = [200,280;700,200];
Clad2PoecilArrow = [440,280;730,210];
Cal2PoecilArrow = [660,280;800,210];
Herb2PoecilArrow = [900,320;850,210];
Mix2HerbArrow = [850,500;850,390];
App2BftArrow = [150,280;460,190];
Clad2BftArrow = [380,280;490,190];
Cal2BftArrow = [620,280;560,190];
Herb2BftArrow = [820,320;600,190];
Mic2BftArrow = [530,465;530,190];
Poecil2BftArrow = [710,170;610,150];

DtmColor=[63,95,41]/256;
CyaColor=[255,175,6]/256;
HBacColor=[80,37,112]/256;
MixColor=[210,238,192]/256;
MicColor=[72,72,72]/256;
HnfColor=[168,208,244]/256;
HerbColor=[5,188,254]/256;
CalColor=[204,6,6]/256;
CladColor=[255,255,32]/256;
AppColor=[121,54,170]/256;
PoecilColor=[255,134,138]/256;
BftColor=[51,91,160]/256;


HBac2Mix=MCMC(intersect(FromHBac,ToMix));
Cya2Mix=MCMC(intersect(FromCya,ToMix));
Nut2Mix=sum(MCMC(intersect(FromNuts,ToMix)));

HBac2Mic=MCMC(intersect(FromHBac,ToMic));
Cya2Mic=MCMC(intersect(FromCya,ToMic));
Dtm2Mic=MCMC(intersect(FromDtm,ToMic));
Mix2Mic=MCMC(intersect(FromMix,ToMic));
Hnf2Mic=MCMC(intersect(FromHnf,ToMic));

HBac2Hnf=MCMC(intersect(FromHBac,ToHnf));
Cya2Hnf=MCMC(intersect(FromCya,ToHnf));

HBac2App=MCMC(intersect(FromHBac,ToApp));
Cya2App=MCMC(intersect(FromCya,ToApp));
Dtm2App=MCMC(intersect(FromDtm,ToApp));
Mix2App=MCMC(intersect(FromMix,ToApp));
Hnf2App=MCMC(intersect(FromHnf,ToApp));

Mic2Clad=MCMC(intersect(FromMic,ToClad));
Dtm2Clad=MCMC(intersect(FromDtm,ToClad));
Mix2Clad=MCMC(intersect(FromMix,ToClad));
Hnf2Clad=MCMC(intersect(FromHnf,ToClad));

Mic2Cal=sum(MCMC(intersect(FromMic,ToCal)));
Dtm2Cal=sum(MCMC(intersect(FromDtm,ToCal)));
Mix2Cal=sum(MCMC(intersect(FromMix,ToCal)));
Hnf2Cal=sum(MCMC(intersect(FromHnf,ToCal)));

Mic2Herb=sum(MCMC(intersect(FromMic,ToHerb)));
Dtm2Herb=sum(MCMC(intersect(FromDtm,ToHerb)));
Mix2Herb=sum(MCMC(intersect(FromMix,ToHerb)));
Hnf2Herb=sum(MCMC(intersect(FromHnf,ToHerb)));

App2Poecil=sum(MCMC(intersect(FromApp,ToPoecil)));
Clad2Poecil=sum(MCMC(intersect(FromClad,ToPoecil)));
Cal2Poecil=sum(MCMC(intersect(FromCal,ToPoecil)));
Herb2Poecil=sum(MCMC(intersect(FromHerb,ToPoecil)));

Mic2Bft=sum(MCMC(intersect(FromMic,ToBft)));
App2Bft=sum(MCMC(intersect(FromApp,ToBft)));
Clad2Bft=sum(MCMC(intersect(FromClad,ToBft)));
Cal2Bft=sum(MCMC(intersect(FromCal,ToBft)));
Herb2Bft=sum(MCMC(intersect(FromHerb,ToBft)));
Poecil2Bft=sum(MCMC(intersect(FromPoecil,ToBft)));


figure('Position',[0 0 length(bkgd(1,:,1)) length(bkgd(:,1,1))])
image(bkgd)
hold on
if strcmp(Taxon,'Cya')
    arrow(Cyano2HnfArrow(1,:),Cyano2HnfArrow(2,:),'EdgeColor',CyaColor,'FaceColor',CyaColor,'Width',2,'Length',20)
    arrow(Cyano2MicArrow(1,:),Cyano2MicArrow(2,:),'EdgeColor',CyaColor,'FaceColor',CyaColor,'Width',2,'Length',20)
    arrow(Cyano2MixArrow(1,:),Cyano2MixArrow(2,:),'EdgeColor',CyaColor,'FaceColor',CyaColor,'Width',2,'Length',20)
    arrow(Cyano2AppArrow(1,:),Cyano2AppArrow(2,:),'EdgeColor',CyaColor,'FaceColor',CyaColor,'Width',2,'Length',20)
elseif strcmp(Taxon,'Dtm')
    arrow(Dtm2MicArrow(1,:),Dtm2MicArrow(2,:),'EdgeColor',DtmColor,'FaceColor',DtmColor,'Width',2,'Length',20)
    arrow(Dtm2AppArrow(1,:),Dtm2AppArrow(2,:),'EdgeColor',DtmColor,'FaceColor',DtmColor,'Width',2,'Length',20)
    arrow(Dtm2CladArrow(1,:),Dtm2CladArrow(2,:),'EdgeColor',DtmColor,'FaceColor',DtmColor,'Width',2,'Length',20)
    arrow(Dtm2CalArrow(1,:),Dtm2CalArrow(2,:),'EdgeColor',DtmColor,'FaceColor',DtmColor,'Width',2,'Length',20)
    arrow(Dtm2HerbArrow(1,:),Dtm2HerbArrow(2,:),'EdgeColor',DtmColor,'FaceColor',DtmColor,'Width',2,'Length',20)
end
arrow(HBac2MixArrow(1,:),HBac2MixArrow(2,:),'EdgeColor',HBacColor,'FaceColor',HBacColor,'Width',2,'Length',20)
arrow(HBac2MicArrow(1,:),HBac2MicArrow(2,:),'EdgeColor',HBacColor,'FaceColor',HBacColor,'Width',2,'Length',20)
arrow(Mix2MicArrow(1,:),Mix2MicArrow(2,:),'EdgeColor',MixColor,'FaceColor',MixColor,'Width',2,'Length',20)
arrow(Hnf2MicArrow(1,:),Hnf2MicArrow(2,:),'EdgeColor',HnfColor,'FaceColor',HnfColor,'Width',2,'Length',20)
arrow(HBac2HnfArrow(1,:),HBac2HnfArrow(2,:),'EdgeColor',HBacColor,'FaceColor',HBacColor,'Width',2,'Length',20)
arrow(HBac2AppArrow(1,:),HBac2AppArrow(2,:),'EdgeColor',HBacColor,'FaceColor',HBacColor,'Width',2,'Length',20)
arrow(Hnf2AppArrow(1,:),Hnf2AppArrow(2,:),'EdgeColor',HnfColor,'FaceColor',HnfColor,'Width',2,'Length',20)
arrow(Mix2AppArrow(1,:),Mix2AppArrow(2,:),'EdgeColor',MixColor,'FaceColor',MixColor,'Width',2,'Length',20)
arrow(Hnf2CladArrow(1,:),Hnf2CladArrow(2,:),'EdgeColor',HnfColor,'FaceColor',HnfColor,'Width',2,'Length',20)
arrow(Mic2CladArrow(1,:),Mic2CladArrow(2,:),'EdgeColor',MicColor,'FaceColor',MicColor,'Width',2,'Length',20)
arrow(Mix2CladArrow(1,:),Mix2CladArrow(2,:),'EdgeColor',MixColor,'FaceColor',MixColor,'Width',2,'Length',20)
arrow(Hnf2CalArrow(1,:),Hnf2CalArrow(2,:),'EdgeColor',HnfColor,'FaceColor',HnfColor,'Width',2,'Length',20)
arrow(Mic2CalArrow(1,:),Mic2CalArrow(2,:),'EdgeColor',MicColor,'FaceColor',MicColor,'Width',2,'Length',20)
arrow(Mix2CalArrow(1,:),Mix2CalArrow(2,:),'EdgeColor',MixColor,'FaceColor',MixColor,'Width',2,'Length',20)
arrow(Hnf2HerbArrow(1,:),Hnf2HerbArrow(2,:),'EdgeColor',HnfColor,'FaceColor',HnfColor,'Width',2,'Length',20)
arrow(Mic2HerbArrow(1,:),Mic2HerbArrow(2,:),'EdgeColor',MicColor,'FaceColor',MicColor,'Width',2,'Length',20)
arrow(Mix2HerbArrow(1,:),Mix2HerbArrow(2,:),'EdgeColor',MixColor,'FaceColor',MixColor,'Width',2,'Length',20)
arrow(App2PoecilArrow(1,:),App2PoecilArrow(2,:),'EdgeColor',AppColor,'FaceColor',AppColor,'Width',2,'Length',20)
arrow(Clad2PoecilArrow(1,:),Clad2PoecilArrow(2,:),'EdgeColor',CladColor,'FaceColor',CladColor,'Width',2,'Length',20)
arrow(Cal2PoecilArrow(1,:),Cal2PoecilArrow(2,:),'EdgeColor',CalColor,'FaceColor',CalColor,'Width',2,'Length',20)
arrow(Herb2PoecilArrow(1,:),Herb2PoecilArrow(2,:),'EdgeColor',HerbColor,'FaceColor',HerbColor,'Width',2,'Length',20)
% arrow(App2BftArrow(1,:),App2BftArrow(2,:),'EdgeColor',AppColor,'FaceColor',AppColor,'Width',2,'Length',20)
% arrow(Clad2BftArrow(1,:),Clad2BftArrow(2,:),'EdgeColor',CladColor,'FaceColor',CladColor,'Width',2,'Length',20)
% arrow(Cal2BftArrow(1,:),Cal2BftArrow(2,:),'EdgeColor',CalColor,'FaceColor',CalColor,'Width',2,'Length',20)
% arrow(Herb2BftArrow(1,:),Herb2BftArrow(2,:),'EdgeColor',HerbColor,'FaceColor',HerbColor,'Width',2,'Length',20)
% arrow(Mic2BftArrow(1,:),Mic2BftArrow(2,:),'EdgeColor',MicColor,'FaceColor',MicColor,'Width',2,'Length',20)
% arrow(Poecil2BftArrow(1,:),Poecil2BftArrow(2,:),'EdgeColor',PoecilColor,'FaceColor',PoecilColor,'Width',2,'Length',20)

axis off
ax = gca;
outerpos = ax.OuterPosition;
ti = [0 0 0 0]; 
left = outerpos(1) + ti(1);
bottom = outerpos(2) + ti(2);
ax_width = outerpos(3) - ti(1) - ti(3);
ax_height = outerpos(4) - ti(2) - ti(4);
ax.Position = [left bottom ax_width ax_height];

frame = getframe(1);
fig = frame2im(frame);
    [imind,cm] = rgb2ind(fig,256);
    imwrite(imind,cm,['Indirect ',Taxon,'.png'],'png');

% 
% %Append Mixotroph Pie Chart
% figure('Position',[400 400 100 100])
% labels={'','',''}
% pie([HBac2Mix,Cya2Mix,Nut2Mix]./sum([HBac2Mix,Cya2Mix,Nut2Mix]),labels)
% set(gcf,'color','w');
% colormap([HBacColor;CyaColor;1,1,1])
% delete(ax.Children([1, 3])) % delete the text object according to index from ax.Children
% frame = getframe(2);
% temppie = frame2im(frame);
% %     [imind,cm] = rgb2ind(temppie,256);
% %     imwrite(imind,cm,'PieMask.jpg','jpg');
% close
% figure(1)
% [fig]=AppendDoodle3(fig,temppie,[760 540],2,piemask,0);
% 
% 
% figure('Position',[400 400 100 100])
% labels={'','','','',''}
% pie([HBac2Mic,Cya2Mic,Dtm2Mic,Mix2Mic,Hnf2Mic]./sum([HBac2Mic,Cya2Mic,Dtm2Mic,Mix2Mic,Hnf2Mic]),labels)
% set(gcf,'color','w');
% colormap([HBacColor;CyaColor;DtmColor;MixColor;HnfColor])
% delete(ax.Children([1, 3])) % delete the text object according to index from ax.Children
% frame = getframe(2);
% temppie = frame2im(frame);
% close
% [fig]=AppendDoodle3(fig,temppie,[460 500],2,piemask,0);
% [imind,cm] = rgb2ind(fig,256);
% imwrite(imind,cm,['Indirect ',Taxon,'.png'],'png');
% 
% 
% 
% figure('Position',[400 400 100 100])
% labels={'',''}
% pie([HBac2Hnf,Cya2Hnf]./sum([HBac2Hnf,Cya2Hnf]),labels)
% set(gcf,'color','w');
% colormap([HBacColor;CyaColor])
% delete(ax.Children([1, 3])) % delete the text object according to index from ax.Children
% frame = getframe(2);
% temppie = frame2im(frame);
% close
% [fig]=AppendDoodle3(fig,temppie,[10 510],2,piemask,0);
% [imind,cm] = rgb2ind(fig,256);
% imwrite(imind,cm,['Indirect ',Taxon,'.png'],'png');


figure('Position',[400 400 100 100])
labels={'',''}
pie([PerApp 1-PerApp],labels)
set(gcf,'color','w');
colormap([AppColor;1,1,1])
delete(ax.Children([1, 3])) % delete the text object according to index from ax.Children
frame = getframe(2);
temppie = frame2im(frame);
close
[fig]=AppendDoodle3(fig,temppie,[60 320],2,piemask,0);
[imind,cm] = rgb2ind(fig,256);
imwrite(imind,cm,['Indirect ',Taxon,'.png'],'png');


figure('Position',[400 400 100 100])
labels={'',''}
pie([PerClad 1-PerClad],labels)
set(gcf,'color','w');
colormap([CladColor;1,1,1])
delete(ax.Children([1, 3])) % delete the text object according to index from ax.Children
frame = getframe(2);
temppie = frame2im(frame);
close
[fig]=AppendDoodle3(fig,temppie,[330 320],2,piemask,0);
[imind,cm] = rgb2ind(fig,256);
imwrite(imind,cm,['Indirect ',Taxon,'.png'],'png');


figure('Position',[400 400 100 100])
labels={'',''}
pie([PerCal 1-PerCal],labels)
set(gcf,'color','w');
colormap([CalColor;1,1,1])
delete(ax.Children([1, 3])) % delete the text object according to index from ax.Children
frame = getframe(2);
temppie = frame2im(frame);
close
[fig]=AppendDoodle3(fig,temppie,[580 320],2,piemask,0);
[imind,cm] = rgb2ind(fig,256);
imwrite(imind,cm,['Indirect ',Taxon,'.png'],'png');


figure('Position',[400 400 100 100])
labels={'',''}
pie([PerHerb 1-PerHerb],labels)
set(gcf,'color','w');
colormap([HerbColor;1,1,1])
delete(ax.Children([1, 3])) % delete the text object according to index from ax.Children
frame = getframe(2);
temppie = frame2im(frame);
close
[fig]=AppendDoodle3(fig,temppie,[800 360],2,piemask,0);
[imind,cm] = rgb2ind(fig,256);
imwrite(imind,cm,['Indirect ',Taxon,'.png'],'png');


figure('Position',[400 400 100 100])
labels={'',''}
pie([PerPoecil 1-PerPoecil],labels)
set(gcf,'color','w');
colormap([PoecilColor;1,1,1])
delete(ax.Children([1, 3])) % delete the text object according to index from ax.Children
frame = getframe(2);
temppie = frame2im(frame);
close
[fig]=AppendDoodle3(fig,temppie,[760 180],2,piemask,0);
[imind,cm] = rgb2ind(fig,256);
imwrite(imind,cm,['Indirect ',Taxon,'.png'],'png');


% 
% figure('Position',[400 400 100 100])
% labels={'','','','','',''}
% pie([Mic2Bft,App2Bft,Clad2Bft,Cal2Bft,Herb2Bft,Poecil2Bft]./sum([Mic2Bft,App2Bft,Clad2Bft,Cal2Bft,Herb2Bft,Poecil2Bft]),labels)
% set(gcf,'color','w');
% colormap([MicColor;AppColor;CladColor;CalColor;HerbColor;PoecilColor])
% delete(ax.Children([1, 3])) % delete the text object according to index from ax.Children
% frame = getframe(2);
% temppie = frame2im(frame);
% close
% [fig]=AppendDoodle3(fig,temppie,[480 160],2,piemask,0);
% [imind,cm] = rgb2ind(fig,256);
% imwrite(imind,cm,['Indirect ',Taxon,'.png'],'png');
% 
% 


figure('Position',[200 200 200 200])
labels={'','','','','',''}
%labels={'Appendicularians','Cladocerans','Calanoids','Herbivores','Poecilostomatoids',''}
pie([PerApp,PerClad,PerCal,PerHerb,PerPoecil,1-sum([PerApp,PerClad,PerCal,PerHerb,PerPoecil])],labels)
set(gcf,'color','w');
colormap([AppColor;CladColor;CalColor;HerbColor;PoecilColor;1,1,1])
delete(ax.Children([1, 3])) % delete the text object according to index from ax.Children
frame = getframe(2);
temppie = frame2im(frame);
close
[fig]=AppendDoodle3(fig,temppie,[410 60],2,piemask_Lg,0);
[imind,cm] = rgb2ind(fig,256);
imwrite(imind,cm,['Indirect ',Taxon,'.png'],'png');


output=0
close all
```
