## Supplementary material for "Comparative food-web analysis of bluefin tuna spawning habitats in the eastern Indian Ocean and Gulf of Mexico": Code Package: FoodwebPathways2Tuna.m

```
clear all
close all

burnin = 0.2;

for Cycle = [0,10]

    load(['N15GoMInverseCycle',num2str(Cycle),'.mat'])
    load(['N15GoMInverseCycle',num2str(Cycle),'Routputs.mat'])
    [num,txt,raw]  = xlsread('N15InverseModelRW.GOM.xlsx','Sheet1','B6:B307');
    Columns = txt;
    clearvars -except Ae0 MCMCmat Aa d15NInputs del15N InputCol wts Cycle burnin Columns
    num=length(MCMCmat(:,1));
    MCMCmat(1:round(num*burnin),:)=[];
    del15N(1:round(num*burnin),:)=[];
    len=1000 %length(MCMCmat(:,1));
    alphabet = 'abcdefghijklmnop';

    for i=1:len
        i
        flows=MCMCmat(i,:)./wts';

        [P,struct,ext,r] = IndirectFlowsRespiration(Ae0,flows);

        [NO3s_col,NH4s_col,Cyas_col,Tris_col,Dtms_col,Flags_col,HNFs_col,MICs_col,HerbNVMs_col,Apps_col,Clads_col,nvmCals_col,Chaetos_col,Poecils_col,GelPreds_col,Plankfishs_col,Preflexs_col,Postflexs_col,Bacs_col,Dets_col,Ldets_col,Dons_col,HerbVM_col,vmCal_col,NO3d_col,NH4d_col,Cyad_col,Trid_col,Dtmd_col,Flagd_col,HNFd_col,MICd_col,HerbNVMd_col,Appd_col,Cladd_col,nvmCald_col,Chaetod_col,Poecild_col,GelPredd_col,Plankfishd_col,Bacd_col,Detd_col,Ldetd_col,Dond_col] = GetColumns(Aa,MCMCmat,del15N(1,:),d15NInputs(1,InputCol),d15NInputs(2,InputCol),d15NInputs(3,InputCol),d15NInputs(4,InputCol),d15NInputs(5,InputCol),d15NInputs(6,InputCol),d15NInputs(7,InputCol));

        Cyanos2Respiration=struct(Cyas_col,:)'.*r;
        CyanosProduction=sum(P(Cyas_col,:));
        Cyanos2RespirationNormalized=Cyanos2Respiration/CyanosProduction;
        Cyanod2Respiration=struct(Cyad_col,:)'.*r;
        CyanodProduction=sum(P(Cyad_col,:));
        Cyanod2RespirationNormalized=Cyanod2Respiration/CyanodProduction;
        Cyano2Preflex(i)=Cyanos2Respiration(Preflexs_col)+Cyanod2Respiration(Preflexs_col);
        Cyano2Postflex(i)=Cyanos2Respiration(Postflexs_col)+Cyanod2Respiration(Postflexs_col);
        Cyanos2Preflex(i)=Cyanos2Respiration(Preflexs_col);
        Cyanos2Postflex(i)=Cyanos2Respiration(Postflexs_col);
        Cyanod2Preflex(i)=Cyanod2Respiration(Preflexs_col);
        Cyanod2Postflex(i)=Cyanod2Respiration(Postflexs_col);
        Cyano2PreflexNormalized(i)=Cyanos2RespirationNormalized(Preflexs_col)+Cyanod2RespirationNormalized(Preflexs_col);
        Cyano2PostflexNormalized(i)=Cyanos2RespirationNormalized(Postflexs_col)+Cyanod2RespirationNormalized(Postflexs_col);
        Cyano2HNFs(i)=Cyanos2Respiration(HNFs_col)+Cyanod2Respiration(HNFs_col);
        Cyano2HNFsNormalized(i)=Cyanos2RespirationNormalized(HNFs_col)+Cyanod2RespirationNormalized(HNFs_col);
        Cyano2Mics(i)=Cyanos2Respiration(MICs_col)+Cyanod2Respiration(MICs_col);
        Cyano2MicsNormalized(i)=Cyanos2RespirationNormalized(MICs_col)+Cyanod2RespirationNormalized(MICs_col);
        Cyano2Apps(i)=Cyanos2Respiration(Apps_col)+Cyanod2Respiration(Apps_col);
        Cyano2AppsNormalized(i)=Cyanos2RespirationNormalized(Apps_col)+Cyanod2RespirationNormalized(Apps_col);
        Cyano2Clads(i)=Cyanos2Respiration(Clads_col)+Cyanod2Respiration(Clads_col);
        Cyano2CladsNormalized(i)=Cyanos2RespirationNormalized(Clads_col)+Cyanod2RespirationNormalized(Clads_col);
        Cyano2nvmCals(i)=Cyanos2Respiration(nvmCals_col)+Cyanod2Respiration(nvmCals_col);
        Cyano2nvmCalsNormalized(i)=Cyanos2RespirationNormalized(nvmCals_col)+Cyanod2RespirationNormalized(nvmCals_col);
        Cyano2Poecils(i)=Cyanos2Respiration(Poecils_col)+Cyanod2Respiration(Poecils_col);
        Cyano2PoecilsNormalized(i)=Cyanos2RespirationNormalized(Poecils_col)+Cyanod2RespirationNormalized(Poecils_col);

        Diatoms2Respiration=struct(Dtms_col,:)'.*r;
        DiatomsProduction=sum(P(Dtms_col,:));
        Diatoms2RespirationNormalized=Diatoms2Respiration/DiatomsProduction;
        Diatomd2Respiration=struct(Dtmd_col,:)'.*r;
        DiatomdProduction=sum(P(Dtmd_col,:));
        Diatomd2RespirationNormalized=Diatomd2Respiration/DiatomdProduction;
        Diatom2Preflex(i)=Diatoms2Respiration(Preflexs_col)+Diatomd2Respiration(Preflexs_col);
        Diatom2Postflex(i)=Diatoms2Respiration(Postflexs_col)+Diatomd2Respiration(Postflexs_col);
        Diatom2PreflexNormalized(i)=Diatoms2RespirationNormalized(Preflexs_col)+Diatomd2RespirationNormalized(Preflexs_col);
        Diatom2PostflexNormalized(i)=Diatoms2RespirationNormalized(Postflexs_col)+Diatomd2RespirationNormalized(Postflexs_col);
        Diatom2HNFs(i)=Diatoms2Respiration(HNFs_col)+Diatomd2Respiration(HNFs_col);
        Diatom2HNFsNormalized(i)=Diatoms2RespirationNormalized(HNFs_col)+Diatomd2RespirationNormalized(HNFs_col);
        Diatom2Mics(i)=Diatoms2Respiration(MICs_col)+Diatomd2Respiration(MICs_col);
        Diatom2MicsNormalized(i)=Diatoms2RespirationNormalized(MICs_col)+Diatomd2RespirationNormalized(MICs_col);
        Diatom2Apps(i)=Diatoms2Respiration(Apps_col)+Diatomd2Respiration(Apps_col);
        Diatom2AppsNormalized(i)=Diatoms2RespirationNormalized(Apps_col)+Diatomd2RespirationNormalized(Apps_col);
        Diatom2Clads(i)=Diatoms2Respiration(Clads_col)+Diatomd2Respiration(Clads_col);
        Diatom2CladsNormalized(i)=Diatoms2RespirationNormalized(Clads_col)+Diatomd2RespirationNormalized(Clads_col);
        Diatom2nvmCals(i)=Diatoms2Respiration(nvmCals_col)+Diatomd2Respiration(nvmCals_col);
        Diatom2nvmCalsNormalized(i)=Diatoms2RespirationNormalized(nvmCals_col)+Diatomd2RespirationNormalized(nvmCals_col);
        Diatom2Poecils(i)=Diatoms2Respiration(Poecils_col)+Diatomd2Respiration(Poecils_col);
        Diatom2PoecilsNormalized(i)=Diatoms2RespirationNormalized(Poecils_col)+Diatomd2RespirationNormalized(Poecils_col);

        Trichos2Respiration=struct(Tris_col,:)'.*r;
        TrichosProduction=sum(P(Tris_col,:));
        Trichos2RespirationNormalized=Trichos2Respiration/TrichosProduction;
        Trichod2Respiration=struct(Trid_col,:)'.*r;
        TrichodProduction=sum(P(Trid_col,:));
        Trichod2RespirationNormalized=Trichod2Respiration/TrichodProduction;
        Tricho2Preflex(i)=Trichos2Respiration(Preflexs_col)+Trichod2Respiration(Preflexs_col);
        Tricho2Postflex(i)=Trichos2Respiration(Postflexs_col)+Trichod2Respiration(Postflexs_col);
        Trichos2Preflex(i)=Trichos2Respiration(Preflexs_col);
        Trichos2Postflex(i)=Trichos2Respiration(Postflexs_col);
        Trichod2Preflex(i)=Trichod2Respiration(Preflexs_col);
        Trichod2Postflex(i)=Trichod2Respiration(Postflexs_col);
        Tricho2PreflexNormalized(i)=Trichos2RespirationNormalized(Preflexs_col)+Trichod2RespirationNormalized(Preflexs_col);
        Tricho2PostflexNormalized(i)=Trichos2RespirationNormalized(Postflexs_col)+Trichod2RespirationNormalized(Postflexs_col);
        Tricho2HNFs(i)=Trichos2Respiration(HNFs_col)+Trichod2Respiration(HNFs_col);
        Tricho2HNFsNormalized(i)=Trichos2RespirationNormalized(HNFs_col)+Trichod2RespirationNormalized(HNFs_col);
        Tricho2Mics(i)=Trichos2Respiration(MICs_col)+Trichod2Respiration(MICs_col);
        Tricho2MicsNormalized(i)=Trichos2RespirationNormalized(MICs_col)+Trichod2RespirationNormalized(MICs_col);
        Tricho2Apps(i)=Trichos2Respiration(Apps_col)+Trichod2Respiration(Apps_col);
        Tricho2AppsNormalized(i)=Trichos2RespirationNormalized(Apps_col)+Trichod2RespirationNormalized(Apps_col);
        Tricho2Clads(i)=Trichos2Respiration(Clads_col)+Trichod2Respiration(Clads_col);
        Tricho2CladsNormalized(i)=Trichos2RespirationNormalized(Clads_col)+Trichod2RespirationNormalized(Clads_col);
        Tricho2nvmCals(i)=Trichos2Respiration(nvmCals_col)+Trichod2Respiration(nvmCals_col);
        Tricho2nvmCalsNormalized(i)=Trichos2RespirationNormalized(nvmCals_col)+Trichod2RespirationNormalized(nvmCals_col);
        Tricho2Poecils(i)=Trichos2Respiration(Poecils_col)+Trichod2Respiration(Poecils_col);
        Tricho2PoecilsNormalized(i)=Trichos2RespirationNormalized(Poecils_col)+Trichod2RespirationNormalized(Poecils_col);

        Flags2Respiration=struct(Flags_col,:)'.*r;
        FlagsProduction=sum(P(Flags_col,:));
        Flags2RespirationNormalized=Flags2Respiration/FlagsProduction;
        Flagd2Respiration=struct(Flagd_col,:)'.*r;
        FlagdProduction=sum(P(Flagd_col,:));
        Flagd2RespirationNormalized=Flagd2Respiration/FlagdProduction;
        Flag2Preflex(i)=Flags2Respiration(Preflexs_col)+Flagd2Respiration(Preflexs_col);
        Flag2Postflex(i)=Flags2Respiration(Postflexs_col)+Flagd2Respiration(Postflexs_col);
        Flag2PreflexNormalized(i)=Flags2RespirationNormalized(Preflexs_col)+Flagd2RespirationNormalized(Preflexs_col);
        Flag2PostflexNormalized(i)=Flags2RespirationNormalized(Postflexs_col)+Flagd2RespirationNormalized(Postflexs_col);
        Flag2HNFs(i)=Flags2Respiration(HNFs_col)+Flagd2Respiration(HNFs_col);
        Flag2HNFsNormalized(i)=Flags2RespirationNormalized(HNFs_col)+Flagd2RespirationNormalized(HNFs_col);
        Flag2Mics(i)=Flags2Respiration(MICs_col)+Flagd2Respiration(MICs_col);
        Flag2MicsNormalized(i)=Flags2RespirationNormalized(MICs_col)+Flagd2RespirationNormalized(MICs_col);
        Flag2Apps(i)=Flags2Respiration(Apps_col)+Flagd2Respiration(Apps_col);
        Flag2AppsNormalized(i)=Flags2RespirationNormalized(Apps_col)+Flagd2RespirationNormalized(Apps_col);
        Flag2Clads(i)=Flags2Respiration(Clads_col)+Flagd2Respiration(Clads_col);
        Flag2CladsNormalized(i)=Flags2RespirationNormalized(Clads_col)+Flagd2RespirationNormalized(Clads_col);
        Flag2nvmCals(i)=Flags2Respiration(nvmCals_col)+Flagd2Respiration(nvmCals_col);
        Flag2nvmCalsNormalized(i)=Flags2RespirationNormalized(nvmCals_col)+Flagd2RespirationNormalized(nvmCals_col);
        Flag2Poecils(i)=Flags2Respiration(Poecils_col)+Flagd2Respiration(Poecils_col);
        Flag2PoecilsNormalized(i)=Flags2RespirationNormalized(Poecils_col)+Flagd2RespirationNormalized(Poecils_col);

        Bacs2Respiration=struct(Bacs_col,:)'.*r;
        BacsProduction=sum(P(Bacs_col,:));
        Bacd2Respiration=struct(Bacd_col,:)'.*r;
        BacdProduction=sum(P(Bacd_col,:));
        Bac2Preflex(i)=Bacs2Respiration(Preflexs_col)+Bacd2Respiration(Preflexs_col);
        Bac2Postflex(i)=Bacs2Respiration(Postflexs_col)+Bacd2Respiration(Postflexs_col);
        Bac2Apps(i)=Bacs2Respiration(Apps_col)+Bacd2Respiration(Apps_col);

            %------------Computing some metrics related to nitrogen fixation--------
    NFixCyanos_col = find(strcmp(Columns,'NfixCyanos'));
    NFixCyanos(i) = flows(NFixCyanos_col);
    NFixCyanod_col = find(strcmp(Columns,'NfixCyanod'));
    NFixCyanod(i) = flows(NFixCyanod_col);
    NO3toCyanos_col = find(strcmp(Columns,'NO3toCyanos'));
    NO3toCyanos(i) = flows(NO3toCyanos_col);
    NH4toCyanos_col = find(strcmp(Columns,'NH4toCyanos'));
    NH4toCyanos(i) = flows(NH4toCyanos_col);
    NO3toCyanod_col = find(strcmp(Columns,'NO3toCyanod'));
    NO3toCyanod(i) = flows(NO3toCyanod_col);
    NH4toCyanod_col = find(strcmp(Columns,'NH4toCyanod'));
    NH4toCyanod(i) = flows(NH4toCyanod_col);
    NFixCyanoPercents = NFixCyanos ./ (NFixCyanos + NH4toCyanos + NO3toCyanos);
    NFixCyanoPercentd = NFixCyanod ./ (NFixCyanod + NH4toCyanod + NO3toCyanod);

    NFixTrichos_col = find(strcmp(Columns,'NfixTrichos'));
    NFixTrichos(i) = flows(NFixTrichos_col);
    NFixTrichod_col = find(strcmp(Columns,'NfixTrichod'));
    NFixTrichod(i) = flows(NFixTrichod_col);
    NO3toTrichos_col = find(strcmp(Columns,'NO3toTrichos'));
    NO3toTrichos(i) = flows(NO3toTrichos_col);
    NH4toTrichos_col = find(strcmp(Columns,'NH4toTrichos'));
    NH4toTrichos(i) = flows(NH4toTrichos_col);
    NO3toTrichod_col = find(strcmp(Columns,'NO3toTrichod'));
    NO3toTrichod(i) = flows(NO3toTrichod_col);
    NH4toTrichod_col = find(strcmp(Columns,'NH4toTrichod'));
    NH4toTrichod(i) = flows(NH4toTrichod_col);
    NFixTrichoPercents = NFixTrichos ./ (NFixTrichos + NH4toTrichos + NO3toTrichos);
    NFixTrichoPercentd = NFixTrichod ./ (NFixTrichod + NH4toTrichod + NO3toTrichod);

    %-------------------Computing some direct tuna diet metrics---------
    ToNH4s = find(Ae0(NH4s_col,:)==1);
    ToPreflex = find(Ae0(Preflexs_col,:)==1);
    ToPostflex = find(Ae0(Postflexs_col,:)==1);
    ToChaeto = find(Ae0(Chaetos_col,:)==1);
    ToPoecil = find(Ae0(Poecils_col,:)==1);
    ToGelPred = find(Ae0(GelPreds_col,:)==1);
    ToPlankfish = find(Ae0(Plankfishs_col,:)==1);
    FromMIC = find(Ae0(MICs_col,:)==-1);
    FromApp = find(Ae0(Apps_col,:)==-1);
    FromClad = find(Ae0(Clads_col,:)==-1);
    FromnvmCal = find(Ae0(nvmCals_col,:)==-1);
    FromPoecil = find(Ae0(Poecils_col,:)==-1);

    MICtoPREFLEX = intersect(ToPreflex,FromMIC);
    MICtoPOSTFLEX = intersect(ToPostflex,FromMIC);
    ApptoCHAETO = intersect(ToChaeto,FromApp);
    ApptoPOECIL = intersect(ToPoecil,FromApp);
    ApptoGEL = intersect(ToGelPred,FromApp);
    ApptoPLANKFISH = intersect(ToPlankfish,FromApp);
    ApptoPREFLEX = intersect(ToPreflex,FromApp);
    ApptoPOSTFLEX = intersect(ToPostflex,FromApp);
    ApptoNH4 = intersect(ToNH4s,FromApp);
    CladtoCHAETO = intersect(ToChaeto,FromClad);
    CladtoPOECIL = intersect(ToPoecil,FromClad);
    CladtoGEL = intersect(ToGelPred,FromClad);
    CladtoPLANKFISH = intersect(ToPlankfish,FromClad);
    CladtoPREFLEX = intersect(ToPreflex,FromClad);
    CladtoPOSTFLEX = intersect(ToPostflex,FromClad);
    CladtoNH4 = intersect(ToNH4s,FromClad);
    nvmCaltoCHAETO = intersect(ToChaeto,FromnvmCal);
    nvmCaltoPOECIL = intersect(ToPoecil,FromnvmCal);
    nvmCaltoGEL = intersect(ToGelPred,FromnvmCal);
    nvmCaltoPLANKFISH = intersect(ToPlankfish,FromnvmCal);
    nvmCaltoPREFLEX = intersect(ToPreflex,FromnvmCal);
    nvmCaltoPOSTFLEX = intersect(ToPostflex,FromnvmCal);
    nvmCaltoNH4 = intersect(ToNH4s,FromnvmCal);
    PoeciltoPREFLEX = intersect(ToPreflex,FromPoecil);
    PoeciltoPOSTFLEX = intersect(ToPostflex,FromPoecil);

    MIC2PREFLEX(i) = flows(MICtoPREFLEX);
    MIC2POSTFLEX(i) = flows(MICtoPOSTFLEX);
    App2PREFLEX(i) = flows(ApptoPREFLEX);
    App2POSTFLEX(i) = flows(ApptoPOSTFLEX);
    Clad2PREFLEX(i) = flows(CladtoPREFLEX);
    Clad2POSTFLEX(i) = flows(CladtoPOSTFLEX);
    nvmCal2PREFLEX(i) = flows(nvmCaltoPREFLEX);
    nvmCal2POSTFLEX(i) = flows(nvmCaltoPOSTFLEX);
    Poecil2PREFLEX(i) = flows(PoeciltoPREFLEX);
    Poecil2POSTFLEX(i) = flows(PoeciltoPOSTFLEX);
    Clad2CHAETO(i) = flows(CladtoCHAETO);
    Clad2POECIL(i) = flows(CladtoPOECIL);
    Clad2GEL(i) = flows(CladtoGEL);
    Clad2PLANKFISH(i) = flows(CladtoPLANKFISH);
    Clad2NH4(i) = flows(CladtoNH4);
    App2CHAETO(i) = flows(ApptoCHAETO);
    App2POECIL(i) = flows(ApptoPOECIL);
    App2GEL(i) = flows(ApptoGEL);
    App2PLANKFISH(i) = flows(ApptoPLANKFISH);
    App2NH4(i) = flows(ApptoNH4);
    nvmCal2CHAETO(i) = flows(nvmCaltoCHAETO);
    nvmCal2POECIL(i) = flows(nvmCaltoPOECIL);
    nvmCal2GEL(i) = flows(nvmCaltoGEL);
    nvmCal2PLANKFISH(i) = flows(nvmCaltoPLANKFISH);
    nvmCal2NH4(i) = flows(nvmCaltoNH4);

    Clad2TUNApercent(i) = (Clad2PREFLEX(i)+Clad2POSTFLEX(i))/( Clad2PREFLEX(i)+Clad2POSTFLEX(i)+Clad2CHAETO(i)+Clad2POECIL(i)+Clad2GEL(i)+Clad2PLANKFISH(i) );    %This is the percentage of cladoceran secondary production consumed by tuna
    App2TUNApercent(i) = (App2PREFLEX(i)+App2POSTFLEX(i))/( App2PREFLEX(i)+App2POSTFLEX(i)+App2CHAETO(i)+App2POECIL(i)+App2GEL(i)+App2PLANKFISH(i) );
    nvmCal2TUNApercent(i) = (nvmCal2PREFLEX(i)+nvmCal2POSTFLEX(i))/( nvmCal2PREFLEX(i)+nvmCal2POSTFLEX(i)+nvmCal2CHAETO(i)+nvmCal2POECIL(i)+nvmCal2GEL(i)+nvmCal2PLANKFISH(i) );

    CladRespProdRatio(i) = (Clad2NH4(i))/( Clad2PREFLEX(i)+Clad2POSTFLEX(i)+Clad2CHAETO(i)+Clad2POECIL(i)+Clad2GEL(i)+Clad2PLANKFISH(i) );    %This is the ratio of cladoceran respiration (NH4 excretion) to cladoceran secondary production
    AppRespProdRatio(i) = (App2NH4(i))/( App2PREFLEX(i)+App2POSTFLEX(i)+App2CHAETO(i)+App2POECIL(i)+App2GEL(i)+App2PLANKFISH(i) );
    nvmCalRespProdRatio(i) = (nvmCal2NH4(i))/( nvmCal2PREFLEX(i)+nvmCal2POSTFLEX(i)+nvmCal2CHAETO(i)+nvmCal2POECIL(i)+nvmCal2GEL(i)+nvmCal2PLANKFISH(i) );

    TUNAconsumption(i) = (Clad2PREFLEX(i)+Clad2POSTFLEX(i) + App2PREFLEX(i)+App2POSTFLEX(i) + nvmCal2PREFLEX(i)+nvmCal2POSTFLEX(i) + Poecil2PREFLEX(i)+Poecil2POSTFLEX(i) + MIC2PREFLEX(i)+MIC2POSTFLEX(i));

    end


    for j=1:2

        fighandle = figure(101+j);
        fighandle.Units = 'inches';
        fighandle.Position = [1 1+Cycle/3 9.5 3];
        if Cycle==0
            col = [0 0 1];
        elseif Cycle==10
            col = [1 0 0];
        end

        subplot(1,8,1)  %----------- N2 fixation ==> Tuna ----------------
        hold on
        if j==1
            y = Cyanos2Preflex.*NFixCyanoPercents  + Cyanod2Preflex.*NFixCyanoPercents  + Trichos2Preflex.*NFixTrichoPercents  + Trichod2Preflex.*NFixTrichoPercents + ...
                Cyanos2Postflex.*NFixCyanoPercents + Cyanod2Postflex.*NFixCyanoPercents + Trichos2Postflex.*NFixTrichoPercents + Trichod2Postflex.*NFixTrichoPercents;
            bar(Cycle/10,mean(y)*10^6,'FaceColor',col);

            set(gca,'XTick',0.5)
            set(gca,'XTickLabel',{'NFix==> ? ==>Tuna'})
            % set(gca,'box','on')
            ylabel(['Nitrogen Flux',char(10),'(nmol N m^-^3 d^-^1)'])
            set(gca,'XTickLabelRotation',60)
            set(gca,'FontSize',9)
            if Cycle==10
                AddPanelLabels('a','topleft','YPADDING',0.01,'XPADDING',0.05,'FONTSIZE',9)
            end

        elseif j==2
            tot = Cyano2Preflex+Cyano2Postflex + Flag2Preflex+Flag2Postflex + Diatom2Preflex+Diatom2Postflex + Tricho2Preflex+Tricho2Postflex;
            y = Cyanos2Preflex.*NFixCyanoPercents  + Cyanod2Preflex.*NFixCyanoPercents  + Trichos2Preflex.*NFixTrichoPercents  + Trichod2Preflex.*NFixTrichoPercents + ...
                Cyanos2Postflex.*NFixCyanoPercents + Cyanod2Postflex.*NFixCyanoPercents + Trichos2Postflex.*NFixTrichoPercents + Trichod2Postflex.*NFixTrichoPercents;
            z = y./tot;
            bar(Cycle/10,mean(z)*100,'FaceColor',col);

            set(gca,'XTick',0.5)
            set(gca,'XTickLabel',{'NFix==> ? ==>Tuna'})
            % set(gca,'box','on')
            %ylabel(['Nitrogen Flux',char(10),'(% of flux to larvae)'])
            set(gca,'XTickLabelRotation',60)
            set(gca,'FontSize',9)
            if Cycle==10
                AddPanelLabels('a','topleft','YPADDING',0.01,'XPADDING',0.05,'FONTSIZE',9,'YPADDING',0.01,'XPADDING',0.05,'FONTSIZE',9)
            end


        end


        subplot(1,8,2)  %----------- Cyano ==> Tuna ----------------
        hold on
        if j==1
            bar(Cycle/10,mean(Cyano2Preflex+Cyano2Postflex)*10^6,'FaceColor',col);

            set(gca,'XTick',0.5)
            set(gca,'XTickLabel',{'Cya==> ? ==>Tuna'})
            % set(gca,'box','on')
            ylabel(['Nitrogen Flux',char(10),'(nmol N m^-^3 d^-^1)'])
            set(gca,'XTickLabelRotation',60)
            set(gca,'FontSize',9)
            if Cycle==10
                AddPanelLabels('a','topleft','YPADDING',0.01,'XPADDING',0.05,'FONTSIZE',9)
            end

        elseif j==2
            tot = Cyano2Preflex+Cyano2Postflex + Flag2Preflex+Flag2Postflex + Diatom2Preflex+Diatom2Postflex + Tricho2Preflex+Tricho2Postflex;
            y = (Cyano2Preflex+Cyano2Postflex);
            z = y./tot;
            bar(Cycle/10,mean(z)*100,'FaceColor',col);

            set(gca,'XTick',0.5)
            set(gca,'XTickLabel',{'Cya==> ? ==>Tuna'})
            % set(gca,'box','on')
            %ylabel(['Nitrogen Flux',char(10),'(% of flux to larvae)'])
            set(gca,'XTickLabelRotation',60)
            set(gca,'FontSize',9)
            ylim([0 60])
            if Cycle==10
                AddPanelLabels('b','topleft','YPADDING',0.01,'XPADDING',0.05,'FONTSIZE',9)
            end


            fn = 'ManuscriptValues.txt'
            fileID = fopen(fn,'a');
            if Cycle==0
                formatSpec =  '\n FoodwebPathways2Tuna.m \n ';
                fprintf(fileID,formatSpec)
            end
            formatSpec =  'For Cycle %f cyanobacteria supported  %.0f +/- %.0f percent of larvae diets \n';
            fprintf(fileID,formatSpec,[Cycle, mean(z)*100, std(z)*100])
        end

        subplot(1,8,3)  %----------- Flagellates ==> Tuna ----------------
        hold on
        if j==1
            bar(Cycle/10,mean(Flag2Preflex+Flag2Postflex)*10^6,'FaceColor',col);

            set(gca,'XTick',0.5)
            set(gca,'XTickLabel',{'Flag==> ? ==>Tuna'})
            % set(gca,'box','on')
            ylabel(['Nitrogen Flux',char(10),'(nmol N m^-^3 d^-^1)'])
            set(gca,'XTickLabelRotation',60)
            set(gca,'FontSize',9)
            if Cycle==10
                AddPanelLabels('b','topleft')
            end

        elseif j==2
            tot = Cyano2Preflex+Cyano2Postflex + Flag2Preflex+Flag2Postflex + Diatom2Preflex+Diatom2Postflex + Tricho2Preflex+Tricho2Postflex;
            y = (Flag2Preflex+Flag2Postflex);
            z=y./tot;
            bar(Cycle/10,mean(z)*100,'FaceColor',col);

            set(gca,'XTick',0.5)
            set(gca,'XTickLabel',{'Flag==> ? ==>Tuna'})
            % set(gca,'box','on')
            %ylabel(['Nitrogen Flux',char(10),'(% of flux to larvae)'])
            set(gca,'XTickLabelRotation',60)
            set(gca,'FontSize',9)
            ylim([0 60])
            if Cycle==10
                AddPanelLabels('c','topleft','YPADDING',0.01,'XPADDING',0.05,'FONTSIZE',9)
            end
        end


        subplot(1,8,4)  %----------- Cya ==> App ==> Tuna ----------------
        hold on
        if j==1
            y = App2TUNApercent.*Cyano2Apps./AppRespProdRatio;
            bar(Cycle/10,mean(y)*10^6,'FaceColor',col);

            set(gca,'XTick',0.5);
            set(gca,'XTickLabel',{'Cya==> App ==>Tuna'});
            % set(gca,'box','on')
            ylabel(['Nitrogen Flux',char(10),'(nmol N m^-^3 d^-^1)']);
            set(gca,'XTickLabelRotation',60);
            set(gca,'FontSize',9);
            if Cycle==10
                AddPanelLabels('a','topleft','YPADDING',0.01,'XPADDING',0.05,'FONTSIZE',9)
            end

        elseif j==2
            tot = TUNAconsumption;
            y = App2TUNApercent.*Cyano2Apps./AppRespProdRatio;
            z = y./tot;
            bar(Cycle/10,mean(z)*100,'FaceColor',col);

            set(gca,'XTick',0.5);
            set(gca,'XTickLabel',{'Cya==> App ==>Tuna'});
            % set(gca,'box','on')
            %ylabel(['Nitrogen Flux',char(10),'(% of flux to larvae)']);
            set(gca,'XTickLabelRotation',60);
            set(gca,'FontSize',9);
            ylim([0 50])
            if Cycle==10
                AddPanelLabels('d','topleft','YPADDING',0.01,'XPADDING',0.05,'FONTSIZE',9);
            end

            formatSpec =  'For Cycle %f cyanobacteria ==> app ==> tuna supported  %.0f +/- %.0f percent of larvae diets \n';
            fprintf(fileID,formatSpec,[Cycle, mean(z)*100, std(z)*100])
        end


        subplot(1,8,5)  %----------- Flag ==> App ==> Tuna ----------------
        hold on
        if j==1
            y = App2TUNApercent.*Flag2Apps./AppRespProdRatio;
            bar(Cycle/10,mean(y)*10^6,'FaceColor',col);

            set(gca,'XTick',0.5);
            set(gca,'XTickLabel',{'Flag==> App ==>Tuna'});
            % set(gca,'box','on')
            ylabel(['Nitrogen Flux',char(10),'(nmol N m^-^3 d^-^1)']);
            set(gca,'XTickLabelRotation',60);
            set(gca,'FontSize',9);
            if Cycle==10
                AddPanelLabels('a','topleft','YPADDING',0.01,'XPADDING',0.05,'FONTSIZE',9)
            end

        elseif j==2
            tot = TUNAconsumption;
            y = App2TUNApercent.*Flag2Apps./AppRespProdRatio;
            z = y./tot;
            bar(Cycle/10,mean(z)*100,'FaceColor',col);

            set(gca,'XTick',0.5);
            set(gca,'XTickLabel',{'Flag==> App ==>Tuna'});
            % set(gca,'box','on')
            %ylabel(['Nitrogen Flux',char(10),'(% of flux to larvae)']);
            set(gca,'XTickLabelRotation',60);
            set(gca,'FontSize',9);
            ylim([0 50])
            if Cycle==10
                AddPanelLabels('e','topleft','YPADDING',0.01,'XPADDING',0.05,'FONTSIZE',9);
            end


            formatSpec =  'For Cycle %f cyanobacteria ==> flag ==> tuna supported  %.0f +/- %.0f percent of larvae diets \n';
            fprintf(fileID,formatSpec,[Cycle, mean(z)*100, std(z)*100])
        end
        % 
        % 
        % subplot(1,8,5)  %----------- Cya ==> Clad ==> Tuna ----------------
        % hold on
        % if j==1
        %     y = Clad2TUNApercent.*Cyano2Clads./CladRespProdRatio;
        %     bar(Cycle/10,mean(y)*10^6,'FaceColor',col);
        % 
        %     set(gca,'XTick',0.5)
        %     set(gca,'XTickLabel',{'Cya==> Clad ==>Tuna'})
        %     % set(gca,'box','on')
        %     ylabel(['Nitrogen Flux',char(10),'(nmol N m^-^3 d^-^1)'])
        %     set(gca,'XTickLabelRotation',60)
        %     set(gca,'FontSize',9)
        %     if Cycle==10
        %         AddPanelLabels('a','topleft','YPADDING',0.01,'XPADDING',0.05,'FONTSIZE',9)
        %     end
        % 
        % elseif j==2
        %     tot = TUNAconsumption;
        %     y = Clad2TUNApercent.*Cyano2Clads./CladRespProdRatio;
        %     z = y./tot;
        %     bar(Cycle/10,mean(z)*100,'FaceColor',col);
        % 
        %     set(gca,'XTick',0.5);
        %     set(gca,'XTickLabel',{'Cya==> Clad ==>Tuna'});
        %     % set(gca,'box','on')
        %     %ylabel(['Nitrogen Flux',char(10),'(% of flux to larvae)']);
        %     set(gca,'XTickLabelRotation',60);
        %     set(gca,'FontSize',9);
        %     if Cycle==10
        %         AddPanelLabels('a','topleft','YPADDING',0.01,'XPADDING',0.05,'FONTSIZE',9);
        %     end
        % end
        % 
        % 
        % 
        % subplot(1,8,6)  %----------- Flag ==> Clad ==> Tuna ----------------
        % hold on
        % if j==1
        %     y = Clad2TUNApercent.*Flag2Clads./CladRespProdRatio;
        %     bar(Cycle/10,mean(y)*10^6,'FaceColor',col);
        % 
        %     set(gca,'XTick',0.5)
        %     set(gca,'XTickLabel',{'Flag==> Clad ==>Tuna'})
        %     % set(gca,'box','on')
        %     ylabel(['Nitrogen Flux',char(10),'(nmol N m^-^3 d^-^1)'])
        %     set(gca,'XTickLabelRotation',60)
        %     set(gca,'FontSize',9)
        %     if Cycle==10
        %         AddPanelLabels('a','topleft','YPADDING',0.01,'XPADDING',0.05,'FONTSIZE',9)
        %     end
        % 
        % elseif j==2
        %     tot = TUNAconsumption;
        %     y = Clad2TUNApercent.*Flag2Clads./CladRespProdRatio;
        %     z = y./tot;
        %     bar(Cycle/10,mean(z)*100,'FaceColor',col);
        % 
        %     set(gca,'XTick',0.5)
        %     set(gca,'XTickLabel',{'Flag==> Clad ==>Tuna'})
        %     % set(gca,'box','on')
        %     %ylabel(['Nitrogen Flux',char(10),'(% of flux to larvae)'])
        %     set(gca,'XTickLabelRotation',60)
        %     set(gca,'FontSize',9)
        %     if Cycle==10
        %         AddPanelLabels('a','topleft','YPADDING',0.01,'XPADDING',0.05,'FONTSIZE',9)
        %     end
        % end


        subplot(1,8,6)  %----------- Cya ==> nvmCal ==> Tuna ----------------
        hold on
        if j==1
            y = nvmCal2TUNApercent.*Cyano2nvmCals./nvmCalRespProdRatio;
            bar(Cycle/10,mean(y)*10^6,'FaceColor',col);

            set(gca,'XTick',0.5)
            set(gca,'XTickLabel',{'Cya==> nvmCal ==>Tuna'})
            % set(gca,'box','on')
            ylabel(['Nitrogen Flux',char(10),'(nmol N m^-^3 d^-^1)'])
            set(gca,'XTickLabelRotation',60)
            set(gca,'FontSize',9)
            if Cycle==10
                AddPanelLabels('f','topleft','YPADDING',0.01,'XPADDING',0.05,'FONTSIZE',9)
            end

        elseif j==2
            tot = TUNAconsumption;
            y = nvmCal2TUNApercent.*Cyano2nvmCals./nvmCalRespProdRatio;
            z = y./tot;
            bar(Cycle/10,mean(z)*100,'FaceColor',col);

            set(gca,'XTick',0.5);
            set(gca,'XTickLabel',{'Cya==> nvmCal ==>Tuna'});
            % set(gca,'box','on')
            %ylabel(['Nitrogen Flux',char(10),'(% of flux to larvae)']);
            set(gca,'XTickLabelRotation',60);
            set(gca,'FontSize',9);
            ylim([0 50])
            if Cycle==10
                AddPanelLabels('f','topleft','YPADDING',0.01,'XPADDING',0.05,'FONTSIZE',9);
            end


            formatSpec =  'For Cycle %f cyanobacteria ==> nvmCal ==> tuna supported  %.0f +/- %.0f percent of larvae diets \n';
            fprintf(fileID,formatSpec,[Cycle, mean(z)*100, std(z)*100])
        end


        subplot(1,8,7)  %----------- Flag ==> nvmCal ==> Tuna ----------------
        hold on
        if j==1
            y = nvmCal2TUNApercent.*Flag2nvmCals.*nvmCalRespProdRatio;
            bar(Cycle/10,mean(y)*10^6,'FaceColor',col);

            set(gca,'XTick',0.5)
            set(gca,'XTickLabel',{'Flag==> nvmCal ==>Tuna'})
            % set(gca,'box','on')
            ylabel(['Nitrogen Flux',char(10),'(nmol N m^-^3 d^-^1)'])
            set(gca,'XTickLabelRotation',60)
            set(gca,'FontSize',9)
            if Cycle==10
                AddPanelLabels('g','topleft','YPADDING',0.01,'XPADDING',0.05,'FONTSIZE',9)
            end

        elseif j==2
            tot = TUNAconsumption;
            y = nvmCal2TUNApercent.*Flag2nvmCals./nvmCalRespProdRatio;   %The fraction of nvmCal Secondary production consumed by tuna X the amount of nitrogen passing through flagellates that is respired by nvmCals X the ratio of respiration to secondary production for nvmCals = the amount of nitrogen that passes through nvmCals that is eventually consumed by tuna.
            z = y./tot;
            bar(Cycle/10,mean(z)*100,'FaceColor',col);

            set(gca,'XTick',0.5)
            set(gca,'XTickLabel',{'Flag==> nvmCal ==>Tuna'})
            % set(gca,'box','on')
            %ylabel(['Nitrogen Flux',char(10),'(% of flux to larvae)'])
            set(gca,'XTickLabelRotation',60)
            set(gca,'FontSize',9)
            ylim([0 50])
            if Cycle==10
                AddPanelLabels('g','topleft','YPADDING',0.01,'XPADDING',0.05,'FONTSIZE',9)
            end

            formatSpec =  'For Cycle %f flag ==> nvmCal ==> tuna supported  %.0f +/- %.0f percent of larvae diets \n';
            fprintf(fileID,formatSpec,[Cycle, mean(z)*100, std(z)*100])

            fclose(fileID)
        end


        % subplot(1,8,5)  %----------- HBac ==> Tuna ----------------
        % hold on
        % if j==1
        %     bar(Cycle/10,mean(Bac2Preflex+Bac2Postflex)*10^6,'FaceColor',col);
        % 
        %     set(gca,'XTick',0.5)
        %     set(gca,'XTickLabel',{'? ==> Bac ==> ? ==>Tuna'})
        %     % set(gca,'box','on')
        %     ylabel(['Nitrogen Flux',char(10),'(nmol N m^-^3 d^-^1)'])
        %     set(gca,'XTickLabelRotation',60)
        %     set(gca,'FontSize',9)
        %     if Cycle==10
        %         AddPanelLabels('b','topleft')
        %     end
        % 
        % elseif j==2
        %     tot = Cyano2Preflex+Cyano2Postflex + Bac2Preflex+Bac2Postflex + Diatom2Preflex+Diatom2Postflex + Tricho2Preflex+Tricho2Postflex;
        %     y = (Bac2Preflex+Bac2Postflex);
        %     z=y./tot;
        %     bar(Cycle/10,mean(z)*100,'FaceColor',col);
        % 
        %     set(gca,'XTick',0.5)
        %     set(gca,'XTickLabel',{'? ==> Bac ==> ? ==>Tuna'})
        %     % set(gca,'box','on')
        %     %ylabel(['Nitrogen Flux',char(10),'(% of flux to larvae)'])
        %     set(gca,'XTickLabelRotation',60)
        %     set(gca,'FontSize',9)
        %     if Cycle==10
        %         AddPanelLabels('a','topleft','YPADDING',0.01,'XPADDING',0.05,'FONTSIZE',9)
        %     end
        % end


    end %j


end

% 
% 
% figure(102)
% fn = ['PlotIndirect2Tuna2.Normalized'];
% exportgraphics(gcf,['../../Figures/',fn,'.png'])

figure(103)
fn = ['FoodwebPathways2Tuna.Percentages'];
exportgraphics(gcf,['../../Figures/',fn,'.png'])
```
