## Supplementary material for "Comparative food-web analysis of bluefin tuna spawning habitats in the eastern Indian Ocean and Gulf of Mexico": Code Package: IndirectFlows.m

```
function [P,struct,r] = IndirectFlows(Ae,flows)

% Note that I may want to rewrite a separate version of this code that
% treats excretion as respiration that removes energy from the ecosystem
% The excretion terms would then become part of the vector r.

% clear all
% close all
% 
% Cycle=1
% 
% load(['N15GoMInverseCycle',num2str(Cycle),'.mat'])
% load(['N15GoMInverseCycle',num2str(Cycle),'Routputs.mat'])
% clearvars -except Ae MCMCmat
% flows=mean(MCMCmat);

%This code is from methods explained in Hannon (1973)

%Calculating the number of different inputs to the ecosystem
tmp=sum(Ae);
numInputs = sum(tmp==1);
clear tmp
numCompartments = length(Ae(:,1));

%Lookup will be a tool to map the solution vector into a matrix
Lookup = zeros(numInputs+numCompartments,numInputs+numCompartments);
r_Lookup = zeros(numInputs+numCompartments,numInputs+numCompartments);
r = zeros(numInputs+numCompartments,numInputs+numCompartments);

whichinput=1;  %Need to keep track of which input we are on
whichoutput=1;  %This is obnoxious, but necessary because we have multiple outputs from some compartments

for i=1:length(Ae(1,:))
    if sum(Ae(:,i))==0
        tmp1=find(Ae(:,i)==1);
        tmp2=find(Ae(:,i)==-1);
        if Lookup(tmp2,tmp1)==0
            Lookup(tmp2,tmp1)=i;
        else
            ['Error at ',num2str(i)]
            Lookup(tmp2,tmp1)=i;
        end
    elseif sum(Ae(:,i))==1
        tmp1=find(Ae(:,i)==1);
        if Lookup(numCompartments+whichinput,tmp1)==0
            Lookup(numCompartments+whichinput,tmp1)=i;
        else
            Lookup(numCompartments+whichinput,tmp1)=i;
            ['Error at ',num2str(i)]
        end
        whichinput=whichinput+1;
    elseif sum(Ae(:,i))==-1
        tmp2=find(Ae(:,i)==-1);
        r_Lookup(tmp2,whichoutput)=i;
        whichoutput=whichoutput+1;
    end
end

P = zeros(numInputs+numCompartments,numInputs+numCompartments);
for i=1:numInputs+numCompartments
    for j=1:numInputs+numCompartments
        if Lookup(i,j)~=0
            P(i,j)=flows(Lookup(i,j));
        end
    end
end

for i=1:length(r_Lookup(:,1))
    for j=1:length(r_Lookup(1,:))
        if r_Lookup(i,j)~=0
            r(i,j)=flows(r_Lookup(i,j));
        end
    end
end
r=sum(r')';

G = zeros(numInputs+numCompartments,numInputs+numCompartments);
for i=1:numInputs+numCompartments
    if sum(P(:,i))>0
        G(:,i)=P(:,i)/sum(P(:,i));
    end
    e(i)=sum(P(:,i));
    p(i)=sum(P(i,:));
end

I = eye(numInputs+numCompartments);

struct=inv(I-G);

% (I-G)^-1 codifies the structural elements of the ecosysten since r
% can be varied to produce changes in e without changing (I-G)-�. Each
% element represents the total energy flow both directly and indirectly from the
% ith to thejth component per unit of respiration of thejth component. These
% elements are the key to understanding the interdependence of each
% component on the other.  (page 537 of Hannon (1973) )
```
