## Supplementary material for "Comparative food-web analysis of bluefin tuna spawning habitats in the eastern Indian Ocean and Gulf of Mexico": Code Package: IndirectFlowsRespiration.m

```
function [P,struct,ext,r] = IndirectFlowsRespiration(Ae_keep,flows)

%Note that this version of the code treats NH4 excretion as a loss from the ecosystem.

% clear all
% close all
% 
% Cycle=1
% load(['N15GoMInverseCycle',num2str(Cycle),'.mat'])
% load(['N15GoMInverseCycle',num2str(Cycle),'Routputs.mat'])
% Ae_keep = Ae;
% flows=mean(MCMCmat);
% clearvars -except Ae_keep flows


Cycle=0
load(['N15GoMInverseCycle',num2str(Cycle),'.mat'])
load(['N15GoMInverseCycle',num2str(Cycle),'Routputs.mat'])
[NO3s_col,NH4s_col,Cyas_col,Tris_col,Dtms_col,Flags_col,HNFs_col,MICs_col,HerbNVMs_col,Apps_col,Clads_col,nvmCals_col,Chaetos_col,Poecils_col,GelPreds_col,Plankfishs_col,Preflexs_col,Postflexs_col,Bacs_col,Dets_col,Ldets_col,Dons_col,HerbVM_col,vmCal_col,NO3d_col,NH4d_col,Cyad_col,Trid_col,Dtmd_col,Flagd_col,HNFd_col,MICd_col,HerbNVMd_col,Appd_col,Cladd_col,nvmCald_col,Chaetod_col,Poecild_col,GelPredd_col,Plankfishd_col,Bacd_col,Detd_col,Ldetd_col,Dond_col] = GetColumns(Aa,MCMCmat,del15N(1,:),d15NInputs(1,InputCol),d15NInputs(2,InputCol),d15NInputs(3,InputCol),d15NInputs(4,InputCol),d15NInputs(5,InputCol),d15NInputs(6,InputCol),d15NInputs(7,InputCol));
clearvars -except NH4s_col NH4d_col flows Ae_keep
Ae=Ae_keep;
clear Ae_keep


%This code is from methods explained in Hannon (1973)


%Calculating the number of different inputs to the ecosystem
tmp=sum(Ae);
numInputs = sum(tmp==1);
clear tmp
numCompartments = length(Ae(:,1));

%Lookup will be a tool to map the solution vector into a matrix
Lookup = zeros(numInputs+numCompartments,numInputs+numCompartments);
r_Lookup = zeros(numInputs+numCompartments,numInputs+numCompartments);
ext_Lookup = zeros(numInputs+numCompartments,numInputs+numCompartments);

whichinput=1;  %Need to keep track of which input we are on
whichoutput=1;  %This is obnoxious, but necessary because we have multiple outputs from some compartments
whichr=1; %This is obnoxious, but necessary because some organisms excrete to surface and others excrete to depth

for i=1:length(Ae(1,:))
    if sum(Ae(:,i))==0
        tmp1=find(Ae(:,i)==1);
        tmp2=find(Ae(:,i)==-1);
        if Lookup(tmp2,tmp1)==0 & tmp1~=NH4s_col & tmp1~=NH4d_col
            Lookup(tmp2,tmp1)=i;
        elseif tmp1==NH4s_col | tmp1==NH4d_col
            r_Lookup(tmp2,whichr)=i;
            whichr=whichr+1;
        else
            ['Error at ',num2str(i)]
            Lookup(tmp2,tmp1)=i;
        end
    elseif sum(Ae(:,i))==1
        tmp1=find(Ae(:,i)==1);
        if Lookup(numCompartments+whichinput,tmp1)==0
            Lookup(numCompartments+whichinput,tmp1)=i;
        else
            Lookup(numCompartments+whichinput,tmp1)=i;
            ['Error at ',num2str(i)]
        end
        whichinput=whichinput+1;
    elseif sum(Ae(:,i))==-1
        tmp2=find(Ae(:,i)==-1);
        ext_Lookup(tmp2,whichoutput)=i;
        whichoutput=whichoutput+1;
    end
end

P = zeros(numInputs+numCompartments,numInputs+numCompartments);
for i=1:numInputs+numCompartments
    for j=1:numInputs+numCompartments
        if Lookup(i,j)~=0
            P(i,j)=flows(Lookup(i,j));
        end
    end
end

ext=zeros(size(ext_Lookup));
for i=1:length(ext_Lookup(:,1))
    for j=1:length(ext_Lookup(1,:))
        if ext_Lookup(i,j)~=0
            ext(i,j)=flows(ext_Lookup(i,j));
        end
    end
end
ext=sum(ext')';

r=zeros(size(r_Lookup));
for i=1:length(r_Lookup(:,1))
    for j=1:length(r_Lookup(1,:))
        if r_Lookup(i,j)~=0
            r(i,j)=flows(r_Lookup(i,j));
        end
    end
end
r=sum(r')';

G = zeros(numInputs+numCompartments,numInputs+numCompartments);
for i=1:numInputs+numCompartments
    if sum(P(:,i))>0
        G(:,i)=P(:,i)/sum(P(:,i));
    end
    e(i)=sum(P(:,i));
    p(i)=sum(P(i,:));
end

I = eye(numInputs+numCompartments);

struct=inv(I-G);

% (I-G)^-1 codifies the structural elements of the ecosysten since r
% can be varied to produce changes in e without changing (I-G)-�. Each
% element represents the total energy flow both directly and indirectly from the
% ith to thejth component per unit of respiration of thejth component. These
% elements are the key to understanding the interdependence of each
% component on the other.  (page 537 of Hannon (1973) )
```
