## Supplementary material for "Comparative food-web analysis of bluefin tuna spawning habitats in the eastern Indian Ocean and Gulf of Mexico": Code Package: Plotdel15N2.m

```
clear all
close all

for Cycle=[1,5]


load(['N15GoMInverseCycle',num2str(Cycle),'.mat'])
load(['N15GoMInverseCycle',num2str(Cycle),'Routputs.mat'])

num=length(MCMCmat(:,1));
MCMCmat(1:round(num*0.2),:)=[];
del15N(1:round(num*0.2),:)=[];
len=length(del15N(:,1));

  del15NupNO3 = del15Nknown(1);
  del15NPreflexs = del15Nknown(2);
  del15NPostflexs = del15Nknown(3);
  del15NLdets = del15Nknown(4);
  del15NLdetd = del15Nknown(5);
  del15NDONs = del15Nknown(6);
  del15NDONd = del15Nknown(7);
  del15NApp = del15Nknown(8);
  del15NCal = del15Nknown(9);
  del15NChaeto = del15Nknown(10);
  del15NHerbVM = del15Nknown(11);
  del15NHerbNVM = del15Nknown(12);
  del15NPoecil = del15Nknown(13);
  del15NClad = del15Nknown(14);

[NO3s_col,NH4s_col,Cyas_col,Tris_col,Dtms_col,Flags_col,HNFs_col,MICs_col,HerbNVMs_col,Apps_col,Clads_col,nvmCals_col,Chaetos_col,Poecils_col,GelPreds_col,Plankfishs_col,Preflexs_col,Postflexs_col,Bacs_col,Dets_col,Ldets_col,Dons_col,HerbVM_col,vmCal_col,NO3d_col,NH4d_col,Cyad_col,Trid_col,Dtmd_col,Flagd_col,HNFd_col,MICd_col,HerbNVMd_col,Appd_col,Cladd_col,nvmCald_col,Chaetod_col,Poecild_col,GelPredd_col,Plankfishd_col,Bacd_col,Detd_col,Ldetd_col,Dond_col] = GetColumns(Aa0,MCMCmat,del15N(1,:),d15NInputs(1,InputCol),d15NInputs(2,InputCol),d15NInputs(3,InputCol),d15NInputs(4,InputCol),d15NInputs(5,InputCol),d15NInputs(6,InputCol),d15NInputs(7,InputCol));

if Cycle==1
Columns = [NO3s_col,NH4s_col,-1,Dets_col,-1,Cyas_col,Tris_col,Dtms_col,Flags_col,Bacs_col,HNFs_col,MICs_col,-1,-1,-1,-1,-1,-1,-1,Poecils_col,GelPreds_col,Plankfishs_col,-1,-1]
Columnsmid = [HerbVM_col,vmCal_col]
Columnsd = [NO3d_col,NH4d_col,-1,Detd_col,-1,Cyad_col,Trid_col,Dtmd_col,Flagd_col,Bacd_col,HNFd_col,MICd_col,-1,-1,-1,-1,-1,-1,-1,Poecild_col,GelPredd_col,Plankfishd_col,-1,-1]
Columnsmidx = [16,18];
elseif Cycle==5
Columns = [NO3s_col,NH4s_col,-1,Dets_col,-1,Cyas_col,Tris_col,Dtms_col,Flags_col,Bacs_col,HNFs_col,MICs_col,-1,-1,-1,-1,-1,-1,-1,-1,GelPreds_col,Plankfishs_col,-1,-1]
Columnsmid = [HerbVM_col,vmCal_col]
Columnsd = [NO3d_col,NH4d_col,-1,Detd_col,-1,Cyad_col,Trid_col,Dtmd_col,Flagd_col,Bacd_col,HNFd_col,MICd_col,-1,-1,-1,-1,-1,-1,-1,-1,GelPredd_col,Plankfishd_col,-1,-1]
Columnsmidx = [16,18];
end

figure('Position',[100 50 1400 600])
hold on
for i=2:2:30
    fill([i-0.5,i-0.5,i+0.5,i+0.5,i-0.5],[-20,40,40,-20,-20],[0.9 0.9 0.9],'FaceColor',[0.9 0.9 0.9],'EdgeColor',[0.9 0.9 0.9])
end
for i=1:2:30
    fill([i-0.5,i-0.5,i+0.5,i+0.5,i-0.5],[-20,40,40,-20,-20],[1 1 1],'FaceColor',[1 1 1],'EdgeColor',[1 1 1])
end
for i=-10:5:30
    plot([0 30],[i i],'-k','Color',[0.5 0.5 0.5])
end
for i=1:length(Columns)
    if Columns(i)>0
        temp=sort(del15N(:,Columns(i)));
        temp = [temp(round(len*0.025)),temp(round(len*0.25)),median(temp), temp(round(len*0.75)),temp(round(len*0.975))];
        plot([i,i]-0.15,[temp(1),temp(5)],'-k')
        fill([i-0.1,i-0.1,i+0.1,i+0.1,i-0.1]-0.15,[temp(2),temp(4),temp(4),temp(2),temp(2)],'c')
        fill([i-0.1,i-0.1,i+0.1,i+0.1,i-0.1]-0.15,[temp(2),temp(3),temp(3),temp(2),temp(2)],'c')
    end
    if Columnsd(i)>0
        temp=sort(del15N(:,Columnsd(i)));
        temp = [temp(round(len*0.025)),temp(round(len*0.25)),median(temp), temp(round(len*0.75)),temp(round(len*0.975))];
        plot([i,i]+0.15,[temp(1),temp(5)],'-k')
        fill([i-0.1,i-0.1,i+0.1,i+0.1,i-0.1]+0.15,[temp(2),temp(4),temp(4),temp(2),temp(2)],'b')
        fill([i-0.1,i-0.1,i+0.1,i+0.1,i-0.1]+0.15,[temp(2),temp(3),temp(3),temp(2),temp(2)],'b')
    end
end
for i=1:length(Columnsmid)
%     temp=sort(del15N(:,Columnsmid(i)));
%     temp = [temp(round(len*0.025)),temp(round(len*0.25)),median(temp), temp(round(len*0.75)),temp(round(len*0.975))];
%     plot([Columnsmidx(i),Columnsmidx(i)],[temp(1),temp(5)],'-k')
%     fill([Columnsmidx(i)-0.1,Columnsmidx(i)-0.1,Columnsmidx(i)+0.1,Columnsmidx(i)+0.1,Columnsmidx(i)-0.1],[temp(2),temp(4),temp(4),temp(2),temp(2)],'r')
%     fill([Columnsmidx(i)-0.1,Columnsmidx(i)-0.1,Columnsmidx(i)+0.1,Columnsmidx(i)+0.1,Columnsmidx(i)-0.1],[temp(2),temp(3),temp(3),temp(2),temp(2)],'r')
end
%Measured values
plot(3-0.15,del15NDONs,'pk','MarkerSize',12,'MarkerFaceColor','c')
plot(3+0.15,del15NDONd,'pk','MarkerSize',12,'MarkerFaceColor','b')
plot(5-0.15,del15NLdets,'pk','MarkerSize',12,'MarkerFaceColor','c')
plot(5+0.15,del15NLdetd,'pk','MarkerSize',12,'MarkerFaceColor','b')
plot(23-0.15,del15NPreflexs,'pk','MarkerSize',12,'MarkerFaceColor','c')
plot(24-0.15,del15NPostflexs,'pk','MarkerSize',12,'MarkerFaceColor','c')
plot(13,del15NApp,'pk','MarkerSize',12,'MarkerFaceColor',[0.1 0.5 1])
plot(14,del15NClad,'pk','MarkerSize',12,'MarkerFaceColor',[0.1 0.5 1])
plot(15,del15NHerbNVM,'pk','MarkerSize',12,'MarkerFaceColor',[0.1 0.5 1])
plot(16,del15NHerbVM,'pk','MarkerSize',12,'MarkerFaceColor',[0.1 0.5 1])
plot(17,del15NCal,'pk','MarkerSize',12,'MarkerFaceColor',[0.1 0.5 1])
plot(18,del15NCal,'pk','MarkerSize',12,'MarkerFaceColor',[0.1 0.5 1])
plot(19,del15NChaeto,'pk','MarkerSize',12,'MarkerFaceColor',[0.1 0.5 1])
if Cycle==5
    plot(20,del15NPoecil,'pk','MarkerSize',12,'MarkerFaceColor',[0.1 0.5 1])
end
if Cycle==5
%Legend
fill([0.7 0.7 11 11 0.7]+13,[17 24 24 17 17]-25,'w')
plot(1+13,23-25,'sc','MarkerEdgeColor','k','MarkerFaceColor','c','MarkerSize',12)
text(1.3+13,23-25,'Mixed Layer','FontSize',16)
plot(1+13,20.7-25,'sb','MarkerEdgeColor','k','MarkerFaceColor','b','MarkerSize',12)
text(1.3+13,20.7-25,'Deep Chl Max','FontSize',16)
plot(1+13,18.4-25,'sr','MarkerEdgeColor','k','MarkerFaceColor','r','MarkerSize',12)
text(1.3+13,18.4-25,'Migratory','FontSize',16)
plot([5.5 6.5]+13,[22 22]-25,'-k')
fill([5.8 5.8 6.2 6.2 5.8]+13,[22.5 21.5 21.5 22.5 22.5]-25,'w') 
fill([5.8 5.8 6 6 5.8]+13,[22.5 21.5 21.5 22.5 22.5]-25,'w') 
text(6.7+13,22-25,'Model Estimate','FontSize',16)
plot(6.2+13,19.4-25,'pk','MarkerSize',12,'MarkerFaceColor','w')
text(6.7+13,19.4-25,'Measured','FontSize',16)
end
%Limits
ylimits=[-10,15];
xlimits=[0.5,24.5];
plot([xlimits(1) xlimits(2) xlimits(2) xlimits(1) xlimits(1)],[ylimits(1) ylimits(1) ylimits(2) ylimits(2) ylimits(1)],'-k')
set(gca,'XTick',1:length(Columns))
set(gca,'XTickLabel',{'Nitrate','Ammonium','DON','Small det','Large det','Cyanobacteria','Trichodesmium','Diatoms','Mixotrophic Flagellates','Het Bacteria','Het Nanoflagellates','Microzooplankton','Appendicularians','Cladocerans','Non-migrating herbivores','Migratory herbivores','Non-migrating calanoids','Migratory Calanoids','Chaetognaths','Poecilostomatoids','Gel Predators','Planktivores','Preflex BFT','Postflex BFT'})
set(gca,'XTickLabelRotation',45)
set(gca,'box','on')
ylim(ylimits)
xlim(xlimits)
set(gca,'FontSize',18)
ylabel('\delta^1^5N')
title(['Cycle = ',num2str(Cycle)])

end
```
