## Supplementary material for "Comparative food-web analysis of bluefin tuna spawning habitats in the eastern Indian Ocean and Gulf of Mexico": Code Package: PlotIndirect2Tuna2.m

```
clear all
close all

burnin = 0.2;

for Cycle = [0,10]

    load(['N15GoMInverseCycle',num2str(Cycle),'.mat'])
    load(['N15GoMInverseCycle',num2str(Cycle),'Routputs.mat'])
    clearvars -except Ae0 MCMCmat Aa d15NInputs del15N InputCol wts Cycle burnin
    num=length(MCMCmat(:,1));
    MCMCmat(1:round(num*burnin),:)=[];
    del15N(1:round(num*burnin),:)=[];
    len=length(MCMCmat(:,1));
    alphabet = 'abcdefghijklmnop';

    for i=1:len
        i
        flows=MCMCmat(i,:)./wts';

        [P,struct,ext,r] = IndirectFlowsRespiration(Ae0,flows);

        [NO3s_col,NH4s_col,Cyas_col,Tris_col,Dtms_col,Flags_col,HNFs_col,MICs_col,HerbNVMs_col,Apps_col,Clads_col,nvmCals_col,Chaetos_col,Poecils_col,GelPreds_col,Plankfishs_col,Preflexs_col,Postflexs_col,Bacs_col,Dets_col,Ldets_col,Dons_col,HerbVM_col,vmCal_col,NO3d_col,NH4d_col,Cyad_col,Trid_col,Dtmd_col,Flagd_col,HNFd_col,MICd_col,HerbNVMd_col,Appd_col,Cladd_col,nvmCald_col,Chaetod_col,Poecild_col,GelPredd_col,Plankfishd_col,Bacd_col,Detd_col,Ldetd_col,Dond_col] = GetColumns(Aa,MCMCmat,del15N(1,:),d15NInputs(1,InputCol),d15NInputs(2,InputCol),d15NInputs(3,InputCol),d15NInputs(4,InputCol),d15NInputs(5,InputCol),d15NInputs(6,InputCol),d15NInputs(7,InputCol));

        Cyanos2Respiration=struct(Cyas_col,:)'.*r;
        CyanosProduction=sum(P(Cyas_col,:));
        Cyanos2RespirationNormalized=Cyanos2Respiration/CyanosProduction;
        Cyanod2Respiration=struct(Cyad_col,:)'.*r;
        CyanodProduction=sum(P(Cyad_col,:));
        Cyanod2RespirationNormalized=Cyanod2Respiration/CyanodProduction;
        Cyano2Preflex(i)=Cyanos2Respiration(Preflexs_col)+Cyanod2Respiration(Preflexs_col);
        Cyano2Postflex(i)=Cyanos2Respiration(Postflexs_col)+Cyanod2Respiration(Postflexs_col);
        Cyano2PreflexNormalized(i)=Cyanos2RespirationNormalized(Preflexs_col)+Cyanod2RespirationNormalized(Preflexs_col);
        Cyano2PostflexNormalized(i)=Cyanos2RespirationNormalized(Postflexs_col)+Cyanod2RespirationNormalized(Postflexs_col);
        Cyano2HNFs(i)=Cyanos2Respiration(HNFs_col)+Cyanod2Respiration(HNFs_col);
        Cyano2HNFsNormalized(i)=Cyanos2RespirationNormalized(HNFs_col)+Cyanod2RespirationNormalized(HNFs_col);
        Cyano2Mics(i)=Cyanos2Respiration(MICs_col)+Cyanod2Respiration(MICs_col);
        Cyano2MicsNormalized(i)=Cyanos2RespirationNormalized(MICs_col)+Cyanod2RespirationNormalized(MICs_col);
        Cyano2Apps(i)=Cyanos2Respiration(Apps_col)+Cyanod2Respiration(Apps_col);
        Cyano2AppsNormalized(i)=Cyanos2RespirationNormalized(Apps_col)+Cyanod2RespirationNormalized(Apps_col);
        Cyano2Clads(i)=Cyanos2Respiration(Clads_col)+Cyanod2Respiration(Clads_col);
        Cyano2CladsNormalized(i)=Cyanos2RespirationNormalized(Clads_col)+Cyanod2RespirationNormalized(Clads_col);
        Cyano2nvmCals(i)=Cyanos2Respiration(nvmCals_col)+Cyanod2Respiration(nvmCals_col);
        Cyano2nvmCalsNormalized(i)=Cyanos2RespirationNormalized(nvmCals_col)+Cyanod2RespirationNormalized(nvmCals_col);
        Cyano2Poecils(i)=Cyanos2Respiration(Poecils_col)+Cyanod2Respiration(Poecils_col);
        Cyano2PoecilsNormalized(i)=Cyanos2RespirationNormalized(Poecils_col)+Cyanod2RespirationNormalized(Poecils_col);

        Diatoms2Respiration=struct(Dtms_col,:)'.*r;
        DiatomsProduction=sum(P(Dtms_col,:));
        Diatoms2RespirationNormalized=Diatoms2Respiration/DiatomsProduction;
        Diatomd2Respiration=struct(Dtmd_col,:)'.*r;
        DiatomdProduction=sum(P(Dtmd_col,:));
        Diatomd2RespirationNormalized=Diatomd2Respiration/DiatomdProduction;
        Diatom2Preflex(i)=Diatoms2Respiration(Preflexs_col)+Diatomd2Respiration(Preflexs_col);
        Diatom2Postflex(i)=Diatoms2Respiration(Postflexs_col)+Diatomd2Respiration(Postflexs_col);
        Diatom2PreflexNormalized(i)=Diatoms2RespirationNormalized(Preflexs_col)+Diatomd2RespirationNormalized(Preflexs_col);
        Diatom2PostflexNormalized(i)=Diatoms2RespirationNormalized(Postflexs_col)+Diatomd2RespirationNormalized(Postflexs_col);
        Diatom2HNFs(i)=Diatoms2Respiration(HNFs_col)+Diatomd2Respiration(HNFs_col);
        Diatom2HNFsNormalized(i)=Diatoms2RespirationNormalized(HNFs_col)+Diatomd2RespirationNormalized(HNFs_col);
        Diatom2Mics(i)=Diatoms2Respiration(MICs_col)+Diatomd2Respiration(MICs_col);
        Diatom2MicsNormalized(i)=Diatoms2RespirationNormalized(MICs_col)+Diatomd2RespirationNormalized(MICs_col);
        Diatom2Apps(i)=Diatoms2Respiration(Apps_col)+Diatomd2Respiration(Apps_col);
        Diatom2AppsNormalized(i)=Diatoms2RespirationNormalized(Apps_col)+Diatomd2RespirationNormalized(Apps_col);
        Diatom2Clads(i)=Diatoms2Respiration(Clads_col)+Diatomd2Respiration(Clads_col);
        Diatom2CladsNormalized(i)=Diatoms2RespirationNormalized(Clads_col)+Diatomd2RespirationNormalized(Clads_col);
        Diatom2nvmCals(i)=Diatoms2Respiration(nvmCals_col)+Diatomd2Respiration(nvmCals_col);
        Diatom2nvmCalsNormalized(i)=Diatoms2RespirationNormalized(nvmCals_col)+Diatomd2RespirationNormalized(nvmCals_col);
        Diatom2Poecils(i)=Diatoms2Respiration(Poecils_col)+Diatomd2Respiration(Poecils_col);
        Diatom2PoecilsNormalized(i)=Diatoms2RespirationNormalized(Poecils_col)+Diatomd2RespirationNormalized(Poecils_col);

        Trichos2Respiration=struct(Tris_col,:)'.*r;
        TrichosProduction=sum(P(Tris_col,:));
        Trichos2RespirationNormalized=Trichos2Respiration/TrichosProduction;
        Trichod2Respiration=struct(Trid_col,:)'.*r;
        TrichodProduction=sum(P(Trid_col,:));
        Trichod2RespirationNormalized=Trichod2Respiration/TrichodProduction;
        Tricho2Preflex(i)=Trichos2Respiration(Preflexs_col)+Trichod2Respiration(Preflexs_col);
        Tricho2Postflex(i)=Trichos2Respiration(Postflexs_col)+Trichod2Respiration(Postflexs_col);
        Tricho2PreflexNormalized(i)=Trichos2RespirationNormalized(Preflexs_col)+Trichod2RespirationNormalized(Preflexs_col);
        Tricho2PostflexNormalized(i)=Trichos2RespirationNormalized(Postflexs_col)+Trichod2RespirationNormalized(Postflexs_col);
        Tricho2HNFs(i)=Trichos2Respiration(HNFs_col)+Trichod2Respiration(HNFs_col);
        Tricho2HNFsNormalized(i)=Trichos2RespirationNormalized(HNFs_col)+Trichod2RespirationNormalized(HNFs_col);
        Tricho2Mics(i)=Trichos2Respiration(MICs_col)+Trichod2Respiration(MICs_col);
        Tricho2MicsNormalized(i)=Trichos2RespirationNormalized(MICs_col)+Trichod2RespirationNormalized(MICs_col);
        Tricho2Apps(i)=Trichos2Respiration(Apps_col)+Trichod2Respiration(Apps_col);
        Tricho2AppsNormalized(i)=Trichos2RespirationNormalized(Apps_col)+Trichod2RespirationNormalized(Apps_col);
        Tricho2Clads(i)=Trichos2Respiration(Clads_col)+Trichod2Respiration(Clads_col);
        Tricho2CladsNormalized(i)=Trichos2RespirationNormalized(Clads_col)+Trichod2RespirationNormalized(Clads_col);
        Tricho2nvmCals(i)=Trichos2Respiration(nvmCals_col)+Trichod2Respiration(nvmCals_col);
        Tricho2nvmCalsNormalized(i)=Trichos2RespirationNormalized(nvmCals_col)+Trichod2RespirationNormalized(nvmCals_col);
        Tricho2Poecils(i)=Trichos2Respiration(Poecils_col)+Trichod2Respiration(Poecils_col);
        Tricho2PoecilsNormalized(i)=Trichos2RespirationNormalized(Poecils_col)+Trichod2RespirationNormalized(Poecils_col);

        Flags2Respiration=struct(Flags_col,:)'.*r;
        FlagsProduction=sum(P(Flags_col,:));
        Flags2RespirationNormalized=Flags2Respiration/FlagsProduction;
        Flagd2Respiration=struct(Flagd_col,:)'.*r;
        FlagdProduction=sum(P(Flagd_col,:));
        Flagd2RespirationNormalized=Flagd2Respiration/FlagdProduction;
        Flag2Preflex(i)=Flags2Respiration(Preflexs_col)+Flagd2Respiration(Preflexs_col);
        Flag2Postflex(i)=Flags2Respiration(Postflexs_col)+Flagd2Respiration(Postflexs_col);
        Flag2PreflexNormalized(i)=Flags2RespirationNormalized(Preflexs_col)+Flagd2RespirationNormalized(Preflexs_col);
        Flag2PostflexNormalized(i)=Flags2RespirationNormalized(Postflexs_col)+Flagd2RespirationNormalized(Postflexs_col);
        Flag2HNFs(i)=Flags2Respiration(HNFs_col)+Flagd2Respiration(HNFs_col);
        Flag2HNFsNormalized(i)=Flags2RespirationNormalized(HNFs_col)+Flagd2RespirationNormalized(HNFs_col);
        Flag2Mics(i)=Flags2Respiration(MICs_col)+Flagd2Respiration(MICs_col);
        Flag2MicsNormalized(i)=Flags2RespirationNormalized(MICs_col)+Flagd2RespirationNormalized(MICs_col);
        Flag2Apps(i)=Flags2Respiration(Apps_col)+Flagd2Respiration(Apps_col);
        Flag2AppsNormalized(i)=Flags2RespirationNormalized(Apps_col)+Flagd2RespirationNormalized(Apps_col);
        Flag2Clads(i)=Flags2Respiration(Clads_col)+Flagd2Respiration(Clads_col);
        Flag2CladsNormalized(i)=Flags2RespirationNormalized(Clads_col)+Flagd2RespirationNormalized(Clads_col);
        Flag2nvmCals(i)=Flags2Respiration(nvmCals_col)+Flagd2Respiration(nvmCals_col);
        Flag2nvmCalsNormalized(i)=Flags2RespirationNormalized(nvmCals_col)+Flagd2RespirationNormalized(nvmCals_col);
        Flag2Poecils(i)=Flags2Respiration(Poecils_col)+Flagd2Respiration(Poecils_col);
        Flag2PoecilsNormalized(i)=Flags2RespirationNormalized(Poecils_col)+Flagd2RespirationNormalized(Poecils_col);


    end

    for j=1:2
        %Figure 1 - Indirect supprt of ABT larvae

        fighandle = figure(101+j);
        fighandle.Units = 'inches';
        fighandle.Position = [1 1 8 5.5];
        col = [0 0 1];
        subplot(2,4,(Cycle)*0.4+1)

        hold on
        if j==2
            ViolinPlot(1,0.5,Tricho2Preflex*10^6,200,[0.9 0.9 0])
            ViolinPlot(2,0.5,Cyano2Preflex*10^6,200,[0.2 1 0])
            ViolinPlot(3,0.5,Flag2Preflex*10^6,200,[0 0.6 0])
            ViolinPlot(4,0.5,Diatom2Preflex*10^6,200,[0 0.4 0.5])

            ViolinPlot(7,0.5,Tricho2Postflex*10^6,200,[0.9 0.9 0])
            ViolinPlot(8,0.5,Cyano2Postflex*10^6,200,[0.2 1 0])
            ViolinPlot(9,0.5,Flag2Postflex*10^6,200,[0 0.6 0])
            ViolinPlot(10,0.5,Diatom2Postflex*10^6,200,[0 0.4 0.5])

            set(gca,'XTick',[2.5 8.5])
            set(gca,'XTickLabel',{'Preflex','Postflex'})
            set(gca,'box','on')
            ylabel(['Indirect support of ABT larvae',char(10),'(nmol N m^-^3 d^-^1)'])
            ylim([0 20])

            set(gca,'XTickLabelRotation',45)
            set(gca,'FontSize',8)
            AddPanelLabels(alphabet((Cycle)*0.3+1),'topleft')

        elseif j==1
            hold on
            ViolinPlot(1,0.5,Tricho2PreflexNormalized,200,[0.9 0.9 0])
            ViolinPlot(2,0.5,Cyano2PreflexNormalized,200,[0.2 1 0])
            ViolinPlot(3,0.5,Flag2PreflexNormalized,200,[0 0.6 0])
            ViolinPlot(4,0.5,Diatom2PreflexNormalized,200,[0 0.4 0.5])

            ViolinPlot(7,0.5,Tricho2PostflexNormalized,200,[0.9 0.9 0])
            ViolinPlot(8,0.5,Cyano2PostflexNormalized,200,[0.2 1 0])
            ViolinPlot(9,0.5,Flag2PostflexNormalized,200,[0 0.6 0])
            ViolinPlot(10,0.5,Diatom2PostflexNormalized,200,[0 0.4 0.5])

            set(gca,'XTick',[2.5 8.5])
            set(gca,'XTickLabel',{'Preflex','Postflex'})
            set(gca,'box','on')
            ylabel(['Indirect support of ABT larvae',char(10),'(Normalized to phytoplankton production)'])
            if Cycle==0
                ylim([0 0.6*10^-4])
            else
                ylim([0 0.6*10^-5])
            end
            set(gca,'XTickLabelRotation',45)
            set(gca,'FontSize',8)
            AddPanelLabels(alphabet(Cycle*0.3+1),'topleft')
        end


        %Figure 3 - Indirect support of protists
        subplot(2,4,(Cycle)*0.4+2)
        hold on
        if j==1  %First (normalized) plot
            ViolinPlot(1,0.5,Tricho2HNFsNormalized,200,[0.9 0.9 0])
            ViolinPlot(2,0.5,Cyano2HNFsNormalized,200,[0.2 1 0])
            ViolinPlot(3,0.5,Flag2HNFsNormalized,200,[0 0.6 0])
            ViolinPlot(4,0.5,Diatom2HNFsNormalized,200,[0 0.4 0.5])

            ViolinPlot(7,0.5,Tricho2MicsNormalized,200,[0.9 0.9 0])
            ViolinPlot(8,0.5,Cyano2MicsNormalized,200,[0.2 1 0])
            ViolinPlot(9,0.5,Flag2MicsNormalized,200,[0 0.6 0])
            ViolinPlot(10,0.5,Diatom2MicsNormalized,200,[0 0.4 0.5])

            ylabel(['Indirect support of heterotrophic protists',char(10),'(Normalized to phytoplankton production)'])
        else  %Second (not normalized) plot

            ViolinPlot(1,0.5,Tricho2HNFs,200,[0.9 0.9 0])
            ViolinPlot(2,0.5,Cyano2HNFs,200,[0.2 1 0])
            ViolinPlot(3,0.5,Flag2HNFs,200,[0 0.6 0])
            ViolinPlot(4,0.5,Diatom2HNFs,200,[0 0.4 0.5])

            ViolinPlot(7,0.5,Tricho2Mics,200,[0.9 0.9 0])
            ViolinPlot(8,0.5,Cyano2Mics,200,[0.2 1 0])
            ViolinPlot(9,0.5,Flag2Mics,200,[0 0.6 0])
            ViolinPlot(10,0.5,Diatom2Mics,200,[0 0.4 0.5])
            ylabel(['Indirect support of heterotrophic protists',char(10),'(mmol N m^-^3 d^-^1)'])
            if Cycle==0
                ylim([0 1])
            else
                ylim([0 0.5])
            end

        end

        set(gca,'XTick',[2.5 8.5])
        set(gca,'XTickLabel',{'Nanoflag','Microzoo'})
        set(gca,'box','on')
        set(gca,'XTickLabelRotation',45)
        set(gca,'FontSize',8)
        AddPanelLabels(alphabet(Cycle*0.3+2),'topleft')

        %Figure 4 - Indirect Support of mesozoos
        subplot(2,2,Cycle/5+2)
        hold on
        if j==1  %First fig (normalized)
            ViolinPlot(13,0.5,Tricho2AppsNormalized,200,[0.9 0.9 0])
            ViolinPlot(14,0.5,Cyano2AppsNormalized,200,[0.2 1 0])
            ViolinPlot(15,0.5,Flag2AppsNormalized,200,[0 0.6 0])
            ViolinPlot(16,0.5,Diatom2AppsNormalized,200,[0 0.4 0.5])

            ViolinPlot(19,0.5,Tricho2CladsNormalized,200,[0.9 0.9 0])
            ViolinPlot(20,0.5,Cyano2CladsNormalized,200,[0.2 1 0])
            ViolinPlot(21,0.5,Flag2CladsNormalized,200,[0 0.6 0])
            ViolinPlot(22,0.5,Diatom2CladsNormalized,200,[0 0.4 0.5])

            ViolinPlot(25,0.5,Tricho2nvmCalsNormalized,200,[0.9 0.9 0])
            ViolinPlot(26,0.5,Cyano2nvmCalsNormalized,200,[0.2 1 0])
            ViolinPlot(27,0.5,Flag2nvmCalsNormalized,200,[0 0.6 0])
            ViolinPlot(28,0.5,Diatom2nvmCalsNormalized,200,[0 0.4 0.5])

            ViolinPlot(31,0.5,Tricho2PoecilsNormalized,200,[0.9 0.9 0])
            ViolinPlot(32,0.5,Cyano2PoecilsNormalized,200,[0.2 1 0])
            ViolinPlot(33,0.5,Flag2PoecilsNormalized,200,[0 0.6 0])
            ViolinPlot(34,0.5,Diatom2PoecilsNormalized,200,[0 0.4 0.5])
            ylabel(['Indirect support of mesozooplankton',char(10),'(Normalized to phytoplankton production)'])
            if Cycle==0
                ylim([0 0.12])
            end
        else  %Second fig (not normalized)

            ViolinPlot(13,0.5,Tricho2Apps,200,[0.9 0.9 0])
            ViolinPlot(14,0.5,Cyano2Apps,200,[0.2 1 0])
            ViolinPlot(15,0.5,Flag2Apps,200,[0 0.6 0])
            ViolinPlot(16,0.5,Diatom2Apps,200,[0 0.4 0.5])

            ViolinPlot(19,0.5,Tricho2Clads,200,[0.9 0.9 0])
            ViolinPlot(20,0.5,Cyano2Clads,200,[0.2 1 0])
            ViolinPlot(21,0.5,Flag2Clads,200,[0 0.6 0])
            ViolinPlot(22,0.5,Diatom2Clads,200,[0 0.4 0.5])

            ViolinPlot(25,0.5,Tricho2nvmCals,200,[0.9 0.9 0])
            ViolinPlot(26,0.5,Cyano2nvmCals,200,[0.2 1 0])
            ViolinPlot(27,0.5,Flag2nvmCals,200,[0 0.6 0])
            ViolinPlot(28,0.5,Diatom2nvmCals,200,[0 0.4 0.5])

            ViolinPlot(31,0.5,Tricho2Poecils,200,[0.9 0.9 0])
            ViolinPlot(32,0.5,Cyano2Poecils,200,[0.2 1 0])
            ViolinPlot(33,0.5,Flag2Poecils,200,[0 0.6 0])
            ViolinPlot(34,0.5,Diatom2Poecils,200,[0 0.4 0.5])
            ylabel(['Indirect support of mesozooplankton',char(10),'(mmol N m^-^3 d^-^1)'])
            if Cycle==0
                ylim([0 0.2])
            end
        end

        if Cycle==0
            if j==1
                upper=0.117;
                lower=0.077;
            else
                upper=0.197;
                lower=0.13;
            end
            fill([12.9 22 22 12.9 12.9],[lower lower upper upper lower],'w')
            fill([13.3 14.2 14.2 13.3 13.3],[upper-(upper-lower)*1/9 upper-(upper-lower)*1/9 upper-(upper-lower)*2/9 upper-(upper-lower)*2/9  upper-(upper-lower)*1/9],'r','FaceColor',[0.9 0.9 0])
            text(14.4,upper-(upper-lower)*1.5/9,'Tricho','FontSize',8)
            fill([13.3 14.2 14.2 13.3 13.3],[upper-(upper-lower)*3/9 upper-(upper-lower)*3/9 upper-(upper-lower)*4/9 upper-(upper-lower)*4/9  upper-(upper-lower)*3/9],'r','FaceColor',[0.2 1 0])
            text(14.4,upper-(upper-lower)*3.5/9,'Picophy','FontSize',8)
            fill([13.3 14.2 14.2 13.3 13.3],[upper-(upper-lower)*5/9 upper-(upper-lower)*5/9 upper-(upper-lower)*6/9 upper-(upper-lower)*6/9  upper-(upper-lower)*5/9],'r','FaceColor',[0 0.6 0])
            text(14.4,upper-(upper-lower)*5.5/9,'Flagellates','FontSize',8)
            fill([13.3 14.2 14.2 13.3 13.3],[upper-(upper-lower)*7/9 upper-(upper-lower)*7/9 upper-(upper-lower)*8/9 upper-(upper-lower)*8/9  upper-(upper-lower)*7/9],'r','FaceColor',[0 0.4 0.5])
            text(14.4,upper-(upper-lower)*7.5/9,'Diatoms','FontSize',8)
        end

        set(gca,'XTick',[14.5 20.5 26.5 32.5])
        set(gca,'XTickLabel',{'Appendicularians','Cladocerans','Calanoids (NVM)','Poecilostomatoids'})
        set(gca,'box','on')
        set(gca,'XTickLabelRotation',45)
        set(gca,'FontSize',8)
        xlim([12.5 34.5])
        AddPanelLabels(alphabet(Cycle*0.3+3),'topright')

    end %j


    Cyano2Preflex=sort(Cyano2Preflex);
    Flag2Preflex=sort(Flag2Preflex);
    Diatom2Preflex=sort(Diatom2Preflex);
    ['Preflexion ABT excreted ',num2str(mean(Flag2Preflex)*10^6,2),' nmol N m^-^2 d^-^1 (',num2str(Flag2Preflex(round(0.025*len))*10^6,2),' - ',...
        num2str(Flag2Preflex(round(0.975*len))*10^6,2),'  nmol N m^-^2 d^-^1)',char(10),'derived from the production of flagellates,    ',...
        num2str(mean(Cyano2Preflex)*10^6,2),' nmol N m^-^2 d^-^1 (',num2str(Cyano2Preflex(round(0.025*len))*10^6,2),' - ',...
        num2str(Cyano2Preflex(round(0.975*len))*10^6,2),')',char(10),'from picophytoplankton, and ',num2str(mean(Diatom2Preflex)*10^6,2),' nmol N m^-^2 d^-^1 (',...
        num2str(Diatom2Preflex(round(0.025*len))*10^6,2),' - ',num2str(Diatom2Preflex(round(0.975*len))*10^6,2),')',char(10),'nmol N m^-^2 d^-^1 from diatoms.']

    Cyano2Postflex=sort(Cyano2Postflex);
    Flag2Postflex=sort(Flag2Postflex);
    Diatom2Postflex=sort(Diatom2Postflex);
    ['Postflexion larvae excreted ',num2str(mean(Flag2Postflex)*10^6,2),' nmol N m^-^2 d^-^1 (',num2str(Flag2Postflex(round(0.025*len))*10^6,2),' - ',...
        num2str(Flag2Postflex(round(0.975*len))*10^6,2),'  nmol N m^-^2 d^-^1)',char(10),'from flagellates,    ',...
        num2str(mean(Cyano2Postflex)*10^6,2),' nmol N m^-^2 d^-^1 (',num2str(Cyano2Postflex(round(0.025*len))*10^6,2),' - ',...
        num2str(Cyano2Postflex(round(0.975*len))*10^6,2),')',char(10),'from picophytoplankton, and ',num2str(mean(Diatom2Postflex)*10^6,2),' nmol N m^-^2 d^-^1 (',...
        num2str(Diatom2Postflex(round(0.025*len))*10^6,2),' - ',num2str(Diatom2Postflex(round(0.975*len))*10^6,2),')',char(10),'nmol N m^-^2 d^-^1 from diatoms.']

    CyanosProduction=sort(CyanosProduction);
    DiatomsProduction=sort(DiatomsProduction);
    FlagsProduction=sort(FlagsProduction);
    ['flagellate, picophytoplankton, and diatom average NPP in the shallow',char(10),'euphotic zone were ',num2str(mean(FlagsProduction),2),', ',...
        num2str(mean(CyanosProduction),2),', and ',num2str(mean(DiatomsProduction),2),' mmol N m-2 d-1, respectively']


    Cyano2PreflexNormalized=sort(Cyano2PreflexNormalized);
    Flag2PreflexNormalized=sort(Flag2PreflexNormalized);
    Diatom2PreflexNormalized=sort(Diatom2PreflexNormalized);
    Tricho2PreflexNormalized=sort(Tricho2PreflexNormalized);
    ['Preflexion ABT excreted ',num2str(mean(Diatom2PreflexNormalized)*100,2),'% (',num2str(Diatom2PreflexNormalized(round(0.025*len))*100,2),' - ',...
        num2str(Diatom2PreflexNormalized(round(0.975*len))*100,2),'%)',char(10),' of diatom NPP and ',...
        num2str(mean(Flag2PreflexNormalized)*100,2),'% (',...
        num2str(Flag2PreflexNormalized(round(0.025*len))*100,2),' - ',num2str(Flag2PreflexNormalized(round(0.975*len))*100,2),'%)',char(10),' of flagellate NPP, compared to ',...
        num2str(mean(Cyano2PreflexNormalized)*100,2),'% (',num2str(Cyano2PreflexNormalized(round(0.025*len))*100,2),' - ',...
        num2str(Cyano2PreflexNormalized(round(0.975*len))*100,2),'%)',char(10),'from picophytoplankton and ',...
        num2str(mean(Tricho2PreflexNormalized)*100,2),'% (',num2str(Tricho2PreflexNormalized(round(0.025*len))*100,2),' - ',...
        num2str(Tricho2PreflexNormalized(round(0.975*len))*100,2),'%) from Trichodesmium']


    Cyano2PostflexNormalized=sort(Cyano2PostflexNormalized);
    Flag2PostflexNormalized=sort(Flag2PostflexNormalized);
    Diatom2PostflexNormalized=sort(Diatom2PostflexNormalized);
    Tricho2PostflexNormalized=sort(Tricho2PostflexNormalized);
    ['Postflexion ABT excreted ',num2str(mean(Diatom2PostflexNormalized)*100,2),'% (',num2str(Diatom2PostflexNormalized(round(0.025*len))*100,2),' - ',...
        num2str(Diatom2PostflexNormalized(round(0.975*len))*100,2),'%)',char(10),' of diatom NPP and ',...
        num2str(mean(Flag2PostflexNormalized)*100,2),'% (',...
        num2str(Flag2PostflexNormalized(round(0.025*len))*100,2),' - ',num2str(Flag2PostflexNormalized(round(0.975*len))*100,2),'%)',char(10),' of flagellate NPP, compaostd to ',...
        num2str(mean(Cyano2PostflexNormalized)*100,2),'% (',num2str(Cyano2PostflexNormalized(round(0.025*len))*100,2),' - ',...
        num2str(Cyano2PostflexNormalized(round(0.975*len))*100,2),'%)',char(10),'from picophytoplankton and ',...
        num2str(mean(Tricho2PostflexNormalized)*100,2),'% (',num2str(Tricho2PostflexNormalized(round(0.025*len))*100,2),' - ',...
        num2str(Tricho2PostflexNormalized(round(0.975*len))*100,2),'%) from Trichodesmium']


    Cyano2HNFsNormalized=sort(Cyano2HNFsNormalized);
    Flag2MicsNormalized=sort(Flag2MicsNormalized);

    ['Heterotrophic nanoflagellates respired ',num2str(mean(Cyano2HNFsNormalized*100),2),'% of picophytoplankton production, and',char(10),...
        'microzooplankton respired ',num2str(mean(Flag2MicsNormalized*100),2),'% of flagellate production']


end


figure(102)
fn = ['PlotIndirect2Tuna2.Normalized'];
exportgraphics(gcf,['../../Figures/',fn,'.png'])


figure(103)
fn = ['PlotIndirect2Tuna2'];
exportgraphics(gcf,['../../Figures/',fn,'.png'])
```
