## Supplementary material for "Comparative food-web analysis of bluefin tuna spawning habitats in the eastern Indian Ocean and Gulf of Mexico": Code Package: PlotIndirectFlows.m

```
clear all
close all

Cycle=1

load(['N15GoMInverseCycle',num2str(Cycle),'.mat'])
load(['N15GoMInverseCycle',num2str(Cycle),'Routputs.mat'])
clearvars -except Ae0 MCMCmat Aa d15NInputs del15N InputCol wts
flows=mean(MCMCmat)./wts';

[P,struct,ext,r] = IndirectFlowsRespiration(Ae0,flows);

[NO3s_col,NH4s_col,Cyas_col,Tris_col,Dtms_col,Flags_col,HNFs_col,MICs_col,HerbNVMs_col,Apps_col,Clads_col,nvmCals_col,Chaetos_col,Poecils_col,GelPreds_col,Plankfishs_col,Preflexs_col,Postflexs_col,Bacs_col,Dets_col,Ldets_col,Dons_col,HerbVM_col,vmCal_col,NO3d_col,NH4d_col,Cyad_col,Trid_col,Dtmd_col,Flagd_col,HNFd_col,MICd_col,HerbNVMd_col,Appd_col,Cladd_col,nvmCald_col,Chaetod_col,Poecild_col,GelPredd_col,Plankfishd_col,Bacd_col,Detd_col,Ldetd_col,Dond_col] = GetColumns(Aa,MCMCmat,del15N(1,:),d15NInputs(1,InputCol),d15NInputs(2,InputCol),d15NInputs(3,InputCol),d15NInputs(4,InputCol),d15NInputs(5,InputCol),d15NInputs(6,InputCol),d15NInputs(7,InputCol));

Cyano2Respiration=struct(Cyas_col,:)'.*r;
CyanoProduction=sum(P(Cyas_col,:));
Cyano2RespirationNormalized=Cyano2Respiration/CyanoProduction;

Cyano2RespirationZL=sum(Cyano2Respiration([HerbNVMs_col,Apps_col,Clads_col,nvmCals_col,Chaetos_col,Poecils_col,HerbVM_col,vmCal_col]));
Cyano2Herb=sum(Cyano2Respiration([HerbNVMs_col,HerbVM_col]))/Cyano2RespirationZL;
Cyano2Cal=sum(Cyano2Respiration([nvmCals_col,vmCal_col]))/Cyano2RespirationZL;
Cyano2App=Cyano2Respiration(Apps_col)/Cyano2RespirationZL;
Cyano2Clad=Cyano2Respiration(Clads_col)/Cyano2RespirationZL;
Cyano2Poecil=Cyano2Respiration(Poecils_col)/Cyano2RespirationZL;


Dtm2Respiration=struct(Dtms_col,:)'.*r;
DtmProduction=sum(P(Dtms_col,:));
Dtm2RespirationNormalized=Dtm2Respiration/DtmProduction;

Dtm2RespirationZL=sum(Dtm2Respiration([HerbNVMs_col,Apps_col,Clads_col,nvmCals_col,Chaetos_col,Poecils_col,HerbVM_col,vmCal_col]));
Dtm2Herb=sum(Dtm2Respiration([HerbNVMs_col,HerbVM_col]))/Dtm2RespirationZL;
Dtm2Cal=sum(Dtm2Respiration([nvmCals_col,vmCal_col]))/Dtm2RespirationZL;
Dtm2App=Dtm2Respiration(Apps_col)/Dtm2RespirationZL;
Dtm2Clad=Dtm2Respiration(Clads_col)/Dtm2RespirationZL;
Dtm2Poecil=Dtm2Respiration(Poecils_col)/Dtm2RespirationZL;


Flag2Respiration=struct(Flags_col,:)'.*r;
FlagProduction=sum(P(Flags_col,:));
Flag2RespirationNormalized=Flag2Respiration/FlagProduction;

Flag2RespirationZL=sum(Flag2Respiration([HerbNVMs_col,Apps_col,Clads_col,nvmCals_col,Chaetos_col,Poecils_col,HerbVM_col,vmCal_col]));
Flag2Herb=sum(Flag2Respiration([HerbNVMs_col,HerbVM_col]))/Flag2RespirationZL;
Flag2Cal=sum(Flag2Respiration([nvmCals_col,vmCal_col]))/Flag2RespirationZL;
Flag2App=Flag2Respiration(Apps_col)/Flag2RespirationZL;
Flag2Clad=Flag2Respiration(Clads_col)/Flag2RespirationZL;
Flag2Poecil=Flag2Respiration(Poecils_col)/Flag2RespirationZL;


[output] = FoodWebIndirectFigure('Cya',Cyano2Herb,Cyano2Cal,Cyano2Clad,Cyano2App,Cyano2Poecil)

[output] = FoodWebIndirectFigure('Dtm',Dtm2Herb,Dtm2Cal,Dtm2Clad,Dtm2App,Dtm2Poecil)
```
