## Supplementary material for "Comparative food-web analysis of bluefin tuna spawning habitats in the eastern Indian Ocean and Gulf of Mexico": Code Package: PlotIndirectNitrogen2Tuna.m

```
clear all
close all

Cycle=5

load(['N15GoMInverseCycle',num2str(Cycle),'.mat'])
load(['N15GoMInverseCycle',num2str(Cycle),'Routputs.mat'])
clearvars -except Ae0 MCMCmat Aa d15NInputs del15N InputCol wts Cycle
num=length(MCMCmat(:,1));
MCMCmat(1:round(num*0.2),:)=[];
del15N(1:round(num*0.2),:)=[];
len=length(MCMCmat(:,1));

for i=1:len
    i
    flows=MCMCmat(i,:)./wts';
    
    [P,struct,r] = IndirectFlows(Ae0,flows);
    
    [NO3s_col,NH4s_col,Cyas_col,Tris_col,Dtms_col,Flags_col,HNFs_col,MICs_col,HerbNVMs_col,Apps_col,Clads_col,nvmCals_col,Chaetos_col,Poecils_col,GelPreds_col,Plankfishs_col,Preflexs_col,Postflexs_col,Bacs_col,Dets_col,Ldets_col,Dons_col,HerbVM_col,vmCal_col,NO3d_col,NH4d_col,Cyad_col,Trid_col,Dtmd_col,Flagd_col,HNFd_col,MICd_col,HerbNVMd_col,Appd_col,Cladd_col,nvmCald_col,Chaetod_col,Poecild_col,GelPredd_col,Plankfishd_col,Bacd_col,Detd_col,Ldetd_col,Dond_col] = GetColumns(Aa,MCMCmat,del15N(1,:),d15NInputs(1,InputCol),d15NInputs(2,InputCol),d15NInputs(3,InputCol),d15NInputs(4,InputCol),d15NInputs(5,InputCol),d15NInputs(6,InputCol),d15NInputs(7,InputCol));
    
    ToNO3d = find(Ae0(NO3d_col,:)==1);
    FromNO3d = find(Ae0(NO3d_col,:)==-1);
    ToNO3s = find(Ae0(NO3s_col,:)==1);
    UpwelledNO3i = ToNO3d(find(sum(Ae0(:,ToNO3d))==1));
    UpwelledShallowNO3i = intersect(FromNO3d,ToNO3s);
    UpwelledNO3 = flows(UpwelledNO3i);            %This is the total amount of upwelling to deep nitrate
    UpwelledNO3Shallow(i) = flows(UpwelledShallowNO3i);            %This is the total amount of upwelling to shallow nitrate
    NO3dProduction=sum(P(NO3d_col,:));            %This is the total production of deep nitrate
    FracUpwelled(i)=UpwelledNO3/NO3dProduction;   %This is the fraction of deep nitrate that came from upwelling
    NO3d2Export=struct(NO3d_col,:)'.*r;
    NO3d2Preflex(i)=NO3d2Export(Preflexs_col);    %This is the total amount of nitrate that goes to preflexion larvae (although I'm not sure if this is total amount exported by Preflex, respired by Preflex, or just gotten by preflex)  but it has units of mmol N m-2 d-1
    NO3d2Postflex(i)=NO3d2Export(Postflexs_col);
    Upwelling2Preflex(i)=NO3d2Preflex(i)*FracUpwelled(i);
    Upwelling2Postflex(i)=NO3d2Postflex(i)*FracUpwelled(i);
    
    ToTrid = find(Ae0(Trid_col,:)==1);
    TrichoNFixi = ToTrid(find(sum(Ae0(:,ToTrid))==1));
    TrichoNFix = flows(TrichoNFixi);          
    TridProduction=sum(P(Trid_col,:));          
    FracTridN2Fix(i)=TrichoNFix/TridProduction;   
    Trid2Export=struct(Trid_col,:)'.*r;
    Trid2Preflex(i)=Trid2Export(Preflexs_col);
    Trid2Postflex(i)=Trid2Export(Postflexs_col);
    TridNFix2Preflex(i)=Trid2Preflex(i)*FracTridN2Fix(i);
    TridNFix2Postflex(i)=Trid2Postflex(i)*FracTridN2Fix(i);
    
    ToTris = find(Ae0(Tris_col,:)==1);
    TrichoNFixsi = ToTris(find(sum(Ae0(:,ToTris))==1));
    TrichoNFixs = flows(TrichoNFixsi);        
    TrisProduction=sum(P(Tris_col,:));          
    FracTrisN2Fix(i)=TrichoNFixs/TrisProduction;   
    Tris2Export=struct(Tris_col,:)'.*r;
    Tris2Preflex(i)=Tris2Export(Preflexs_col);
    Tris2Postflex(i)=Tris2Export(Postflexs_col);
    TrisNFix2Preflex(i)=Tris2Preflex(i)*FracTrisN2Fix(i);
    TrisNFix2Postflex(i)=Tris2Postflex(i)*FracTrisN2Fix(i);
    
    ToCyad = find(Ae0(Cyad_col,:)==1);
    CyanoNFixi = ToCyad(find(sum(Ae0(:,ToCyad))==1));
    CyanoNFix = flows(CyanoNFixi);        
    CyadProduction=sum(P(Cyad_col,:));          
    FracCyadN2Fix(i)=CyanoNFix/CyadProduction;   
    Cyad2Export=struct(Cyad_col,:)'.*r;
    Cyad2Preflex(i)=Cyad2Export(Preflexs_col);
    Cyad2Postflex(i)=Cyad2Export(Postflexs_col);
    CyadNFix2Preflex(i)=Cyad2Preflex(i)*FracCyadN2Fix(i);
    CyadNFix2Postflex(i)=Cyad2Postflex(i)*FracCyadN2Fix(i);
    
    ToCyas = find(Ae0(Cyas_col,:)==1);
    CyanoNFixsi = ToCyas(find(sum(Ae0(:,ToCyas))==1));
    CyanoNFixs = flows(CyanoNFixsi);     
    CyasProduction=sum(P(Cyas_col,:));          
    FracCyasN2Fix(i)=CyanoNFixs/CyasProduction;  
    Cyas2Export=struct(Cyas_col,:)'.*r;
    Cyas2Preflex(i)=Cyas2Export(Preflexs_col);
    Cyas2Postflex(i)=Cyas2Export(Postflexs_col);
    CyasNFix2Preflex(i)=Cyas2Preflex(i)*FracCyasN2Fix(i);
    CyasNFix2Postflex(i)=Cyas2Postflex(i)*FracCyasN2Fix(i);
    
    NFixs(i) = CyanoNFixs + TrichoNFixs;
    
    NFix2Preflex(i) = TridNFix2Preflex(i) + TrisNFix2Preflex(i) + CyadNFix2Preflex(i) + CyasNFix2Preflex(i);
    NFix2Postflex(i) = TridNFix2Postflex(i) + TrisNFix2Postflex(i) + CyadNFix2Postflex(i) + CyasNFix2Postflex(i);
    
    ToDons = find(Ae0(Dons_col,:)==1);
    LateralDONi = ToDons(find(sum(Ae0(:,ToDons))==1));
    LateralDON = flows(LateralDONi);     
    DonsProduction=sum(P(Dons_col,:));          
    FracDonsLateral(i)=LateralDON/DonsProduction;  
    Dons2Export=struct(Dons_col,:)'.*r;
    Dons2Preflex(i)=Dons2Export(Preflexs_col);
    Dons2Postflex(i)=Dons2Export(Postflexs_col);
    LateralDONFlux(i)=LateralDON;
    
    ToDets = find(Ae0(Dets_col,:)==1);
    LateralDeti = ToDets(find(sum(Ae0(:,ToDets))==1));
    LateralDet = flows(LateralDeti);   
    DetsProduction=sum(P(Dets_col,:));          
    FracDetsLateral(i)=LateralDet/DetsProduction;  
    Dets2Export=struct(Dets_col,:)'.*r;
    Dets2Preflex(i)=Dets2Export(Preflexs_col);
    Dets2Postflex(i)=Dets2Export(Postflexs_col);
    LateralDetFlux(i)=LateralDet;
    
    Lateral2Preflex(i) = Dons2Preflex(i) + Dets2Preflex(i);
    Lateral2Postflex(i) = Dons2Postflex(i) + Dets2Postflex(i);
    
    FracLateralPreflex(i)= Lateral2Preflex(i)/(Lateral2Preflex(i)+NFix2Preflex(i)+Upwelling2Preflex(i));
    FracLateralPostflex(i)= Lateral2Postflex(i)/(Lateral2Postflex(i)+NFix2Postflex(i)+Upwelling2Postflex(i));
    
    FracNFixPreflex(i)= NFix2Preflex(i)/(Lateral2Preflex(i)+NFix2Preflex(i)+Upwelling2Preflex(i));
    FracNFixPostflex(i)= NFix2Postflex(i)/(Lateral2Postflex(i)+NFix2Postflex(i)+Upwelling2Postflex(i));
    
    FracUpwellingPreflex(i)= Upwelling2Preflex(i)/(Lateral2Preflex(i)+NFix2Preflex(i)+Upwelling2Preflex(i));
    FracUpwellingPostflex(i)= Upwelling2Postflex(i)/(Lateral2Postflex(i)+NFix2Postflex(i)+Upwelling2Postflex(i));
    
    ToBacs=find(Ae0(Bacs_col,:)==1);
    FromDons = find(Ae0(Dons_col,:)==-1);
    FromBacs=find(Ae0(Bacs_col,:)==-1);
    ToNH4s = find(Ae0(NH4s_col,:)==1);
    BacUptakei=intersect(FromDons,ToBacs);
    BacRespirationi=intersect(FromBacs,ToNH4s);
    BacUptake(i)=flows(BacUptakei);
    BacRespiration(i)=flows(BacRespirationi);
    BacGGE(i)=1-BacRespiration(i)/BacUptake(i);
    
    FromNH4s = find(Ae0(NH4s_col,:)==-1);
    ShallowNitrificationi=intersect(FromNH4s,ToNO3s);
    ShallowNitrification(i)=flows(ShallowNitrificationi);
end


%Figure 1 - Indirect supprt of ABT larvae
figure('Position',[50 50 800 600])
hold on
ViolinPlot(1,0.5,FracLateralPreflex,200,[0.5 0 0])
ViolinPlot(2,0.5,FracNFixPreflex,200,[0.9 0.9 0])
ViolinPlot(3,0.5,FracUpwellingPreflex,200,[0 0.6 0])

ViolinPlot(7,0.5,FracLateralPostflex,200,[0.5 0 0])
ViolinPlot(8,0.5,FracNFixPostflex,200,[0.9 0.9 0])
ViolinPlot(9,0.5,FracUpwellingPostflex,200,[0 0.6 0])

set(gca,'XTick',[2.5 8.5])
set(gca,'XTickLabel',{'Preflex','Postflex'})
set(gca,'box','on')
ylabel(['Indirect support of ABT larvae',char(10),'(mmol N m^-^3 d^-^1)'])
ylim([0 1])
xlim([0 16.5])
set(gca,'XTickLabelRotation',45)
set(gca,'FontSize',17)

upper=0.97;
lower=0.7;
fill([9 16 16 9 9],[lower lower upper upper lower],'w')
fill([9.2 10 10 9.2 9.2],[upper-(upper-lower)*1/9 upper-(upper-lower)*1/9 upper-(upper-lower)*2/9 upper-(upper-lower)*2/9  upper-(upper-lower)*1/9],'r','FaceColor',[0.5 0 0])
text(10.4,upper-(upper-lower)*1.5/9,'Lateral Transport','FontSize',16)
fill([9.2 10 10 9.2 9.2],[upper-(upper-lower)*3/9 upper-(upper-lower)*3/9 upper-(upper-lower)*4/9 upper-(upper-lower)*4/9  upper-(upper-lower)*3/9],'r','FaceColor',[0.9 0.9 0])
text(10.4,upper-(upper-lower)*3.5/9,'N2 Fixation','FontSize',15)
fill([9.2 10 10 9.2 9.2],[upper-(upper-lower)*5/9 upper-(upper-lower)*5/9 upper-(upper-lower)*6/9 upper-(upper-lower)*6/9  upper-(upper-lower)*5/9],'r','FaceColor',[0 0.6 0])
text(10.4,upper-(upper-lower)*5.5/9,'Upwelling','FontSize',16)


FracLateralPreflex=sort(FracLateralPreflex);
FracLateralPostflex=sort(FracLateralPostflex);

['Bluefin tuna production was supported primarily by nitrogen derived from lateral transport',char(10),...
    'Preflexion Larvae derived ',num2str(mean(FracLateralPreflex*100),3),'% (',num2str(FracLateralPreflex(round(len*0.025))*100,3),' - ',num2str(FracLateralPreflex(round(len*0.975))*100,3),'%) and',char(10),...
    'Postflexion Larvae derived ',num2str(mean(FracLateralPostflex*100),3),'% (',num2str(FracLateralPostflex(round(len*0.025))*100,3),' - ',num2str(FracLateralPostflex(round(len*0.975))*100,3),'%) of their N from lateral transport'    ]

FracNFixPreflex=sort(FracNFixPreflex);
FracNFixPostflex=sort(FracNFixPostflex);
['Preflexion Larvae derived ',num2str(mean(FracNFixPreflex*100),3),'% (',num2str(FracNFixPreflex(round(len*0.025))*100,3),' - ',num2str(FracNFixPreflex(round(len*0.975))*100,3),'%) and',char(10),...
    'Postflexion Larvae derived ',num2str(mean(FracNFixPostflex*100),3),'% (',num2str(FracNFixPostflex(round(len*0.025))*100,3),' - ',num2str(FracNFixPostflex(round(len*0.975))*100,3),'%) of their N from N2 fixation'    ]


FracUpwellingPreflex=sort(FracUpwellingPreflex);
FracUpwellingPostflex=sort(FracUpwellingPostflex);
['Preflexion Larvae derived ',num2str(mean(FracUpwellingPreflex*100),3),'% (',num2str(FracUpwellingPreflex(round(len*0.025))*100,3),' - ',num2str(FracUpwellingPreflex(round(len*0.975))*100,3),'%) and',char(10),...
    'Postflexion Larvae derived ',num2str(mean(FracUpwellingPostflex*100),3),'% (',num2str(FracUpwellingPostflex(round(len*0.025))*100,3),' - ',num2str(FracUpwellingPostflex(round(len*0.975))*100,3),'%) of their N from Nitrate upwelled from beneath the euphotic zone'    ]


UpwelledNO3Shallow=sort(UpwelledNO3Shallow);
['the ecosystem is not substantially supported by locally-upwelled nitrate',char(10),'(which supplied ',num2str(mean(UpwelledNO3Shallow),2),...
    ' (',num2str(UpwelledNO3Shallow(round(len*0.025)),2),' - ',num2str(UpwelledNO3Shallow(round(len*0.975)),2),')',char(10),' mmol N m-2 d-1 ',...
    'to the upper euphotic zone during C',num2str(Cycle)]
    

NFixs = sort(NFixs);
['or by nitrogen fixation (which supplied ',num2str(mean(NFixs),2),...
    ' (',num2str(NFixs(round(len*0.025)),2),' - ',num2str(NFixs(round(len*0.975)),2),')',...
    char(10),' mmol N m-2 d-1 to the upper euphotic zone during C',num2str(Cycle)]


LateralDetFlux=sort(LateralDetFlux);
['Rather, nitrogen entered the ecosystem primarily through lateral advection of PON',char(10),'(',num2str(mean(LateralDetFlux),2),...
    ' (',num2str(LateralDetFlux(round(len*0.025)),2),' - ',num2str(LateralDetFlux(round(len*0.975)),2),') mmol N m-2 d-1)']


LateralDONFlux=sort(LateralDONFlux);
[' and DON (',num2str(mean(LateralDONFlux),2),...
    ' (',num2str(LateralDONFlux(round(len*0.025)),2),' - ',num2str(LateralDONFlux(round(len*0.975)),2),') mmol N m-2 d-1) during C',num2str(Cycle)]


BacGGE=sort(BacGGE);
['bacteria had a gross growth efficiency of only ',num2str(100*mean(BacGGE),2),...
    '% (',num2str(100*BacGGE(round(len*0.025)),2),' - ',num2str(100*BacGGE(round(len*0.975)),2),'%) during C',num2str(Cycle)]


ShallowNitrification=sort(ShallowNitrification);
['Nitrification rates in the upper euphotic zone were ',num2str(mean(ShallowNitrification),2),...
    ' (',num2str(ShallowNitrification(round(len*0.025)),2),' - ',num2str(ShallowNitrification(round(len*0.975)),2),' mmol N m-2 d-1 during C',num2str(Cycle)]
['This equates to nitrification rates of ',num2str(mean(ShallowNitrification)/50*10^3,2),' nmol N L-1 d-1, respectively']


data=xlsread('..\..\..\2018 Cruise\Nitrate Uptake\NF17&18 - In Situ 15NO3 and H13CO3 - 10-24-19.xlsx','Nutrients','V15:Z16');
Nutrientdata = array2table(data,'VariableNames',{'Cycle','UEZ_NH4','LEZ_NH4','UEZ_NO3','LEZ_NO3'}); %Units are mmol / m2

if Cycle==1
NO3UEZ=data(Cycle,4);
NH4UEZ=data(Cycle,2);
elseif Cycle==5
NO3UEZ=data(2,4);
NH4UEZ=data(2,2);
end
    

['Nitrate in the surface ocean was able to be regenerated every ',num2str(NO3UEZ/mean(ShallowNitrification),2),' days during C',num2str(Cycle)]

['this suggests a specific ammonium-oxidation rate of ',num2str(mean(ShallowNitrification)/NH4UEZ,2),' d-1 during C',num2str(Cycle)]
```
