## Supplementary material for "Comparative food-web analysis of bluefin tuna spawning habitats in the eastern Indian Ocean and Gulf of Mexico": Code Package: PlotMeasurements2Cycles.m

```
clearvars
close all

burnin = 0.2;

for Cycle=[0,10]
    clearvars -except Cycle burnin RateTable0 MeanTable KEEP

    load(['N15GoMInverseCycle',num2str(Cycle),'.mat'])
    load(['N15GoMInverseCycle',num2str(Cycle),'Routputs.mat'])
    num=length(MCMCmat(:,1));
    MCMCmat(1:round(num*burnin),:)=[];
    del15N(1:round(num*burnin),:)=[];
    %     [num,txt,raw]  = xlsread('N15InverseModelRW.GOM.xlsx','Sheet1','B6:B307');
    %     Columns = txt;
    %     save('Column Names.mat','Columns')
    load('Column Names.mat')
    meanvals = mean(MCMCmat)'./wts;
    stdvals = std(MCMCmat)'./wts;
    meantable = table(Columns,meanvals,stdvals);
    len=length(MCMCmat(:,1));

    for i=1:length(MCMCmat(:,1))

        MCMC=MCMCmat(i,:);
        d15N=del15N(i,:);

    if Cycle<=5
        [Aa] = ResetRN15(Aa0,MCMC,wts,d15N,d15NInputs(1,InputCol),d15NInputs(2,InputCol),d15NInputs(3,InputCol),d15NInputs(4,InputCol),d15NInputs(5,InputCol),d15NInputs(6,InputCol),d15NInputs(7,InputCol),d15NInputs(10,InputCol),d15NInputs(11,InputCol),d15NInputs(12,InputCol),d15NInputs(13,InputCol),d15NInputs(14,InputCol),d15NInputs(15,InputCol),d15NInputs(16,InputCol));
    else
        [Aa] = ResetRN15IO(Aa0,MCMC,wts,d15N,d15NInputs(1,InputCol),d15NInputs(2,InputCol),d15NInputs(3,InputCol),d15NInputs(4,InputCol),d15NInputs(5,InputCol),d15NInputs(6,InputCol),d15NInputs(7,InputCol),d15NInputs(10,InputCol),d15NInputs(11,InputCol),d15NInputs(12,InputCol),d15NInputs(13,InputCol),d15NInputs(14,InputCol),d15NInputs(15,InputCol),d15NInputs(16,InputCol));
    end

    ba_tmp = ba;
    sdba_tmp = sdba;
    % %The following code snippets gets rid of the constraint for
    % %phaeopigment flux since this was not actually included in the model
    % ind = find(sum(Aa')==0);
    % ind = find(ind>26);
    % Aa(ind,:) = [];
    % ba_tmp(ind) = [];
    % sdba_tmp(ind) = [];


        Measurement(:,i)=Aa*MCMC';
        Error(:,i)=Aa*MCMC'-ba_tmp;
        ErrorNorm(:,i)=Error(:,i)./sdba_tmp;
        SSR(:,i)=sum(ErrorNorm(:,i).^2);
        InequalError(:,i)=G*MCMC'-h;


    end


    if length(Aa(:,1))==72
        Labels1={'NPPs','NPPd','NO3ups','NO3upd','NH4ups','NH4upd','CyanoNPPs','CyanoNPPd','DtmNPPs','DtmNPPd',...
            'CyanoMorts','CyanoMortd','DtmMorts','DtmMortd','nvmMesoGr','vmMesoGr','STFluxs','STfluxd','Mic2Pre','Mic2Post',...
            'App2Pre','App2Post','Clad2Pre','Clad2Post','Cal2Pre','Cal2Post','Poecil2Pre','Poecil2Post'};
        Labels2={'NO3','NH4','Cyano','Tricho','Dtm','Flag','HNF','Mic','HerbNVM','App','Clad','nvmCal','Chaeto','Poecil','GelPred','PlankFish','Preflex','Postflex','Bac','Det','LDet','DON',...
            'HerbVM','vmCal',...
            'NO3','NH4','Cyano','Tricho','Dtm','Flag','HNF','Mic','HerbNVM','App','Clad','nvmCal','Chaeto','Poecil','GelPred','PlankFish','Bac','Det','LDet','DON'};
        Labels3={'Exc_HNF','Exc_Mic','Exc_Herb','Exc_App','Exc_Clad','Exc_Cal','Exc_Chaeto','Exc_Poecil','Exc_Gel','Exc_Plankfish','Exc_Preflex','Exc_Postflex',...
            'Exc_HerbVM','Exc_vmCal',...
            'Exc_HNF','Exc_Mic','Exc_Herb','Exc_App','Exc_Clad','Exc_Cal','Exc_Chaeto','Exc_Poecil','Exc_Gel','Exc_Plankfish',...
            'Resp_Herb','Resp_App','Resp_Clad','Resp_Cal','Resp_Chaeto','Resp_Poecil','Resp_HerbVM','Resp_vmCal',...
            'Resp_upper/lower','Resp_upper/lower'};
    elseif length(Aa(:,1))==80
        Labels1={'NPPs','NPPd','NO3ups','NO3upd','NH4ups','NH4upd','CyanoNPPs','CyanoNPPd','FlagNPPs','FlagNPPd',...
            'DtmNPPs','DtmNPPd','CyanoMorts','CyanoMortd','FlagMorts','FlagMortd','DtmMorts','DtmMortd','nvmMesoGr','vmMesoGr',...
            'STFluxs','STfluxd','ChlSinks','ChlSinkd','PhaeoSinks','PhaeoSinkd','Mic2Pre','Mic2Post','App2Pre','App2Post',...
            'Clad2Pre','Clad2Post','Cal2Pre','Cal2Post','Poecil2Pre','Poecil2Post'};
        Labels2={'NO3','NH4','Cyano','Tricho','Dtm','Flag','HNF','Mic','HerbNVM','App','Clad','nvmCal','Chaeto','Poecil','GelPred','PlankFish','Preflex','Postflex','Bac','Det','LDet','DON',...
            'HerbVM','vmCal',...
            'NO3','NH4','Cyano','Tricho','Dtm','Flag','HNF','Mic','HerbNVM','App','Clad','nvmCal','Chaeto','Poecil','GelPred','PlankFish','Bac','Det','LDet','DON'};
        Labels3={'Exc_HNF','Exc_Mic','Exc_Herb','Exc_App','Exc_Clad','Exc_Cal','Exc_Chaeto','Exc_Poecil','Exc_Gel','Exc_Plankfish','Exc_Preflex','Exc_Postflex',...
            'Exc_HerbVM','Exc_vmCal',...
            'Exc_HNF','Exc_Mic','Exc_Herb','Exc_App','Exc_Clad','Exc_Cal','Exc_Chaeto','Exc_Poecil','Exc_Gel','Exc_Plankfish',...
            'Resp_Herb','Resp_App','Resp_Clad','Resp_Cal','Resp_Chaeto','Resp_Poecil','Resp_HerbVM','Resp_vmCal',...
            'Resp_upper/lower','Resp_upper/lower'};
    elseif length(Aa(:,1))==82
        Labels1={'NPPs','NPPd','f-ratio s','f-ratio d','ProtistGrazings','ProtistGrazingd','CyanoNPPs','CyanoNPPd','FlagNPPs','FlagNPPd',...
            'DtmNPPs','DtmNPPd','HBacProds','HBacProdd','CyanoMorts','CyanoMortd','FlagMorts','FlagMortd','DtmMorts','DtmMortd','HBacGrazs','HBacGrazd','nvmMesoGr','vmMesoGr',...
            'STFluxs','STfluxd','ChlSinks','ChlSinkd','Mic2Pre','Mic2Post','App2Pre','App2Post',...
            'Clad2Pre','Clad2Post','Cal2Pre','Cal2Post','Poecil2Pre','Poecil2Post'};
        Labels2={'NO3','NH4','Cyano','Tricho','Dtm','Flag','HNF','Mic','HerbNVM','App','Clad','nvmCal','Chaeto','Poecil','GelPred','PlankFish','Preflex','Postflex','Bac','Det','LDet','DON',...
            'HerbVM','vmCal',...
            'NO3','NH4','Cyano','Tricho','Dtm','Flag','HNF','Mic','HerbNVM','App','Clad','nvmCal','Chaeto','Poecil','GelPred','PlankFish','Bac','Det','LDet','DON'};
        Labels3={'Exc_HNF','Exc_Mic','Exc_Herb','Exc_App','Exc_Clad','Exc_Cal','Exc_Chaeto','Exc_Poecil','Exc_Gel','Exc_Plankfish','Exc_Preflex','Exc_Postflex',...
            'Exc_HerbVM','Exc_vmCal',...
            'Exc_HNF','Exc_Mic','Exc_Herb','Exc_App','Exc_Clad','Exc_Cal','Exc_Chaeto','Exc_Poecil','Exc_Gel','Exc_Plankfish',...
            'Resp_Herb','Resp_App','Resp_Clad','Resp_Cal','Resp_Chaeto','Resp_Poecil','Resp_HerbVM','Resp_vmCal',...
            'Resp_upper/lower','Resp_upper/lower'};
    elseif length(Aa(:,1))==84 & Cycle==0
        Labels1={'NPPs','NPPd','f-ratio s','f-ratio d','ProtistGrazings','ProtistGrazingd','CyanoNPPs','CyanoNPPd','FlagNPPs','FlagNPPd',...
            'DtmNPPs','DtmNPPd','HBacProds','HBacProdd','CyanoMorts','CyanoMortd','FlagMorts','FlagMortd','DtmMorts','DtmMortd','HBacGrazs','HBacGrazd','nvmMesoGr','vmMesoGr',...
            'STFluxs','STfluxd','ChlSinks','ChlSinkd','PhaeoSinks','PhaeoSinkd','Mic2Pre','Mic2Post','App2Pre','App2Post',...
            'Clad2Pre','Clad2Post','Cal2Pre','Cal2Post','Poecil2Pre','Poecil2Post'};
        Labels2={'NO3','NH4','Cyano','Tricho','Dtm','Flag','HNF','Mic','HerbNVM','App','Clad','nvmCal','Chaeto','Poecil','GelPred','PlankFish','Preflex','Postflex','Bac','Det','LDet','DON',...
            'HerbVM','vmCal',...
            'NO3','NH4','Cyano','Tricho','Dtm','Flag','HNF','Mic','HerbNVM','App','Clad','nvmCal','Chaeto','Poecil','GelPred','PlankFish','Bac','Det','LDet','DON'};
        Labels3={'Exc_HNF','Exc_Mic','Exc_Herb','Exc_App','Exc_Clad','Exc_Cal','Exc_Chaeto','Exc_Poecil','Exc_Gel','Exc_Plankfish','Exc_Preflex','Exc_Postflex',...
            'Exc_HerbVM','Exc_vmCal',...
            'Exc_HNF','Exc_Mic','Exc_Herb','Exc_App','Exc_Clad','Exc_Cal','Exc_Chaeto','Exc_Poecil','Exc_Gel','Exc_Plankfish',...
            'Resp_Herb','Resp_App','Resp_Clad','Resp_Cal','Resp_Chaeto','Resp_Poecil','Resp_HerbVM','Resp_vmCal',...
            'Resp_upper/lower','Resp_upper/lower'};
    elseif length(Aa(:,1))==84 & Cycle==10
        Labels1={'NPPs','NPPd','f-ratio s','f-ratio d','NFix s','NFix d','ProtistGrazings','ProtistGrazingd','CyanoNPPs','CyanoNPPd','FlagNPPs','FlagNPPd',...
            'DtmNPPs','DtmNPPd','HBacProds','HBacProdd','CyanoMorts','CyanoMortd','FlagMorts','FlagMortd','DtmMorts','DtmMortd','HBacGrazs','HBacGrazd','nvmMesoGr','vmMesoGr',...
            'STFluxs','STfluxd','ChlSinks','ChlSinkd','Mic2Pre','Mic2Post','App2Pre','App2Post',...
            'Clad2Pre','Clad2Post','Cal2Pre','Cal2Post','Poecil2Pre','Poecil2Post'};
        Labels2={'NO3','NH4','Cyano','Tricho','Dtm','Flag','HNF','Mic','HerbNVM','App','Clad','nvmCal','Chaeto','Poecil','GelPred','PlankFish','Preflex','Postflex','Bac','Det','LDet','DON',...
            'HerbVM','vmCal',...
            'NO3','NH4','Cyano','Tricho','Dtm','Flag','HNF','Mic','HerbNVM','App','Clad','nvmCal','Chaeto','Poecil','GelPred','PlankFish','Bac','Det','LDet','DON'};
        Labels3={'Exc_HNF','Exc_Mic','Exc_Herb','Exc_App','Exc_Clad','Exc_Cal','Exc_Chaeto','Exc_Poecil','Exc_Gel','Exc_Plankfish','Exc_Preflex','Exc_Postflex',...
            'Exc_HerbVM','Exc_vmCal',...
            'Exc_HNF','Exc_Mic','Exc_Herb','Exc_App','Exc_Clad','Exc_Cal','Exc_Chaeto','Exc_Poecil','Exc_Gel','Exc_Plankfish',...
            'Resp_Herb','Resp_App','Resp_Clad','Resp_Cal','Resp_Chaeto','Resp_Poecil','Resp_HerbVM','Resp_vmCal',...
            'Resp_upper/lower','Resp_upper/lower'};
    elseif length(Aa(:,1))==86
        Labels1={'NPPs','NPPd','f-ratio s','f-ratio d','NFix s','NFix d','ProtistGrazings','ProtistGrazingd','CyanoNPPs','CyanoNPPd','FlagNPPs','FlagNPPd',...
            'DtmNPPs','DtmNPPd','HBacProds','HBacProdd','CyanoMorts','CyanoMortd','FlagMorts','FlagMortd','DtmMorts','DtmMortd','HBacGrazs','HBacGrazd','nvmMesoGr','vmMesoGr',...
            'STFluxs','STfluxd','ChlSinks','ChlSinkd','PhaeoSinks','PhaeoSinkd','Mic2Pre','Mic2Post','App2Pre','App2Post',...
            'Clad2Pre','Clad2Post','Cal2Pre','Cal2Post','Poecil2Pre','Poecil2Post'};
        Labels2={'NO3','NH4','Cyano','Tricho','Dtm','Flag','HNF','Mic','HerbNVM','App','Clad','nvmCal','Chaeto','Poecil','GelPred','PlankFish','Preflex','Postflex','Bac','Det','LDet','DON',...
            'HerbVM','vmCal',...
            'NO3','NH4','Cyano','Tricho','Dtm','Flag','HNF','Mic','HerbNVM','App','Clad','nvmCal','Chaeto','Poecil','GelPred','PlankFish','Bac','Det','LDet','DON'};
        Labels3={'Exc_HNF','Exc_Mic','Exc_Herb','Exc_App','Exc_Clad','Exc_Cal','Exc_Chaeto','Exc_Poecil','Exc_Gel','Exc_Plankfish','Exc_Preflex','Exc_Postflex',...
            'Exc_HerbVM','Exc_vmCal',...
            'Exc_HNF','Exc_Mic','Exc_Herb','Exc_App','Exc_Clad','Exc_Cal','Exc_Chaeto','Exc_Poecil','Exc_Gel','Exc_Plankfish',...
            'Resp_Herb','Resp_App','Resp_Clad','Resp_Cal','Resp_Chaeto','Resp_Poecil','Resp_HerbVM','Resp_vmCal',...
            'Resp_upper/lower','Resp_upper/lower'};
    end


    if Cycle==0
        
        fighandle = figure(1);
        fighandle.Units = 'inches';
        fighandle.Position = [1 1 3.5 6];
    end
    for j=1:2
        subplot(2,1,j)
        hold on
        plot([-12 1],[-12 1],'-k')
        for i=1:length(Labels1)
            temp=sort(Measurement(i,:));
            temp=[mean(temp),temp(round(len*0.025)),temp(round(len*0.975))];
            plot([log10(ba(i)),log10(ba(i))],[log10(temp(2)),log10(temp(3))],'-k');
            if ba(i)-sdba(i)>0
                plot([log10(ba(i)-sdba(i)*1.95),log10(ba(i)+sdba(i)*1.95)],[log10(temp(1)),log10(temp(1))],'-k');
            else
                plot([-11,log10(ba(i)+sdba(i)*1.95)],[log10(temp(1)),log10(temp(1))],'-k');
            end
            if Cycle==0
                plot(log10(ba(i)),log10(temp(1)),'dk','MarkerFaceColor','b','MarkerSize',8);
            else
                plot(log10(ba(i)),log10(temp(1)),'ok','MarkerFaceColor','r','MarkerSize',6);
            end
        end
        set(gca,'box','on')
        xlabel('Measurement (mmol N m^-^2 d^-^1)')
        ylabel('Model Estimate (mmol N m^-^2 d^-^1)')
        set(gca,'XTick',log10([10^-12:10^-12:10^-11,2*10^-11:10^-11:10^-10,2*10^-10:10^-10:10^-9,2*10^-9:10^-9:10^-8,2*10^-8:10^-8:10^-7,2*10^-7:10^-7:10^-6,2*10^-6:10^-6:10^-5,2*10^-5:10^-5:10^-4,2*10^-4:10^-4:10^-3, ...
            2*10^-3:10^-3:10^-2,2*10^-2:10^-2:10^-1,2*10^-1:10^-1:10^0,2*10^0:10^0:10^1]))
        set(gca,'XTickLabel',{'10^-^1^2','','','','','','','','','10^-^1^1','','','','','','','','','10^-^1^0','','','','','','','','','10^-^9','','','','','','','','','10^-^8','','','','','','','','','10^-^7','','','','','','','','','10^-^6','','','','','','','','','10^-^5',...
            '','','','','','','','','10^-^4','','','','','','','','','10^-^3','','','','','','','','','10^-^2',...
            '','','','','','','','','10^-^1','','','','','','','','','1','','','','','','','','','10'})
        set(gca,'YTick',log10([10^-12:10^-12:10^-11,2*10^-11:10^-11:10^-10,2*10^-10:10^-10:10^-9,2*10^-9:10^-9:10^-8,2*10^-8:10^-8:10^-7,2*10^-7:10^-7:10^-6,2*10^-6:10^-6:10^-5,2*10^-5:10^-5:10^-4,2*10^-4:10^-4:10^-3, ...
            2*10^-3:10^-3:10^-2,2*10^-2:10^-2:10^-1,2*10^-1:10^-1:10^0,2*10^0:10^0:10^1]))
        set(gca,'YTickLabel',{'10^-^1^2','','','','','','','','','10^-^1^1','','','','','','','','','10^-^1^0','','','','','','','','','10^-^9','','','','','','','','','10^-^8','','','','','','','','','10^-^7','','','','','','','','','10^-^6','','','','','','','','','10^-^5',...
            '','','','','','','','','10^-^4','','','','','','','','','10^-^3','','','','','','','','','10^-^2',...
            '','','','','','','','','10^-^1','','','','','','','','','1','','','','','','','','','10'})
    end
    subplot(2,1,1)
    xlim([-4 1])
    ylim([-4 1])
    subplot(2,1,2)
    xlim([-10.8, -4])
    ylim([-10.8, -4])


    %The following code snippets gets rid of the constraint for
    %phaeopigment flux since this was not actually included in the model
    ind = find(sum(Aa')==0);
    ind = ind(find(ind>26));
    Measurement(ind,:) = [];
    ErrorNorm(ind,:) = [];
    Labels1(ind) = [];
    ba(ind) = [];
    sdba(ind) = [];

sqrtmeanErrorNorm2 = sqrt(mean(ErrorNorm'.^2));
    ['The square root mean squared error, which can be thought of ',char(10),...
        'as the average number of standard errors that model estimates',char(10),...
        'were from the measurements was ',num2str(mean(sqrtmeanErrorNorm2(1:length(Labels1)))), ...
        ' if we consider ',char(10),' only the field rate measurements']
    
    ['It was ',num2str(mean(sqrtmeanErrorNorm2)),' for all measurements (including ',...
        'd^1^5N mass balance equations)']


    fn = 'ManuscriptValues.txt'
    fileID = fopen(fn,'a');
    if Cycle==0
        formatSpec =  '\n PlotMeasurements2Cycles.m \n ';
        fprintf(fileID,formatSpec)
    end
    formatSpec =  'The square root mean squared error was %f for Cycle %f if we consider only the field rate measurements \n';
    fprintf(fileID,formatSpec,[mean(sqrt(mean(ErrorNorm(1:length(Labels1),:)'.^2))), Cycle])
    formatSpec = 'It was %f for all measurements (including d^1^5N mass balance equations) for Cycle %f \n';
    fprintf(fileID,formatSpec,[mean(sqrt(mean(ErrorNorm'.^2))) Cycle])
    fclose(fileID)

    ModelEstimate = mean(Measurement(1:length(Labels1),:)')';
    ModelEstimate_sd = std(Measurement(1:length(Labels1),:)')';
    Observation = ba(1:length(Labels1));
    Observation_sd = sdba(1:length(Labels1));
    NormalizedError = sqrt(mean(ErrorNorm(1:length(Labels1),:)'.^2))';
    Labels = Labels1';
    if Cycle==0
        RateTable0 = table(Labels,ModelEstimate,ModelEstimate_sd,Observation,Observation_sd,NormalizedError);
        MeanTable = meantable;
        KEEP.MesoGraz0 = [mean(Measurement(23,:)+Measurement(24,:)),std(Measurement(23,:)+Measurement(24,:))]
    elseif Cycle==10
        RateTable10 = table(Labels,ModelEstimate,ModelEstimate_sd,Observation,Observation_sd,NormalizedError);
        MeanTable.IO_mean = meantable.meanvals;
        MeanTable.IO_std = meantable.stdvals;
        KEEP.MesoGraz10 = [mean(Measurement(25,:)+Measurement(26,:)),std(Measurement(25,:)+Measurement(26,:))]
    end

    

end
subplot(2,1,1)
plot(log10([1.2*10^-4,4*10^-2,4*10^-2,1.2*10^-4,1.2*10^-4]),log10([0.7 0.7 8 8 0.7]),'-k')
plot(log10([1.9*10^-4,1.9*10^-4]),log10([2.5 6]),'-k')
plot(log10([1.9*10^-4,1.9*10^-4]),log10([0.9 2]),'-k')
plot(log10([1.5*10^-4,2.4*10^-4]),log10([4 4]),'-k')
plot(log10([1.5*10^-4,2.4*10^-4]),log10([1.4 1.4]),'-k')
plot(log10(1.9*10^-4),log10(4),'dk','MarkerFaceColor','b','MarkerSize',8)
plot(log10(1.9*10^-4),log10(1.4),'ok','MarkerFaceColor','r','MarkerSize',6)
text(log10(2.8*10^-4),log10(4),'Gulf of Mexico','FontSize',9)
text(log10(2.8*10^-4),log10(1.4),'Argo Basin','FontSize',9)
set(gca,'FontSize',9)
AddPanelLabels('a','bottomright','YPADDING',0.04,'XPADDING',0.04,'FONTSIZE',9)
subplot(2,1,2)
set(gca,'FontSize',9)
AddPanelLabels('b','bottomright','YPADDING',0.04,'XPADDING',0.04,'FONTSIZE',9)


fn = ['PlotMeasurements2Cycles'];
exportgraphics(gcf,['../../Figures/',fn,'.png'])


    fn = 'ManuscriptValues.txt'
    fileID = fopen(fn,'a');
    formatSpec =  'The greatest square root mean squared errors were %f and %f for NPPs and NPPd on AAP, respectively \n';
    fprintf(fileID,formatSpec,[RateTable10.NormalizedError(1),RateTable10.NormalizedError(2) ])
    formatSpec = 'For shallow NPP the model estimate was %f +/- %f and the observed NPP was %f +/- %f \n';
    fprintf(fileID,formatSpec,[RateTable10.ModelEstimate(1),RateTable10.ModelEstimate_sd(1),RateTable10.Observation(1),RateTable10.Observation_sd(1)])
    formatSpec = 'For deep NPP the model estimate was %f +/- %f and the observed NPP was %f +/- %f \n';
    fprintf(fileID,formatSpec,[RateTable10.ModelEstimate(2),RateTable10.ModelEstimate_sd(2),RateTable10.Observation(2),RateTable10.Observation_sd(2)])
    x = (KEEP.MesoGraz0(1) - (RateTable0.Observation(23)+RateTable0.Observation(24)))/(RateTable0.Observation(23)+RateTable0.Observation(24))*100;
    y = (KEEP.MesoGraz10(1) - (RateTable10.Observation(25)+RateTable10.Observation(26)))/(RateTable10.Observation(25)+RateTable10.Observation(26))*100;
    formatSpec = 'For GoM the model overestimated mesozoo grazing by %f percent \n and for AAP it overestimated it by %f percent \n';
    fprintf(fileID,formatSpec,[x,y])
    x = RateTable10.ModelEstimate(9)/(RateTable10.ModelEstimate(11)+RateTable10.ModelEstimate(13));
    y = RateTable10.Observation(9)/(RateTable10.Observation(11)+RateTable10.Observation(13));
    formatSpec = 'For AAP the model estimate of the ratio of cyanobacteria NPP to eukaryotic phytoplankton NPP was %f \n while observations suggested that the ratio should be %f \n';
    fprintf(fileID,formatSpec,[x,y])


    fclose(fileID)


% 
% 
% breakitty
% 
% 
% 
% 
% 
% for Cycle=[0,10]
%     clearvars -except Cycle burnin
% 
%     load(['N15GoMInverseCycle',num2str(Cycle),'.mat'])
%     load(['N15GoMInverseCycle',num2str(Cycle),'Routputs.mat'])
%     num=length(MCMCmat(:,1));
%     MCMCmat(1:round(num*burnin),:)=[];
%     del15N(1:round(num*0.2),:)=[];
% %     [num,txt,raw]  = xlsread('N15InverseModelRW.GOM.xlsx','Sheet1','B6:B307');
% %     Columns = txt;
% %     save('Column Names.mat','Columns')
%     load('Column Names.mat')
% meanvals = mean(MCMCmat)'./wts;
% stdvals = std(MCMCmat)'./wts;
% meantable = table(Columns,meanvals,stdvals);
%     len=length(MCMCmat(:,1));
% 
%     for i=1:length(MCMCmat(:,1))
% 
%         MCMC=MCMCmat(i,:);
%         d15N=del15N(i,:);
% 
%         [Aa] = ResetRN15(Aa0,MCMC,wts,d15N,d15NInputs(1,InputCol),d15NInputs(2,InputCol),d15NInputs(3,InputCol),d15NInputs(4,InputCol),d15NInputs(5,InputCol),d15NInputs(6,InputCol),d15NInputs(7,InputCol));
%         Measurement(:,i)=Aa*MCMC';
%         Error(:,i)=Aa*MCMC'-ba;
%         ErrorNorm(:,i)=Error(:,i)./sdba;
%         SSR(:,i)=sum(ErrorNorm(:,i).^2);
%         InequalError(:,i)=G*MCMC'-h;
% 
% 
%     end
% 
% 
%     if length(Aa(:,1))==72
%         Labels1={'NPPs','NPPd','NO3ups','NO3upd','NH4ups','NH4upd','CyanoNPPs','CyanoNPPd','DtmNPPs','DtmNPPd',...
%             'CyanoMorts','CyanoMortd','DtmMorts','DtmMortd','nvmMesoGr','vmMesoGr','STFluxs','STfluxd','Mic2Pre','Mic2Post',...
%             'App2Pre','App2Post','Clad2Pre','Clad2Post','Cal2Pre','Cal2Post','Poecil2Pre','Poecil2Post'};
%         Labels2={'NO3','NH4','Cyano','Tricho','Dtm','Flag','HNF','Mic','HerbNVM','App','Clad','nvmCal','Chaeto','Poecil','GelPred','PlankFish','Preflex','Postflex','Bac','Det','LDet','DON',...
%             'HerbVM','vmCal',...
%             'NO3','NH4','Cyano','Tricho','Dtm','Flag','HNF','Mic','HerbNVM','App','Clad','nvmCal','Chaeto','Poecil','GelPred','PlankFish','Bac','Det','LDet','DON'};
%         Labels3={'Exc_HNF','Exc_Mic','Exc_Herb','Exc_App','Exc_Clad','Exc_Cal','Exc_Chaeto','Exc_Poecil','Exc_Gel','Exc_Plankfish','Exc_Preflex','Exc_Postflex',...
%             'Exc_HerbVM','Exc_vmCal',...
%             'Exc_HNF','Exc_Mic','Exc_Herb','Exc_App','Exc_Clad','Exc_Cal','Exc_Chaeto','Exc_Poecil','Exc_Gel','Exc_Plankfish',...
%             'Resp_Herb','Resp_App','Resp_Clad','Resp_Cal','Resp_Chaeto','Resp_Poecil','Resp_HerbVM','Resp_vmCal',...
%             'Resp_upper/lower','Resp_upper/lower'};
%     elseif length(Aa(:,1))==80
%         Labels1={'NPPs','NPPd','NO3ups','NO3upd','NH4ups','NH4upd','CyanoNPPs','CyanoNPPd','FlagNPPs','FlagNPPd',...
%             'DtmNPPs','DtmNPPd','CyanoMorts','CyanoMortd','FlagMorts','FlagMortd','DtmMorts','DtmMortd','nvmMesoGr','vmMesoGr',...
%             'STFluxs','STfluxd','ChlSinks','ChlSinkd','PhaeoSinks','PhaeoSinkd','Mic2Pre','Mic2Post','App2Pre','App2Post',...
%             'Clad2Pre','Clad2Post','Cal2Pre','Cal2Post','Poecil2Pre','Poecil2Post'};
%         Labels2={'NO3','NH4','Cyano','Tricho','Dtm','Flag','HNF','Mic','HerbNVM','App','Clad','nvmCal','Chaeto','Poecil','GelPred','PlankFish','Preflex','Postflex','Bac','Det','LDet','DON',...
%             'HerbVM','vmCal',...
%             'NO3','NH4','Cyano','Tricho','Dtm','Flag','HNF','Mic','HerbNVM','App','Clad','nvmCal','Chaeto','Poecil','GelPred','PlankFish','Bac','Det','LDet','DON'};
%         Labels3={'Exc_HNF','Exc_Mic','Exc_Herb','Exc_App','Exc_Clad','Exc_Cal','Exc_Chaeto','Exc_Poecil','Exc_Gel','Exc_Plankfish','Exc_Preflex','Exc_Postflex',...
%             'Exc_HerbVM','Exc_vmCal',...
%             'Exc_HNF','Exc_Mic','Exc_Herb','Exc_App','Exc_Clad','Exc_Cal','Exc_Chaeto','Exc_Poecil','Exc_Gel','Exc_Plankfish',...
%             'Resp_Herb','Resp_App','Resp_Clad','Resp_Cal','Resp_Chaeto','Resp_Poecil','Resp_HerbVM','Resp_vmCal',...
%             'Resp_upper/lower','Resp_upper/lower'};
%     elseif length(Aa(:,1))==84
%         Labels1={'NPPs','NPPd','f-ratio s','f-ratio d','NFix s','NFix d','CyanoNPPs','CyanoNPPd','FlagNPPs','FlagNPPd',...
%             'DtmNPPs','DtmNPPd','CyanoMorts','CyanoMortd','FlagMorts','FlagMortd','DtmMorts','DtmMortd','nvmMesoGr','vmMesoGr',...
%             'STFluxs','STfluxd','ChlSinks','ChlSinkd','PhaeoSinks','PhaeoSinkd','Mic2Pre','Mic2Post','App2Pre','App2Post',...
%             'Clad2Pre','Clad2Post','Cal2Pre','Cal2Post','Poecil2Pre','Poecil2Post'};
%         Labels2={'NO3','NH4','Cyano','Tricho','Dtm','Flag','HNF','Mic','HerbNVM','App','Clad','nvmCal','Chaeto','Poecil','GelPred','PlankFish','Preflex','Postflex','Bac','Det','LDet','DON',...
%             'HerbVM','vmCal',...
%             'NO3','NH4','Cyano','Tricho','Dtm','Flag','HNF','Mic','HerbNVM','App','Clad','nvmCal','Chaeto','Poecil','GelPred','PlankFish','Bac','Det','LDet','DON'};
%         Labels3={'Exc_HNF','Exc_Mic','Exc_Herb','Exc_App','Exc_Clad','Exc_Cal','Exc_Chaeto','Exc_Poecil','Exc_Gel','Exc_Plankfish','Exc_Preflex','Exc_Postflex',...
%             'Exc_HerbVM','Exc_vmCal',...
%             'Exc_HNF','Exc_Mic','Exc_Herb','Exc_App','Exc_Clad','Exc_Cal','Exc_Chaeto','Exc_Poecil','Exc_Gel','Exc_Plankfish',...
%             'Resp_Herb','Resp_App','Resp_Clad','Resp_Cal','Resp_Chaeto','Resp_Poecil','Resp_HerbVM','Resp_vmCal',...
%             'Resp_upper/lower','Resp_upper/lower'};
%     elseif length(Aa(:,1))==86
%         Labels1={'NPPs','NPPd','f-ratio s','f-ratio d','NFix s','NFix d','CyanoNPPs','CyanoNPPd','FlagNPPs','FlagNPPd',...
%             'DtmNPPs','DtmNPPd','CyanoMorts','CyanoMortd','FlagMorts','FlagMortd','DtmMorts','DtmMortd','nvmMesoGr','vmMesoGr',...
%             'STFluxs','STfluxd','ChlSinks','ChlSinkd','PhaeoSinks','PhaeoSinkd','Mic2Pre','Mic2Post','App2Pre','App2Post',...
%             'Clad2Pre','Clad2Post','Cal2Pre','Cal2Post','Poecil2Pre','Poecil2Post'};
%         Labels2={'NO3','NH4','Cyano','Tricho','Dtm','Flag','HNF','Mic','HerbNVM','App','Clad','nvmCal','Chaeto','Poecil','GelPred','PlankFish','Preflex','Postflex','Bac','Det','LDet','DON',...
%             'HerbVM','vmCal',...
%             'NO3','NH4','Cyano','Tricho','Dtm','Flag','HNF','Mic','HerbNVM','App','Clad','nvmCal','Chaeto','Poecil','GelPred','PlankFish','Bac','Det','LDet','DON'};
%         Labels3={'Exc_HNF','Exc_Mic','Exc_Herb','Exc_App','Exc_Clad','Exc_Cal','Exc_Chaeto','Exc_Poecil','Exc_Gel','Exc_Plankfish','Exc_Preflex','Exc_Postflex',...
%             'Exc_HerbVM','Exc_vmCal',...
%             'Exc_HNF','Exc_Mic','Exc_Herb','Exc_App','Exc_Clad','Exc_Cal','Exc_Chaeto','Exc_Poecil','Exc_Gel','Exc_Plankfish',...
%             'Resp_Herb','Resp_App','Resp_Clad','Resp_Cal','Resp_Chaeto','Resp_Poecil','Resp_HerbVM','Resp_vmCal',...
%             'Resp_upper/lower','Resp_upper/lower'};
%     end
% 
% 
%     if Cycle==1
%         figure('Position',[200 50 900 600])
%     end
%     for j=1:1
%         %subplot(2,1,j)
%         hold on
%         plot([-10 1],[-10 1],'-k')
%         for i=1:length(Labels1)
%             temp=sort(Measurement(i,:));
%             temp=[mean(temp),temp(round(len*0.025)),temp(round(len*0.975))];
%             plot([log10(ba(i)),log10(ba(i))],[log10(temp(2)),log10(temp(3))],'-k');
%             if ba(i)-sdba(i)>0
%                 plot([log10(ba(i)-sdba(i)*1.95),log10(ba(i)+sdba(i)*1.95)],[log10(temp(1)),log10(temp(1))],'-k');
%             else
%                 plot([-9,log10(ba(i)+sdba(i)*1.95)],[log10(temp(1)),log10(temp(1))],'-k');
%             end
%             if Cycle==1
%                 plot(log10(ba(i)),log10(temp(1)),'dk','MarkerFaceColor','b','MarkerSize',8);
%             else
%                 plot(log10(ba(i)),log10(temp(1)),'ok','MarkerFaceColor','r','MarkerSize',6);
%             end
%             text(log10(ba(i)),log10(temp(1)),num2str(i))
%         end
%         set(gca,'box','on')
%         xlabel('Measurement (mmol N m^-^2 d^-^1)')
%         ylabel('Model Estimate (mmol N m^-^2 d^-^1)')
%         set(gca,'XTick',log10([10^-9:10^-9:10^-8,2*10^-8:10^-8:10^-7,2*10^-7:10^-7:10^-6,2*10^-6:10^-6:10^-5,2*10^-5:10^-5:10^-4,2*10^-4:10^-4:10^-3, ...
%             2*10^-3:10^-3:10^-2,2*10^-2:10^-2:10^-1,2*10^-1:10^-1:10^0,2*10^0:10^0:10^1]))
%         set(gca,'XTickLabel',{'10^-^9','','','','','','','','','10^-^8','','','','','','','','','10^-^7','','','','','','','','','10^-^6','','','','','','','','','10^-^5',...
%             '','','','','','','','','10^-^4','','','','','','','','','10^-^3','','','','','','','','','10^-^2',...
%             '','','','','','','','','10^-^1','','','','','','','','','1','','','','','','','','','10'})
%         set(gca,'YTick',log10([10^-9:10^-9:10^-8,2*10^-8:10^-8:10^-7,2*10^-7:10^-7:10^-6,2*10^-6:10^-6:10^-5,2*10^-5:10^-5:10^-4,2*10^-4:10^-4:10^-3, ...
%             2*10^-3:10^-3:10^-2,2*10^-2:10^-2:10^-1,2*10^-1:10^-1:10^0,2*10^0:10^0:10^1]))
%         set(gca,'YTickLabel',{'10^-^9','','','','','','','','','10^-^8','','','','','','','','','10^-^7','','','','','','','','','10^-^6','','','','','','','','','10^-^5',...
%             '','','','','','','','','10^-^4','','','','','','','','','10^-^3','','','','','','','','','10^-^2',...
%             '','','','','','','','','10^-^1','','','','','','','','','1','','','','','','','','','10'})
%     end
%     xlim([-4 1])
%     ylim([-4 1])
% 
% 
% end
% plot(log10([1.2*10^-4,2*10^-3,2*10^-3,1.2*10^-4,1.2*10^-4]),log10([0.7 0.7 8 8 0.7]),'-k')
% plot(log10([1.9*10^-4,1.9*10^-4]),log10([2.5 6]),'-k')
% plot(log10([1.9*10^-4,1.9*10^-4]),log10([0.9 2]),'-k')
% plot(log10([1.5*10^-4,2.4*10^-4]),log10([4 4]),'-k')
% plot(log10([1.5*10^-4,2.4*10^-4]),log10([1.4 1.4]),'-k')
% plot(log10(1.9*10^-4),log10(4),'dk','MarkerFaceColor','b','MarkerSize',8)
% plot(log10(1.9*10^-4),log10(1.4),'ok','MarkerFaceColor','r','MarkerSize',6)
% text(log10(2.8*10^-4),log10(4),'Cycle 1','FontSize',14)
% text(log10(2.8*10^-4),log10(1.4),'Cycle 5','FontSize',14)
% set(gca,'FontSize',14)
```
