## Supplementary material for "Comparative food-web analysis of bluefin tuna spawning habitats in the eastern Indian Ocean and Gulf of Mexico": Code Package: PlotTL.m

```
clear all
close all

burnin = 0.2;
cols = [1 0.8 0.3; ...
        0.4 0.9 0.3; ...
        0 0.5 0.4];

for Cycle=[0,10]
    
    %--------------------------------------------------
    %-----------Data loading and processing------------
    %--------------------------------------------------
    
    clearvars -except cols burnin Cycle
    load(['N15GoMInverseCycle',num2str(Cycle),'.mat'])
    load(['N15GoMInverseCycle',num2str(Cycle),'Routputs.mat'])
    
    num=length(MCMCmat(:,1));
    MCMCmat(1:round(num*burnin),:)=[];
    del15N(1:round(num*burnin),:)=[];
    len=length(MCMCmat(:,1));
    
    
    [NO3s_col,NH4s_col,Cyas_col,Tris_col,Dtms_col,Flags_col,HNFs_col,MICs_col,HerbNVMs_col,Apps_col,Clads_col,nvmCals_col,Chaetos_col,Poecils_col,GelPreds_col,Plankfishs_col,Preflexs_col,Postflexs_col,Bacs_col,Dets_col,Ldets_col,Dons_col,HerbVM_col,vmCal_col,NO3d_col,NH4d_col,Cyad_col,Trid_col,Dtmd_col,Flagd_col,HNFd_col,MICd_col,HerbNVMd_col,Appd_col,Cladd_col,nvmCald_col,Chaetod_col,Poecild_col,GelPredd_col,Plankfishd_col,Bacd_col,Detd_col,Ldetd_col,Dond_col] = GetColumns(Aa0,MCMCmat,del15N(1,:),d15NInputs(1,InputCol),d15NInputs(2,InputCol),d15NInputs(3,InputCol),d15NInputs(4,InputCol),d15NInputs(5,InputCol),d15NInputs(6,InputCol),d15NInputs(7,InputCol));
    
    for i=1:length(MCMCmat(:,1))
        i
        flows=MCMCmat(i,:)./wts';
        TL0 = CalcTL(flows,del15N,d15NInputs,Ae0,InputCol);
        TL_track(:,i)=TL0;
    end
    
    Columns = [HNFs_col,MICs_col,Apps_col,HerbNVMs_col,Clads_col,nvmCals_col,HerbVM_col,vmCal_col,Chaetos_col,Poecils_col,GelPreds_col,Preflexs_col,Postflexs_col,Plankfishs_col];
    Columnsd = [HNFd_col,MICd_col,Appd_col,HerbNVMd_col,Cladd_col,nvmCald_col,HerbVM_col,vmCal_col,Chaetod_col,Poecild_col,GelPredd_col,Plankfishd_col];
    
    %-------------------------------------------------
    %-------------Main TL Plot------------------------
    %--------------------------------------------------
    
    %Mixed layer
    if Cycle==0
        fighandle = figure(1);
        fighandle.Units = 'inches';
        fighandle.Position = [1 1 6.5 6.5];
        subplot(2,2,1)
    else
        figure(1)
        subplot(2,2,3)
    end
    ymax=5.5;
    ymin=1.8;
    %boxplot(TL_track(Columns,:)','Symbol','')
    for i=1:length(Columns)
        if i==2 | i==3 | i==5 | i==6 | i==10
            col = cols(2,:);
        elseif i==12 | i==13
            col = cols(3,:);
        else
            col = cols(1,:);
        end
        ViolinPlot(i,0.5,TL_track(Columns(i),:),200,col)
    end
    set(gca,'XTick',1:length(Columns))
    set(gca,'XTickLabel',{'Nanoflag','Microzoo','Appendicularians','Herb (NVM)','Cladocerans','Calanoids (NVM)','Herb (VM)','Calanoids (VM)','Chaetognaths','Poecilostomatoids','Gel Predators','Preflex ABT','Postflex ABT','Planktivores'})
    set(gca,'XTickLabelRotation',45)
    ylim([ymin ymax])
    set(gca,'FontSize',9)
    ylabel('Trophic Level')
    if Cycle==0
        title(['Upper Euphotic Zone',char(10),'Gulf of Mexico'])
        AddPanelLabels('a','bottomright')
    else
        title(['Upper Euphotic Zone',char(10),'Argo Basin'])
        AddPanelLabels('c','bottomright')
    end
    xlim([0.5, length(Columns)+0.5])
    
    %Lower Euphotic Zone
    if Cycle==0
        subplot(2,2,2)
    else
        subplot(2,2,4)
    end
    ymax=5.5;
    ymin=1.8;
    %boxplot(TL_track(Columnsd,:)','Symbol','')
    for i=1:length(Columnsd)
        if i==2 | i==3 | i==5 | i==6 | i==10
            col = cols(2,:);
        elseif i>14
            col = cols(3,:);
        else
            col = cols(1,:);
        end
        ViolinPlot(i,0.5,TL_track(Columnsd(i),:),200,col)
    end
    set(gca,'XTick',1:length(Columnsd))
    set(gca,'XTickLabel',{'Nanoflag','Microzoo','Appendicularians','Herb (NVM)','Cladocerans','Calanoids (NVM)','Herb (VM)','Calanoids (VM)','Chaetognaths','Poecilostomatoids','Gel Predators','Planktivores'})
    set(gca,'XTickLabelRotation',45)
    ylim([ymin ymax]);
    set(gca,'FontSize',9)
    ylabel('Trophic Level')
    title(['Deep Chlorophyll Max - Cycle ',num2str(Cycle)])
    xlim([0.5, length(Columnsd)+0.5]);
    if Cycle==0
        title(['Deep Chlorophyll Maximum',char(10),'Gulf of Mexico'])
        AddPanelLabels('b','bottomright')
    else
        title(['Deep Chlorophyll Maximum',char(10),'Argo Basin'])
        AddPanelLabels('d','bottomright')
    end
    


    %-------------------------------------------------
    %-------------Subsetted side-by-side TL figure------------------------
    %--------------------------------------------------
    
    %Mixed layer
    if Cycle==0
        fighandle = figure(202);
        fighandle.Units = 'inches';
        fighandle.Position = [1 1 3.5 3];
    else
        figure(202)
    end
    ymax=5;
    ymin=1.9;


    XLOCS = [-10, ... %nanoflags
             1, ... %microzoos
             2, ... %apps
             -10, ... %herb (nvm)
             3, ... %Cladocerans
             4, ... %Calanoids (nvm)
             -10, ... %herbivores (vm)
             -10, ... %calanoids (vm)
             -10, ... %chaetognaths
             5, ... %poecilostomatoids
             -10, ... %gel predators
             6, ... %preflex ABT
             7, ... %postflex ABT
             -10];    %planktivores


    for i=1:length(XLOCS)
       if Cycle==0
           col = [0 0 1];
       else
           col = [1 0 0];
       end
        ViolinPlot(XLOCS(i)+Cycle/50,0.2,TL_track(Columns(i),:),200,col)
    end
    xlim([0.5 7.5])

    h = fill([0.6 0.6 4.2 4.2 0.6],[ymax-0.05 ymax-0.65 ymax-0.65    ymax-0.05  ymax-0.05],'w')
    h=fill([1 1 0.75 0.75 1],[ymax-0.1 ymax-0.3 ymax-0.3    ymax-0.1  ymax-0.1],'w')
    set(h,'FaceColor','b')
    text(1.1,ymax-0.2,'Gulf of Mexico','FontSize',9)
    h=fill([1 1 0.75 0.75 1],[ymax-0.6 ymax-0.4 ymax-0.4    ymax-0.6  ymax-0.6],'w')
    set(h,'FaceColor','r')
    text(1.1,ymax-0.5,'Argo Basin','FontSize',9)

    set(gca,'XTick',[1:max(XLOCS)]+0.1)
    set(gca,'XTickLabel',{'Microzoo','Appendicularians','Cladocerans','Calanoids (NVM)','Poecilostomatoids','Preflex ABT','Postflex ABT'})
    set(gca,'XTickLabelRotation',30)
    ylim([ymin ymax]);
    set(gca,'FontSize',9)
    ylabel('Trophic Level')


    %-------------------------------------------------
    %-------Max food chain length plot----------------------------
    %-------------------------------------------------
    maxfoodchainlength = [3, 4, 4, 5, 5, 5, 5, 5, 6, 6, 7, 7, 7, 7];
    minfoodchainlength = [2, 2, 2, 2, 2, 2, 2, 2, 3, 3, 3, 3, 3, 3];
    
    for i=1:length(TL_track(:,1))
        temp = sort(TL_track(i,:));
        TL(i,:) = [mean(temp),temp(round(0.025*length(temp))),temp(round(0.975*length(temp)))];
    end
    
    col = [1 1 0;
        0 0 1;
        0 0 1;
        1 1 0;
        0 0 1;
        0 0 1;
        1 1 0;
        1 1 0;
        1 1 0;
        0 0 1;
        1 1 0;
        1 0 0;
        1 0 0;
        1 1 0];
    
    figure('Position',[100 50 600 600])
    for i=1:length(Columns)
        hold on
        ind = Columns(i);
        plot([TL(ind,1) TL(ind,1)],[TL(ind,2), TL(ind,3)]/maxfoodchainlength(i),'-k')
        plot([TL(ind,2) TL(ind,3)],[TL(ind,1), TL(ind,1)]/maxfoodchainlength(i),'-k')
        plot(TL(ind,1),TL(ind,1)/maxfoodchainlength(i),'dk','MarkerFaceColor',col(i,:),'MarkerSize',12);
    end
    fill([4, 4, 4.95, 4.95, 4],[0.41 0.51 0.51 0.41 0.41],'w')
    set(gca,'box','on')
    xlabel('Trophic Level')
    ylabel('Trophic Level / Maximum Trophic Level')
    plot(4.1,0.49,'dk','MarkerFaceColor',col(2,:),'MarkerSize',12)
    text(4.2,0.49,'ABT Prey','FontSize',16)
    plot(4.1,0.46,'dk','MarkerFaceColor',col(12,:),'MarkerSize',12)
    text(4.2,0.46,'Larval ABT','FontSize',16)
    plot(4.1,0.43,'dk','MarkerFaceColor',col(1,:),'MarkerSize',12)
    text(4.2,0.43,'Others','FontSize',16)
    set(gca,'FontSize',16)
    title(['Cycle ',num2str(Cycle)])
    
    
    figure('Position',[100 50 600 600])
    for i=1:length(Columns)
        hold on
        ind = Columns(i);
        if Cycle==1
            plot([TL(ind,1) TL(ind,1)],[TL(ind,2), TL(ind,3)]/maxfoodchainlength(i),'-k')
            plot([TL(ind,2) TL(ind,3)],[TL(ind,1), TL(ind,1)]/maxfoodchainlength(i),'-k')
            plot(TL(ind,1),TL(ind,1)/maxfoodchainlength(i),'dk','MarkerFaceColor',col(i,:),'MarkerSize',12);
        elseif Cycle==5
            plot([TL(ind,1) TL(ind,1)],[TL(ind,2), TL(ind,3)]/maxfoodchainlength(i),'-k')
            plot([TL(ind,2) TL(ind,3)],[TL(ind,1), TL(ind,1)]/maxfoodchainlength(i),'-k')
            plot(TL(ind,1),TL(ind,1)/maxfoodchainlength(i),'sk','MarkerFaceColor',col(i,:),'MarkerSize',10);
        end
    end
    fill([3.2, 3.2, 5.05, 5.05, 3.2],[0.41 0.51 0.51 0.41 0.41],'w')
    set(gca,'box','on')
    xlabel('Trophic Level')
    ylabel('Trophic Level / Maximum Trophic Level')
    plot(3.3,0.49,'ok','MarkerFaceColor',col(2,:),'MarkerSize',12)
    text(3.4,0.49,'ABT Prey','FontSize',16)
    plot(3.3,0.46,'ok','MarkerFaceColor',col(12,:),'MarkerSize',12)
    text(3.4,0.46,'Larval ABT','FontSize',16)
    plot(3.3,0.43,'ok','MarkerFaceColor',col(1,:),'MarkerSize',12)
    text(3.4,0.43,'Others','FontSize',16)
    plot(4.3,0.48,'dk','MarkerFaceColor',[0.5 0.5 0.5],'MarkerSize',12)
    text(4.4,0.48,'Cycle 1','FontSize',16)
    plot(4.3,0.44,'sk','MarkerFaceColor',[0.5 0.5 0.5],'MarkerSize',10)
    text(4.4,0.44,'Cycle 2','FontSize',16)
    set(gca,'FontSize',16)
    xlim([2 5.1])
    ylim([0.4 0.9])
    
    
    figure('Position',[500 50 600 600])
    for i=1:length(Columns)
        hold on
        ind = Columns(i);
        plot([TL(ind,1) TL(ind,1)],[TL(ind,2), TL(ind,3)]/minfoodchainlength(i),'-k')
        plot([TL(ind,2) TL(ind,3)],[TL(ind,1), TL(ind,1)]/minfoodchainlength(i),'-k')
        plot(TL(ind,1),TL(ind,1)/minfoodchainlength(i),'dk','MarkerFaceColor',col(i,:),'MarkerSize',12);
    end
    fill([4, 4, 4.95, 4.95, 4],[1.02, 1.22, 1.22, 1.02, 1.02],'w')
    set(gca,'box','on')
    xlabel('Trophic Level')
    ylabel('Trophic Level / Minimum Trophic Level')
    plot(4.1,1.18,'dk','MarkerFaceColor',col(2,:),'MarkerSize',12)
    text(4.2,1.18,'ABT Prey','FontSize',16)
    plot(4.1,1.12,'dk','MarkerFaceColor',col(12,:),'MarkerSize',12)
    text(4.2,1.12,'Larval ABT','FontSize',16)
    plot(4.1,1.06,'dk','MarkerFaceColor',col(1,:),'MarkerSize',12)
    text(4.2,1.06,'Others','FontSize',16)
    set(gca,'FontSize',16)
    title(['Cycle ',num2str(Cycle)])
    
    
    
    
    
    figure('Position',[500 50 600 600])
    for i=1:length(Columns)
        hold on
        ind = Columns(i);
        plot([TL(ind,1) TL(ind,1)],([TL(ind,2), TL(ind,3)]-minfoodchainlength(i))/(maxfoodchainlength(i)-minfoodchainlength(i)),'-k')
        plot([TL(ind,2) TL(ind,3)],([TL(ind,1), TL(ind,1)]-minfoodchainlength(i))/(maxfoodchainlength(i)-minfoodchainlength(i)),'-k')
        plot(TL(ind,1),(TL(ind,1)-minfoodchainlength(i))/(maxfoodchainlength(i)-minfoodchainlength(i)),'dk','MarkerFaceColor',col(i,:),'MarkerSize',12);
    end
    fill([4, 4, 4.95, 4.95, 4],[0.8 0.98 0.98 0.8 0.8],'w')
    set(gca,'box','on')
    xlabel('Trophic Level')
    ylabel('(TL - TL_m_i_n)/(TL_m_a_x - TL_m_i_n) ')
    plot(4.1,0.95,'dk','MarkerFaceColor',col(2,:),'MarkerSize',12)
    text(4.2,0.95,'ABT Prey','FontSize',16)
    plot(4.1,0.89,'dk','MarkerFaceColor',col(12,:),'MarkerSize',12)
    text(4.2,0.89,'Larval ABT','FontSize',16)
    plot(4.1,0.83,'dk','MarkerFaceColor',col(1,:),'MarkerSize',12)
    text(4.2,0.83,'Others','FontSize',16);
    set(gca,'FontSize',16);
    ylim([0 1]);
    title(['Cycle ',num2str(Cycle)]);
    
    
    
    
    
    
    
    
    
    
    
    TLpreflex = sort(TL_track(Preflexs_col,:));
    TLpostflex = sort(TL_track(Postflexs_col,:));
    ['Preflexion ABT had a TL of ',num2str(mean(TLpreflex),2),' (95% C.I. = ',num2str(TLpreflex(round(0.025*len)),2),' - ',...
        num2str(TLpreflex(round(0.975*len)),2),'), while',char(10),'postflexion ABT had a TL of ',num2str(mean(TLpostflex),2),...
        '(',num2str(TLpostflex(round(0.025*len)),2),' - ',num2str(TLpreflex(round(0.975*len)),2),')']


    fn = 'ManuscriptValues.txt'
    fileID = fopen(fn,'a');
    if Cycle==0
        formatSpec =  '\n PlotTL.m \n ';
        fprintf(fileID,formatSpec)
    end
    formatSpec =  'For Cycle %f Preflexion ABT had a TL of %f +/- %f and Postflexion ABT had a TL of %f +/- %f \n';
    fprintf(fileID,formatSpec,[Cycle, mean(TLpreflex), std(TLpreflex), mean(TLpostflex), std(TLpostflex)])
    fclose(fileID)
    
    TLpoecils = sort(TL_track(Poecils_col,:));
    TLchaetos = sort(TL_track(Chaetos_col,:));
    TLgelpreds = sort(TL_track(GelPreds_col,:));
    TLpoecild = sort(TL_track(Poecild_col,:));
    TLchaetod = sort(TL_track(Chaetod_col,:));
    TLgelpredd = sort(TL_track(GelPredd_col,:));
    ['Predatory zooplankton (poecilostomatoid copepods, chaetognaths, and gelatinous predators)',char(10),...
        ' had particularly high trophic positions of ',num2str(mean(TLpoecils),2),', ',num2str(mean(TLchaetos),2),', and ',num2str(mean(TLgelpreds),2),', respectively, in the upper euphotic zone.']
    ['Their trophic positions in the deep euphotic zone were ',char(10),...
        num2str(mean(TLpoecild),2),', ',num2str(mean(TLchaetod),2),', and ',num2str(mean(TLgelpredd),2),', respectively, in the deep euphotic zone.']
    
    
end

figure(1)
fn = ['PlotTL'];
exportgraphics(gcf,['../../Figures/',fn,'.png'])

figure(202)
fn = ['PlotTL.SideBySide'];
exportgraphics(gcf,['../../Figures/',fn,'.png'])
```
