## Supplementary material for "Comparative food-web analysis of bluefin tuna spawning habitats in the eastern Indian Ocean and Gulf of Mexico": Code Package: PlotTL_ICCAT.m

```
clear all
close all

Cycle=1

load(['N15GoMInverseCycle',num2str(Cycle),'.mat'])
load(['N15GoMInverseCycle',num2str(Cycle),'Routputs.mat'])

num=length(MCMCmat(:,1));
MCMCmat(1:round(num*0.2),:)=[];
del15N(1:round(num*0.2),:)=[];


[NO3s_col,NH4s_col,Cyas_col,Tris_col,Dtms_col,Flags_col,HNFs_col,MICs_col,HerbNVMs_col,Apps_col,Clads_col,nvmCals_col,Chaetos_col,Poecils_col,GelPreds_col,Plankfishs_col,Preflexs_col,Postflexs_col,Bacs_col,Dets_col,Ldets_col,Dons_col,HerbVM_col,vmCal_col,NO3d_col,NH4d_col,Cyad_col,Trid_col,Dtmd_col,Flagd_col,HNFd_col,MICd_col,HerbNVMd_col,Appd_col,Cladd_col,nvmCald_col,Chaetod_col,Poecild_col,GelPredd_col,Plankfishd_col,Bacd_col,Detd_col,Ldetd_col,Dond_col] = GetColumns(Aa0,MCMCmat,del15N(1,:),d15NInputs(1,InputCol),d15NInputs(2,InputCol),d15NInputs(3,InputCol),d15NInputs(4,InputCol),d15NInputs(5,InputCol),d15NInputs(6,InputCol),d15NInputs(7,InputCol));

for i=1:length(MCMCmat(:,1))
    i
    flows=MCMCmat(i,:)./wts';
    TL0 = CalcTL(flows,del15N,d15NInputs,Ae0,InputCol);
    TL_track(:,i)=TL0;
end

Columns = [HNFs_col,MICs_col,Apps_col,HerbNVMs_col,Clads_col,nvmCals_col,HerbVM_col,vmCal_col,Chaetos_col,Poecils_col,GelPreds_col,Preflexs_col,Postflexs_col,Plankfishs_col]
    


figure('Position',[100 50 1000 600])
ymax=5.5;
ymin=1.8;
boxplot(TL_track(Columns,:)')
set(gca,'XTick',1:length(Columns))
set(gca,'XTickLabel',{'Nanoflag','Microzoo','Appendicularians','Herb (NVM)','Cladocerans','Calanoids (NVM)','Herb (VM)','Calanoids (VM)','Chaetognaths','Poecilostomatoids','Gel Predators','Preflex ABT','Postflex ABT','Planktivorous fish'})
set(gca,'XTickLabelRotation',45)
ylim([ymin ymax])
set(gca,'FontSize',20)
ylabel('Trophic Level')

figure('Position',[100 50 1000 600])
ymax=5.5;
ymin=1.8;
Prey=[2,3,5,6,10];
hold on
for i=1:length(Prey)
    h=fill([Prey(i)-0.4,Prey(i)-0.4,Prey(i)+0.4,Prey(i)+0.4,Prey(i)-0.4],[ymin+0.05 ymax-0.05 ymax-0.05 ymin+0.05 ymin+0.05],'y','EdgeColor','none')
    set(h,'FaceColor',[1 1 0.5])
end
BFT=[12,13];
for i=1:length(BFT)
    h=fill([BFT(i)-0.4,BFT(i)-0.4,BFT(i)+0.4,BFT(i)+0.4,BFT(i)-0.4],[ymin+0.05 ymax-0.05 ymax-0.05 ymin+0.05 ymin+0.05],'c','EdgeColor','none')
    set(h,'FaceColor',[0.7 0.9 1])
end
h=boxplot(TL_track(Columns,:)','Symbol','')
set(h,'LineWidth',1.5)
set(gca,'XTick',1:length(Columns))
set(gca,'XTickLabel',{'Nanoflag','Microzoo','Appendicularians','Herb (NVM)','Cladocerans','Calanoids (NVM)','Herb (VM)','Calanoids (VM)','Chaetognaths','Poecilostomatoids','Gel Predators','Preflex ABT','Postflex ABT','Planktivorous fish'})
set(gca,'XTickLabelRotation',45)
ylim([ymin ymax])
set(gca,'FontSize',20)
ylabel('Trophic Level')

origin = 'GoM Nitrogen & Trophic - NOAA RESTORE\Inverse Model\R Code\Test 50\PlotTLICCAT.m'
save('TrophicLevel.mat','BFT','Prey','origin','TL_track','Columns')
```
