## Supplementary material for "Comparative food-web analysis of bluefin tuna spawning habitats in the eastern Indian Ocean and Gulf of Mexico": Code Package: PlotTunaDiet.m

```
clear all
close all

for Cycle=[0,10]

    load(['N15GoMInverseCycle',num2str(Cycle),'.mat'])
    load(['N15GoMInverseCycle',num2str(Cycle),'Routputs.mat'])
    clearvars -except Ae0 MCMCmat Aa d15NInputs del15N InputCol wts Cycle
    num=length(MCMCmat(:,1));
    MCMCmat(1:round(num*0.2),:)=[];
    del15N(1:round(num*0.2),:)=[];
    len=length(MCMCmat(:,1));
    [num,txt,raw]  = xlsread('N15InverseModelRW.GOM.xlsx','Sheet1','B6:B307');
    Columns = txt;
    meanvals = mean(MCMCmat)'./wts;
    stdvals = std(MCMCmat)'./wts;
    meantable = table(Columns,meanvals,stdvals);

    [NO3s_col,NH4s_col,Cyas_col,Tris_col,Dtms_col,Flags_col,HNFs_col,MICs_col,HerbNVMs_col,Apps_col,Clads_col,nvmCals_col,Chaetos_col,Poecils_col,GelPreds_col,Plankfishs_col,Preflexs_col,Postflexs_col,Bacs_col,Dets_col,Ldets_col,Dons_col,HerbVM_col,vmCal_col,NO3d_col,NH4d_col,Cyad_col,Trid_col,Dtmd_col,Flagd_col,HNFd_col,MICd_col,HerbNVMd_col,Appd_col,Cladd_col,nvmCald_col,Chaetod_col,Poecild_col,GelPredd_col,Plankfishd_col,Bacd_col,Detd_col,Ldetd_col,Dond_col] = GetColumns(Aa,MCMCmat,del15N(1,:),d15NInputs(1,InputCol),d15NInputs(2,InputCol),d15NInputs(3,InputCol),d15NInputs(4,InputCol),d15NInputs(5,InputCol),d15NInputs(6,InputCol),d15NInputs(7,InputCol));
    ToPreflex = find(Ae0(Preflexs_col,:)==1);
    ToPostflex = find(Ae0(Postflexs_col,:)==1);
    FromMIC = find(Ae0(MICs_col,:)==-1);
    FromApp = find(Ae0(Apps_col,:)==-1);
    FromClad = find(Ae0(Clads_col,:)==-1);
    FromnvmCal = find(Ae0(nvmCals_col,:)==-1);
    FromPoecil = find(Ae0(Poecils_col,:)==-1);

    MICtoPREFLEX = intersect(ToPreflex,FromMIC);
    MICtoPOSTFLEX = intersect(ToPostflex,FromMIC);
    ApptoPREFLEX = intersect(ToPreflex,FromApp);
    ApptoPOSTFLEX = intersect(ToPostflex,FromApp);
    CladtoPREFLEX = intersect(ToPreflex,FromClad);
    CladtoPOSTFLEX = intersect(ToPostflex,FromClad);
    nvmCaltoPREFLEX = intersect(ToPreflex,FromnvmCal);
    nvmCaltoPOSTFLEX = intersect(ToPostflex,FromnvmCal);
    PoeciltoPREFLEX = intersect(ToPreflex,FromPoecil);
    PoeciltoPOSTFLEX = intersect(ToPostflex,FromPoecil);


    for i=1:len
        flows=MCMCmat(i,:)./wts';

        mic2preflex_percent(i)=flows(MICtoPREFLEX)./sum(flows(ToPreflex))*100;
        app2preflex_percent(i)=flows(ApptoPREFLEX)./sum(flows(ToPreflex))*100;
        clad2preflex_percent(i)=flows(CladtoPREFLEX)./sum(flows(ToPreflex))*100;
        cal2preflex_percent(i)=flows(nvmCaltoPREFLEX)./sum(flows(ToPreflex))*100;
        poecil2preflex_percent(i)=flows(PoeciltoPREFLEX)./sum(flows(ToPreflex))*100;

        mic2postflex_percent(i)=flows(MICtoPOSTFLEX)./sum(flows(ToPostflex))*100;
        app2postflex_percent(i)=flows(ApptoPOSTFLEX)./sum(flows(ToPostflex))*100;
        clad2postflex_percent(i)=flows(CladtoPOSTFLEX)./sum(flows(ToPostflex))*100;
        cal2postflex_percent(i)=flows(nvmCaltoPOSTFLEX)./sum(flows(ToPostflex))*100;
        poecil2postflex_percent(i)=flows(PoeciltoPOSTFLEX)./sum(flows(ToPostflex))*100;


    end
    
    if Cycle==0
                fighandle = figure(1);
                fighandle.Units = 'inches';
                fighandle.Position = [1 1 8 3.5];
                col = [0 0 1];
    else
        col = [1 0 0];
    end
    subplot(1,2,1)
    hold on
    ViolinPlot(1+0.03*Cycle-0.15,0.4,mic2preflex_percent,200,col)
    ViolinPlot(2+0.03*Cycle-0.15,0.4,app2preflex_percent,200,col)
    ViolinPlot(3+0.03*Cycle-0.15,0.4,clad2preflex_percent,200,col)
    ViolinPlot(4+0.03*Cycle-0.15,0.4,cal2preflex_percent,200,col)
    ViolinPlot(5+0.03*Cycle-0.15,0.4,poecil2preflex_percent,200,col)
    set(gca,'box','on')
    set(gca,'XTick',1:5)
    set(gca,'XTickLabel',{'Microzooplankton','Appendicularians','Cladocerans','Calanoids (NVM)','Poecilostomatoids'})
    set(gca,'XTickLabelRotation',45)
    ylabel('Percentage of Diet')
    set(gca,'FontSize',9)
    %title('Preflex ABT')
    ylim([0 100])
    xlim([0.5 5.5])
    AddPanelLabels('a) Preflexion bluefin larvae','topleft','YPADDING',0.01,'XPADDING',0.01,'FONTSIZE',9)

    subplot(1,2,2)
    hold on
    ViolinPlot(1+0.03*Cycle-0.15,0.4,mic2postflex_percent,200,col)
    ViolinPlot(2+0.03*Cycle-0.15,0.4,app2postflex_percent,200,col)
    ViolinPlot(3+0.03*Cycle-0.15,0.4,clad2postflex_percent,200,col)
    ViolinPlot(4+0.03*Cycle-0.15,0.4,cal2postflex_percent,200,col)
    ViolinPlot(5+0.03*Cycle-0.15,0.4,poecil2postflex_percent,200,col)
    set(gca,'box','on')
    set(gca,'XTick',1:5)
    set(gca,'XTickLabel',{'Microzooplankton','Appendicularians','Cladocerans','Calanoids (NVM)','Poecilostomatoids'})
    set(gca,'XTickLabelRotation',45)
    set(gca,'FontSize',9)
    ylabel('Percentage of Diet')
    %title('Postflex ABT')
    ylim([0 100])
    xlim([0.5 5.5])
    AddPanelLabels('b) Postflexion bluefin larvae','topleft','YPADDING',0.01,'XPADDING',0.01,'FONTSIZE',9)

        upper=99;
        lower=80;
        left = 3.6;
        right = 5.45;
    fill([left right right left left],[lower lower upper upper lower],'w')
    fill([left+0.3 left+0.1 left+0.1 left+0.3 left+0.3],[upper-(upper-lower)*1/9 upper-(upper-lower)*1/9 upper-(upper-lower)*4/9 upper-(upper-lower)*4/9  upper-(upper-lower)*1/9],'r','FaceColor',[0 0 1])
    text(left+0.35,upper-(upper-lower)*2.5/9,'Gulf of Mexico','FontSize',8)
    fill([left+0.3 left+0.1 left+0.1 left+0.3 left+0.3],[upper-(upper-lower)*5/9 upper-(upper-lower)*5/9 upper-(upper-lower)*8/9 upper-(upper-lower)*8/9  upper-(upper-lower)*5/9],'r','FaceColor',[1 0 0])
    text(left+0.35,upper-(upper-lower)*6.5/9,'Argo Basin','FontSize',8)

    origin = 'GoM Nitrogen & Trophic - NOAA RESTORE\Inverse Model\R Code\Test 50\PlotTunaDiet.m'
    save('TunaDiet.mat','mic2preflex_percent','app2preflex_percent','clad2preflex_percent','cal2preflex_percent','poecil2preflex_percent',...
        'mic2postflex_percent','app2postflex_percent','clad2postflex_percent','cal2postflex_percent','poecil2postflex_percent',...
        'origin')


    mic2preflex_percent=sort(mic2preflex_percent);
    app2preflex_percent=sort(app2preflex_percent);
    clad2preflex_percent=sort(clad2preflex_percent);
    cal2preflex_percent=sort(cal2preflex_percent);
    poecil2preflex_percent=sort(poecil2preflex_percent);
    ['Calanoid copepods comprised ',num2str(mean(cal2preflex_percent)),'% of ',char(10),...
        'preflexion ABT diets (95% C. I. = ',num2str(cal2preflex_percent(round(len*0.025))),' - ',num2str(cal2preflex_percent(round(len*0.975))),'%)']
    ['Microzooplankton were ',num2str(mean(mic2preflex_percent),3),'% of ',...
        'preflexion ABT diets (95% C. I. = ',num2str(mic2preflex_percent(round(len*0.025)),3),' - ',num2str(mic2preflex_percent(round(len*0.975)),3),'%)']
    ['Appendicularians were ',num2str(mean(app2preflex_percent),3),'% of ',...
        'preflexion ABT diets (95% C. I. = ',num2str(app2preflex_percent(round(len*0.025)),3),' - ',num2str(app2preflex_percent(round(len*0.975)),3),'%)']
    ['Cladocerans were ',num2str(mean(clad2preflex_percent),3),'% of ',...
        'preflexion ABT diets (95% C. I. = ',num2str(clad2preflex_percent(round(len*0.025)),3),' - ',num2str(clad2preflex_percent(round(len*0.975)),3),'%)']
    ['Poecilostomatoid copepods were ',num2str(mean(poecil2preflex_percent),3),'% of ',...
        'preflexion ABT diets (95% C. I. = ',num2str(poecil2preflex_percent(round(len*0.025)),3),' - ',num2str(poecil2preflex_percent(round(len*0.975)),3),'%)']


    mic2postflex_percent=sort(mic2postflex_percent);
    app2postflex_percent=sort(app2postflex_percent);
    clad2postflex_percent=sort(clad2postflex_percent);
    cal2postflex_percent=sort(cal2postflex_percent);
    poecil2postflex_percent=sort(poecil2postflex_percent);
    ['Although calanoid copepods were also the dominant dietary source for',char(10),'postflexion ABT (',num2str(mean(cal2postflex_percent)),'%',...
        '; 95% C. I. = ',num2str(cal2postflex_percent(round(len*0.025))),' - ',num2str(cal2postflex_percent(round(len*0.975))),'%)']
    ['Microzooplankton were ',num2str(mean(mic2postflex_percent),3),'% of ',...
        'postflexion ABT diets (95% C. I. = ',num2str(mic2postflex_percent(round(len*0.025)),3),' - ',num2str(mic2postflex_percent(round(len*0.975)),3),'%)']
    ['Appendicularians were ',num2str(mean(app2postflex_percent),3),'% of ',...
        'postflexion ABT diets (95% C. I. = ',num2str(app2postflex_percent(round(len*0.025)),3),' - ',num2str(app2postflex_percent(round(len*0.975)),3),'%)']
    ['Cladocerans were ',num2str(mean(clad2postflex_percent),3),'% of ',...
        'postflexion ABT diets (95% C. I. = ',num2str(clad2postflex_percent(round(len*0.025)),3),' - ',num2str(clad2postflex_percent(round(len*0.975)),3),'%)']
    ['Poecilostomatoid copepods were ',num2str(mean(poecil2postflex_percent),3),'% of ',...
        'postflexion ABT diets (95% C. I. = ',num2str(poecil2postflex_percent(round(len*0.025)),3),' - ',num2str(poecil2postflex_percent(round(len*0.975)),3),'%)']


    fn = 'ManuscriptValues.txt'
    fileID = fopen(fn,'a');
    if Cycle==0
        formatSpec =  '\n PlotTunaDiet.m \n ';
        fprintf(fileID,formatSpec)
    end
    formatSpec =  'For Cycle %.0f Preflexion ABT consumed %.0f percent calanoids, %.0f percent appendicularians %.1f percent cladocerans, and %.1f percent poecilostomatoids \n';
    fprintf(fileID,formatSpec,[Cycle, mean(cal2preflex_percent), mean(app2preflex_percent), mean(clad2preflex_percent), mean(poecil2preflex_percent)])
    formatSpec =  'For Cycle %.0f Postflexion ABT consumed %.0f percent calanoids, %.0f percent appendicularians %.1f percent cladocerans, and %.1f percent poecilostomatoids \n';
    fprintf(fileID,formatSpec,[Cycle, mean(cal2postflex_percent), mean(app2postflex_percent), mean(clad2postflex_percent), mean(poecil2postflex_percent)])
    fclose(fileID)

    ToMIC = find(Ae0(MICs_col,:)==1);
    ToApp = find(Ae0(Apps_col,:)==1);
    ToClad = find(Ae0(Clads_col,:)==1);
    TonvmCal = find(Ae0(nvmCals_col,:)==1);
    ToPoecil = find(Ae0(Poecils_col,:)==1);
    FromDtm = find(Ae0(Dtms_col,:)==-1);
    FromFlag = find(Ae0(Flags_col,:)==-1);
    FromCya = find(Ae0(Cyas_col,:)==-1);
    FromBac = find(Ae0(Bacs_col,:)==-1);
    FromHNF = find(Ae0(HNFs_col,:)==-1);

    MICtoNVMCAL = intersect(FromMIC,TonvmCal);
    HNFtoNVMCAL = intersect(FromHNF,TonvmCal);
    FLAGtoNVMCAL = intersect(FromFlag,TonvmCal);
    DTMtoNVMCAL = intersect(FromDtm,TonvmCal);

    HNFtoAPP = intersect(FromHNF,ToApp);
    FLAGtoAPP = intersect(FromFlag,ToApp);
    DTMtoAPP = intersect(FromDtm,ToApp);
    CYAtoAPP = intersect(FromCya,ToApp);
    BACtoAPP = intersect(FromBac,ToApp);

    HNFtoMIC = intersect(FromHNF,ToMIC);
    FLAGtoMIC = intersect(FromFlag,ToMIC);
    DTMtoMIC = intersect(FromDtm,ToMIC);
    CYAtoMIC = intersect(FromCya,ToMIC);
    BACtoMIC = intersect(FromBac,ToMIC);

    for i=1:len
        flows=MCMCmat(i,:)./wts';

        mic2nvmcal_percent(i)=flows(MICtoNVMCAL)./sum(flows(TonvmCal))*100;
        hnf2nvmcal_percent(i)=flows(HNFtoNVMCAL)./sum(flows(TonvmCal))*100;
        flag2nvmcal_percent(i)=flows(FLAGtoNVMCAL)./sum(flows(TonvmCal))*100;
        dtm2nvmcal_percent(i)=flows(DTMtoNVMCAL)./sum(flows(TonvmCal))*100;

        hnf2app_percent(i)=flows(HNFtoAPP)./sum(flows(ToApp))*100;
        flag2app_percent(i)=flows(FLAGtoAPP)./sum(flows(ToApp))*100;
        dtm2app_percent(i)=flows(DTMtoAPP)./sum(flows(ToApp))*100;
        cya2app_percent(i)=flows(CYAtoAPP)./sum(flows(ToApp))*100;
        bac2app_percent(i)=flows(BACtoAPP)./sum(flows(ToApp))*100;

        hnf2mic_percent(i)=flows(HNFtoMIC)./sum(flows(ToMIC))*100;
        flag2mic_percent(i)=flows(FLAGtoMIC)./sum(flows(ToMIC))*100;
        dtm2mic_percent(i)=flows(DTMtoMIC)./sum(flows(ToMIC))*100;
        cya2mic_percent(i)=flows(CYAtoMIC)./sum(flows(ToMIC))*100;
        bac2mic_percent(i)=flows(BACtoMIC)./sum(flows(ToMIC))*100;

    end

    dtm2nvmcal_percent = sort(dtm2nvmcal_percent);
    flag2nvmcal_percent = sort(flag2nvmcal_percent);
    hnf2nvmcal_percent = sort(hnf2nvmcal_percent);
    mic2nvmcal_percent = sort(hnf2nvmcal_percent);

    hnf2app_percent = sort(hnf2app_percent);
    flag2app_percent = sort(flag2app_percent);
    dtm2app_percent = sort(dtm2app_percent);
    cya2app_percent = sort(cya2app_percent);
    bac2app_percent = sort(bac2app_percent);

    hnf2mic_percent = sort(hnf2mic_percent);
    flag2mic_percent = sort(flag2mic_percent);
    dtm2mic_percent = sort(dtm2mic_percent);
    cya2mic_percent = sort(cya2mic_percent);
    bac2mic_percent = sort(bac2mic_percent);


    mic2nvmcal_meanpercent=meanvals(MICtoNVMCAL)./sum(meanvals(TonvmCal))*100;
    hnf2nvmcal_meanpercent=meanvals(HNFtoNVMCAL)./sum(meanvals(TonvmCal))*100;
    dtm2nvmcal_meanpercent=meanvals(DTMtoNVMCAL)./sum(meanvals(TonvmCal))*100;
    flag2nvmcal_meanpercent=meanvals(MICtoNVMCAL)./sum(meanvals(TonvmCal))*100;

    ['Diatoms comprised ',num2str(dtm2nvmcal_meanpercent,2),'% of nvmCal diets (',num2str(dtm2nvmcal_percent(0.025*len),2),' - ',num2str(dtm2nvmcal_percent(0.975*len),2),'%)']
    ['Mixotrophic flagellates comprised ',num2str(flag2nvmcal_meanpercent,2),'% of nvmCal diets (',num2str(flag2nvmcal_percent(0.025*len),2),' - ',num2str(flag2nvmcal_percent(0.975*len),2),'%)']
    ['HNF comprised ',num2str(hnf2nvmcal_meanpercent,2),'% of nvmCal diets (',num2str(hnf2nvmcal_percent(0.025*len),2),' - ',num2str(hnf2nvmcal_percent(0.975*len),2),'%)']
    ['Microzooplankton comprised ',num2str(mic2nvmcal_meanpercent,2),'% of nvmCal diets (',num2str(mic2nvmcal_percent(0.025*len),2),' - ',num2str(mic2nvmcal_percent(0.975*len),2),'%)']

    ['Cyanobacteria comprised ',num2str(mean(cya2app_percent),2),'% of appendicularian diets (',num2str(cya2app_percent(0.025*len),2),' - ',num2str(cya2app_percent(0.975*len),2),'%)']
    ['Diatoms comprised ',num2str(mean(dtm2app_percent),2),'% of appendicularian diets (',num2str(dtm2app_percent(0.025*len),2),' - ',num2str(dtm2app_percent(0.975*len),2),'%)']
    ['Flagellates comprised ',num2str(mean(flag2app_percent),2),'% of appendicularian diets (',num2str(flag2app_percent(0.025*len),2),' - ',num2str(flag2app_percent(0.975*len),2),'%)']
    ['HNF comprised ',num2str(mean(hnf2app_percent),2),'% of appendicularian diets (',num2str(hnf2app_percent(0.025*len),2),' - ',num2str(hnf2app_percent(0.975*len),2),'%)']
    ['Bacterial comprised ',num2str(mean(bac2app_percent),2),'% of appendicularian diets (',num2str(bac2app_percent(0.025*len),2),' - ',num2str(bac2app_percent(0.975*len),2),'%)']


    ['Cyanobacteria comprised ',num2str(mean(cya2mic_percent),2),'% of microzooplankton diets (',num2str(cya2mic_percent(0.025*len),2),' - ',num2str(cya2mic_percent(0.975*len),2),'%)']
    ['Diatoms comprised ',num2str(mean(dtm2mic_percent),2),'% of microzooplankton diets (',num2str(dtm2mic_percent(0.025*len),2),' - ',num2str(dtm2mic_percent(0.975*len),2),'%)']
    ['Flagellates comprised ',num2str(mean(flag2mic_percent),2),'% of microzooplankton diets (',num2str(flag2mic_percent(0.025*len),2),' - ',num2str(flag2mic_percent(0.975*len),2),'%)']
    ['HNF comprised ',num2str(mean(hnf2mic_percent),2),'% of microzooplankton diets (',num2str(hnf2mic_percent(0.025*len),2),' - ',num2str(hnf2mic_percent(0.975*len),2),'%)']
    ['Bacterial comprised ',num2str(mean(bac2mic_percent),2),'% of microzooplankton diets (',num2str(bac2mic_percent(0.025*len),2),' - ',num2str(bac2mic_percent(0.975*len),2),'%)']

end


figure(1)
fn = ['PlotTunaDiet'];
exportgraphics(gcf,['../../Figures/',fn,'.png'])
```
