## Supplementary material for "Comparative food-web analysis of bluefin tuna spawning habitats in the eastern Indian Ocean and Gulf of Mexico": Code Package: RunN15InverseRWGoM.R

```
#Note that first you have to set the directory: Session>Set Working Directory
rm(list=ls())
library(R.matlab)
library(limSolve)
library(MASS)
source('xsampleN15outputs.r')
source('ExternalFunctionsGoM.r')
Cycle <- 0

tmp <- paste('N15GoMInverseCycle',Cycle,'.mat',sep="")
data <- readMat(tmp, fixNames=TRUE)
h <- data[['h']]
b <- data[['b']]
ba <- data[['ba']]
be <- data[['be']]
Ae <- data[['Ae']]
Aa <- data[['Aa']]
Aa0 <- data[['Aa0']]
G <- data[['G']]
sdba <- data[['sdba']]
InputCol <- data[['InputCol']]
A <- data[['A']]
Inputs <- data[['Inputs']]
d15NInputs <- data[['d15NInputs']]
wts <- data[['wts']]
#W <- rep(1,39)
BurninLength1 <- 1000
BurninLength2 <- 1000
IterLength <- 500000
OutputLength <- IterLength/10000
jmpLength <- 0.003
jmpLength15 <- 0.02

Aa2 <- Aa[1:36,]
ba2 <- ba[1:36]
sdba2 <- sdba[1:36]
#Aa2 <- Aa[2:3,]
#ba2 <- ba[2:3]
#sdba2 <- sdba[2:3]
#G <- G[1:50,]
#h <- h[1:50]

d15N0 <- c(0,0,0,0,0,0,0,0,0,0,
           0,0,0,0,0,0,0,0,0,0,
           0,0,0,0,0,0,0,0,0,0,
           0,0,0,0,0,0,0,0,0,0,
           0,0,0,0,0,0)
d15N0 <- d15N0 + d15NInputs[1,InputCol]


upNO315N=d15NInputs[1,InputCol]
pre15N=d15NInputs[2,InputCol]
post15N=d15NInputs[3,InputCol]
LDetS15N=d15NInputs[4,InputCol]
LDetD15N=d15NInputs[5,InputCol]
DONS15N=d15NInputs[6,InputCol]
DOND15N=d15NInputs[7,InputCol]
AppD15N=d15NInputs[10,InputCol]
CalD15N=d15NInputs[11,InputCol]
ChaetoD15N=d15NInputs[12,InputCol]
HerbVMD15N=d15NInputs[13,InputCol]
HerbNVMD15N=d15NInputs[14,InputCol]
PoecilD15N=d15NInputs[15,InputCol]
CladD15N=d15NInputs[16,InputCol]
del15Nknown = c(upNO315N,pre15N,post15N,LDetS15N,LDetD15N,DONS15N,DOND15N,AppD15N,CalD15N,ChaetoD15N,HerbVMD15N,HerbNVMD15N,PoecilD15N,CladD15N)


#G <- G[76:183,]
#h <- h[76:183]

#L2MN Solution
test <- lsei(A = Aa2, B = ba2, E = Ae, F = be, G = G, H = h, type=2)
X <- test[['X']]
solNorm <- test[['solutionNorm']]
lseisol <- as.matrix(X)
rm(test, X)
fileout <- paste('N15GoMInverseLSEICycle',Cycle,'Routputs.mat',sep="")
writeMat(fileout, Aa = Aa2, ba = ba2, Ae = Ae, be = be, G = G, h = h, lseisol = lseisol, wts=wts)


#Central Value Solution
center <- xranges(E = Ae, F = be, G = G, H = h,
                  ispos = FALSE, tol = 1e-8, central = TRUE, full=FALSE)
centralval <- center[,3]


#Burnin - Plain
test2 <- xsample(A = Aa2, B = ba2, E = Ae, F = be, G = G,
                 H = h, sdB = sdba2*10, iter = BurninLength1, type="mirror", jmp=jmpLength+runif(1, 0, 1)*jmpLength/5, x0 = centralval, fulloutput='TRUE')
Burninmat <- test2[['X']]
Startpt <- Burninmat[BurninLength1,]
rm(Burninmat)

# test2 <- xsample(A = Aa2, B = ba2, E = Ae, F = be, G = G,
#                  H = h, sdB = sdba2*10, iter = BurninLength2, type="mirror", jmp=jmpLength+runif(1, 0, 1)*jmpLength/5, x0 = Startpt, fulloutput='TRUE')
# Burninmat <- test2[['X']]
# Startpt <- Burninmat[BurninLength2,]
# rm(Burninmat)

test2 <- xsample(A = Aa2, B = ba2, E = Ae, F = be, G = G,
                 H = h, sdB = sdba2, iter = IterLength/10, outputlength = OutputLength, type="mirror", jmp=jmpLength+runif(1, 0, 1)*jmpLength/5, x0 = Startpt, fulloutput='TRUE')

MCMCmatplain <- test2[['X']]

MCMCmatmean <- colMeans(MCMCmatplain)


Eps_Remin <- -1
Eps_Eg <- -2
Eps_NH4up <- -10
sdbafactor <- -Eps_Remin
#sdPPF <- Inputs[84,InputCol+1]
#sdHFPF <- Inputs[83,InputCol+1]
#sdba[20] <- sdPPF
#sdba[21] <- sdHFPF
sdba[41] <- sdbafactor
sdba[42] <- sdbafactor
sdba[43] <- sdbafactor
sdba[44] <- sdbafactor
sdba[45] <- sdbafactor
sdba[46] <- sdbafactor
sdba[47] <- sdbafactor
sdba[48] <- sdbafactor
sdba[49] <- sdbafactor
sdba[50] <- sdbafactor
sdba[51] <- sdbafactor
sdba[52] <- sdbafactor
sdba[53] <- sdbafactor
sdba[54] <- sdbafactor
sdba[55] <- sdbafactor
sdba[56] <- sdbafactor
sdba[57] <- sdbafactor
sdba[58] <- sdbafactor
sdba[59] <- sdbafactor
sdba[60] <- sdbafactor
sdba[61] <- sdbafactor
sdba[62] <- sdbafactor
sdba[63] <- sdbafactor
sdba[64] <- sdbafactor
sdba[65] <- sdbafactor
sdba[66] <- sdbafactor
sdba[67] <- sdbafactor
sdba[68] <- sdbafactor
sdba[69] <- sdbafactor
sdba[70] <- sdbafactor
sdba[71] <- sdbafactor
sdba[72] <- sdbafactor
sdba[73] <- sdbafactor
sdba[74] <- sdbafactor
sdba[75] <- sdbafactor
sdba[76] <- sdbafactor
sdba[77] <- sdbafactor
sdba[78] <- sdbafactor
sdba[79] <- sdbafactor
sdba[80] <- sdbafactor
sdba[81] <- sdbafactor
sdba[82] <- sdbafactor
sdba[83] <- sdbafactor
sdba[84] <- sdbafactor


#Burnin
test2 <- xsampleN15outputs(A = Aa0, B = ba, E = Ae, F = be, G = G,
                 H = h, sdB = sdba*10, wts = wts, iter = BurninLength2, burninlength = BurninLength1, type="mirror", jmp=jmpLength+runif(1, 0, 1)*jmpLength/5, jmp2=jmpLength15/5, x0 = MCMCmatmean, del15N1=d15N0, del15Nknown=del15Nknown, fulloutput='TRUE')
Burninmat <- test2[['X']]
del15Ntemp <- test2[['del15Ntrack']]
d15N0 <- del15Ntemp[BurninLength2,]
Startpt <- Burninmat[BurninLength2,]
rm(del15Ntemp, Burninmat)

test2 <- xsampleN15outputs(A = Aa0, B = ba, E = Ae, F = be, G = G,
                 H = h, sdB = sdba, wts=wts, iter = IterLength, outputlength = OutputLength, type="mirror", jmp=jmpLength+runif(1, 0, 1)*jmpLength/5, jmp2=jmpLength15, x0 = Startpt, del15N1=d15N0, del15Nknown=del15Nknown, fulloutput='TRUE')
# Xa1 <- test2[['Xa1']]
# Xa2 <- test2[['Xa2']]
# Aa1 <- test2[['Aa1']]
# Aa2 <- test2[['Aa2']]
# SSRq1 <- test2[['SSRq1']]
# SSRq2 <- test2[['SSRq2']]

MCMCmat <- test2[['X']]
MCMCmat_rejects <- test2[['x_rejects']]
del15N <- test2[['del15Ntrack']]
del15N_rejects <- test2[['del15Ntrack_rejects']]
randomnumber <- test2[['randomnumber']]
p <- test2[['p']]
p_rejects <- test2[['p_rejects']]
err <- test2[['err']]
acceptedratio <- test2[['acceptedratio']]
fileout <- paste('N15GoMInverseCycle',Cycle,'Routputs.mat',sep="")
writeMat(fileout, MCMCmat = MCMCmat, MCMCmat_rejects = MCMCmat_rejects, MCMCmatplain = MCMCmatplain, del15N=del15N, del15N_rejects=del15N_rejects, Aa = Aa, ba = ba, Ae = Ae, be = be, G = G, h = h, sdba = sdba, Inputs = Inputs, InputCol = InputCol, jmpLength = jmpLength, jmpLength15 = jmpLength15, acceptedratio = acceptedratio, Startpt = Startpt, centralval = centralval, p=p, p_rejects=p_rejects, err=err, del15Nknown=del15Nknown, lseisol=lseisol, wts=wts)

# N <- 1:(IterLength/100)
# N <- N*100
# MCMCmat <- MCMCmattemp[N,]
# rm(MCMCmattemp,N)
# 
# 
#writeMat('N15InverseModel.NEMURO.Coastal.ROutputs.mat', MCMCmat = MCMCmat, del15N=del15N, Xa1=Xa1, Xa2=Xa2, Aa1=Aa1, Aa2=Aa2, SSRq1=SSRq1, SSRq2=SSRq2, ba=ba, sdba=sdba, randomnumber=randomnumber)
#plot(MCMCmat[,1], type="o", col="blue")

# Aa3 = ResetRN15(Aa,del15N[OutputLength,],del15Nknown[1],del15Nknown[2],del15Nknown[3],del15Nknown[4],del15Nknown[5])
# errors=(Aa3%*%MCMCmat[OutputLength,]-ba)/sdba
# sum(errors^2)
```
