## Supplementary material for "Comparative food-web analysis of bluefin tuna spawning habitats in the eastern Indian Ocean and Gulf of Mexico": Code Package: SecondaryProductionDeterminants3.m

```
clearvars
close all

burnin = 0.2;
PlotPhyEatenPercent = 0;

for Cycle=[0,10]
    clearvars -except Cycle burnin keep PlotPhyEatenPercent

    load(['N15GoMInverseCycle',num2str(Cycle),'.mat'])
    load(['N15GoMInverseCycle',num2str(Cycle),'Routputs.mat'])
    clearvars -except Ae0 MCMCmat Aa d15NInputs del15N InputCol wts Cycle burnin keep PlotPhyEatenPercent
    num=length(MCMCmat(:,1));
    MCMCmat(1:round(num*burnin),:)=[];
    del15N(1:round(num*burnin),:)=[];
    len=length(MCMCmat(:,1));
    [num,txt,raw]  = xlsread('N15InverseModelRW.GOM.xlsx','Sheet1','B6:B307');
    Columns = txt;
    meanvals = mean(MCMCmat)'./wts;
    stdvals = std(MCMCmat)'./wts;
    meantable = table(Columns,meanvals,stdvals);

    [NO3s_col,NH4s_col,Cyas_col,Tris_col,Dtms_col,Flags_col,HNFs_col,MICs_col,HerbNVMs_col,Apps_col,Clads_col,nvmCals_col,Chaetos_col,Poecils_col,GelPreds_col,Plankfishs_col,Preflexs_col,Postflexs_col,Bacs_col,Dets_col,Ldets_col,Dons_col,HerbVM_col,vmCal_col,NO3d_col,NH4d_col,Cyad_col,Trid_col,Dtmd_col,Flagd_col,HNFd_col,MICd_col,HerbNVMd_col,Appd_col,Cladd_col,nvmCald_col,Chaetod_col,Poecild_col,GelPredd_col,Plankfishd_col,Bacd_col,Detd_col,Ldetd_col,Dond_col] = GetColumns(Aa,MCMCmat,del15N(1,:),d15NInputs(1,InputCol),d15NInputs(2,InputCol),d15NInputs(3,InputCol),d15NInputs(4,InputCol),d15NInputs(5,InputCol),d15NInputs(6,InputCol),d15NInputs(7,InputCol));
    
    

    for i=1:length(MCMCmat(:,1))
        i
        flows=MCMCmat(i,:)./wts';
        TL0 = CalcTL(flows,del15N,d15NInputs,Ae0,InputCol);
        TL_track(:,i)=TL0;
    end

    
    ToMesozoos = sort([find(Ae0(HerbNVMs_col,:)==1),find(Ae0(Apps_col,:)==1),find(Ae0(Clads_col,:)==1),find(Ae0(nvmCals_col,:)==1),find(Ae0(Chaetos_col,:)==1),find(Ae0(Poecils_col,:)==1),find(Ae0(HerbVM_col,:)==1),find(Ae0(vmCal_col,:)==1)]);
    ToMesozood = sort([find(Ae0(HerbNVMd_col,:)==1),find(Ae0(Appd_col,:)==1),find(Ae0(Cladd_col,:)==1),find(Ae0(nvmCald_col,:)==1),find(Ae0(Chaetod_col,:)==1),find(Ae0(Poecild_col,:)==1),find(Ae0(HerbVM_col,:)==1),find(Ae0(vmCal_col,:)==1)]);
    FromMesozoos = sort([find(Ae0(HerbNVMs_col,:)==-1),find(Ae0(Apps_col,:)==-1),find(Ae0(Clads_col,:)==-1),find(Ae0(nvmCals_col,:)==-1),find(Ae0(Chaetos_col,:)==-1),find(Ae0(Poecils_col,:)==-1),find(Ae0(HerbVM_col,:)==-1),find(Ae0(vmCal_col,:)==-1)]);
    FromMesozood = sort([find(Ae0(HerbNVMd_col,:)==-1),find(Ae0(Appd_col,:)==-1),find(Ae0(Cladd_col,:)==-1),find(Ae0(nvmCald_col,:)==-1),find(Ae0(Chaetod_col,:)==-1),find(Ae0(Poecild_col,:)==-1),find(Ae0(HerbVM_col,:)==-1),find(Ae0(vmCal_col,:)==-1)]);
    ToHTLs = sort([find(Ae0(GelPreds_col,:)==1),find(Ae0(Preflexs_col,:)==1),find(Ae0(Postflexs_col,:)==1),find(Ae0(Plankfishs_col,:)==1)]);
    ToHTLd = sort([find(Ae0(HerbNVMd_col,:)==1),find(Ae0(Plankfishd_col,:)==1)]);

    ToPreflexs = find(Ae0(Preflexs_col,:)==1);
    ToPostflexs = find(Ae0(Postflexs_col,:)==1);
    ToGelPreds = find(Ae0(GelPreds_col,:)==1);
    ToPlankfishs = find(Ae0(Plankfishs_col,:)==1);
    ToGelPredd = find(Ae0(GelPredd_col,:)==1);
    ToPlankfishd = find(Ae0(Plankfishd_col,:)==1);
    

    FromHTLs = sort([find(Ae0(GelPreds_col,:)==-1),find(Ae0(Preflexs_col,:)==-1),find(Ae0(Postflexs_col,:)==-1),find(Ae0(Plankfishs_col,:)==-1)]);
    FromHTLd = sort([find(Ae0(HerbNVMd_col,:)==-1),find(Ae0(Plankfishd_col,:)==-1)]);
    FromPhys = sort([find(Ae0(Cyas_col,:)==-1),find(Ae0(Tris_col,:)==-1),find(Ae0(Dtms_col,:)==-1)]);
    FromPhyd = sort([find(Ae0(Cyad_col,:)==-1),find(Ae0(Trid_col,:)==-1),find(Ae0(Dtmd_col,:)==-1)]);
    FromMixs = find(Ae0(Flags_col,:)==-1);
    FromMixd = find(Ae0(Flagd_col,:)==-1);
    ToMixs = find(Ae0(Flags_col,:)==1);
    ToMixd = find(Ae0(Flagd_col,:)==1);
    FromCyas = find(Ae0(Cyas_col,:)==-1);
    FromCyad = find(Ae0(Cyad_col,:)==-1);
    ToCyas = find(Ae0(Cyas_col,:)==1);
    ToCyad = find(Ae0(Cyad_col,:)==1);
    FromNO3s = find(Ae0(NO3s_col,:)==-1);
    FromNO3d = find(Ae0(NO3d_col,:)==-1);
    FromNH4s = find(Ae0(NH4s_col,:)==-1);
    FromNH4d = find(Ae0(NH4d_col,:)==-1);
    ToNH4s = find(Ae0(NH4s_col,:)==1);
    ToNH4d = find(Ae0(NH4d_col,:)==1);
    ToDons = find(Ae0(Dons_col,:)==1);
    ToDond = find(Ae0(Dond_col,:)==1);
    ToDets = find(Ae0(Dets_col,:)==1);
    ToLdets = find(Ae0(Ldets_col,:)==1);
    ToDetd = find(Ae0(Detd_col,:)==1);
    ToLdetd = find(Ae0(Ldetd_col,:)==1);
    FromBacs = find(Ae0(Bacs_col,:)==-1);
    FromBacd = find(Ae0(Bacd_col,:)==-1);
    ToBacs = find(Ae0(Bacs_col,:)==1);
    ToBacd = find(Ae0(Bacd_col,:)==1);
    FromProtists = sort([find(Ae0(HNFs_col,:)==-1),find(Ae0(MICs_col,:)==-1)]);
    FromProtistd = sort([find(Ae0(HNFd_col,:)==-1),find(Ae0(MICd_col,:)==-1)]);
    ToProtists = sort([find(Ae0(HNFs_col,:)==1),find(Ae0(MICs_col,:)==1)]);
    ToTops = sort([find(Ae0(GelPreds_col,:)==1),find(Ae0(Plankfishs_col,:)==1),find(Ae0(Preflexs_col,:)==1),find(Ae0(Postflexs_col,:)==1)]);
    ToTopd = sort([find(Ae0(GelPredd_col,:)==1),find(Ae0(Plankfishd_col,:)==1)]);
    FromTops = sort([find(Ae0(GelPreds_col,:)==-1),find(Ae0(Plankfishs_col,:)==-1),find(Ae0(Preflexs_col,:)==-1),find(Ae0(Postflexs_col,:)==-1)]);
    FromTopd = sort([find(Ae0(GelPredd_col,:)==-1),find(Ae0(Plankfishd_col,:)==-1)]);
    ToProtistd = sort([find(Ae0(HNFd_col,:)==1),find(Ae0(MICd_col,:)==1)]);
    FromDtms = find(Ae0(Dtms_col,:)==-1);
    FromDtmd = find(Ae0(Dtmd_col,:)==-1);
    ToDtms = find(Ae0(Dtms_col,:)==1);
    ToDtmd = find(Ae0(Dtmd_col,:)==1);
    FromTris = find(Ae0(Tris_col,:)==-1);
    FromTrid = find(Ae0(Trid_col,:)==-1);
    ToTris = find(Ae0(Tris_col,:)==1);
    ToTrid = find(Ae0(Trid_col,:)==1);
    ToMesozoo = sort([ find(Ae0(HerbNVMs_col,:)==1), find(Ae0(Apps_col,:)==1), find(Ae0(Clads_col,:)==1), find(Ae0(nvmCals_col,:)==1), find(Ae0(Chaetos_col,:)==1), find(Ae0(Poecils_col,:)==1), ...
        find(Ae0(HerbVM_col,:)==1), find(Ae0(vmCal_col,:)==1), ...
        find(Ae0(HerbNVMd_col,:)==1), find(Ae0(Appd_col,:)==1), find(Ae0(Cladd_col,:)==1), find(Ae0(nvmCald_col,:)==1), find(Ae0(Chaetod_col,:)==1), find(Ae0(Poecild_col,:)==1)     ]);
    FromMesozoo = sort([ find(Ae0(HerbNVMs_col,:)==-1), find(Ae0(Apps_col,:)==-1), find(Ae0(Clads_col,:)==-1), find(Ae0(nvmCals_col,:)==-1), find(Ae0(Chaetos_col,:)==-1), find(Ae0(Poecils_col,:)==-1), ...
        find(Ae0(HerbVM_col,:)==-1), find(Ae0(vmCal_col,:)==-1), ...
        find(Ae0(HerbNVMd_col,:)==-1), find(Ae0(Appd_col,:)==-1), find(Ae0(Cladd_col,:)==-1), find(Ae0(nvmCald_col,:)==-1), find(Ae0(Chaetod_col,:)==-1), find(Ae0(Poecild_col,:)==-1)     ]);

    PHYtoMESOZOOs = intersect(ToMesozoos,FromPhys);
    PHYtoMESOZOOd = intersect(ToMesozood,FromPhyd);
    PHYtoNONLIVING = intersect([ToDons,ToDond,ToDets,ToDetd,ToLdets,ToLdetd],[FromPhys,FromPhyd]);
    PHYtoDETRITUS = intersect([ToDets,ToDetd,ToLdets,ToLdetd],[FromPhys,FromPhyd]);
    FLAGtoMESOZOOs = intersect(ToMesozoos,FromMixs);
    FLAGtoMESOZOOd = intersect(ToMesozood,FromMixd);
    BACtoFLAGs = intersect(FromBacs,ToMixs);
    BACtoFLAGd = intersect(FromBacd,ToMixd);
    CYAtoFLAGs = intersect(FromCyas,ToMixs);
    CYAtoFLAGd = intersect(FromCyad,ToMixd);
    NUTtoFLAGs = intersect([FromNO3s,FromNH4s],ToMixs);
    NUTtoFLAGd = intersect([FromNO3d,FromNH4d],ToMixd);
    FLAGtoDONs = intersect(FromMixs,ToDons);
    FLAGtoDONd = intersect(FromMixd,ToDond);
    CYAtoDONs = intersect(FromCyas,ToDons);
    CYAtoDONd = intersect(FromCyad,ToDond);
    DTMtoDONs = intersect(FromDtms,ToDons);
    DTMtoDONd = intersect(FromDtmd,ToDond);
    TRICHOtoDONs = intersect(FromTris,ToDons);
    TRICHOtoDONd = intersect(FromTrid,ToDond);
    NUTtoDTMs = intersect([FromNO3s,FromNH4s],ToDtms);
    NUTtoDTMd = intersect([FromNO3d,FromNH4d],ToDtmd);
    DTMtoDONs = intersect(FromDtms,ToDons);
    DTMtoDONd = intersect(FromDtmd,ToDond);
    NUTtoCYAs = intersect([FromNO3s,FromNH4s],ToCyas);
    NUTtoCYAd = intersect([FromNO3d,FromNH4d],ToCyad);
    FLAGtoCYAs = intersect(FromCyas,ToDons);
    FLAGtoCYAd = intersect(FromCyad,ToDond);
    NUTtoTRICHOs = intersect([FromNO3s,FromNH4s],ToTris);
    NUTtoTRICHOd = intersect([FromNO3d,FromNH4d],ToTrid);
    FLAGtoTRICHOs = intersect(FromTris,ToDons);
    FLAGtoTRICHOd = intersect(FromTrid,ToDond);
    PROTISTtoMESOZOOs = intersect(ToMesozoos,FromProtists);
    PROTISTtoMESOZOOd = intersect(ToMesozood,FromProtistd);
    BACtoNH4s = intersect(FromBacs,ToNH4s);
    BACtoNH4d = intersect(FromBacd,ToNH4d);
    BACtoPROTISTs = intersect(FromBacs,ToProtists);
    BACtoPROTISTd = intersect(FromBacd,ToProtistd);
    PROTISTtoDONs = intersect(FromProtists,ToDons);
    PROTISTtoDONd = intersect(FromProtistd,ToDond);
    PROTISTtoNH4s = intersect(FromProtists,ToNH4s);
    PROTISTtoNH4d = intersect(FromProtistd,ToNH4d);
    NO3toPHYs = intersect(FromNO3s,[ToDtms,ToCyas,ToTris,ToMixs]);
    NO3toPHYd = intersect(FromNO3d,[ToDtmd,ToCyad,ToTris,ToMixd]);
    NH4toPHYs = intersect(FromNH4s,[ToDtms,ToCyas,ToTris,ToMixs]);
    NH4toPHYd = intersect(FromNH4d,[ToDtmd,ToCyad,ToTris,ToMixd]);
    PHYtoPROTIST = intersect([FromPhys,FromMixs,FromPhyd,FromMixd],[ToProtists,ToProtistd]);
    PHYtoDONs = intersect(FromPhys,ToDons);
    MIXtoDONs = intersect(FromMixs,ToDons);
    MESOZOOtoDONs = intersect(FromMesozoos,ToDons);
    HTLtoDONs = intersect(FromHTLs,ToDons);
    BACtoNH4s = intersect(FromBacs,ToNH4s);
    MIXtoNH4s = intersect(FromMixs,ToNH4s);
    MESOZOOtoNH4s = intersect(FromMesozoos,ToNH4s);
    HTLtoNH4s = intersect(FromHTLs,ToNH4s);

    MESOZOO_production = intersect(FromMesozoo,[ToMesozoo,ToHTLs,ToHTLd]);
    PROTIST_production = intersect([FromProtists,FromProtistd],[ToProtists,ToProtistd,ToMesozoo,ToHTLs,ToHTLd]);
    PLANKFISH_2PROD = find(strcmp(meantable.Columns,'plankfishTOhtls') | strcmp(meantable.Columns,'plankfishTOhtld'));
    GELPRED_2PROD = find(strcmp(meantable.Columns,'gelTOhtls') | strcmp(meantable.Columns,'gelTOhtld'));
    PREFLEX_2PROD = find(strcmp(meantable.Columns,'preflexTOgrowths') | strcmp(meantable.Columns,'preflexTOmorts'));
    POSTFLEX_2PROD = find(strcmp(meantable.Columns,'postflexTOgrowths') | strcmp(meantable.Columns,'postflexTOmorts'));
    PLANKFISH_2PRODs = find(strcmp(meantable.Columns,'plankfishTOhtls'));
    GELPRED_2PRODs = find(strcmp(meantable.Columns,'gelTOhtls'));
    PLANKFISH_2PRODd = find(strcmp(meantable.Columns,'plankfishTOhtld'));
    GELPRED_2PRODd = find(strcmp(meantable.Columns,'gelTOhtld'));


    for i=1:len
        i
        flows=MCMCmat(i,:)./wts';

        CYAnpps(i) = sum(flows(ToCyas)) - flows(CYAtoDONs);
        TRInpps(i) = sum(flows(ToTris)) - flows(TRICHOtoDONs);
        DTMnpps(i) = sum(flows(NUTtoDTMs)) - flows(DTMtoDONs);
        FLAGnpps(i) = sum(flows(NUTtoFLAGs)) - flows(FLAGtoDONs);
        CYAnppd(i) = sum(flows(ToCyad)) - flows(CYAtoDONd);
        TRInppd(i) = sum(flows(ToTrid)) - flows(TRICHOtoDONd);
        DTMnppd(i) = sum(flows(NUTtoDTMd)) - flows(DTMtoDONd);
        FLAGnppd(i) = sum(flows(NUTtoFLAGd)) - flows(FLAGtoDONd);

        CYAgpps(i) = sum(flows(ToCyas)) ;
        TRIgpps(i) = sum(flows(ToTris)) ;
        DTMgpps(i) = sum(flows(NUTtoDTMs)) ;
        FLAGgpps(i) = sum(flows(NUTtoFLAGs)) ;
        CYAgppd(i) = sum(flows(ToCyad)) ;
        TRIgppd(i) = sum(flows(ToTrid)) ;
        DTMgppd(i) = sum(flows(NUTtoDTMd)) ;
        FLAGgppd(i) = sum(flows(NUTtoFLAGd)) ;
        GPP(i) = CYAgpps(i) + TRIgpps(i) + DTMgpps(i) + FLAGgpps(i) + CYAgppd(i) + TRIgppd(i) + DTMgppd(i) + FLAGgppd(i);
        PHY2NONLIVING(i) = sum(flows(PHYtoNONLIVING));
        PHY2DETRITUS(i) = sum(flows(PHYtoDETRITUS));

        FLAGmixs = flows(BACtoFLAGs)+flows(CYAtoFLAGs);
        HerbFoodChains(i) = sum(flows(PHYtoMESOZOOs)) + sum(flows(FLAGtoMESOZOOs))*FLAGnpps(i)/(FLAGnpps(i)+FLAGmixs);
        FLAGmixd = flows(BACtoFLAGd)+flows(CYAtoFLAGd);
        HerbFoodChaind(i) = sum(flows(PHYtoMESOZOOd)) + sum(flows(FLAGtoMESOZOOd))*FLAGnppd(i)/(FLAGnppd(i)+FLAGmixd);

        MultivFoodChains(i) = sum(flows(PROTISTtoMESOZOOs)) + sum(flows(FLAGtoMESOZOOs))*sum(flows(CYAtoFLAGs))/(FLAGnpps(i)+FLAGmixs);
        MultivFoodChaind(i) = sum(flows(PROTISTtoMESOZOOd)) + sum(flows(FLAGtoMESOZOOd))*sum(flows(CYAtoFLAGd))/(FLAGnppd(i)+FLAGmixd);

        ProtistRespirationPercents = sum(flows(PROTISTtoNH4s))/sum(flows(ToProtists));
        MicrobialLoops(i) = flows(BACtoNH4s) + sum(flows(BACtoPROTISTs))*ProtistRespirationPercents;
        ProtistRespirationPercentd = sum(flows(PROTISTtoNH4d))/sum(flows(ToProtistd));
        MicrobialLoopd(i) = flows(BACtoNH4d) + sum(flows(BACtoPROTISTd))*ProtistRespirationPercentd;
        BacDemands(i) = flows(ToBacs);
        BacDemandd(i) = flows(ToBacd);
        BacRespirations(i) = flows(BACtoNH4s);
        BacRespirationd(i) = flows(BACtoNH4d);

        NPPs(i) = CYAnpps(i) + TRInpps(i) + DTMnpps(i) + FLAGnpps(i);
        NPPd(i) = CYAnppd(i) + TRInppd(i) + DTMnppd(i) + FLAGnppd(i);
        NPP(i) = NPPs(i) + NPPd(i);

        NO3UPTAKEs(i) = sum(flows(NO3toPHYs));
        NO3UPTAKEd(i) = sum(flows(NO3toPHYd));
        NH4UPTAKEs(i) = sum(flows(NH4toPHYs));
        NH4UPTAKEd(i) = sum(flows(NH4toPHYd));
        N2FIXs(i) = sum(flows([ToTris,ToCyas])) - sum(flows([NUTtoCYAs,NUTtoTRICHOs]));
        N2FIXd(i) = sum(flows([ToTrid,ToCyad])) - sum(flows([NUTtoCYAd,NUTtoTRICHOd]));

        PROTISTGRAZING(i) = sum(flows(PHYtoPROTIST));
        %MESOZOOGRAZING(i) = sum(flows(PHYtoMESOZOOs)) + sum(flows(FLAGtoMESOZOOs)) + sum(flows(PHYtoMESOZOOd)) + sum(flows(FLAGtoMESOZOOd));

        PhytoplanktonDONproductions(i)=sum(flows(PHYtoDONs)) + sum(flows(MIXtoDONs))*FLAGnpps(i)/(FLAGnpps(i)+FLAGmixs);
        ProtistDONproductions(i)=sum(flows(PROTISTtoDONs)) + sum(flows(MIXtoDONs))*FLAGmixs/(FLAGnpps(i)+FLAGmixs);
        MesozooDONproductions(i)=sum(flows(MESOZOOtoDONs));
        HTLDONproductions(i)=sum(flows(HTLtoDONs));
        TotalDONproductions(i)=sum(flows(ToDons));

        BacNH4productions(i)=sum(flows(BACtoNH4s));
        ProtistNH4productions(i)=sum(flows(PROTISTtoNH4s)) + sum(flows(MIXtoNH4s))*FLAGmixs/(FLAGnpps(i)+FLAGmixs);
        MesozooNH4productions(i)=sum(flows(MESOZOOtoNH4s));
        HTLNH4productions(i)=sum(flows(HTLtoNH4s));
        TotalNH4productions(i)=sum(flows(ToNH4s));

        HTLproduction(i) = sum(flows(PLANKFISH_2PROD)) + sum(flows(GELPRED_2PROD)) + sum(flows(PREFLEX_2PROD)) + sum(flows(POSTFLEX_2PROD));
        PREFLEXproduction(i) = sum(flows(PREFLEX_2PROD));
        POSTFLEXproduction(i) = sum(flows(POSTFLEX_2PROD));
        GELPREDproductions(i) = sum(flows(GELPRED_2PRODs));
        GELPREDproductiond(i) = sum(flows(GELPRED_2PRODd));
        PLANKFISHproductions(i) = sum(flows(PLANKFISH_2PRODs));
        PLANKFISHproductiond(i) = sum(flows(PLANKFISH_2PRODd));

        Mesozoo_consumption(i) = sum(flows(ToMesozoo));
        Mesozoo_production(i) = sum(flows(MESOZOO_production));


        Protist_consumption(i) = sum(flows([ToProtists,ToProtistd]));
        Protist_production(i) = sum(flows(PROTIST_production));
    end

    %---------------------------------------------------------------
    %------------Start Figure---------------------------------------
    %---------------------------------------------------------------


    numsubplots = 10;
    for j=1:1
        if j==1
            if Cycle==0
                fighandle = figure(1);
                fighandle.Units = 'inches';
                fighandle.Position = [1 1 11 4];
            else
                figure(1)
            end
        else
            if Cycle==0
                fighandle = figure(2);
                fighandle.Units = 'inches';
                fighandle.Position = [2.6 1 6.5 3];
            else
                figure(2)
            end
        end
        if Cycle==0
            col = [0 0 1];
        else
            col = [1 0 0];
        end

        %-------------------------Ecosystem Production--------------------------------
        subplot(1,numsubplots,2)
        hold on
        ViolinPlot(Cycle*0.1+1,0.5,HTLproduction,200,col)

        set(gca,'box','on')
        %set(gca,'XTick',[2,6])
        %set(gca,'XTickLabel',{['Shallow Eup Zone'],'Deep Chl Max'})
        %set(gca,'XTickLabelRotation',20)
        ylabel('Ecosystem Production (mmol N m^-^2 d^-^1)')
        set(gca,'FontSize',9)
        if Cycle==0
            keep.maxHTLprod = max(HTLproduction);
        else
            keep.maxHTLprod = max([HTLproduction,max(HTLproduction)]);
        end
        ylim([0 max([0,keep.maxHTLprod]) ])
        set(gca,'XTick',1)
        set(gca,'XTickLabel',{''})
        if Cycle==10
            AddPanelLabels('a','bottomright','YPADDING',0.02,'XPADDING',0.15,'FONTSIZE',9)
        end

    fn = 'ManuscriptValues.txt'
    fileID = fopen(fn,'a');
    if Cycle==0
        formatSpec =  '\n FoodwebIndices.m \n ';
        fprintf(fileID,formatSpec)
    end
    formatSpec =  'For Cycle %f Ecosystem Production was mmol N m-2 d-1  %f +/- %f \n';
    fprintf(fileID,formatSpec,[Cycle, mean(HTLproduction), std(HTLproduction)])


        %-------------------------NPP--------------------------------
        subplot(1,numsubplots,3)
        hold on
        ViolinPlot(Cycle*0.1+1,0.5,NPPs+NPPd,200,col)

        set(gca,'box','on')
        %set(gca,'XTick',[2,6])
        %set(gca,'XTickLabel',{['Shallow Eup Zone'],'Deep Chl Max'})
        %set(gca,'XTickLabelRotation',20)
        ylabel('Net Primary Production (mmol N m^-^2 d^-^1)')
        set(gca,'FontSize',9)
        if Cycle==0
            keep.maxNPP = max(NPPs+NPPd);
        else
            keep.maxNPP = max([keep.maxNPP,max(NPPs+NPPd)]);
        end
        ylim([0 max([0,keep.maxNPP]) ])
        set(gca,'XTick',1)
        set(gca,'XTickLabel',{''})
        if Cycle==10
            AddPanelLabels('b','bottomright','YPADDING',0.02,'XPADDING',0.15,'FONTSIZE',9)
        end

    formatSpec =  'For Cycle %f NPP was %f +/- %f mmol N m-2 d-1 \n';
    fprintf(fileID,formatSpec,[Cycle, mean(NPPs+NPPd), std(NPPs+NPPd)])


        %-------------------------Ecosystem Efficiency--------------------------------
        subplot(1,numsubplots,4)
        hold on
        ViolinPlot(Cycle*0.1+1,0.5,HTLproduction./(NPPs+NPPd),200,col)

        set(gca,'box','on')
        %set(gca,'XTick',[2,6])
        %set(gca,'XTickLabel',{['Shallow Eup Zone'],'Deep Chl Max'})
        %set(gca,'XTickLabelRotation',20)
        ylabel('Ecosystem Efficiency')
        set(gca,'FontSize',9)
        ylim([0 0.16])
        set(gca,'XTick',[])
        set(gca,'XTickLabel',{''})
        if Cycle==10
            AddPanelLabels('c','bottomright','YPADDING',0.02,'XPADDING',0.15,'FONTSIZE',9)
        end


    formatSpec =  'For Cycle %f ecosystem efficiency was %f +/- %f \n';
    fprintf(fileID,formatSpec,[Cycle, mean(HTLproduction./(NPPs+NPPd)), std(HTLproduction./(NPPs+NPPd))])
    fclose(fileID)


        %-------------------------Bacteria GGE--------------------------------
        subplot(1,numsubplots/2,3)
        hold on
        x = 1 - (BacRespirations+BacRespirationd)./(BacDemands+BacDemandd);
        ViolinPlot(Cycle*0.1+1,0.5,x,200,col)

        if Cycle==0
            keep.maxBacGGE = max(x);
        else
            keep.maxBacGGE = max([keep.maxBacGGE,max(x)]);
        end
            


        %-------------------------Protist GGE--------------------------------
        subplot(1,numsubplots/2,3)
        hold on
        x = Protist_production./Protist_consumption;
        ViolinPlot(Cycle*0.1+6,0.5,x,200,col)

        if Cycle==0
            keep.maxProtistGGE = max(x);
        else
            keep.maxProtistGGE = max([keep.maxProtistGGE,max(x)]);
        end


        %-------------------------Mesozoo GGE--------------------------------
        subplot(1,numsubplots/2,3)
        hold on
        x = Mesozoo_production./Mesozoo_consumption;
        ViolinPlot(Cycle*0.1+11,0.5,x,200,col)

        set(gca,'box','on')
        %set(gca,'XTick',[2,6])
        %set(gca,'XTickLabel',{['Shallow Eup Zone'],'Deep Chl Max'})
        %set(gca,'XTickLabelRotation',20)
        ylabel(' ')
        set(gca,'FontSize',9)
        if Cycle==0
            keep.maxMesogooGGE = max(x);
        else
            keep.maxMesogooGGE = max([keep.maxMesogooGGE,max(x)]);
        end
        set(gca,'YTick',[0.1:0.1:0.4])
        ylim([0.1 0.4])
            
        set(gca,'XTick',[1:5:100])
        set(gca,'XTickLabel',{'Bacteria GGE','Protist GGE','Mesozoo GGE'})
        if Cycle==10
            AddPanelLabels('d','bottomright','YPADDING',0.02,'XPADDING',0.05,'FONTSIZE',9)
        end


        %-------------------------Large PhyNPP / Total NPP--------------------------------
        subplot(1,numsubplots/2,4)
        hold on
        x = (FLAGnpps + FLAGnppd + DTMnpps + DTMnppd)./(NPPs+NPPd)
        ViolinPlot(Cycle*0.1+0,0.5,x,200,col)

        if Cycle==0
            keep.maxNPPfrac = max(x);
        else
            keep.maxNPPfrac = max([keep.maxNPPfrac,max(x)]);
        end
            


        %-------------------------Herbivory--------------------------------

        subplot(1,numsubplots/2,4)
        hold on
        x = (HerbFoodChains + HerbFoodChaind)./(NPPs+NPPd)
        ViolinPlot(Cycle*0.1+5,0.5,x,200,col)
        set(gca,'box','on')
        %ylabel('Mean top trophic level')
        set(gca,'FontSize',9)
        set(gca,'XTick',[1:5:100])
        set(gca,'XTickLabel',{'Nano + Micro Phy NPP','Herbivory/NPP'})
        set(gca,'YTick',[0:0.1:0.6])
        if Cycle==10
            AddPanelLabels('e','bottomright','YPADDING',0.02,'XPADDING',0.05,'FONTSIZE',9)
        end
        


        %-------------------------Percentage of phytoplankton eaten--------------------------------

        if PlotPhyEatenPercent==1
            subplot(1,numsubplots,9)
            PercentageGPPconsumed = (NPP - PHY2DETRITUS)./NPP;
            x = PercentageGPPconsumed;
            ViolinPlot(Cycle*0.1,0.5,x,200,col)
            set(gca,'box','on')
            ylabel('Fraction of GPP Consumed')
            set(gca,'FontSize',9)
            set(gca,'XTick',[])
            AddPanelLabels('f','bottomright','YPADDING',0.02,'XPADDING',0.15,'FONTSIZE',9)
        end


        %-------------------------Trophic Level--------------------------------
        if PlotPhyEatenPercent==0
        subplot(1,numsubplots,9)
        elseif PlotPhyEatenPercent==1
        subplot(1,numsubplots,10)
        end
        hold on

        TL_mean = (TL_track(Plankfishs_col).*PLANKFISHproductions + ...
                   TL_track(Plankfishd_col).*PLANKFISHproductiond + ...
                   TL_track(GelPreds_col).*GELPREDproductions + ...
                   TL_track(GelPredd_col).*GELPREDproductiond + ...
                   TL_track(Preflexs_col).*PREFLEXproduction + ...
                   TL_track(Postflexs_col).*POSTFLEXproduction ) ./ ...
                   (PLANKFISHproductions + ...
                   PLANKFISHproductiond + ...
                   GELPREDproductions + ...
                   GELPREDproductiond + ...
                   PREFLEXproduction + ...
                   POSTFLEXproduction );

        x = TL_mean;
        ViolinPlot(Cycle*0.1,0.5,x,200,col)
        set(gca,'box','on')
        ylabel('Food chain length')
        set(gca,'FontSize',9)
        set(gca,'XTick',[])
        set(gca,'YTick',[3:0.2:5])
        set(gca,'YTickLabel',{'3','','','','','4','','','','','5'})
        ylim([3 5])
        if Cycle==10 & PlotPhyEatenPercent==0
            AddPanelLabels('f','bottomright','YPADDING',0.02,'XPADDING',0.15,'FONTSIZE',9)
        elseif Cycle==10 & PlotPhyEatenPercent==0
            AddPanelLabels('g','bottomright','YPADDING',0.02,'XPADDING',0.15,'FONTSIZE',9)
        end


        % if j==2 & Cycle==10
        %     subplot(1,2,1)
        %     AddPanelLabels('b','bottomright')
        %     AddPanelLabels('a','bottomright','YPADDING',0.05,'XPADDING',0.54)
        %     subplot(1,2,2)
        %     AddPanelLabels('d','bottomright')
        %     AddPanelLabels('c','bottomright','YPADDING',0.05,'XPADDING',0.54)
        % end
        %
        % if Cycle==10 & j==2
        %     fill([4.1 4.1 7.9 7.9 4.1],[2.75 3.95 3.95 2.75 2.75],'w')
        %     h=fill([4.2 4.2 4.4 4.4 4.2],[3.6 3.9 3.9 3.6 3.6],'w')
        %     set(h,'FaceColor',colVertex1)
        %     text(4.5,3.75,'Herbiv Food Chain','FontSize',8)
        %     h=fill([4.2 4.2 4.4 4.4 4.2],[3.5 3.2 3.2 3.5 3.5],'w')
        %     set(h,'FaceColor',colVertex2)
        %     text(4.5,3.35,'Multiv Food Chain','FontSize',8)
        %     h=fill([4.2 4.2 4.4 4.4 4.2],[3.1 2.8 2.8 3.1 3.1],'w')
        %     set(h,'FaceColor',colVertex3)
        %     text(4.5,2.95,'Microbial Loop','FontSize',8)
        % end

    end
    %breakitty
end


fn1 = ['SecondaryProductionDeterminants3']
exportgraphics(gcf,['../../Figures/',fn1,'.png'],'Resolution',600)
```
