## Supplementary material for "Comparative food-web analysis of bluefin tuna spawning habitats in the eastern Indian Ocean and Gulf of Mexico": Code Package: SetMatricesN15.m

```
clear all
close all

Cycle=10;
if Cycle==0  %BLOOFINZ-GoM Mean Values
    col=28;
elseif Cycle==10 %BLOOFINZ-IO Mean Values
    col=31;
else
    col = Cycle*2-1;
end

% first, get the name of spreadsheet with data
%sheet=['N15InverseModelRW.GOM-C5.xlsx']
if Cycle<=5
    sheet=['N15InverseModelRW.GOM.xlsx']
else
    sheet=['N15InverseModelRW.IO.xlsx']
end

workbook='Sheet1'

% now start reading in the data, starting with sizes
datsize=xlsread(sheet,workbook,'B2:E2');
neeq=datsize(1);   %Number of exact equalities
naeq=datsize(2);  %Number of approximate equalities
ngt0=datsize(3);  %Number of inequalities
nvar=datsize(4);  %Number of variables (flows)

ngt=ngt0+nvar;		% total no. ineqs, including >0
neq=neeq+naeq;      % total number of equalities

rskip=5;
cskip=5;
ccl=cskip+1;	% upper left number to be read
crl=rskip+1;
crr=rskip+nvar+1;
ccr=cskip+neq+ngt0+2;
crr2=rskip+nvar+2;
cread=[char(ExcelCol(ccl)),num2str(crl),':',char(ExcelCol(ccr)),num2str(crr)];
M0=xlsread(sheet,workbook,cread);
cread=[char(ExcelCol(ccl-1)),num2str(crl),':',char(ExcelCol(ccl-1)),num2str(crr-1)];
wts=xlsread(sheet,workbook,cread);

% % now, sort out
% % do some minimal prcocessing
M=[M0(:,1:(neq+ngt0)),eye(nvar+1,nvar)];
A=M(1:nvar,1:neq)';
G=M(1:nvar,neq+1:neq+ngt)';
b=M(nvar+1,1:neq)';
Ae=A(1:neeq,:);
Aa=A(neeq+1:neeq+naeq,:);
be=b(1:neeq);
ba=b(neeq+1:neeq+naeq);
sdba=NaN(size(ba));

Inputs=xlsread('Inputs.xlsx','Inputs','D3:AI46');

CN=106/16;
diff=2*10^-4*24*60*60;

%A(end,105)=Inputs(40,col)/1.2;  %POC:DOC vert grad ratio (assuming EqPac DOC vert grad from Carlson et al 1995)


ba(1)=Inputs(1,col);  %NPP (shallow)
ba(2)=Inputs(2,col);  %NPP (deep)
ba(3)=0;  %f-ratio mass balance (shallow)
ba(4)=0;  %f-ratio mass balance (deep)
frats = Inputs(3,col);
fratd = Inputs(4,col);
Aa(3,6) = frats - 1;
Aa(3,7) = frats;
Aa(3,15) = frats - 1;
Aa(3,16) = frats;
Aa(3,19) = frats - 1;
Aa(3,20) = frats;
Aa(3,30) = frats - 1;
Aa(3,31) = frats;
Aa(4,185) = fratd - 1;
Aa(4,186) = fratd;
Aa(4,194) = fratd - 1;
Aa(4,195) = fratd;
Aa(4,198) = fratd - 1;
Aa(4,199) = fratd;
Aa(4,209) = fratd - 1;
Aa(4,210) = fratd;
Aa0(3,6) = frats - 1;
Aa0(3,7) = frats;
Aa0(3,15) = frats - 1;
Aa0(3,16) = frats;
Aa0(3,19) = frats - 1;
Aa0(3,20) = frats;
Aa0(3,30) = frats - 1;
Aa0(3,31) = frats;
Aa0(4,185) = fratd - 1;
Aa0(4,186) = fratd;
Aa0(4,194) = fratd - 1;
Aa0(4,195) = fratd;
Aa0(4,198) = fratd - 1;
Aa0(4,199) = fratd;
Aa0(4,209) = fratd - 1;
Aa0(4,210) = fratd;
ba(5)=Inputs(43,col);  %Nfix (shallow)
ba(6)=Inputs(44,col);  %Nfix (deep)
ba(5+2)=Inputs(5,col);  %Total protistan grazing (shallow)
ba(6+2)=Inputs(6,col);  %Total protistan grazing (deep)
ba(7+2)=Inputs(7,col);  %Cyano NPP (shallow)
ba(8+2)=Inputs(8,col);  %Cyano NPP (deep)
ba(9+2)=Inputs(9,col);  %Flagellate NPP (shallow)
ba(10+2)=Inputs(10,col);  %Flagellate NPP (deep)
ba(11+2)=Inputs(11,col);  %Diatom NPP (shallow)
ba(12+2)=Inputs(12,col);    %Diatom NPP (deep)
ba(13+2)=Inputs(39,col);  %HBac Production (shallow)
ba(14+2)=Inputs(40,col);  %HBac Production (shallow)
ba(15+2)=Inputs(13,col);  %Cyano Mortality (shallow)
ba(16+2)=Inputs(14,col);  %Cyano Mortality (deep)
ba(17+2)=Inputs(15,col);  %Flagellate Mortality (shallow)
ba(18+2)=Inputs(16,col);  %Flagellate Mortality (deep)
ba(19+2)=Inputs(17,col);    %Diatom Mortality (shallow)
ba(20+2)=Inputs(18,col);   %Diatom Mortality (deep)
ba(21+2)=Inputs(41,col);  %HBac Mortality due to grazing (shallow)
ba(22+2)=Inputs(42,col);  %HBac Mortality due to grazing (shallow)
ba(23+2)=Inputs(19,col);   %NVM Mesozoo grazing
ba(24+2)=Inputs(20,col);   %VM Mesozoo grazing
ba(25+2)=Inputs(21,col);   %Sedtrap Flux (shallow)
ba(26+2)=Inputs(22,col);   %SedTrap Flux (deep)
ba(27+2)=Inputs(23,col);   %Thorium Flux (shallow)
ba(28+2)=Inputs(24,col);   %Thorium Flux (deep)
ba(29+2)=Inputs(25,col);   %Chl Sinking (shallow)
ba(30+2)=Inputs(26,col);   %Chl Sinking (deep)
ba(31+2)=Inputs(27,col);   %Fecal Pellet Sinking (shallow)
ba(32+2)=Inputs(28,col);   %Fecal Pellet Sinking (deep)
ba(33+2)=Inputs(29,col);   %Microzoo to Preflex
ba(34+2)=Inputs(30,col);   %Microzoo to Postflex
ba(35+2)=Inputs(31,col);   %Appendicularian to Preflex
ba(36+2)=Inputs(32,col);   %Appendicularian to Postflex
ba(37+2)=Inputs(33,col);   %Cladoceran to Preflex
ba(38+2)=Inputs(34,col);   %Cladoceran to Postflex
ba(39+2)=Inputs(35,col);   %Calanoids to Preflex
ba(40+2)=Inputs(36,col);   %Calanoids to Postflex
ba(41+2)=Inputs(37,col);   %Poecilostomatoids to Preflex
ba(42+2)=Inputs(38,col);   %Poecilostomatoids to Postflex

sdba(1)=Inputs(1,col+1);  %NPP (shallow)
sdba(2)=Inputs(2,col+1);  %NPP (deep)
%Note that the actual f-ratio equation is (f-1)NO3up + fNH4up, which is
%equal to 2(f-1)fNtot, so the uncertainty is 2f(f-1)Ntot*sqrt((sdf/f)^2+(sdf/f)^2+(sdNtot/Ntot)^2)
Ntot=Inputs(1,col);
sdNtot=Inputs(1,col+1);
sdfrat=Inputs(3,col+1);
sdba(3)=2*frats*(1-frats)*Ntot*sqrt( (sdfrat/frats)^2 + (sdfrat/frats)^2 + (sdNtot/Ntot)^2 );
Ntot=Inputs(2,col);
sdNtot=Inputs(2,col+1);
sdfrat=Inputs(4,col+1);
sdba(4)=2*fratd*(1-fratd)*Ntot*sqrt( (sdfrat/fratd)^2 + (sdfrat/fratd)^2 + (sdNtot/Ntot)^2 );
sdba(5)=Inputs(43,col+1);  %Nfix (shallow)
sdba(6)=Inputs(44,col+1);  %Nfix (deep)
sdba(5+2)=Inputs(5,col+1);  %NH4 uptake (shallow)
sdba(6+2)=Inputs(6,col+1);  %NH4 uptake (deep)
sdba(7+2)=Inputs(7,col+1);  %Cyano NPP (shallow)
sdba(8+2)=Inputs(8,col+1);  %Cyano NPP (deep)
sdba(9+2)=Inputs(9,col+1);  %Flagellate NPP (shallow)
sdba(10+2)=Inputs(10,col+1);  %Flagellate NPP (deep)
sdba(11+2)=Inputs(11,col+1);  %Diatom NPP (shallow)
sdba(12+2)=Inputs(12,col+1);    %Diatom NPP (deep)
sdba(13+2)=Inputs(39,col+1);  %HBac Production (shallow)
sdba(14+2)=Inputs(40,col+1);  %HBac Production (shallow)
sdba(15+2)=Inputs(13,col+1);  %Cyano Mortality (shallow)
sdba(16+2)=Inputs(14,col+1);  %Cyano Mortality (deep)
sdba(17+2)=Inputs(15,col+1);  %Flagellate Mortality (shallow)
sdba(18+2)=Inputs(16,col+1);  %Flagellate Mortality (deep)
sdba(19+2)=Inputs(17,col+1);    %Diatom Mortality (shallow)
sdba(20+2)=Inputs(18,col+1);   %Diatom Mortality (deep)
sdba(21+2)=Inputs(41,col+1);  %HBac Mortality due to grazing (shallow)
sdba(22+2)=Inputs(42,col+1);  %HBac Mortality due to grazing (shallow)
sdba(23+2)=Inputs(19,col+1);   %NVM Mesozoo grazing
sdba(24+2)=Inputs(20,col+1);   %VM Mesozoo grazing
sdba(25+2)=Inputs(21,col+1);   %Sedtrap Flux (shallow)
sdba(26+2)=Inputs(22,col+1);   %SedTrap Flux (deep)
sdba(27+2)=Inputs(23,col+1);   %Thorium Flux (shallow)
sdba(28+2)=Inputs(24,col+1);   %Thorium Flux (deep)
sdba(29+2)=Inputs(25,col+1);   %Chl Sinking (shallow)
sdba(30+2)=Inputs(26,col+1);   %Chl Sinking (deep)
sdba(31+2)=Inputs(27,col+1);   %Fecal Pellet Sinking (shallow)
sdba(32+2)=Inputs(28,col+1);   %Fecal Pellet Sinking (deep)
sdba(33+2)=Inputs(29,col+1);   %Microzoo to Preflex
sdba(34+2)=Inputs(30,col+1);   %Microzoo to Postflex
sdba(35+2)=Inputs(31,col+1);   %Appendicularian to Preflex
sdba(36+2)=Inputs(32,col+1);   %Appendicularian to Postflex
sdba(37+2)=Inputs(33,col+1);   %Cladoceran to Preflex
sdba(38+2)=Inputs(34,col+1);   %Cladoceran to Postflex
sdba(39+2)=Inputs(35,col+1);   %Calanoids to Preflex
sdba(40+2)=Inputs(36,col+1);   %Calanoids to Postflex
sdba(41+2)=Inputs(37,col+1);   %Poecilostomatoids to Preflex
sdba(42+2)=Inputs(38,col+1);   %Poecilostomatoids to Postflex
sdba(43+2:86+2)=NaN;
 
b=[be;ba];

InEqualInputs = xlsread('Inputs.xlsx','Inputs','D49:AI96');

h=M(nvar+1,neq+1:neq+ngt)';
Temp1=InEqualInputs(1,col);
Temp2=InEqualInputs(2,col);
Temp3=InEqualInputs(3,col);

HerbNVMbiomass=InEqualInputs(4,col);
Appbiomass=InEqualInputs(5,col);
Cladbiomass=InEqualInputs(6,col);
NVMCalbiomass=InEqualInputs(7,col);
Chaetobiomass=InEqualInputs(8,col);
Poecilbiomass=InEqualInputs(9,col);
GelPredbiomass=InEqualInputs(10,col);
Preflexbiomass=InEqualInputs(11,col);
Postflexbiomass=InEqualInputs(12,col);
HerbVMbiomass=InEqualInputs(13,col);
vmCalbiomass=InEqualInputs(14,col);
HerbNVMsize=InEqualInputs(34,col);
Appsize=InEqualInputs(35,col);
Cladsize=InEqualInputs(36,col);
NVMCalsize=InEqualInputs(37,col);
Chaetosize=InEqualInputs(38,col);
Poecilsize=InEqualInputs(39,col);
GelPredsize=InEqualInputs(40,col);
Preflexsize=InEqualInputs(41,col);
Postflexsize=InEqualInputs(42,col);
HerbVMsize=InEqualInputs(43,col);
vmCalsize=InEqualInputs(44,col);


Cyanobiomass=InEqualInputs(22,col);
Trichobiomass=InEqualInputs(23,col);
Diatombiomass=InEqualInputs(24,col);
Flagbiomass=InEqualInputs(25,col);
HNFbiomass=InEqualInputs(26,col);
MICbiomass=InEqualInputs(27,col);
Cyanobiomassd=InEqualInputs(28,col);
Trichobiomassd=InEqualInputs(29,col);
Diatombiomassd=InEqualInputs(30,col);
Flagbiomassd=InEqualInputs(31,col);
HNFbiomassd=InEqualInputs(32,col);
MICbiomassd=InEqualInputs(33,col);

%The following are minimum respiration from Ikeda (1985) - they
%parameterize ammonium excretion from carbon weight (in units of mg ind-1)
a0=-2.1763;
a1=0.8293;
a2=0.0648;
CN=6.625;
% CWsmall=0.0038;
% CWlarge=0.19;
CW=HerbNVMsize;
factor=24/14*exp(a0)*exp(a1*log(CW))*exp(a2*Temp1)/CW;  %Multiply by 24 hours divide by 14 g/mol divide by carbon weight, so this has units of umol N / day / mg C
h(25)=factor*(HerbNVMbiomass*CN*12)/1000;                       %Need to multiply by CN*12, because factor is in units of per mg carbon, while biomass is in units of mmol N.  Need to divide by 1000 to go from umol N / day to mmol N / day
CW=Appsize;
factor=24/14*exp(a0)*exp(a1*log(CW))*exp(a2*Temp1)/CW;  %Multiply by 24 hours divide by 14 g/mol
h(26)=factor*(Appbiomass*CN*12)/1000; 
CW=Cladsize;
factor=24/14*exp(a0)*exp(a1*log(CW))*exp(a2*Temp1)/CW;  %Multiply by 24 hours divide by 14 g/mol
h(27)=factor*(Cladbiomass*CN*12)/1000; 
CW=NVMCalsize;
factor=24/14*exp(a0)*exp(a1*log(CW))*exp(a2*Temp1)/CW;  %Multiply by 24 hours divide by 14 g/mol
h(28)=factor*(NVMCalbiomass*CN*12)/1000; 
CW=Chaetosize;
factor=24/14*exp(a0)*exp(a1*log(CW))*exp(a2*Temp1)/CW;  %Multiply by 24 hours divide by 14 g/mol
h(29)=factor*(Chaetobiomass*CN*12)/1000; 
CW=Poecilsize;
factor=24/14*exp(a0)*exp(a1*log(CW))*exp(a2*Temp1)/CW;  %Multiply by 24 hours divide by 14 g/mol
h(30)=factor*(Poecilbiomass*CN*12)/1000; 
CW=HerbVMsize;
factor=24/14*exp(a0)*exp(a1*log(CW))*exp(a2*Temp1)/CW;  %Multiply by 24 hours divide by 14 g/mol
h(33)=factor*(HerbVMbiomass*CN*12)/1000; 
CW=vmCalsize;
factor=24/14*exp(a0)*exp(a1*log(CW))*exp(a2*Temp1)/CW;  %Multiply by 24 hours divide by 14 g/mol
h(34)=factor*(vmCalbiomass*CN*12)/1000; 

%Constraining respiration to be higher in the surface than at depth
herbvmexc_col=41;
vmCalexc_col=42;
herbvmTOsnh4 = 145;
herbvmTOdnh4 = 152;
herbvmTOexportnh4 = 155;
vmCalTOsnh4 = 163;
vmCalTOdnh4 = 170;
vmCalTOexportnh4 = 173;

G(herbvmexc_col,herbvmTOsnh4)=exp(a2*Temp3);
G(herbvmexc_col,herbvmTOdnh4)=exp(a2*Temp3);
G(herbvmexc_col,herbvmTOexportnh4)=-exp(a2*Temp1);
G(vmCalexc_col,vmCalTOsnh4)=exp(a2*Temp3);
G(vmCalexc_col,vmCalTOdnh4)=exp(a2*Temp3);
G(vmCalexc_col,vmCalTOexportnh4)=-exp(a2*Temp1);

herbvmexcdom_col=109;
vmCalexcdom_col=110;
herbvmTOsdom = 146;
herbvmTOddom = 153;
herbvmTOexportdom = 156;
vmCalTOsdom = 164;
vmCalTOddom = 171;
vmCalTOexportdom = 174;

G(herbvmexcdom_col,herbvmTOsdom)=exp(a2*Temp3);
G(herbvmexcdom_col,herbvmTOddom)=exp(a2*Temp3);
G(herbvmexcdom_col,herbvmTOexportdom)=-exp(a2*Temp1);
G(vmCalexcdom_col,vmCalTOsdom)=exp(a2*Temp3);
G(vmCalexcdom_col,vmCalTOddom)=exp(a2*Temp3);
G(vmCalexcdom_col,vmCalTOexportdom)=-exp(a2*Temp1);


AllPhytoNPPs = Inputs(1,col);
CyanoNPPs = Inputs(7,col);
DtmNPPs = Inputs(11,col);
FlagNPPs = Inputs(9,col);
AllPhytoNPPd = Inputs(1,col);
CyanoNPPd = Inputs(7,col);
DtmNPPd = Inputs(11,col);
FlagNPPd = Inputs(9,col);

% h(43)=0.02*AllPhytoNPPs;
% h(51)=-0.55*AllPhytoNPPs;
% h(44)=0.02*CyanoNPPs;
% h(52)=-0.55*CyanoNPPs;
% h(45)=0.02*DtmNPPs;
% h(53)=-0.55*DtmNPPs;
% h(46)=0.02*FlagNPPs;
% h(54)=-0.55*FlagNPPs;
% h(47)=0.02*AllPhytoNPPd;
% h(55)=-0.55*AllPhytoNPPd;
% h(48)=0.02*CyanoNPPd;
% h(56)=-0.55*CyanoNPPd;
% h(49)=0.02*DtmNPPd;
% h(57)=-0.55*DtmNPPd;
% h(40)=0.02*FlagNPPd;
% h(58)=-0.55*FlagNPPd;

%The following are max ingestion constraints from Hansen
a6 = 0.1+0.24;  %THis is the value for protozooplankton from Hansen et al (1997, p. 11) plus one s.e.
b6 = -0.20;   %This is the value for protozooplankton from Hansen et al. (1997, p. 11) )

%Shallow
%Mixotrophic Flagellates
PredVol=1413.7;  %um^3
h(215)=-24*10^(a6+b6*log10(PredVol)+log10(2.8)*(Temp1-20)/10)*Flagbiomass;
%Heterotrophic Nanoflagellates
PredVol=14.1;
h(216)=-24*10^(a6+b6*log10(PredVol)+log10(2.8)*(Temp1-20)/10)*HNFbiomass;
%Microzoos
PredVol=1413.7;
h(217)=-24*10^(a6+b6*log10(PredVol)+log10(2.8)*(Temp1-20)/10)*MICbiomass;

a6 = -1.78+0.47;  %THis is the value for metazooplankton from Hansen et al (1997, p. 11) plus one s.e.
b6 = 0.08;   %This is the value for metazooplankton from Hansen et al. (1997, p. 11))
%HerbNVM
DWC=0.4; %Dry weight to carbon conversion
WWDW = 0.2;  %Wet weight to Dryweight conversion
PredVol=HerbNVMsize/DWC/WWDW*10^9;   %10^9 goes from mm^3 to um^3
h(218)=-24*10^(a6+b6*log10(PredVol)+log10(2.8)*(Temp1-20)/10)*HerbNVMbiomass;
%Appendicularians
DWC=0.4; %Dry weight to carbon conversion
WWDW = 0.2;  %Wet weight to Dryweight conversion
PredVol=Appsize/DWC/WWDW*10^9;   %10^9 goes from mm^3 to um^3
h(219)=-24*10^(a6+b6*log10(PredVol)+log10(2.8)*(Temp1-20)/10)*Appbiomass;
%Cladocerans
DWC=0.4; %Dry weight to carbon conversion
WWDW = 0.2;  %Wet weight to Dryweight conversion
PredVol=Cladsize/DWC/WWDW*10^9;   %10^9 goes from mm^3 to um^3
h(220)=-24*10^(a6+b6*log10(PredVol)+log10(2.8)*(Temp1-20)/10)*Cladbiomass;
%nvmCal
DWC=0.4; %Dry weight to carbon conversion
WWDW = 0.2;  %Wet weight to Dryweight conversion
PredVol=NVMCalsize/DWC/WWDW*10^9;   %10^9 goes from mm^3 to um^3
h(221)=-24*10^(a6+b6*log10(PredVol)+log10(2.8)*(Temp1-20)/10)*NVMCalbiomass;
%Chaeto
DWC=0.4; %Dry weight to carbon conversion
WWDW = 0.05;  %Wet weight to Dryweight conversion
PredVol=Chaetosize/DWC/WWDW*10^9;   %10^9 goes from mm^3 to um^3
h(222)=-24*10^(a6+b6*log10(PredVol)+log10(2.8)*(Temp1-20)/10)*Chaetobiomass;
%Poecil
DWC=0.4; %Dry weight to carbon conversion
WWDW = 0.2;  %Wet weight to Dryweight conversion
PredVol=Poecilsize/DWC/WWDW*10^9;   %10^9 goes from mm^3 to um^3
h(223)=-24*10^(a6+b6*log10(PredVol)+log10(2.8)*(Temp1-20)/10)*Poecilbiomass;
%Gelatinous Predators
DWC=0.4; %Dry weight to carbon conversion
WWDW = 0.2;  %Wet weight to Dryweight conversion
PredVol=GelPredsize/DWC/WWDW*10^9;   %10^9 goes from mm^3 to um^3
h(224)=-24*10^(a6+b6*log10(PredVol)+log10(2.8)*(Temp1-20)/10)*GelPredbiomass;
%Preflex
DWC=0.4; %Dry weight to carbon conversion
WWDW = 0.2;  %Wet weight to Dryweight conversion
PredVol=Preflexsize/DWC/WWDW*10^9;   %10^9 goes from mm^3 to um^3
h(226)=-24*10^(a6+b6*log10(PredVol)+log10(2.8)*(Temp1-20)/10)*Preflexbiomass;
%Postflex
DWC=0.4; %Dry weight to carbon conversion
WWDW = 0.2;  %Wet weight to Dryweight conversion
PredVol=Postflexsize/DWC/WWDW*10^9;   %10^9 goes from mm^3 to um^3
h(227)=-24*10^(a6+b6*log10(PredVol)+log10(2.8)*(Temp1-20)/10)*Postflexbiomass;
%Vertical Migrators
%herbvm
DWC=0.4; %Dry weight to carbon conversion
WWDW = 0.2;  %Wet weight to Dryweight conversion
PredVol=HerbVMsize/DWC/WWDW*10^9;   %10^9 goes from mm^3 to um^3
h(228)=-24*10^(a6+b6*log10(PredVol)+log10(2.8)*(Temp1-20)/10)*HerbVMbiomass;
%vmCal
DWC=0.4; %Dry weight to carbon conversion
WWDW = 0.2;  %Wet weight to Dryweight conversion
PredVol=vmCalsize/DWC/WWDW*10^9;   %10^9 goes from mm^3 to um^3
h(229)=-24*10^(a6+b6*log10(PredVol)+log10(2.8)*(Temp1-20)/10)*vmCalbiomass;
%Deep Euphotic


h(241)=-2*Cyanobiomass;  %Cyano max production at double biomass
h(242)=-0.24*Trichobiomass;  %Tricho max production at double biomass   %From Mulholland et al. (2006)
h(243)=-2*Diatombiomass;  %Diatom max production at double biomass
h(244)=-2*Flagbiomass;  %Flag max production at double biomass
h(245)=-2*Cyanobiomassd;  %Cyano max production at double biomass
h(246)=-0.24*Trichobiomassd;  %Tricho max production    %From Mulholland et al. (2006)
h(247)=-2*Diatombiomassd;  %Diatom max production at double biomass
h(248)=-2*Flagbiomassd;  %Flag max production at double biomass

MaxUpwellings = InEqualInputs(45,col);
MaxUpwellingd = InEqualInputs(46,col);
%Next is mixing
h(249)=-MaxUpwellings;
h(250)=-MaxUpwellingd;

%Next is maximum Tricho N2fixation
h(251)=-0.15*Trichobiomass;  %Tricho max production at double biomass   %From Mulholland et al. (2006)
h(252)=-0.15*Trichobiomassd;  %Tricho max production at double biomass   %From Mulholland et al. (2006)

MaxLateralPON = InEqualInputs(47,col);
MaxLateralDON = InEqualInputs(48,col);
%Next is lateral input
h(253)=-MaxLateralPON;
h(254)=-MaxLateralDON;


inds=find(isnan(h));
G(inds,:)=[];
h(inds)=[];
inds=find(isnan(ba));
Aa(inds,:)=[];
ba(inds)=[];
sdba(inds)=[];


A0 = A;
G0 = G;
Ae0=Ae;
Aa0=Aa;
for i=1:length(wts)
    A(:,i)=A(:,i)/wts(i);
    Aa(:,i)=Aa(:,i)/wts(i);
    Ae(:,i)=Ae(:,i)/wts(i);
    G(:,i)=G(:,i)/wts(i);
end


d15NInputs=xlsread('Inputs.xlsx','Inputs','D99:AI114');


%Just clearing variables that we don't need to see
clear Araw
clear Awt
clear Graw
clear M
clear M0
clear captions
clear cc2
clear cc3
clear ccl
clear ccr
clear chsheet
clear comment
clear constraintout
clear constraints
clear cread
clear cread2
clear cread3
clear cread4
clear crl
clear crr
clear crr2
clear crr3
clear crr4
clear cskip
clear datsize
clear equalities
clear equalitiesout
clear linsum
clear neq
clear naeq
clear neeq
clear ngt
clear ngt0
clear norm2
clear npar
clear nvar
clear outgate
clear rc1
clear cnorm
clear rr
clear rrout
clear rrtrans
clear rskip
clear sheet
clear sols
clear sqrterror
clear temp
clear time1
clear weight
clear SD


InputCol = col;

if length(ba)>0
    save(['N15GoMInverseCycle',num2str(Cycle),'.mat'],'A','Ae','Aa','G','b','be','ba','h','Inputs','sdba','InputCol','d15NInputs','wts','A0','Ae0','Aa0','G0')
else
    save([sheetp,'.',Model,'.mat'],'A','G','b','h','wts')
end
```
