## Supplementary material for "Comparative food-web analysis of bluefin tuna spawning habitats in the eastern Indian Ocean and Gulf of Mexico": Code Package: TernaryPlot.m

```
function [outputs]=TernaryPlot(Vertex1,Vertex2,Vertex3,LABELS,TYPE,DotColor,MarkSize,LargeFont,SmallFont,overlay,translucency)

%Typiceal call: TernaryPlot(Vertex1,Vertex2,Vertex3,Labels,'Circle',[1 0 0],10,12,9)

% clearvars
% close all
% Vertex1 = rand(10,1)+2;
% Vertex2 = rand(10,1)+0.5;
% Vertex3 = rand(10,1);
% LargeFont = 12;
% SmallFont = 9;
% MarkSize = 10;
% DotColor = [1 0 0];
% TYPE = 'Circle';  %Options are 'Circle', 'Dot', 'Histogram'

colVertex1 = [0 0.6 0];
colVertex2 = [0.7 0 1];
colVertex3 = [0.4 0.3 0];


plot([-1/sqrt(2) 0 1/sqrt(2) -1/sqrt(2)],[0 1 0 0],'-k','LineWidth',2)
hold on
for i=0.2:0.2:0.8
    Vertex3fracRemaining=-0.15*i:0.01:1;
    PerVertex1=i*ones(size(Vertex3fracRemaining));
    NonVertex1=1-PerVertex1;
    x = (1 - Vertex3fracRemaining*2) .* NonVertex1 / sqrt(2);
    plot(x,PerVertex1,'-b','Color',colVertex1,'LineWidth',2)
    text(max(x)+0.01,i+0.01,[num2str(i*100),'%'],'Color',colVertex1,'FontSize',SmallFont)
end
text(0,1.03,LABELS{1},'HorizontalAlignment','center','FontSize',LargeFont,'Color',colVertex1)
for i=0.2:0.2:0.8
    PerVertex1=0:0.01:1-i+0.03;
    NonVertex1=1-PerVertex1;
    Vertex3fracRemaining=i./(1-PerVertex1);
    x = (1 - Vertex3fracRemaining*2) .* NonVertex1 / sqrt(2);
    plot(x,PerVertex1,'-m','Color',colVertex3,'LineWidth',2)
    text(min(x)-0.03,max(PerVertex1)+0.02,[num2str(i*100),'%'],'HorizontalAlignment','center','Color',colVertex3,'FontSize',SmallFont)
end
text(-1/sqrt(2),-0.08,LABELS{3},'HorizontalAlignment','center','FontSize',LargeFont,'Color',colVertex3)
for i=0.2:0.2:0.8
    PerVertex1=-0.02:0.01:1-i;
    NonVertex1=1-PerVertex1;
    Vertex3fracRemaining=(1-i-PerVertex1)./(1-PerVertex1);
    x = (1 - Vertex3fracRemaining*2) .* NonVertex1 / sqrt(2);
    plot(x,PerVertex1,'-r','Color',colVertex2,'LineWidth',2)
    text(min(x)-0.01,-0.04,[num2str(i*100),'%'],'HorizontalAlignment','center','Color',colVertex2,'FontSize',SmallFont)
end
text(1/sqrt(2),-0.03,LABELS{2},'HorizontalAlignment','center','FontSize',LargeFont,'Color',colVertex2)
set(gca,'box','off')
set(gca,'xcolor',[1 1 1])
set(gca,'ycolor',[1 1 1])
set(gcf,'Color','w')
set(gca,'XTick',[])
set(gca,'Ytick',[])
xlim([-1/sqrt(2) 1/sqrt(2)])


%------------------------------------------------------------------------
%------------- First Figure (not histogram) -----------------------------
%------------------------------------------------------------------------


Total = Vertex1 + Vertex3 + Vertex2;
PerVertex3 = Vertex3./Total
PerVertex2 = Vertex2./Total
PerVertex1 = Vertex1./Total
PerVertex3=reshape(PerVertex3,length(PerVertex3(:,1))*length(PerVertex3(1,:)),1);
PerVertex2=reshape(PerVertex2,length(PerVertex2(:,1))*length(PerVertex2(1,:)),1);
PerVertex1=reshape(PerVertex1,length(PerVertex1(:,1))*length(PerVertex1(1,:)),1);
NonVertex1=1-PerVertex1;
Vertex3fracRemaining = PerVertex3./(PerVertex3+PerVertex2);
x = (1 - Vertex3fracRemaining*2) .* NonVertex1 / sqrt(2);
Vertex2=reshape(Vertex2,length(Vertex2(:,1))*length(Vertex2(1,:)),1);
if strcmp(TYPE,'Dot')
    if translucency==1
        plot(x,PerVertex1,'.r','Color',DotColor)
    else
        h = scatter(x,PerVertex1,MarkSize,'o','filled','MarkerFaceColor',DotColor,'MarkerEdgeColor',DotColor,'MarkerFaceAlpha',translucency,'MarkerEdgeAlpha',0);
    end
elseif strcmp(TYPE,'Circle')
    plot(x,PerVertex1,'ok','MarkerFaceColor',DotColor,'MarkerEdgeColor','k','MarkerSize',MarkSize)
elseif strcmp(TYPE,'Square')
    plot(x,PerVertex1,'sk','MarkerFaceColor',DotColor,'MarkerEdgeColor','k','MarkerSize',MarkSize)
elseif strcmp(TYPE,'Diamond')
    plot(x,PerVertex1,'dk','MarkerFaceColor',DotColor,'MarkerEdgeColor','k','MarkerSize',MarkSize)
elseif strcmp(TYPE,'Hexagram')
    plot(x,PerVertex1,'hexagram','MarkerFaceColor',DotColor,'MarkerEdgeColor','k','MarkerSize',MarkSize)
elseif strcmp(TYPE,'Histogram')
    len=length(x);
    data = [x,PerVertex1];
    [N,c]=hist3(data,[99,100]);
    N(find(N==0))=NaN;
    N = N/len;
    pcolor(c{1,1},c{1,2},log10(N'));
    shading interp
    h=colorbar('southoutside');
    set(h,'YTick',log10([1:10,20:10:100,200:100:1000,2000:1000:10^4,2*10^4:10^4:10^5,2*10^5:10^5:10^6,2*10^6:10^6:10^7,2*10^7:10^7:10^8]/10^8));
    set(h,'YTickLabel',{'10^-^8','','','','','','','','','10^-^7','','','','','','','','','10^-^6',...
        '','','','','','','','','10^-^5','','','','','','','','','10^-^4',...
        '','','','','','','','','10^-^3','','','','','','','','','10^-^2',...
        '','','','','','','','','10^-^1','','','','','','','','','1'});
end


if overlay==1
    plot([-1/sqrt(2) 0 1/sqrt(2) -1/sqrt(2)],[0 1 0 0],'-k','LineWidth',2)
    hold on
    for i=0.2:0.2:0.8
        Vertex3fracRemaining=-0.15*i:0.01:1;
        PerVertex1=i*ones(size(Vertex3fracRemaining));
        NonVertex1=1-PerVertex1;
        x = (1 - Vertex3fracRemaining*2) .* NonVertex1 / sqrt(2);
        plot(x,PerVertex1,'-b','Color',colVertex1,'LineWidth',2)
        text(max(x)+0.01,i+0.01,[num2str(i*100),'%'],'Color',colVertex1,'FontSize',SmallFont)
    end
    %text(0,1.03,LABELS{1},'HorizontalAlignment','center','FontSize',LargeFont,'Color',colVertex1)
    for i=0.2:0.2:0.8
        PerVertex1=0:0.01:1-i+0.03;
        NonVertex1=1-PerVertex1;
        Vertex3fracRemaining=i./(1-PerVertex1);
        x = (1 - Vertex3fracRemaining*2) .* NonVertex1 / sqrt(2);
        plot(x,PerVertex1,'-m','Color',colVertex3,'LineWidth',2)
        text(min(x)-0.03,max(PerVertex1)+0.02,[num2str(i*100),'%'],'HorizontalAlignment','center','Color',colVertex3,'FontSize',SmallFont)
    end
    %text(-1/sqrt(2),-0.08,LABELS{3},'HorizontalAlignment','center','FontSize',LargeFont,'Color',colVertex3)
    for i=0.2:0.2:0.8
        PerVertex1=-0.02:0.01:1-i;
        NonVertex1=1-PerVertex1;
        Vertex3fracRemaining=(1-i-PerVertex1)./(1-PerVertex1);
        x = (1 - Vertex3fracRemaining*2) .* NonVertex1 / sqrt(2);
        plot(x,PerVertex1,'-r','Color',colVertex2,'LineWidth',2)
        text(min(x)-0.01,-0.04,[num2str(i*100),'%'],'HorizontalAlignment','center','Color',colVertex2,'FontSize',SmallFont)
    end
    %text(1/sqrt(2),-0.03,LABELS{2},'HorizontalAlignment','center','FontSize',LargeFont,'Color',colVertex2)
end
```
