## Supplementary material for "Comparative food-web analysis of bluefin tuna spawning habitats in the eastern Indian Ocean and Gulf of Mexico": Code Package: TurnOffWarnings.m

```
%Query to find last warning is: warning('query','last')

id = 'MATLAB:table:ModifiedAndSavedVarnames';
warning('off',id);
id = 'MATLAB:table:RowsAddedExistingVars';
warning('off',id);
id = 'MATLAB:scatteredInterpolant:DupPtsAvValuesWarnId';
warning('off',id);
id = 'MATLAB:griddata:DuplicateDataPoints';
warning('off',id);
id = 'MATLAB:interp1:NaNstrip';
warning('off',id);
```
