## Supplementary material for "Comparative food-web analysis of bluefin tuna spawning habitats in the eastern Indian Ocean and Gulf of Mexico": Code Package: ViolinPlot.m

```
function [output]  = ViolinPlot(x,wid,data,num,col)

% clearvars
% x=1;
% wid=0.5;
% num=300;
% load('E:\Stukel Work\Science\OneDrive - Florida State University\GoM Nitrogen & Trophic - NOAA RESTORE\Inverse Model\R Code\Test 18\N15GoMInverseCycle1Routputs.mat')
% data=MCMCmat(:,22);
% col=[1 0 0];

ymin=min(data)
ymax=max(data)
for i=1:num
    lowerlim=ymin+(ymax-ymin)/num*(i-1);
    upperlim=ymin+(ymax-ymin)/num*i;
    yint(i)=(lowerlim+upperlim)/2;
    ind=find(data>lowerlim & data<upperlim);
    histog(i)=length(ind);
end
thick=histog*wid/max(histog);

plot([x,x],[ymin,ymax],'-k','Color',col)
hold on
% for i=1:length(thick)
%     plot([x-thick(i),x+thick(i)],[yint(i) yint(i)],'-k','Color',col)
% end
h=fill([thick,fliplr(-thick)]+x,[yint,fliplr(yint)],'r');
set(h,'LineStyle','none')
set(h,'FaceColor',col)
```
