## Supplementary material for "Comparative food-web analysis of bluefin tuna spawning habitats in the eastern Indian Ocean and Gulf of Mexico": Code Package: xsampleN15.r

```
##==============================================================================
## xsample  : Samples linear problem with equality and inequality constraints ##
##==============================================================================

xsampleN15 <- function(A=NULL, B=NULL, E=NULL, F=NULL, G=NULL, H=NULL,
                     sdB=NULL, W=1, iter=3000, outputlength = iter,
                     burninlength = NULL, type="mirror", jmp=NULL, jmp2=NULL,
                     tol=sqrt(.Machine$double.eps), x0=NULL, del15N1=NULL,
					 del15Nknown=NULL, HFPFfrac=NULL, PPFfrac=NULL,
                     fulloutput=FALSE, test=TRUE)   {
  
  #####   1. definition of internal functions    #####
  
  ## function ensuring that a jump from q1 to q2
  ## fulfills all inequality constraints formulated in g and h
  ## gq=h can be seen as equations for planes that are considered mirrors.
  ## when a jump crosses one or more of these mirrors,
  ## the vector describing the jump is deviated according to rules of mirroring.
  ## the resulting new vector q will always be within the subspace of R^n for
  ## which all inequalities are met.
  ## also are the requirements for a MCMC met: the probability in the subspace
  ## is constant, the probability out of the subspace is 0.
  ## q1 has to fulfill constraints by default!
  ## Karel Van den Meersche 20070921
  
  
  ## Stukel adding a function to allow delN15 to vary uniformly
  
  new15N <- function(d15N,jmp2){
    l15N <- length(d15N)
    d15N <- d15N + (runif(l15N)-0.5)*jmp2
  }
  
  
  mirror <- function(q1,g,h,k=length(q),jmp)   {
    
    ##if (any((g%*%q1)<h)) stop("starting point of mirroring is not in feasible space")
    q2 <- rnorm(k,q1,jmp)
    if (!is.null(g))    {
      
      residual <- g%*%q2-h
      q10 <- q1
      
      while (any(residual<0))  {              #mirror
        epsilon <- q2-q10                     #vector from q1 to q2: our considered light-ray that will be mirrored at the boundaries of the space
        w <- which(residual<0)                #which mirrors are hit?
        alfa <- ((h-g%*%q10)/g%*%epsilon)[w]  #alfa: at which point does the light-ray hit the mirrors? g*(q1+alfa*epsilon)-h=0
        whichminalfa <- which.min(alfa)
        j <- w[whichminalfa]                  #which smallest element of alfa: which mirror is hit first?
        d <- -residual[j]/sum(g[j,]^2)        #add to q2 a vector d*Z[j,] which is oriented perpendicular to the plane Z[j,]%*%x+p; the result is in the plane.
        q2 <- q2+2*d*g[j,]                    #mirrored point
        residual <- g%*%q2-h
        q10 <- q10+alfa[whichminalfa]*epsilon #point of reflection
      }
    }
    q2
  }
  
  ## hit-and-run algorithms
  ## modeled after Smith, R.L. Efficient Monte Carlo Procedures for
  ## Generating Points Uniformly Distributed over Bounded Regions. Operations Research 32, pp. 1296-1308,1984.
  
  ## 1. coordinates direction algorithm
  
  cda <- function(q,g,h,k=length(q),...)    {
    # samples a new point in the feasible range of the solution space
    # along a random coordinate
    
    i <- sample(1:k,1)                  #
    h1 <- h-as.matrix(g[,-i])%*%q[-i]              # g[,i]q[i]>h1
    maxqi <- min((h1/g[,i])[g[,i]<0])
    minqi <- max((h1/g[,i])[g[,i]>0])
    q[i]  <-  runif(1,minqi,maxqi)
    return(q)
  }
  
  ## 2. random direction algorithm
  rda <- function(q,g,h,k=length(q),...)   {
    #samples a new point in the feasible range of the solution space
    #in a random direction
    ##if (any((g%*%q)<h)) stop("starting point is not in feasible space")
    e <- rnorm(k)
    d <- e/norm(e)                        #d: random direction vector; q2 = q + alfa*d
    
    alfa <- ((h-g%*%q)/g%*%d)             #
    if (any(alfa>0)) alfa.u <- min(alfa[alfa>0]) else alfa.u <- 0
    if (any(alfa<0)) alfa.l <- max(alfa[alfa<0]) else alfa.l <- 0
    
    q.u <- q+alfa.u*d
    q.l <- q+alfa.l*d
    
    if (any(g%*%q.u<h)) alfa.u <- 0
    if (any(g%*%q.l<h)) alfa.l <- 0
    q <- q+runif(1,alfa.l,alfa.u)*d
    return(q)
  }
  
  norm <- function(x) sqrt(x%*%x)
  
  automatedjump <- function(a,b,g,h,g.scale=5,a.scale=1)   {
    if (is.null(g)) s1 <- rep(NA,k)
    else  {
      q_ranges <- xranges(E=NULL,F=NULL,g,h)
      s1 <- abs(q_ranges[,1]-q_ranges[,2])/g.scale
    }
    s2 <- rep(NA,k)
    if (!is.null(A)) {
      if (estimate_sdB) {
        estVar <- SS0/(lb-lx)*solve(t(a)%*%a) # estimated variance on the parameters, simplified from Brun et al 2001
        estSd  <- sqrt(diag(estVar))
        s2 <- estSd/a.scale
      }
      if (qr(A)$rank==lx&lx==lb)
        s2 <- sdB*.2
    }
    s <- pmin(s1,s2,na.rm=T)
    s[s>tol^-2] <- NA
    if (any (is.na(s)))  {
      if (all(is.na(s)))  {
        warning(" problem is unbounded - all jump lengths are set to 1")
        s[] <- 1
      } else {
        warning(" problem is unbounded - some jump lengths are set arbitrarily")
        s[is.na(s)] <- mean(s,na.rm=T)*100
      }
    }
    return(s)
  }
  
  
  #### 2. the xsample function ####
  
  ## conversions vectors to matrices and checks
  if (is.data.frame(A)) A <- as.matrix(A)
  if (is.data.frame(E)) E <- as.matrix(E)
  if (is.data.frame(G)) G <- as.matrix(G)
  if (is.vector(A)) A <- t(A)
  if (is.vector(E)) E <- t(E)
  if (is.vector(G)) G <- t(G)
  
  

  del15NupNO3 <- del15Nknown[1]
  del15NPreflexs <- del15Nknown[2]
  del15NPostflexs <- del15Nknown[3]
  del15NLdets <- del15Nknown[4]
  del15NLdetd <- del15Nknown[5]
  del15NDONs <- del15Nknown[6]
  del15NDONd <- del15Nknown[7]
  
  
  if ( !is.null(A) )   {
    lb <- length(B)
    lx <- ncol(A)
    ## system overdetermined?
	  A2 <- ResetRN15(Aa,x0,del15N1,del15NupNO3,del15NPreflexs,del15NPostflexs,del15NLdets,del15NLdetd,del15NDONs,del15NDONd)
    M <- rbind(cbind(A2,B),cbind(E,F))
    overdetermined <- !qr(M)$rank<=lx
    
    if (overdetermined & is.null(sdB)) {
      warning("The given linear problem is overdetermined. A standard deviation for the data vector B is incorporated in the MCMC as a model parameter.")
      estimate_sdB=TRUE
      A <- A*W
      B <- B*W
    } else {
      estimate_sdB=FALSE
      if (overdetermined)
        warning("The given linear problem is overdetermined. Giving fixed standard deviations for the data vector B can lead to dubious results. Maybe you want to set sdB=NULL and estimate the data error.")
      if (!length(sdB)%in%c(1,lb))
        stop("sdB does not have the correct length")
#      A <- A/sdB
#      B <- B/sdB           # set sd = 1 in Ax = N(B,sd)
      
    }
  } else {  #is.null A
    estimate_sdB=FALSE
  }
  
  ## find a particular solution x0
  if (is.null(x0))  {
    l <- lsei(A=A,B=B,E=E,F=F,G=G,H=H,type=2)
    if (l$residualNorm>1e-6)
      stop("no particular solution found;incompatible constraints")
    else
      x0 <- l$X
  }
  lx <- length(x0)
  
  ## additional checks for equalities, hidden in inequalities... (Karline S.)
  if (test && !is.null(G))   {
    xv <- varranges(E,F,G,H,EqA=G)
    ii <- which (xv[,1]-xv[,2]==0)
    if (length(ii)>0) { # if they exist: add regular equalities !
      E  <- rbind(E,G[ii,])
      F  <- c(F,xv[ii,1])
      
      G  <- G[-ii,]
      H  <- H[-ii]
      if (length(H)==0)
        G <- H <- NULL
    }
    xr <- xranges(E,F,G,H)
    ii <- which (xr[,1]-xr[,2]==0)
    if (length(ii)>0) { # if they exist: add regular equalities !
      dia <- diag(nrow=nrow(xr))
      E  <- rbind(E,dia[ii,])
      F  <- c(F,xr[ii,1])
    }
  }
  
  ## Z is an orthogonal matrix for which E%*%Z=0;
  ## it can serve as basis for the null space of E.
  ## all solutions for the equalities have the form x = x0 + Zq
  ## with q a random vector.
  ## the vector q is varied in a random walk, using a MCMC with
  ## acceptance rate = 1. The inequality constraints Gx>H
  ## can be rewritten as Gx0-H + (GZ)q >0
  
  if (!is.null(E))  {
    Z <- Null(t(E)); Z[abs(Z)<tol] <- 0  #x=x0+Zq ; EZ=0
  } else { Z <- diag(lx) }
  
  if (length(Z)==0)  {
    warning("the problem has a single solution; this solution is returned as function value")
    return(x0)
  }
  k <- ncol(Z)
  
  if (!is.null(G))   {
    g <- G%*%Z
    h <- H-G%*%x0                                            #gq-h>=0
    g[abs(g)<tol] <- 0
    h[abs(h)<tol] <- 0
    
  } else { g <- G; h <- H }
  
  
  if (!is.null(A))   {
    # a <- A%*%Z
    # b <- B-A%*%x0                          #aq-b~=0
    # v <- svd(a,nv=k)$v                     #transformation q <- t(v)q for better convergence
    # a <- a%*%v                             #transformation a <- av
    # if (!is.null(G)) g <- g%*%v            #transformation g <- gv
    # Z <- Z%*%v                             #transformation Z <- Zv
    # 
    # ## if overdetermined, calculate posterior distribution of S in Ax=N(B,S)
    # ## Marko Laine 2008, thesis on adaptive mcmc
    # ## S = 1/sd^2 of model
    # ## prior n0=lb
    # ## prior SSR0=n0*s0^2=sum((Ax0-B)^2)=sum(b^2)
    # ## if underdetermined: S=1
    # ## if overdetermined: S is sampled from a
    # ## posterior gamma distribution (Laine 2008) and
    # ## standard deviations of data are S^-.5
    if (estimate_sdB)    {
      q0 <- lsei(a,b)$X
      SS0 <- sum((a%*%q0-b)^2)
      S <- lb/SS0
    } else {
      S <- 1
    }
    
    # sum of squared residuals
    SSR <- function(q,d15N,Aa,Ba,sdB){ 
      Xa <- x0+Z%*%q
      Aa <- ResetRN15(Aa,Xa,d15N,del15NupNO3,del15NPreflexs,del15NPostflexs,del15NLdets,del15NLdetd,del15NDONs,del15NDONd)
      ssr <- sum(((Aa%*%Xa-Ba)/sdB)^2)
      return(ssr)
    }	  
    
    ##        prob <- function(q) prod(dnorm(b,a%*%q,S^-.5))
    prob <- function(q,del15N1,A,B,sdB) exp(-.5*S*SSR(q,del15N1,A,B,sdB))
    ## test <- function(q2) (prob(q2)/prob(q1))>runif(1) #metropolis criterion
    test <- function(q1,q2,del15N1,del15N2,A,B,sdB) exp(-.5*S*(SSR(q2,del15N2,A,B,sdB)-SSR(q1,del15N1,A,B,sdB))) > runif(1)
    
  } else {
    prob <- function(q) 1
    test <- function(q2) TRUE
    S <- 1
    overdetermined <- FALSE
  }
  
  outputlength <- min (outputlength,iter)
  ou <- ceiling(iter/outputlength)
  
  q1 <- rep(0,k)
  x <- matrix(nrow=outputlength,ncol=lx,dimnames=list(NULL,colnames(A)))
  x[1,] <- x0
  naccepted <- 1
  p <- vector(length=outputlength) # probability distribution
  p[1] <- prob(q1,del15N1,A,B,sdB)
  del15Ntrack <- matrix(nrow=outputlength,ncol=46)
  del15Ntrack[1,] <- del15N1
  
  if (fulloutput)   {
    q <- matrix(nrow=outputlength, ncol=k)
    q[1,] <- q1
  }
  
  if (is.null(jmp)) jmp <- automatedjump(a,b,g,h) # automatedjump(g,h)
  if (type=="mirror") newq <- mirror
  if (type=="rda") newq <- rda
  if (type=="cda") newq <- cda
  
  ## ##################
  ## the random walk ##
  ## ##################
  
  if (!is.null(burninlength))
    
    for (i in 1:burninlength)  {
      q2 <- newq(q1,g,h,k,jmp)
      del15N2 <- new15N(del15N1,jmp2)
      if (test(q1,q2,del15N1,del15N2,A,B,sdB)){
        q1 <- q2
        del15N1 <- del15N2
      } 
    }
  
  for (i in 2:outputlength)  {
    for (ii in 1:ou)  {
      if (estimate_sdB) ## sd is estimated using Laine 2008
        S <- rgamma(1,shape=lb,rate=0.5*(SS0+SSR(q1)))
      q2 <- newq(q1,g,h,k,jmp)
      del15N2 <- new15N(del15N1,jmp2)
      test(q1,q2,del15N1,del15N2,A,B,sdB)
      if (test(q1,q2,del15N1,del15N2,A,B,sdB)) {
        q1 <- q2
        del15N1 <- del15N2
        naccepted <- naccepted+1
      }
    }
    notify(i, outputlength)     ##added function that adds progress bar functionality (from Tom)
    x[i,] <- x0+Z%*%q1
    p[i] <- prob(q1,del15N1,A,B,sdB)
    del15Ntrack[i,] <- del15N1
    if (fulloutput)  q[i,] <- q1
  }
  ## ##################
  ## end random walk ##
  ## ##################
  
  xnames <- colnames(A)
  if (is.null(xnames)) xnames <- colnames(E)
  if (is.null(xnames)) xnames <- colnames(G)
  colnames (x) <- xnames
  
  xsampleN15 <- list(X=x,acceptedratio=naccepted/iter,p=p,jmp=jmp,del15Ntrack=del15Ntrack)
  if (fulloutput) xsampleN15 <- list(X=x,acceptedratio=naccepted/iter,
                                   Q=q,p=p,jmp=jmp,del15Ntrack=del15Ntrack)
  
  return(xsampleN15)
}
```
